# Quantification of heterogeneity in human CD8^+^ T cell responses to vaccine antigens: an HLA-guided perspective

**DOI:** 10.1101/2024.07.02.601716

**Authors:** Duane C. Harris, Apoorv Shanker, Makaela M. Montoya, Trent R. Llewellyn, Anna R. Matuszak, Aditi Lohar, Jessica Z. Kubicek-Sutherland, Ying Wai Li, Kristen Wilding, Ben Mcmahon, Sandrasegaram Gnanakaran, Ruy M. Ribeiro, Alan S. Perelson, Carmen Molina-París

## Abstract

Vaccines have historically played a pivotal role in controlling epidemics. Effective vaccines for viruses causing significant human disease, *e.g*., Ebola, Lassa fever, or Crimean Congo hemorrhagic fever virus, would be invaluable to public health strategies and counter-measure development missions. Here, we propose coverage metrics to quantify vaccine-induced CD8^+^T cell-mediated immune protection, as well as metrics to characterize immuno-dominant epitopes, in light of human genetic heterogeneity and viral evolution. Proof-of-principle of our approach and methods will be demonstrated for Ebola virus, SARS-CoV-2, and *Burkholderia pseudomallei* (vaccine) proteins.

## 1 INTRODUCTION

Vaccines exploit the exceptional ability of the adaptive immune system to respond to, and remember, encounters with pathogens [1]. Novel vaccine technologies (*e.g*., viral vector, DNA, or RNA) enable a “plug and play” approach to *immunogen* (part of the pathogen that can be recognized by the immune system) design [2]. These technical advances inherently raise a number of challenges in vaccine immunology. First, the genetic diversity of highly variable pathogens makes it difficult to identify an immunogen that can be used in a vaccine to protect against infection. Second, in addition to targeting the genetic diversity of the pathogen, the most effective route to vaccine efficacy and protection is to engage multiple arms of the immune system [1]. Thus, a first challenge is: given a pathogen, how to optimize the choice of immunogens.

A second challenge relates to the (molecular or cellular) mechanisms that mediate immune protection after vaccination or infection. Finding an immune response that correlates with protection can accelerate the development of new vaccines [3]. Unfortunately, there exist significant gaps in our immunological knowledge of *correlates of (vaccine-mediated) protection*. Most current vaccine strategies aim to confer protection through antibodies (humoral response), which are produced by B cells. Yet, there exists substantial evidence of protective *cellular immunity* correlated with CD8^+^ T cell-mediated responses to *conserved regions* of the genome of HIV-1 [4], Lassa virus [5], SARS-CoV-2 [6, 7], pandemic influenza [8], and Ebola virus [9]. Hence, a third challenge is to quantify the potential of CD8^+^ T cells to induce vaccine-mediated immune responses, and if possible, to identify viral immuno-dominant epitopes in these responses. CD8^+^ T cells (or cytotoxic T cells that kill infected cells) express a unique receptor on their surface: the T cell receptor (TCR). The binding of TCRs to immunogens on the surface of infected cells initiates an immune response (see Fig. 1). In the case of CD8^+^ T cells, the immunogen is a bi-molecular complex composed of a viral *peptide* (a short protein fragment) bound to a major histocompatibility complex (MHC) class I molecule, referred to as a pMHC complex. In humans, the MHC molecule is also called human leukocyte antigen (HLA) [10, 11]. This constitutes the *MHC-restriction* of TCR immunogen recognition. MHC-restriction brings additional challenges to the study of CD8^+^ T cell responses, since the HLA locus is the most polymorphic gene cluster of the entire human genome [10], and genome-wide association studies of host and virus genomes have shown that different HLA alleles exert selective pressure, driving *in vivo* viral evolution (*e.g*., hepatitis C virus [11, 12] and HIV-1 [13]). Our objective in this manuscript is to define novel metrics to quantify CD8^+^ T cell-mediated vaccine protein coverage, in light of human HLA heterogeneity, viral evolution, and immuno-dominant epitopes.

**Figure 1.**
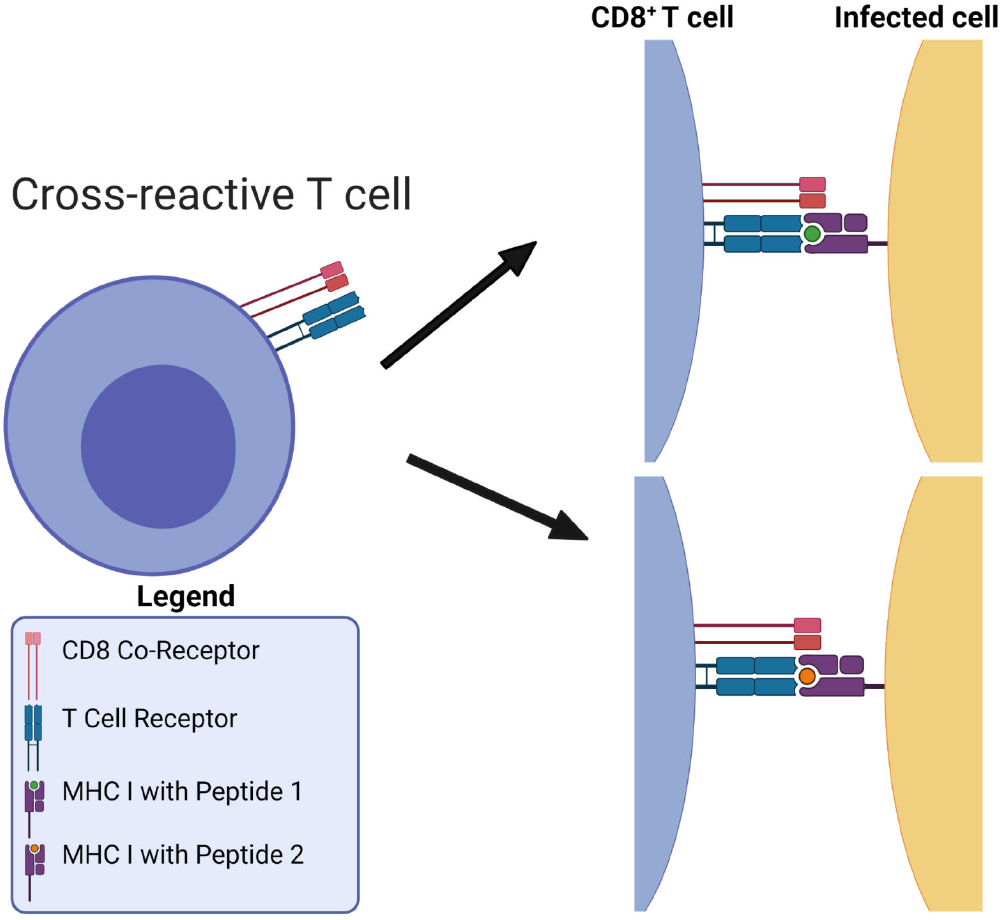
MHC-restriction in T cell receptor recognition of peptide-MHC complexes. T cell receptors are cross-reactive: they can bind to many different viral pMHCs. Figure reproduced from Ref. [14] (Figure 1) with permission under the terms and conditions of the Creative Commons Attribution license CC BY 4.0.

Desirable in a vaccine-induced CD8^+^ T cell immune response is for it to be broad and directed against several immunogens, ideally from conserved genome regions, to reduce the possibility of selecting viral escape variants, and to make it more difficult for the virus to exhaust that response. We hypothesize that the problem to *i)* optimize CD8^+^ T cell-mediated vaccine coverage across the human population, while *ii)* minimizing viral escape is best, and naturally, posed in terms of a multi-partite graph, given the HLA genetic heterogeneity, the bi-molecular nature of T cell immunogens, and that immunogen recognition by TCRs is inherently cross-reactive (see Fig. 1). Thus, we propose to represent CD8^+^ T cell viral immunogen recognition as a multi-partite graph, 𝒢, with four different sets of nodes (see Fig. 2). The first set, ℛ, corresponds to eleven geographical regions covering the world’s human population [15], so that ℛ = {*r*_1_, *r*_2_, …, *r*_*K*_ } (*K* = 11); the second set, 𝒜, to *M* different HLA alleles in the human population (of a given region), so that 𝒜 = {*a*_1_, *a*_2_, …, *a*_*M*_}; the third set, 𝒫, to *N* different peptides (9 amino acids long derived from the vaccine protein of interest), so that 𝒫 = {*p*_1_, *p*_2_, …, *p*_*N*_ }; and the fourth set, 𝒯, to *D* different possible TCR molecular structures, so that 𝒯 = {*t*_1_, *t*_2_, …, *t*_*D*_}. Edges between nodes (from different sets) are as follows: *i)* an edge between a geographical region and an HLA allele encodes the frequency of that allele in the region (see Section 2.1.1), *i.e*., 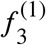 is the frequency in *r*_1_ of allele *a*_3_; *ii)* an edge between an HLA allele and a peptide encodes the binding score of the HLA allele to the peptide and thus, represents the stability of this interaction (see Section 2.1.2), *i.e*., *s*_51_ is the binding score of allele *a*_5_ to peptide *p*_1_; and *iii)* an edge between a peptide and a TCR encodes the binding score of the peptide to the TCR and thus, represents the immunogenicity of the peptide (see Section 2.1.3), *i.e*., *g*_41_ is the immunogenicity of peptide *p*_4_ as measured by TCR *t*_1_ (see Fig. 2). This novel graph approach allows us to address the above challenges: *1)* viral genetic diversity of the pathogen is represented in the set of peptides, 𝒫, so that wild type and all circulating (or predicted) variants can be analyzed, *2)* HLA variability is considered with regard to geographical regions ℛ, HLA alleles 𝒜, and their frequencies within each region, and *3)* TCR recognition variability is accounted for by *peptide immunogenicity* [16]. Finally, the entire multi-partite graph, 𝒢, straightforwardly provides a *metric* to quantify *vaccine coverage* (see Section 2.2), and the framework to characterize *immuno-dominant* peptides (experimentally identified) and to predict *viral immune escape* from CD8^+^ T cell recognition [17] (see Section 4). Our methods will be applied to Ebola virus, SARS-CoV-2, and *Burkholderia pseudomallei* vaccine proteins.

**Figure 2.**
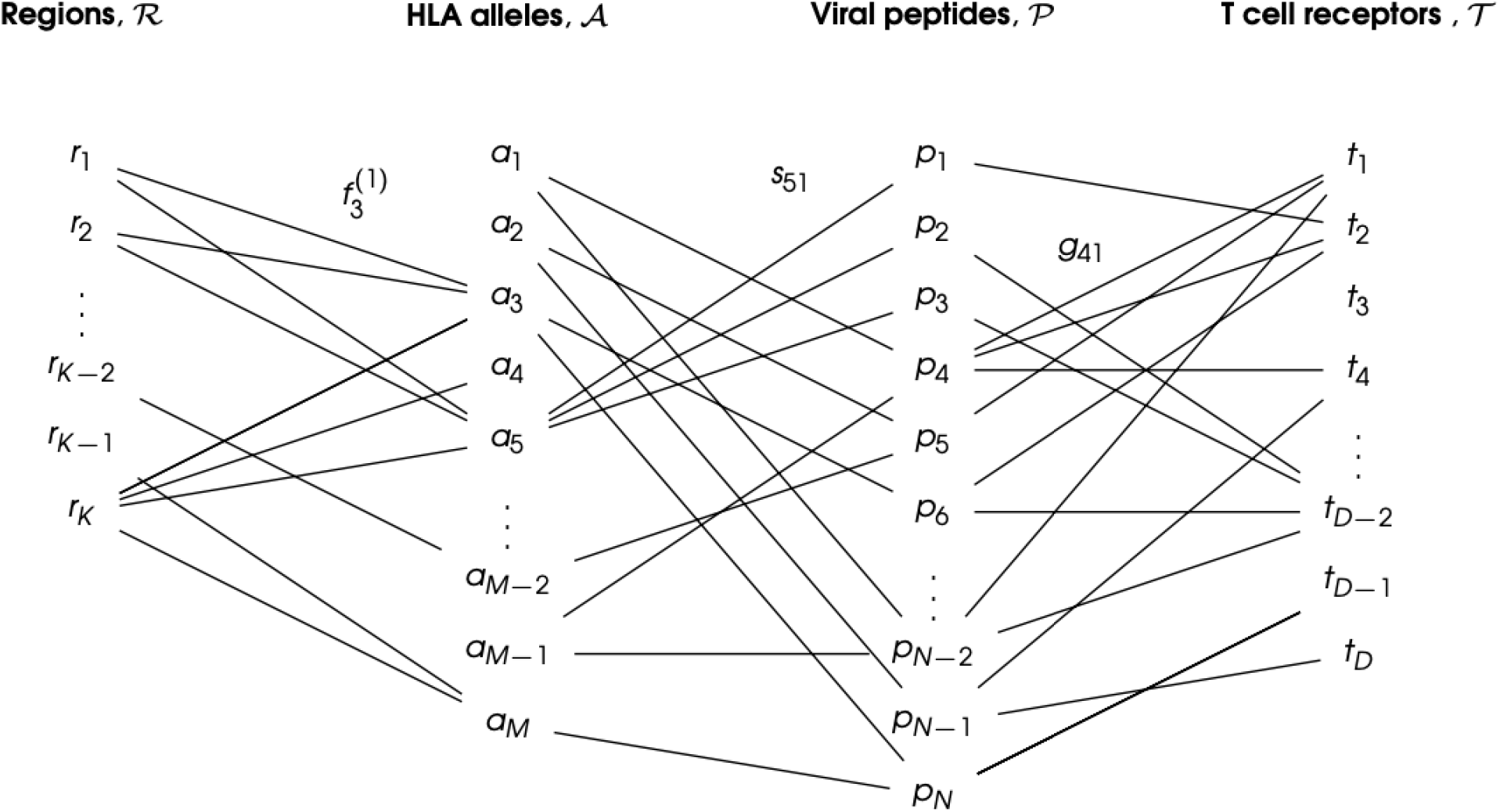
CD8^+^ T cell immunogen recognition as a multi-partite graph, 𝒢, to account for geographical HLA allele variation. Only a subset of the edges is shown for clarity.

A wide range of extremely valuable computational tools have already been developed to accelerate T cell epitope discovery and vaccine design, *e.g*., Predivac-3.0, a proteome-wide bioinformatics tool [18], Epigraph, a graph-based algorithm to optimize potential T cell epitope coverage [19], OptiTope, a web server for the selection of an optimal set of peptides for epitope-based vaccines [20, 21], or PEPVAC, a web server for multi-epitope vaccine development based on the prediction of MHC supertype ligands [22]. Our interest and objective is slightly different; we want to capture the contributions of human HLA class I heterogeneity, petide:TCR interaction, and the more often studied HLA allele:peptide interaction, to CD8^+^ T cell responses to vaccine proteins. We note that immunogenicity of a peptide as defined in Refs. [20, 21, 18] is based on MHC class I binding affinity prediction methods, but not on the contribution of T cell receptor binding as considered in this manuscript [23]. Furthermore, PEPVAC’s predictions of promiscuous epitopes are focused on five HLA I supertypes (HLA-A and HLA-B genes) [22], while we are interested in individual HLA class I allele frequencies in a given human population. Thus, in this paper we present a framework to characterize CD8^+^ T cell immunogen recognition, based on a multi-partite graph representation (see Fig. 2), which can account for geographical variation in HLA class I allele frequencies (for each HLA allele type), HLA allele and peptide interaction, as well as peptide and T cell receptor interaction. The paper is organized as follows. Section 2 describes our methods and approaches; in particular, it presents the details of data acquisition, definition of the coverage metrics, regional and individual, to quantify HLA-driven variability of CD8^+^ T cell responses, as well as metrics to characterize and compare immuno-dominant CD8^+^ T cell epitopes. Results are presented in Section 3, where we focus our attention to the North America region. We have analysed all regions and those results are included as Supplementary Material. We conclude with a discussion and plans for future work.

## 2 MATERIALS AND METHODS

### 2.1 Data acquisition

The generation of the multi-partite graph, 𝒢, requires the following steps. **Step I:** make use of existing databases, such as Allele Frequency Net Database, to obtain HLA class I allele frequencies for the eleven different geographical regions (see section 2.1.1): Australia, Europe, North Africa, North America, North-East Asia, Oceania, South and Central America, South Asia, South-East Asia, Sub-Saharan Africa, and Western Asia. This will determine the elements in sets ℛ and ℛ, as well as the edges between them. **Step II:** choose a vaccine protein and make use of the database, Immune Epitope Database, to obtain binding scores for pairs of HLA class I alleles and 9-mer peptides (or nonamers) (see section 2.1.2). This determines the elements in set 𝒫, as well as the edges between elements of 𝒜 and 𝒫. **Step III:** compute the immunogenicity of elements in the set 𝒫 making use of methods described in Ref. [16] (see section 2.1.3). In this way, we obtain the edges between elements of 𝒫 and a representative element of 𝒯. We now describe in greater detail these steps, in particular how we collect data directly from databases (see sections 2.1.1 and 2.1.2), and how *mean immunogenicity* is computed based on the approach from Ref. [16] (see section 2.1.3).

#### 2.1.1 HLA class I allele frequencies

Every individual has a total of six (classical) HLA class I alleles: two HLA-A, two HLA-B, and two HLA-C alleles [10]. Here, we are interested in defining coverage metrics for each HLA type, *i.e*., A, B, or C, so that they can be compared. Thus, in what follows we consider each allele type (A, B, or C) separately.

Allele frequency data were obtained from the Allele Frequency Net Database [24, 25]. We have restricted our analysis to studies with a gold or silver population standard ^1^, and have considered HLA class I alleles with two sets of digits, *e.g*., HLA-B∗35:05. This nomenclature indicates the HLA molecule of gene B, with the first two numbers representing the serologic assignment, and the last two, the unique sequence [27]. No allele suffix has been included in our results to indicate its expression status [28]. It is out of the scope of this paper to consider differences in expression levels of the different HLA types [29]. The HLA database divides its data into eleven geographical regions, and each of these regions is subdivided into a number of locations ^2^. Independent studies (from peer-reviewed publications, HLA and immuno-genetics workshops, individual laboratories, and short publication reports in collaboration with the *Human Immunology* journal) were conducted to determine allele frequencies at each location. The database contains local (at the location of the study) allele frequencies, calculated using the following equation

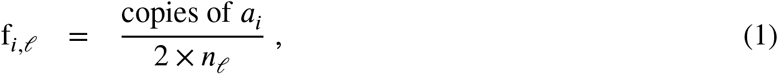

where f_*i,𝓁*_ is the frequency of allele *a*_*i*_ at location *𝓁*, “copies of *a*_*i*_” refers to the total number of copies of allele *a*_*i*_ in the population sample at the given location, and *n*_*𝓁*_ to the sample size of the population in the local study (at location *𝓁*). The factor two is required since humans are diploids, and thus, there are two alleles for each gene [10]. We note that Eq. (1) will be used for each HLA type (A, B, or C). To compute the regional allele frequency based on the frequency data provided for each location, we take the weighted average of the local frequencies; that is, if we denote by ℛ = {*r*_1_, …, *r*_*K*_ }, with *K* = 11, the different regions, the frequency of allele *a*_*i*_ in 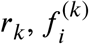, with 1 ≤ *k* ≤ *K*, is given by

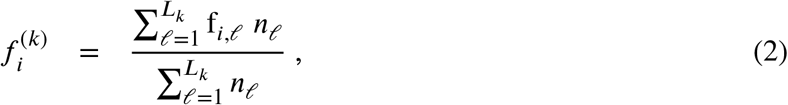

where *L*_*k*_ is the total number of study locations in region *r*_*k*_, f_*i,𝓁*_ the frequency of allele *a*_*i*_ at location *𝓁* (defined in Eq. (1)), and *n*_*𝓁*_ the sample size at location *𝓁*. We note that once the regional frequency of each allele is calculated, the sum (over alleles) of their regional frequencies is close to one, but not necessarily equal to one [26]. Therefore, we define

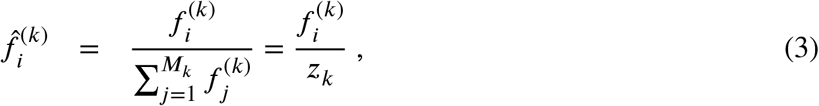

where 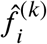 is the normalized frequency of allele *a*_*i*_ in region *r*_*k*_, *M*_*k*_ the number of different unique alleles found in region *r*_k_, and we have introduced the variable 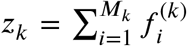. We note that both *M* and *z* depend on the region under consideration, and thus, our choice of notation includes this fact (as a lower index). Table 5 in section 3 provides the values of *M*_*k*_ and *z*_*k*_ for each region and allele type (HLA-A, HLA-B, and HLA-C).

#### 2.1.2 Binding scores of HLA class I alleles to 9-mer peptides

The next step is to choose a protein, under consideration for use in a vaccine, and analyze all its (linear) 9-mer (9 amino acids long) peptides (or nonamers), which can be potential CD8^+^ T cell epitopes. We note that if the protein is *P* amino acids long, there will be a total of *P* − 9 + 1(= *P* − 8) 9-mer peptides. For the protein of interest, we denote the set of such nonamers by 𝒫 = {*p*_1_, …, *p*_*N*_ } with *N* = *P* − 8. HLA class I allele binding scores (for each HLA type) to CD8^+^ T cell epitopes can be generated with the Immune Epitope Database (IEDB) [30]. Let us consider HLA class I allele *a*_*i*_ and epitope *p*_*j*_ (from a vaccine protein). Given *a*_*i*_ and *p*_*j*_, the IEDB database provides a binding score, *s*_*ij*_, for the pair (*a*_*i*_, *p*_*j*_). The predictions are made with the NetMHCpan-4.1 method [31]. Binding scores range from 0 to 1, with higher scores correlating with greater affinity between the HLA class I allele *a*_*i*_ and the peptide *p*_*j*_. Thus, for a given peptide *p*_*j*_, we shall obtain binding scores for HLA class I alleles of type A, B, and C.

#### 2.1.3 Immunogenicity of CD8^**+**^ T cell epitopes

We now discuss the concept of immunogenicity: a variable to quantify the likelihood that a CD8^+^ T cell receptor will recognize a nonamer [16]. This quantity proposed in Ref. [16] is calculated based on the preference that T cell receptors have for certain amino acids (or enrichment score), and the positions of those amino acids within the nonamer peptide chain. Enrichment scores, as provided in Ref. [16], correspond to logarithmic enrichment values per amino acid, which we denote by *q*_*β*_, with 1 ≤ *β* ≤ 20. Since our aim is to define a non-negative vaccine coverage metric, it is useful to convert such amino acid logarithmic enrichment scores into non-negative and normalized enrichment scores, 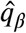, with 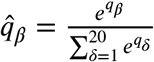. Table 1 provides both the set of values 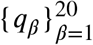 and 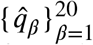. A second contribution to the mean TCR immunogenicity of a 9-mer peptide comes from the specific positions of its amino acids within the nonamer chain. Ref. [16] provides the relative weight (or importance) of position *α* in the nonamer chain, *w*_*α*_, with 1 ≤ *α* ≤ 9. Again, since we are interested in defining a non-negative vaccine coverage metric and the binding scores belong to the interval [0, 1] (see section 2.1.2), it is appropriate to normalize these weights. We, thus, introduce 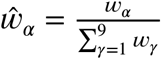. Table 2 provides both the set of values 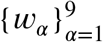 and 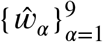. We note that amino acids in positions 1, 2 or 9 do not contribute to the immunogenicity of the nonamer, since these positions are anchor residues, which interact with the MHC molecule. We now can define the immunogenicity of a nonamer.

**Table 1.**
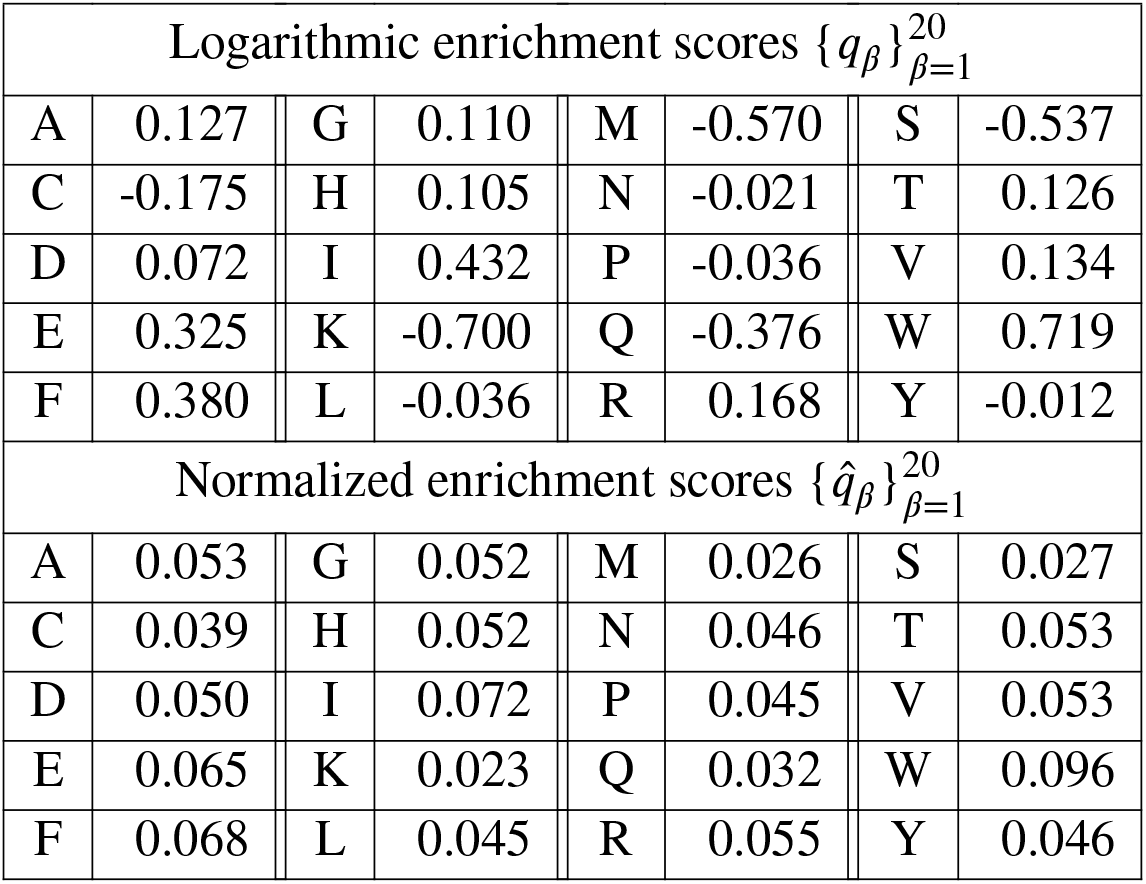
Logarithmic (*q*) and normalized 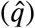 amino acid enrichment scores.

**Table 2.**
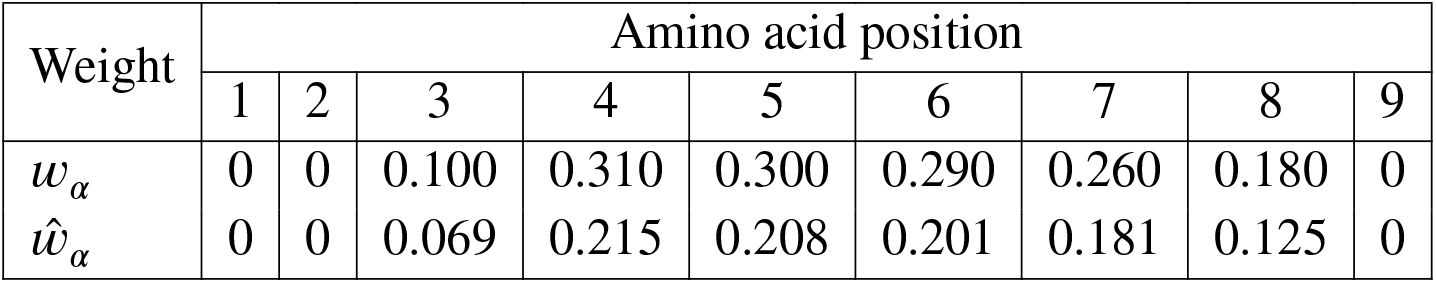
Weights of each position in the nonamer: not normalized (*w*) and normalized 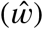.

| Weight | Amino acid position |  |  |  |  |  |  |  |  |
| --- | --- | --- | --- | --- | --- | --- | --- | --- | --- |
|  | 1 | 2 | 3 | 4 | 5 | 6 | 7 | 8 | 9 |
| $w_\alpha$ | 0 | 0 | 0.100 | 0.310 | 0.300 | 0.290 | 0.260 | 0.180 | 0 |
| $\hat{w}_\alpha$ | 0 | 0 | 0.069 | 0.215 | 0.208 | 0.201 | 0.181 | 0.125 | 0 |

The immunogenicity, *g*_*j*_, of nonamer *p*_*j*_, with 1 ≤ *j* ≤ *N*, is given by

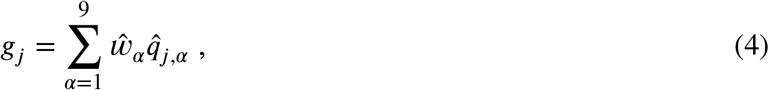

where 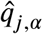 is the normalized enrichment score of the amino acid of peptide *p*_*j*_ in position *α*, with 1 ≤ *α* ≤ 9 and 1 ≤ *j* ≤ *N*, and 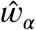 is given in Table 2.

We conclude this section with a few observations. The normalizations proposed ensure that the immunogenicity is positive definite, as is the case for the binding scores presented in the previous section. Its values range from 0.023 (when the epitope consists of lysine only) to 0.096 (when the nonamer consists of tryptophan only). Finally, we note that current estimates of the human TCR diversity in a given individual are of the order of 10^7^ − 10^8^ [32, 33, 34], and thus, we do not have precise knowledge of specific TCR sequences; that is, for a given individual, we cannot enumerate the set 𝒯 = {*t*_1_, *t*_2_, …, *t*_*D*_}. Without this enumeration we are not able to define edges between elements in the sets 𝒫 and 𝒯, and the best we can do is to compute the immunogenicity of an element in 𝒫. It is, then, out of the scope of this paper to consider these edges in the multi-partite graph (see Fig. 2). Our analysis will proceed on the basis of a multi-partite graph with sets ℛ, 𝒜, and 𝒫, with mean immunogenicity of a peptide *p*_*j*_ to a *representative T cell receptor* as a proxy for the edges to elements in the set 𝒯.

### 2.2 Coverage metric to quantify HLA-driven variability of CD8^+^ T cell responses

We now have all the ingredients to define a coverage metric to quantify HLA-driven variability of CD8^+^ T cell responses to a (vaccine) protein. We first introduce a *mean regional coverage metric*, and then we propose, since an individual only expresses two alleles of a given HLA class I, an *individual regional coverage metric* and a corresponding *mean individual regional coverage metric*. We shall show that in the absence of correlations between HLA alleles, or allele associations, the mean regional and the mean individual regional coverage metrics are the same.

#### 2.2.1 Mean regional coverage metric: a definition

We define, for a given (vaccine) protein, its mean regional coverage metric in region *r*_*k*_, 𝒞_*k*_, as follows

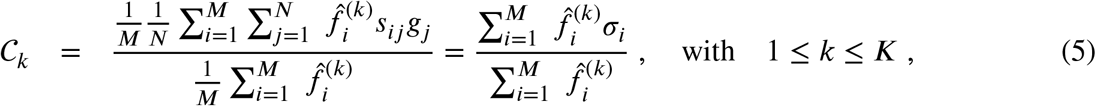

where *M* is the number of alleles considered (*M* = 25 in what follows, and we note that *M* ≠ *M*_*k*_, see section 3), index *i* sums over alleles, 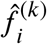 the normalized frequency of allele *a*_*i*_ in region *r*_*k*_ (defined in Eq. (3)), *N* the total number of nonamer (linear) epitopes that can be formed from the (vaccine) protein under consideration, index *j* sums over nonamers, *s*_*ij*_ the binding score of the interaction between allele *a*_*i*_ and nonamer *p*_*j*_ (defined in section 2.1.2), and *g*_*j*_ the immunogenicity of *p*_*j*_ (defined in Eq. (4)). We have introduced σ_*i*_, for 1 ≤ *i* ≤ *M*, defined by

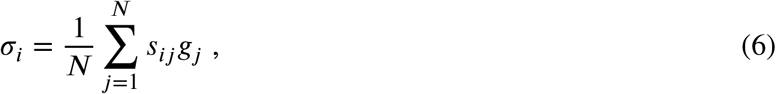

and which measures how well (on average) allele *a*_*i*_ binds to the nonamers from the vaccine protein of interest, with binding score weighted by nonamer immunogenicity to CD8^+^ T cell receptors. Eq. (5) will be used for each HLA class I allele type separately; that is, for a given region and vaccine protein, we shall obtain three different values for HLA-A, HLA-B, and HLA-C alleles. We note that our choice for *M* is discussed in section 3.

#### 2.2.2 Individual regional coverage metric: two definitions

We note that 𝒞_*k*_, as defined by Eq. 5, does not consider the fact that an individual only presents two alleles of each type, and not *M*. In order to account for this fact, we now turn to define an individual regional coverage metric. To this end, each individual in a region will be described by an allele pair (for each type), drawn out the *M* different alleles in the region. For the purposes of this study, we have chosen *M* = 25 for each region and allele type (see section 3). This implies that we confine our analysis to individuals whose alleles are drawn from a list of the top *M* (most frequent) alleles (of each type) in their region. We note that for each allele type (A, B, or C), there are a total of 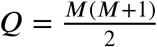 different allele pairs, each of them representing an individual in region *r*_*k*_. We define the *individual regional coverage metric*, 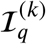, for an individual of region *r*_*k*_, with allele pair *q*, where 1 ≤ *q* ≤ *Q*, as follows

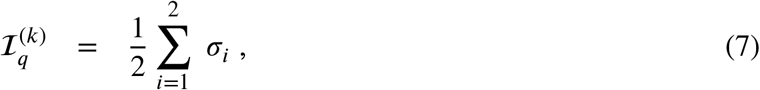

where the sum over *i* corresponds to each of the alleles in the pair *q*, drawn from region *r*_*k*_. Next, making use of the regional frequencies for each allele (see section 2.1.1), we compute the regional frequency of each individual; that is, the regional frequency of each allele pair (for a given type). Let 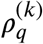 represent the regional frequency (in region *r*_*k*_) of an individual with allele pair *q*. If the individual has two copies of a given allele, *q* = (*a, a*), with 1 ≤ *i* ≤ *M*, then we have 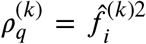. If the two alleles are different, *q* = (*a, a*), with 1 ≤ *i, j* ≤ *M*, and *i* ≠ *j*, then we have 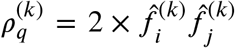, since an individual with allele pair (*a*_*i*_, *a*_*j*_), is equivalent to one with allele pair (*a*_*j*_, *a*_*i*_). We note that this analysis does not account for potential correlations between HLA alleles, or allele associations. With these considerations, we can now define the *mean individual regional coverage metric*, ℐ_*k*_, in region *r*_*k*_ as the weighted average of the coverage metric for each individual in the population; that is, we can write

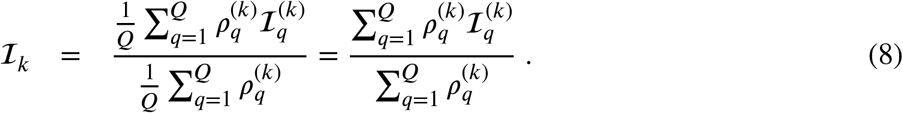

#### 2.2.3 𝒞 _*k*_ and ℐ_*k*_ are equal in the absence of HLA allele associations

We now show that in the absence of HLA allele associations, one has 𝒞 _*k*_ = ℐ _*k*_. We present the proof for the case of a population with three alleles (of a given type). The arguments of the proof can be generalized to any number of alleles. Without lack of generality and to simplify the notation, we drop the regional index and the normalization symbols for the allele frequencies. We denote the three alleles by *a*_1_, *a*_2_, *a*_3_, and their individual frequencies by *f*_1_, *f*_2_, *f*_3_, respectively. Thus, the mean regional coverage metric (see Eq. 5) is given by

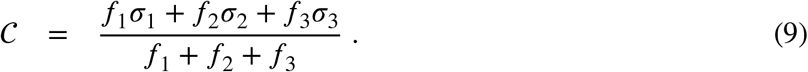

In a population with three alleles, we have 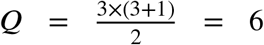 different allele pairs given by: (1, 1), (1, 2), (1, 3), (2, 2), (2, 3), (3, 3). Let us denote by *ρ*_(*n,m*)_ the frequency of allele pair (*n, m*), with *n* ≤ *m* and 1 ≤ *n* ≤ 3. In the absence of HLA allele associations these frequencies are given by

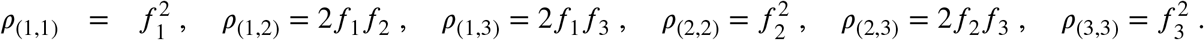

We now make use of Eq. (7) to write

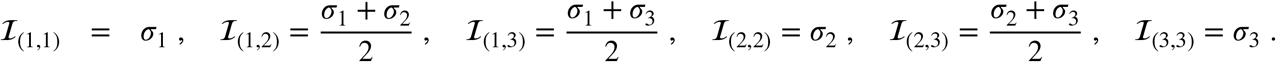

We note that the denominator of Eq. (8) is equal to (*f*_1_ + *f*_2_ + *f*_3_)^2^, so that we can write

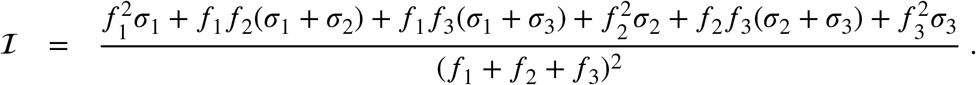

We now collect the factors of σ_1_, σ_2_, σ_3_ as follows

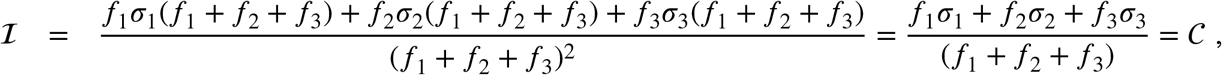

as we wanted to show. The arguments of the proof can also be generalised to *M* alleles, making use of an induction argument.

From now on, we shall compute 𝒞_*k*_ for the different regions, HLA alleles, and vaccine proteins of interest, since it is simpler than ℐ_*k*_, and we have shown that ℐ_*k*_ is equal to 𝒞_*k*_, under the assumption of no HLA allele associations. Were we to be provided with *true* allele pair frequencies, then those could be directly introduced in Eq. (8) to obtain ℐ_*k*_. It is interesting to observe that the difference between 𝒞_*k*_ and ℐ_*k*_ will encode inherent HLA allele associations, and thus, it is a measure of such correlations [11].

### 2.3 Metrics to characterise and compare immuno-dominant CD8^+^ T cell epitopes

In the previous section we have defined two coverage metrics (mean regional and mean individual regional) to quantify CD8^+^ T cell responses to (vaccine) proteins and their linear 9-mer peptides, as well as their HLA class I heterogeneity based on regional allele frequency differences. As described and reviewed in Ref. [10], not only is the quality of a CD8^+^ T cell response a strong correlate of immune protection, but the relative contribution from the different potential 9-mer peptides (derived from a single protein) can be important to identify immune protection. In fact, it is well known that CD8^+^ T cell responses are generally characterized by an *immuno-dominance hierarchy* of the different nonamers [10], which leads to CD8^+^ T cell responses focused on a small subset of epitopes. A wide range of factors regulate these hierarchies for a given (vaccine) protein: from antigen processing and presentation, to the affinity of the nonamer for MHC class I molecules and the stability of these pMHC complexes, the expression levels of MHC molecules, the affinity of the pMHC complex for TCR molecules and the stability of these complexes, and to CD8^+^ T cell competition [10, 11, 29]. It is clearly out of the scope of this manuscript to consider all of these factors. Our aim here is to investigate *i)* the contribution of known *immuno-dominant* epitopes to the coverage metrics defined earlier, and *ii)* where the known immuno-dominant epitopes fall in suitably defined distributions. In what follows we restrict our study to the SARS-CoV-2 spike protein and Ebola glycoprotein (GP) immuno-dominant nonamers found in Refs. [35, 36], respectively. SARS-CoV-2 spike protein immuno-dominant nonamers (obtained from Table 1 of Ref. [35]) are presented in Table 3 and those for Ebola GP protein (obtained from Table 1 of Ref. [36]) in Table 4.

**Table 3.**
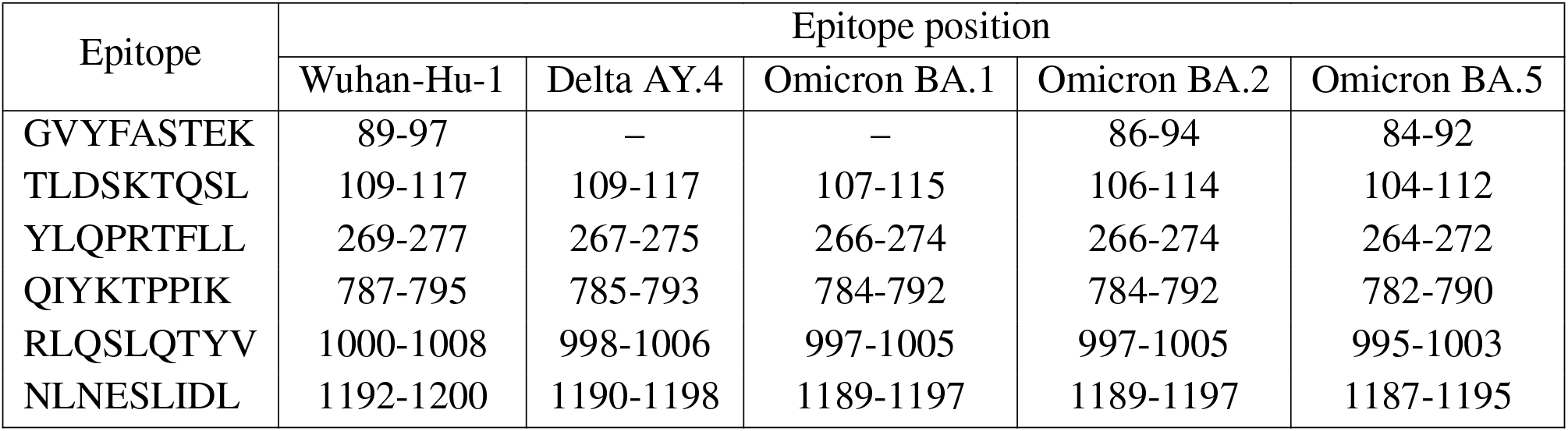
SARS-CoV-2 spike protein immuno-dominant epitopes from Table 1 of Ref. [35], and their presence (or absence) in five different SARS-CoV-2 strains.

**Table 4.** Ebola GP protein immuno-dominant epitopes from Table 1 of Ref. [36], and their presence (or absence) in two different Ebola strains (Sudan and Zaire).

| Epitope | Epitope position |  | Epitope | Epitope position |  |
| --- | --- | --- | --- | --- | --- |
|  | Sudan | Zaire |  | Sudan | Zaire |
| ATDVPSATK | – | 76-84 | DTTIGEWAF | – | 282-290 |
| TDVPSATKR | – | 77-85 | TTIGEWAFW | – | 283-291 |
| GFRSGVPPK | 87-95 | 87-95 | NQDGLICGL | – | 550-558 |
| AENCYNLEI | 105-113 | 105-113 | TELRTFSIL | – | 577-585 |
| RLASTVIYR | 164-172 | 164-172 | ALFCICKFV | – | 667-675 |
| TEDPSSGYY | – | 206-214 | LFCICKFVF | – | 668-676 |

We notice that different viral strains have a different number, *η*, of immuno-dominant epitopes. We have *η* = 6, 5, 5, 6, 6, 12, 3 for SARS-CoV-2 Wuhan-Hu-1, SARS-CoV-2 Delta AY.4, SARS-CoV-2 Omicron BA.1, SARS-CoV-2 Omicron BA.2, SARS-CoV-2 Omicron BA.5 spike, Ebola (Zaire) GP, and Ebola (Sudan) GP, respectively. We first evaluate the contribution of known *immuno-dominant* epitopes to the coverage metrics defined earlier, by defining (for a given protein) the immuno-dominant mean regional coverage metric, 𝒞_*k,D*_, as follows

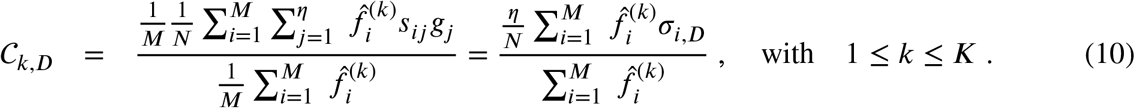

We are, in fact, interested in the ratio

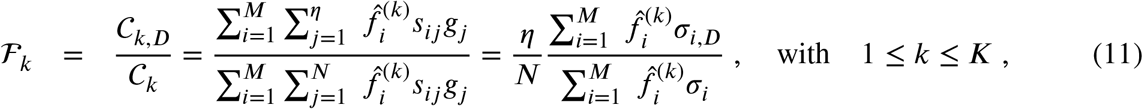

where we have introduced the notation 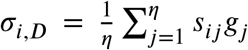, which is the contribution to σ_*i*_ from the immuno-dominant epitopes.

The previous approach can be (easily) extended to the individual regional coverage metric, to evaluate the contribution to this variable from the subset of immuno-dominant epitopes. Let us define for an allele pair *q* (see notation in section 2.2.2), 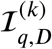, as follows

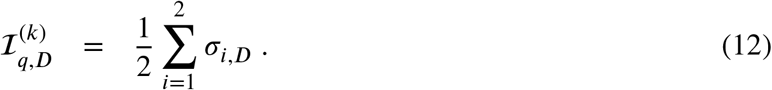

We now introduce the immuno-dominant mean individual regional coverage metric, ℐ _*k,D*_, given by

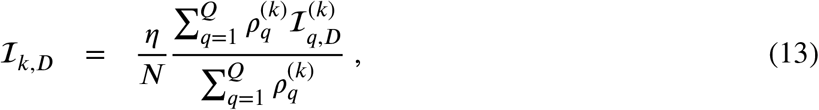

and the ratio ℋ _*k*_, with 1 ≤ *k* ≤ *K*, defined as

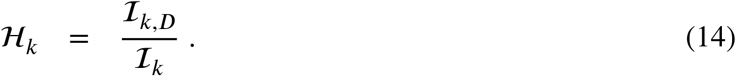

We note that ℐ_*k,D*_ = 𝒞_*k,D*_, and ℋ_*k*_ = ℱ_*k*_, since we have assumed no HLA allele associations. Yet, we point out that if frequencies of allele pairs were available, it would be valuable to compute ℐ_*k,D*_ and ℋ_*k*_ to characterize and quantify the role of HLA allele correlations in the contribution of the immuno-dominant CD8^+^ T cell epitopes to the mean individual regional coverage. The contribution of immuno-dominant nonamers to the mean regional coverage metric is presented in section 3.4.

We now turn to show that the known immuno-dominant epitopes (for the vaccine proteins considered in this section) belong to the tail of suitably defined distributions (these results are provided in section 3). We, thus, define for any *p*_*j*_ ∈ 𝒫, the following variables (averaging over the top *M* alleles in a given region) ^3^:

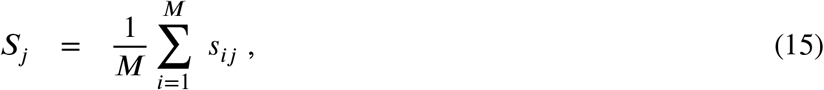

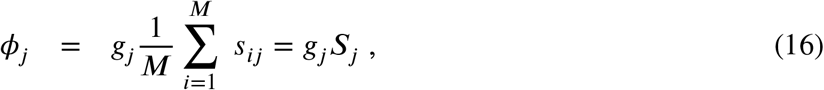

and *g*_*j*_ given by Eq. (4), with 1 ≤ *j* ≤ *N*. We note that *g*_*j*_ only depends on the vaccine protein of interest and is independent of the geographical region considered. On the other hand, *S*_*j*_ and *ϕ*_*j*_ depend on the geographical region considered, since the sum over alleles is different for each region, and on HLA class I allele type. Thus, for a given vaccine protein, we have generated the probability distributions for the variables 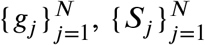, and 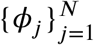, and evaluated where in these distributions the corresponding immuno-dominant epitopes fall (see section 3.5).

## 3 RESULTS

As a demonstration of the methods introduced and discussed in Section 2, we apply them to exemplar pathogens and corresponding proteins. We chose one bacterium (*Burkholderia pseudomallei*) and two viruses (a widespread virus, SARS-CoV-2, and a geographically restricted one, Ebola) to explore different and interesting cases. Specifically, we shall analyze the following proteins: *i) Burkholderia pseudomallei* Hcp1 (A5PM44), *ii)* Ebola (Zaire) GP (Q05320), *iii)* Ebola (Sudan) GP (Q7T9D9), *iv)* Ebola (Zaire) NP (P18272), *v)* Ebola (Sudan) NP (A0A6M2Y086), *vi)* SARS-CoV-2 Wuhan-Hu-1 spike (EPI_ISL_402124), *vii)* SARS-CoV-2 Delta AY.4 spike (EPI_ISL_1758376), *viii)* SARS-CoV-2 Omicron BA.1 spike (EPI_ISL_6795848), *ix)* SARS-CoV-2 Omicron BA.2 spike (EPI_ISL_8135710), and *x)* SARS-CoV-2 Omicron BA.5 spike (EPI_ISL_411542604). In brackets we have provided UniProt accession numbers for the first five proteins, and GISAID accession numbers for the last five. The values of *P* (see Section 2.1.2) are given by *P* = 169, 676, 676, 739, 738, 1273, 1271, 1270, 1270, and 1268, respectively. In our HLA analysis, we have chosen *M* to be equal to 25 (the top 25 most frequent alleles per region) for all regions and HLA class I types, except for HLA-C in Australia, where *M* = 22, since that was the total number of alleles available in the database. The values of *M*_*k*_ and *z*_*k*_ are provided in Table 5. The top 25 alleles per region and per HLA class I type are provided in Table 6 for HLA-A, Table 7 for HLA-B, and Table 8 for HLA-C, respectively.

**Table 5.**
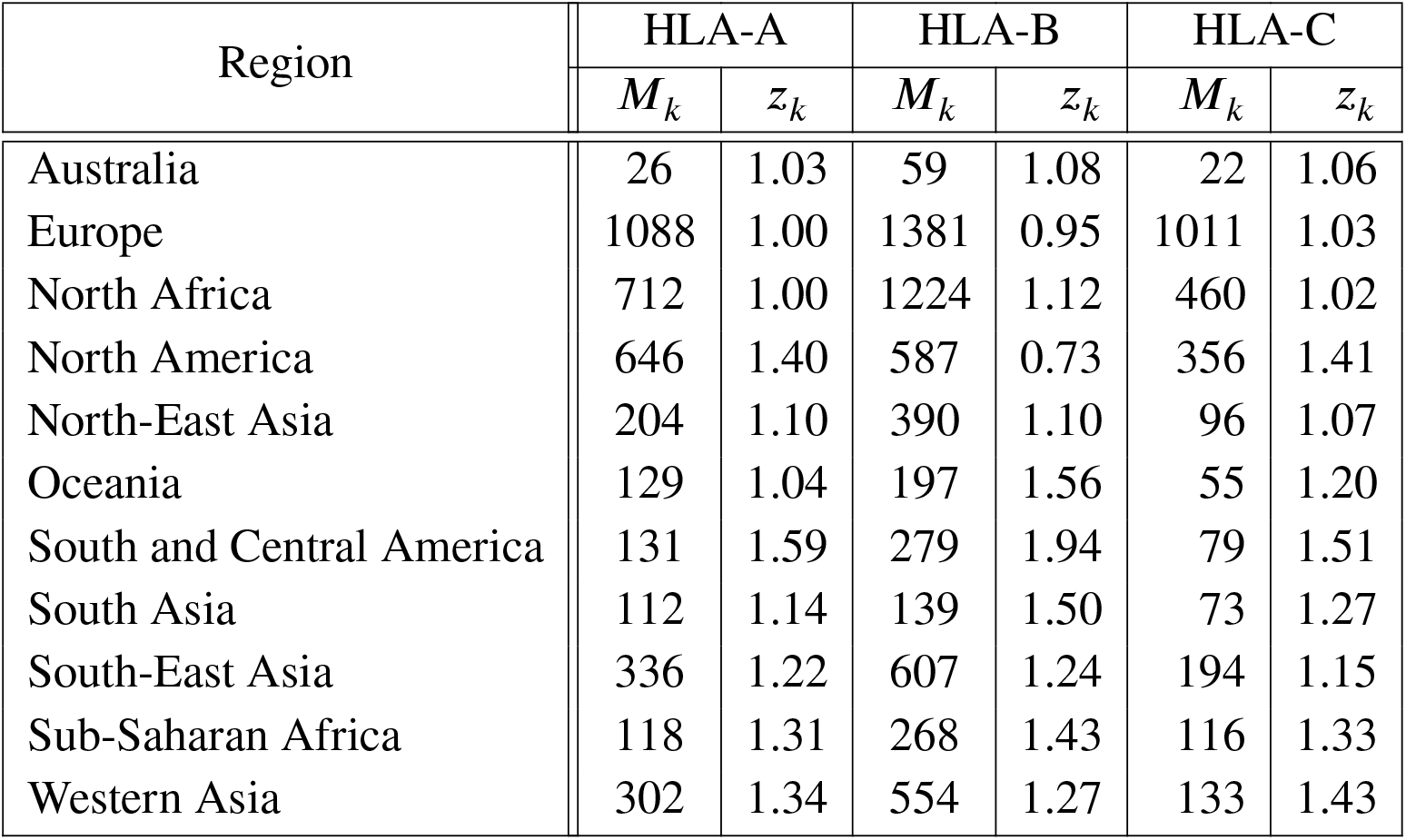
Values of *M*_*k*_ and *z*_*k*_ for every region and HLA class I type. These values were used to compute the normalized regional allele frequencies (see Section 2.1.1).

| Region | HLA-A |  | HLA-B |  | HLA-C |  |
| --- | --- | --- | --- | --- | --- | --- |
| | $M_k$ | $z_k$ | $M_k$ | $z_k$ | $M_k$ | $z_k$ |
| Australia | 26 | 1.03 | 59 | 1.08 | 22 | 1.06 |
| Europe | 1088 | 1.00 | 1381 | 0.95 | 1011 | 1.03 |
| North Africa | 712 | 1.00 | 1224 | 1.12 | 460 | 1.02 |
| North America | 646 | 1.40 | 587 | 0.73 | 356 | 1.41 |
| North-East Asia | 204 | 1.10 | 390 | 1.10 | 96 | 1.07 |
| Oceania | 129 | 1.04 | 197 | 1.56 | 55 | 1.20 |
| South and Central America | 131 | 1.59 | 279 | 1.94 | 79 | 1.51 |
| South Asia | 112 | 1.14 | 139 | 1.50 | 73 | 1.27 |
| South-East Asia | 336 | 1.22 | 607 | 1.24 | 194 | 1.15 |
| Sub-Saharan Africa | 118 | 1.31 | 268 | 1.43 | 116 | 1.33 |
| Western Asia | 302 | 1.34 | 554 | 1.27 | 133 | 1.43 |

**Table 6.** Top 25 most frequent HLA-A alleles for the eleven regions considered, in order of decreasing frequency.

| Australia | Europe | North Africa | North America | North-East Asia | Oceania |
| --- | --- | --- | --- | --- | --- |
| HLA-A*34:01 | HLA-A*02:01 | HLA-A*02:01 | HLA-A*02:01 | HLA-A*24:02 | HLA-A*24:02 |
| HLA-A*24:02 | HLA-A*01:01 | HLA-A*23:01 | HLA-A*01:01 | HLA-A*02:01 | HLA-A*11:01 |
| HLA-A*02:01 | HLA-A*03:01 | HLA-A*30:01 | HLA-A*24:02 | HLA-A*33:03 | HLA-A*34:01 |
| HLA-A*11:01 | HLA-A*24:02 | HLA-A*01:01 | HLA-A*03:01 | HLA-A*11:01 | HLA-A*26:03 |
| HLA-A*01:01 | HLA-A*11:01 | HLA-A*03:01 | HLA-A*31:29 | HLA-A*02:06 | HLA-A*02:06 |
| HLA-A*03:01 | HLA-A*32:01 | HLA-A*68:02 | HLA-A*11:01 | HLA-A*31:01 | HLA-A*24:07 |
| HLA-A*32:01 | HLA-A*68:01 | HLA-A*24:02 | HLA-A*03:27 | HLA-A*26:01 | HLA-A*11:02 |
| HLA-A*68:01 | HLA-A*26:01 | HLA-A*30:02 | HLA-A*24:41 | HLA-A*02:07 | HLA-A*02:01 |
| HLA-A*29:02 | HLA-A*25:01 | HLA-A*29:02 | HLA-A*29:25 | HLA-A*25:01 | HLA-A*26:01 |
| HLA-A*24:13 | HLA-A*31:01 | HLA-A*32:01 | HLA-A*29:50 | HLA-A*29:10 | HLA-A*01:01 |
| HLA-A*26:01 | HLA-A*29:02 | HLA-A*33:03 | HLA-A*68:01 | HLA-A*26:03 | HLA-A*02:05 |
| HLA-A*25:01 | HLA-A*23:01 | HLA-A*33:01 | HLA-A*23:01 | HLA-A*26:02 | HLA-A*24:08 |
| HLA-A*23:01 | HLA-A*30:01 | HLA-A*02:05 | HLA-A*33:03 | HLA-A*03:01 | HLA-A*02:12 |
| HLA-A*24:06 | HLA-A*33:01 | HLA-A*30:04 | HLA-A*29:02 | HLA-A*01:01 | HLA-A*02:07 |
| HLA-A*68:02 | HLA-A*02:05 | HLA-A*34:02 | HLA-A*31:01 | HLA-A*30:01 | HLA-A*24:10 |
| HLA-A*30:01 | HLA-A*68:02 | HLA-A*68:01 | HLA-A*26:01 | HLA-A*24:20 | HLA-A*68:01 |
| HLA-A*30:02 | HLA-A*30:02 | HLA-A*02:02 | HLA-A*32:01 | HLA-A*02:46 | HLA-A*33:03 |
| HLA-A*02:07 | HLA-A*66:01 | HLA-A*11:01 | HLA-A*02:240 | HLA-A*01:134 | HLA-A*68:03 |
| HLA-A*02:05 | HLA-A*33:03 | HLA-A*31:01 | HLA-A*30:01 | HLA-A*23:01 | HLA-A*66:01 |
| HLA-A*33:03 | HLA-A*29:01 | HLA-A*26:01 | HLA-A*30:02 | HLA-A*02:10 | HLA-A*24:04 |
| HLA-A*30:04 | HLA-A*03:02 | HLA-A*03:02 | HLA-A*24:143 | HLA-A*02:04 | HLA-A*31:01 |
| HLA-A*29:01 | HLA-A*02:06 | HLA-A*74:01 | HLA-A*68:02 | HLA-A*68:02 | HLA-A*02:119 |
| HLA-A*26:03 | HLA-A*24:03 | HLA-A*66:01 | HLA-A*24:242 | HLA-A*32:01 | HLA-A*03:01 |
| HLA-A*24:10 | HLA-A*30:04 | HLA-A*80:01 | HLA-A*02:06 | HLA-A*30:04 | HLA-A*02:10 |
| HLA-A*02:06 | HLA-A*23:02 | HLA-A*30:10 | HLA-A*25:01 | HLA-A*01:28 | HLA-A*30:02 |

| South and Central America | South-East Asia | South Asia | Sub-Saharan Africa | Western Asia |
| --- | --- | --- | --- | --- |
| HLA-A*24:02 | HLA-A*24:02 | HLA-A*11:01 | HLA-A*02:01 | HLA-A*01:01 |
| HLA-A*02:01 | HLA-A*11:01 | HLA-A*24:02 | HLA-A*23:01 | HLA-A*02:01 |
| HLA-A*02:12 | HLA-A*01:01 | HLA-A*02:01 | HLA-A*68:02 | HLA-A*03:02 |
| HLA-A*31:01 | HLA-A*33:03 | HLA-A*02:07 | HLA-A*30:02 | HLA-A*26:01 |
| HLA-A*68:01 | HLA-A*02:11 | HLA-A*33:03 | HLA-A*30:01 | HLA-A*24:02 |
| HLA-A*03:01 | HLA-A*03:01 | HLA-A*02:03 | HLA-A*01:01 | HLA-A*31:03 |
| HLA-A*01:01 | HLA-A*68:01 | HLA-A*11:02 | HLA-A*29:02 | HLA-A*11:01 |
| HLA-A*02:19 | HLA-A*02:01 | HLA-A*02:06 | HLA-A*74:01 | HLA-A*02:02 |
| HLA-A*11:01 | HLA-A*26:01 | HLA-A*26:01 | HLA-A*03:01 | HLA-A*31:08 |
| HLA-A*23:01 | HLA-A*31:01 | HLA-A*30:01 | HLA-A*02:02 | HLA-A*32:01 |
| HLA-A*29:02 | HLA-A*32:01 | HLA-A*31:01 | HLA-A*23:17 | HLA-A*23:01 |
| HLA-A*02:22 | HLA-A*31:08 | HLA-A*33:19 | HLA-A*66:01 | HLA-A*02:52 |
| HLA-A*68:02 | HLA-A*02:06 | HLA-A*24:94 | HLA-A*02:05 | HLA-A*68:02 |
| HLA-A*68:47 | HLA-A*01:06 | HLA-A*33:01 | HLA-A*34:02 | HLA-A*33:01 |
| HLA-A*02:64 | HLA-A*24:07 | HLA-A*01:01 | HLA-A*33:03 | HLA-A*29:01 |
| HLA-A*68:03 | HLA-A*30:01 | HLA-A*03:01 | HLA-A*36:01 | HLA-A*30:01 |
| HLA-A*68:17 | HLA-A*26:03 | HLA-A*11:12 | HLA-A*68:01 | HLA-A*03:01 |
| HLA-A*30:02 | HLA-A*02:03 | HLA-A*24:07 | HLA-A*24:02 | HLA-A*30:02 |
| HLA-A*33:01 | HLA-A*29:01 | HLA-A*32:01 | HLA-A*32:01 | HLA-A*02:34 |
| HLA-A*30:01 | HLA-A*66:01 | HLA-A*11:10 | HLA-A*11:01 | HLA-A*02:17 |
| HLA-A*26:01 | HLA-A*02:02 | HLA-A*24:20 | HLA-A*29:11 | HLA-A*25:01 |
| HLA-A*33:18 | HLA-A*03:02 | HLA-A*03:08 | HLA-A*24:23 | HLA-A*02:61 |
| HLA-A*32:01 | HLA-A*32:04 | HLA-A*29:01 | HLA-A*30:10 | HLA-A*02:48 |
| HLA-A*02:13 | HLA-A*24:33 | HLA-A*31:18 | HLA-A*26:01 | HLA-A*01:03 |
| HLA-A*24:03 | HLA-A*68:02 | HLA-A*01:26 | HLA-A*32:106 | HLA-A*69:01 |

**Table 7.** Top 25 most frequent HLA-B alleles for the eleven regions considered, in order of decreasing frequency.

| Australia | Europe | North Africa | North America | North-East Asia | Oceania |
| --- | --- | --- | --- | --- | --- |
| HLA-B*13:01 | HLA-B*07:02 | HLA-B*35:01 | HLA-B*07:02 | HLA-B*52:01 | HLA-B*40:02 |
| HLA-B*40:02 | HLA-B*08:01 | HLA-B*50:01 | HLA-B*08:01 | HLA-B*51:01 | HLA-B*35:01 |
| HLA-B*56:01 | HLA-B*44:02 | HLA-B*51:01 | HLA-B*35:01 | HLA-B*15:01 | HLA-B*56:01 |
| HLA-B*40:01 | HLA-B*15:01 | HLA-B*08:01 | HLA-B*15:01 | HLA-B*35:01 | HLA-B*15:06 |
| HLA-B*15:21 | HLA-B*35:01 | HLA-B*53:01 | HLA-B*40:01 | HLA-B*40:02 | HLA-B*40:01 |
| HLA-B*56:02 | HLA-B*51:01 | HLA-B*45:01 | HLA-B*18:01 | HLA-B*44:03 | HLA-B*13:01 |
| HLA-B*08:01 | HLA-B*40:01 | HLA-B*52:01 | HLA-B*13:38 | HLA-B*54:01 | HLA-B*15:02 |
| HLA-B*07:02 | HLA-B*18:01 | HLA-B*15:03 | HLA-B*14:02 | HLA-B*07:02 | HLA-B*59:01 |
| HLA-B*15:25 | HLA-B*44:03 | HLA-B*42:01 | HLA-B*27:05 | HLA-B*40:01 | HLA-B*27:04 |
| HLA-B*44:02 | HLA-B*27:05 | HLA-B*44:02 | HLA-B*40:02 | HLA-B*46:01 | HLA-B*55:02 |
| HLA-B*15:01 | HLA-B*13:02 | HLA-B*07:02 | HLA-B*13:02 | HLA-B*40:06 | HLA-B*39:01 |
| HLA-B*58:01 | HLA-B*35:03 | HLA-B*18:01 | HLA-B*35:61 | HLA-B*39:01 | HLA-B*15:13 |
| HLA-B*39:01 | HLA-B*38:01 | HLA-B*49:01 | HLA-B*35:03 | HLA-B*48:01 | HLA-B*54:01 |
| HLA-B*51:01 | HLA-B*14:02 | HLA-B*58:01 | HLA-B*38:01 | HLA-B*55:02 | HLA-B*56:02 |
| HLA-B*35:01 | HLA-B*40:02 | HLA-B*41:01 | HLA-B*15:03 | HLA-B*59:01 | HLA-B*40:10 |
| HLA-B*27:05 | HLA-B*55:01 | HLA-B*14:02 | HLA-B*07:105 | HLA-B*58:01 | HLA-B*48:01 |
| HLA-B*18:01 | HLA-B*39:01 | HLA-B*41:02 | HLA-B*37:01 | HLA-B*15:18 | HLA-B*48:03 |
| HLA-B*44:03 | HLA-B*37:01 | HLA-B*38:01 | HLA-B*39:01 | HLA-B*13:01 | HLA-B*15:21 |
| HLA-B*38:01 | HLA-B*49:01 | HLA-B*78:01 | HLA-B*40:06 | HLA-B*67:01 | HLA-B*58:01 |
| HLA-B*35:03 | HLA-B*50:01 | HLA-B*13:02 | HLA-B*35:02 | HLA-B*13:02 | HLA-B*35:05 |
| HLA-B*55:01 | HLA-B*52:01 | HLA-B*51:33 | HLA-B*15:231 | HLA-B*15:11 | HLA-B*08:01 |
| HLA-B*14:01 | HLA-B*35:02 | HLA-B*39:10 | HLA-B*14:01 | HLA-B*35:03 | HLA-B*15:31 |
| HLA-B*39:06 | HLA-B*27:02 | HLA-B*44:03 | HLA-B*07:05 | HLA-B*35:02 | HLA-B*15:35 |
| HLA-B*14:02 | HLA-B*14:01 | HLA-B*82:02 | HLA-B*15:02 | HLA-B*44:02 | HLA-B*15:18 |
| HLA-B*57:01 | HLA-B*35:08 | HLA-B*15:10 | HLA-B*39:06 | HLA-B*27:02 | HLA-B*55:04 |

| South and Central America | South-East Asia | South Asia | Sub-Saharan Africa | Western Asia |
| --- | --- | --- | --- | --- |
| HLA-B*35:99 | HLA-B*40:06 | HLA-B*40:01 | HLA-B*53:01 | HLA-B*38:01 |
| HLA-B*40:02 | HLA-B*57:01 | HLA-B*46:01 | HLA-B*58:02 | HLA-B*35:08 |
| HLA-B*35:43 | HLA-B*51:01 | HLA-B*58:01 | HLA-B*15:03 | HLA-B*44:03 |
| HLA-B*35:19 | HLA-B*52:01 | HLA-B*13:01 | HLA-B*58:01 | HLA-B*18:01 |
| HLA-B*35:01 | HLA-B*35:03 | HLA-B*15:02 | HLA-B*45:01 | HLA-B*14:02 |
| HLA-B*48:03 | HLA-B*44:03 | HLA-B*38:02 | HLA-B*42:01 | HLA-B*35:01 |
| HLA-B*51:01 | HLA-B*58:01 | HLA-B*51:01 | HLA-B*07:02 | HLA-B*52:01 |
| HLA-B*44:03 | HLA-B*35:01 | HLA-B*15:01 | HLA-B*35:01 | HLA-B*13:02 |
| HLA-B*35:05 | HLA-B*44:06 | HLA-B*54:01 | HLA-B*15:10 | HLA-B*35:27 |
| HLA-B*07:02 | HLA-B*37:01 | HLA-B*55:02 | HLA-B*44:03 | HLA-B*08:01 |
| HLA-B*44:02 | HLA-B*07:02 | HLA-B*27:04 | HLA-B*08:01 | HLA-B*49:01 |
| HLA-B*39:05 | HLA-B*07:05 | HLA-B*13:02 | HLA-B*18:01 | HLA-B*41:01 |
| HLA-B*14:02 | HLA-B*14:05 | HLA-B*35:01 | HLA-B*49:01 | HLA-B*51:01 |
| HLA-B*18:01 | HLA-B*18:07 | HLA-B*39:01 | HLA-B*44:10 | HLA-B*07:02 |
| HLA-B*35:102 | HLA-B*08:01 | HLA-B*35:89 | HLA-B*57:03 | HLA-B*50:01 |
| HLA-B*35:12 | HLA-B*51:10 | HLA-B*40:02 | HLA-B*81:01 | HLA-B*15:17 |
| HLA-B*08:01 | HLA-B*55:01 | HLA-B*52:12 | HLA-B*51:01 | HLA-B*57:01 |
| HLA-B*35:48 | HLA-B*56:03 | HLA-B*40:06 | HLA-B*14:02 | HLA-B*35:02 |
| HLA-B*39:03 | HLA-B*53:03 | HLA-B*48:01 | HLA-B*41:01 | HLA-B*55:01 |
| HLA-B*40:10 | HLA-B*42:01 | HLA-B*52:01 | HLA-B*40:06 | HLA-B*53:01 |
| HLA-B*40:64 | HLA-B*13:01 | HLA-B*51:02 | HLA-B*52:01 | HLA-B*58:01 |
| HLA-B*39:09 | HLA-B*44:04 | HLA-B*44:03 | HLA-B*13:02 | HLA-B*49:02 |
| HLA-B*15:01 | HLA-B*15:18 | HLA-B*15:11 | HLA-B*47:03 | HLA-B*44:02 |
| HLA-B*49:01 | HLA-B*15:02 | HLA-B*15:32 | HLA-B*13:01 | HLA-B*07:05 |
| HLA-B*08:50 | HLA-B*15:01 | HLA-B*56:01 | HLA-B*27:03 | HLA-B*40:46 |

**Table 8.** Top 25 most frequent HLA-C alleles for the eleven regions considered, in order of decreasing frequency.

| Australia | Europe | North Africa | North America | North-East Asia | Oceania |
| --- | --- | --- | --- | --- | --- |
| HLA-C*04:01 | HLA-C*07:01 | HLA-C*06:02 | HLA-C*01:57 | HLA-C*01:02 | HLA-C*01:02 |
| HLA-C*01:02 | HLA-C*07:02 | HLA-C*04:01 | HLA-C*04:01 | HLA-C*07:02 | HLA-C*04:03 |
| HLA-C*15:02 | HLA-C*04:01 | HLA-C*07:01 | HLA-C*07:02 | HLA-C*03:03 | HLA-C*07:02 |
| HLA-C*04:03 | HLA-C*06:02 | HLA-C*16:01 | HLA-C*07:01 | HLA-C*03:04 | HLA-C*04:01 |
| HLA-C*07:02 | HLA-C*03:04 | HLA-C*12:03 | HLA-C*06:02 | HLA-C*12:02 | HLA-C*03:04 |
| HLA-C*03:03 | HLA-C*05:01 | HLA-C*02:02 | HLA-C*04:43 | HLA-C*08:01 | HLA-C*03:03 |
| HLA-C*07:01 | HLA-C*12:03 | HLA-C*17:01 | HLA-C*03:135 | HLA-C*14:03 | HLA-C*15:02 |
| HLA-C*12:03 | HLA-C*03:03 | HLA-C*08:02 | HLA-C*03:04 | HLA-C*14:02 | HLA-C*08:01 |
| HLA-C*05:01 | HLA-C*02:02 | HLA-C*07:02 | HLA-C*05:01 | HLA-C*04:01 | HLA-C*14:02 |
| HLA-C*06:02 | HLA-C*01:02 | HLA-C*05:01 | HLA-C*01:02 | HLA-C*15:02 | HLA-C*12:02 |
| HLA-C*03:04 | HLA-C*08:02 | HLA-C*15:02 | HLA-C*02:02 | HLA-C*17:03 | HLA-C*03:07 |
| HLA-C*08:02 | HLA-C*15:02 | HLA-C*17:03 | HLA-C*16:01 | HLA-C*06:02 | HLA-C*12:03 |
| HLA-C*07:04 | HLA-C*16:01 | HLA-C*12:02 | HLA-C*03:03 | HLA-C*08:03 | HLA-C*07:04 |
| HLA-C*16:01 | HLA-C*07:04 | HLA-C*03:04 | HLA-C*12:03 | HLA-C*07:01 | HLA-C*05:01 |
| HLA-C*08:01 | HLA-C*14:02 | HLA-C*15:05 | HLA-C*08:02 | HLA-C*07:04 | HLA-C*15:07 |
| HLA-C*02:02 | HLA-C*17:03 | HLA-C*14:02 | HLA-C*15:02 | HLA-C*03:02 | HLA-C*06:02 |
| HLA-C*16:02 | HLA-C*02:09 | HLA-C*16:02 | HLA-C*17:01 | HLA-C*03:05 | HLA-C*14:03 |
| HLA-C*14:02 | HLA-C*17:01 | HLA-C*18:01 | HLA-C*14:02 | HLA-C*12:03 | HLA-C*07:01 |
| HLA-C*03:02 | HLA-C*12:02 | HLA-C*02:10 | HLA-C*08:01 | HLA-C*05:01 | HLA-C*04:07 |
| HLA-C*15:05 | HLA-C*16:02 | HLA-C*18:02 | HLA-C*12:02 | HLA-C*08:22 | HLA-C*01:03 |
| HLA-C*12:02 | HLA-C*03:02 | HLA-C*16:09 | HLA-C*03:02 | HLA-C*02:02 | HLA-C*15:05 |
| HLA-C*17:01 | HLA-C*15:05 | HLA-C*07:04 | HLA-C*07:270 | HLA-C*16:02 | HLA-C*08:02 |
|  | HLA-C*07:18 | HLA-C*04:04 | HLA-C*07:04 | HLA-C*16:01 | HLA-C*15:08 |
|  | HLA-C*16:04 | HLA-C*16:04 | HLA-C*07:248 | HLA-C*16:74 | HLA-C*15:03 |
|  | HLA-C*07:03 | HLA-C*03:03 | HLA-C*15:05 | HLA-C*02:08 | HLA-C*02:02 |

| South and Central America | South-East Asia | South Asia | Sub-Saharan Africa | Western Asia |
| --- | --- | --- | --- | --- |
| HLA-C*04:03 | HLA-C*06:02 | HLA-C*07:02 | HLA-C*06:02 | HLA-C*05:09 |
| HLA-C*04:01 | HLA-C*07:02 | HLA-C*01:02 | HLA-C*04:01 | HLA-C*04:01 |
| HLA-C*07:02 | HLA-C*04:01 | HLA-C*08:01 | HLA-C*07:01 | HLA-C*06:02 |
| HLA-C*01:02 | HLA-C*15:02 | HLA-C*03:04 | HLA-C*17:01 | HLA-C*07:01 |
| HLA-C*07:01 | HLA-C*07:01 | HLA-C*03:02 | HLA-C*16:01 | HLA-C*07:02 |
| HLA-C*03:04 | HLA-C*12:02 | HLA-C*04:01 | HLA-C*02:02 | HLA-C*12:03 |
| HLA-C*03:05 | HLA-C*14:02 | HLA-C*03:03 | HLA-C*03:04 | HLA-C*15:02 |
| HLA-C*06:02 | HLA-C*03:02 | HLA-C*06:02 | HLA-C*02:10 | HLA-C*02:03 |
| HLA-C*05:01 | HLA-C*12:03 | HLA-C*07:17 | HLA-C*07:02 | HLA-C*12:02 |
| HLA-C*16:01 | HLA-C*01:02 | HLA-C*14:02 | HLA-C*08:02 | HLA-C*08:02 |
| HLA-C*08:02 | HLA-C*05:09 | HLA-C*12:02 | HLA-C*07:04 | HLA-C*02:02 |
| HLA-C*15:02 | HLA-C*07:06 | HLA-C*15:02 | HLA-C*18:01 | HLA-C*03:02 |
| HLA-C*12:03 | HLA-C*16:02 | HLA-C*04:03 | HLA-C*03:02 | HLA-C*17:01 |
| HLA-C*02:02 | HLA-C*07:04 | HLA-C*12:03 | HLA-C*07:18 | HLA-C*07:18 |
| HLA-C*02:07 | HLA-C*03:06 | HLA-C*07:01 | HLA-C*18:02 | HLA-C*15:05 |
| HLA-C*03:57 | HLA-C*08:01 | HLA-C*07:04 | HLA-C*07:06 | HLA-C*03:03 |
| HLA-C*03:03 | HLA-C*03:04 | HLA-C*07:03 | HLA-C*12:03 | HLA-C*05:01 |
| HLA-C*02:10 | HLA-C*04:03 | HLA-C*15:05 | HLA-C*07:328 | HLA-C*16:02 |
| HLA-C*01:06 | HLA-C*15:08 | HLA-C*03:16 | HLA-C*05:01 | HLA-C*08:01 |
| HLA-C*07:08 | HLA-C*08:06 | HLA-C*06:06 | HLA-C*04:07 | HLA-C*14:02 |
| HLA-C*15:03 | HLA-C*03:03 | HLA-C*07:06 | HLA-C*15:02 | HLA-C*08:13 |
| HLA-C*17:01 | HLA-C*15:03 | HLA-C*08:03 | HLA-C*03:03 | HLA-C*01:02 |
| HLA-C*08:01 | HLA-C*18:01 | HLA-C*01:03 | HLA-C*14:03 | HLA-C*16:04 |
| HLA-C*07:14 | HLA-C*03:19 | HLA-C*03:09 | HLA-C*08:04 | HLA-C*16:01 |
| HLA-C*03:02 | HLA-C*04:07 | HLA-C*08:22 | HLA-C*15:07 | HLA-C*07:04 |

### 3.1 Mean regional coverage metric

We compute the mean regional coverage metric, 𝒞_*k*_, shown in Fig. 3, grouped by region and for the chosen ten different vaccine proteins. The top panel corresponds to HLA-A, middle one to HLA-B, and bottom to HLA-C alleles, respectively. From left to right, the bars for each region represent Ebola GP (Zaire), Ebola GP (Sudan), Ebola NP (Zaire), Ebola NP (Sudan), SARS-CoV-2 spike (Wuhan-Hu-1), SARS-CoV-2 spike (Delta AY.4), SARS-CoV-2 spike (Omicron BA.1), SARS-CoV-2 spike (Omicron BA.2), SARS-CoV-2 spike (Omicron BA.5), and *Burkholderia* Hcp1. We observe that HLA-C values are (overall) lower than those for HLA-A and HLA-B alleles; this implies that for the studied proteins CD8^+^ T cell responses will be dominated (on average) by T cell receptors binding to HLA-A or HLA-B pMHC complexes. If we now turn our attention to HLA-A alleles (top panel), for almost all regions, the largest values correspond to SARS-CoV-2 spike (Omicron BA.1), SARS-CoV-2 spike (Omicron BA.2), and SARS-CoV-2 spike (Omicron BA.5), followed by SARS-CoV-2 spike (Wuhan-Hu-1) and SARS-CoV-2 spike (Delta AY.4), and then *Burkholderia* Hcp1. Lower values correspond to Ebola GP (Zaire), Ebola GP (Sudan), Ebola NP (Zaire), and Ebola NP (Sudan), with a small overall dominance of Ebola NP (Zaire). Europe does not follow this precise pattern with a large value for *Burkholderia* Hcp1. It is also interesting to note that HLA-A Ebola GP (Zaire) is comparable to, or even larger than, Ebola NP (Zaire) in Australia, North-East Asia, Oceania, South and Central America, South Asia, and South-East Asia. For HLA-B alleles, coverage values are dominated by Ebola NP (Sudan), followed closely by Ebola NP (Zaire), followed by *Burkholderia* Hcp1, then the five different SARS-CoV-2 spike proteins (with similar magnitude), with lowest values for Ebola GP (Sudan) and Ebola GP (Zaire). We note that Ebola NP (nucleoprotein) is not a surface protein, as is the case of GP or SARS-CoV-2 spike. We also note the rather large value of Hcp1 for North America for HLA-B (middle panel).

**Figure 3.**
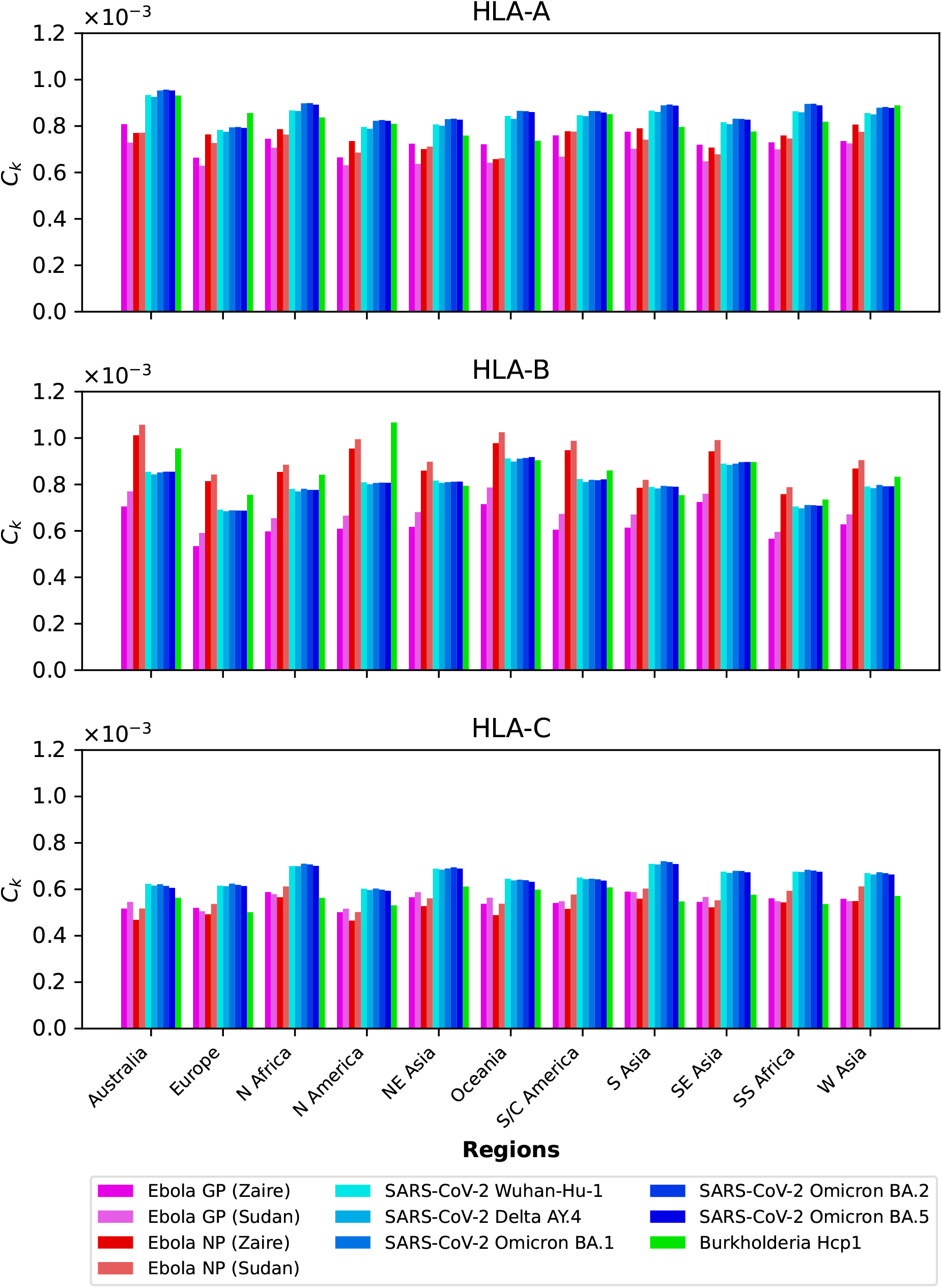
Mean regional coverage metric, 𝒞_*k*_, grouped by region and for ten different proteins. The top panel corresponds to HLA-A, middle one to HLA-B, and bottom to HLA-C alleles, respectively. From left to right, the bars for each region represent Ebola GP (Zaire), Ebola GP (Sudan), Ebola NP (Zaire), Ebola NP (Sudan), SARS-CoV-2 spike (Wuhan-Hu-1), SARS-CoV-2 spike (Delta AY.4), SARS-CoV-2 spike (Omicron BA.1), SARS-CoV-2 spike (Omicron BA.2), SARS-CoV-2 spike (Omicron BA.5), and Burkholderia Hcp1.

We next show in Fig. 4 the mean regional coverage metric, 𝒞_*k*_, grouped by pathogen and for eleven different regions. We observe that for HLA-A and HLA-B alleles, Australia has the largest values, but that is not the case for HLA-C, with North Africa, North-East Asia and South Asia dominating the scores. For HLA-B alleles, Oceania and South-East Asia have overall second largest scores, but for this HLA type the patterns of dominance depend on the specific protein under consideration. For instance, for *Burkholderia* Hcp1 North America clearly dominates, but that is not the case for SARS-CoV-2 spike (overall for the different variants), where Oceania takes the lead. It is interesting to note that for HLA-B the largest values overall are obtained for Ebola NP (Sudan). The results for HLA-C (bottom panel) for a given vaccine protein do not show great variation between geographical regions. North Africa tends to dominate, followed closely by North-East Asia and South Asia. It is interesting to observe that this pattern is broken for Hcp1, where North-East Asia, Oceania, and South and Central America take the lead.

**Figure 4.**
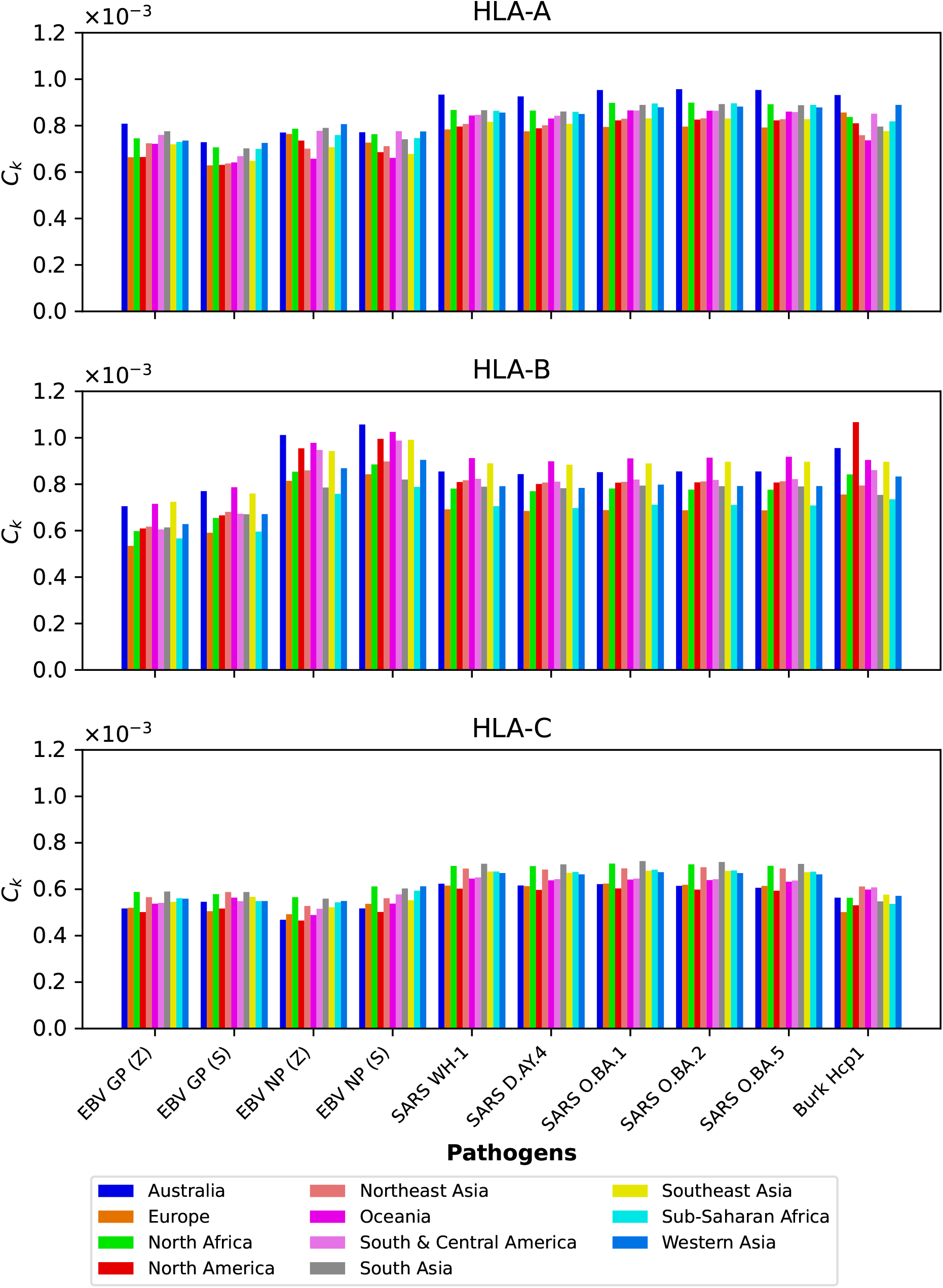
Mean regional coverage metric, 𝒞_*k*_, grouped by pathogen and for eleven different regions. The top panel corresponds to HLA-A, middle one to HLA-B, and bottom to HLA-C alleles, respectively. From left to right, the bars for each protein represent Australia, Europe, North Africa, North America, North-East Asia, Oceania, South and Central America, South Asia, South-East Asia, Sub-Saharan Africa, and Western Asia.

### 3.2 Dissecting the mean regional coverage metric

We now want to dissect the results from the previous section by evaluating the contribution to the mean regional coverage metric from allele frequencies on the one hand, and from HLA allele-peptide binding and peptide immunogenicity, on the other (see Eq. 5). To that end, we focus on North America, and provide plots of the contributions to 𝒞_*k*_ from the normalized allele frequencies and from the binding scores and peptide immunogenicity, as encoded in the variable σ_*i*_ (see Eq. 6). Fig. 5, Fig. 6, and Fig. 7 show on the *x* axis individual alleles (top panel represents HLA-A, middle one HLA-B, and bottom one HLA-C alleles, respectively), on the left *y* axis normalized regional frequencies, and on the right *y* axis the σ_*i*_ value of each allele, for Ebola GP and NP (Sudan and Zaire), SARS-CoV-2 spike (five different variants), and Burkholderia Hcp1 proteins.

**Figure 5.**
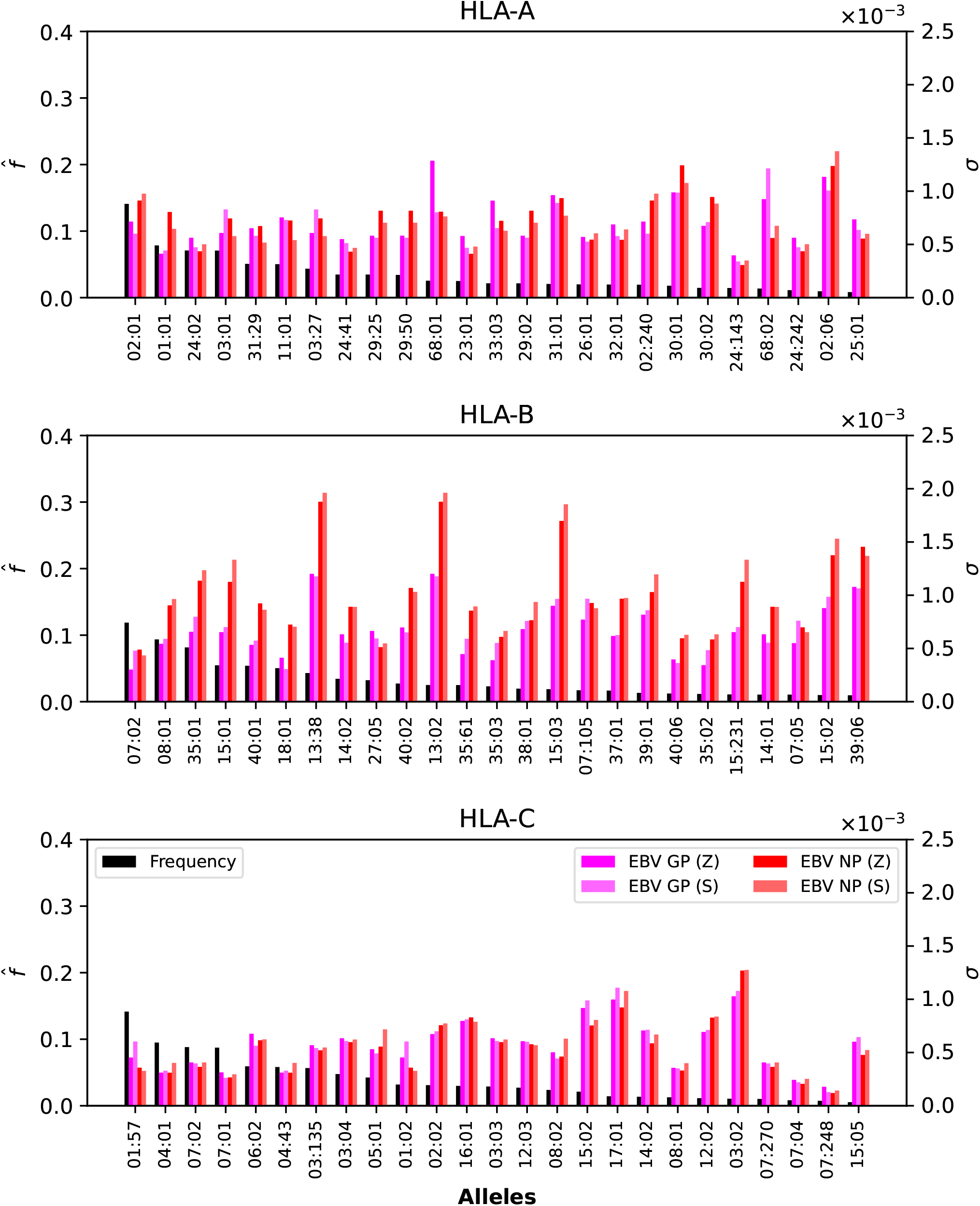
Normalized regional frequencies (left *y* axis), 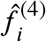, and Ebola σ_*i*_ values (right *y* axis) for the top 25 most frequent alleles of each type in North America (*x* axis). The top panel represents HLA-A, the middle HLA-B, and the bottom HLA-C alleles, respectively.

**Figure 6.**
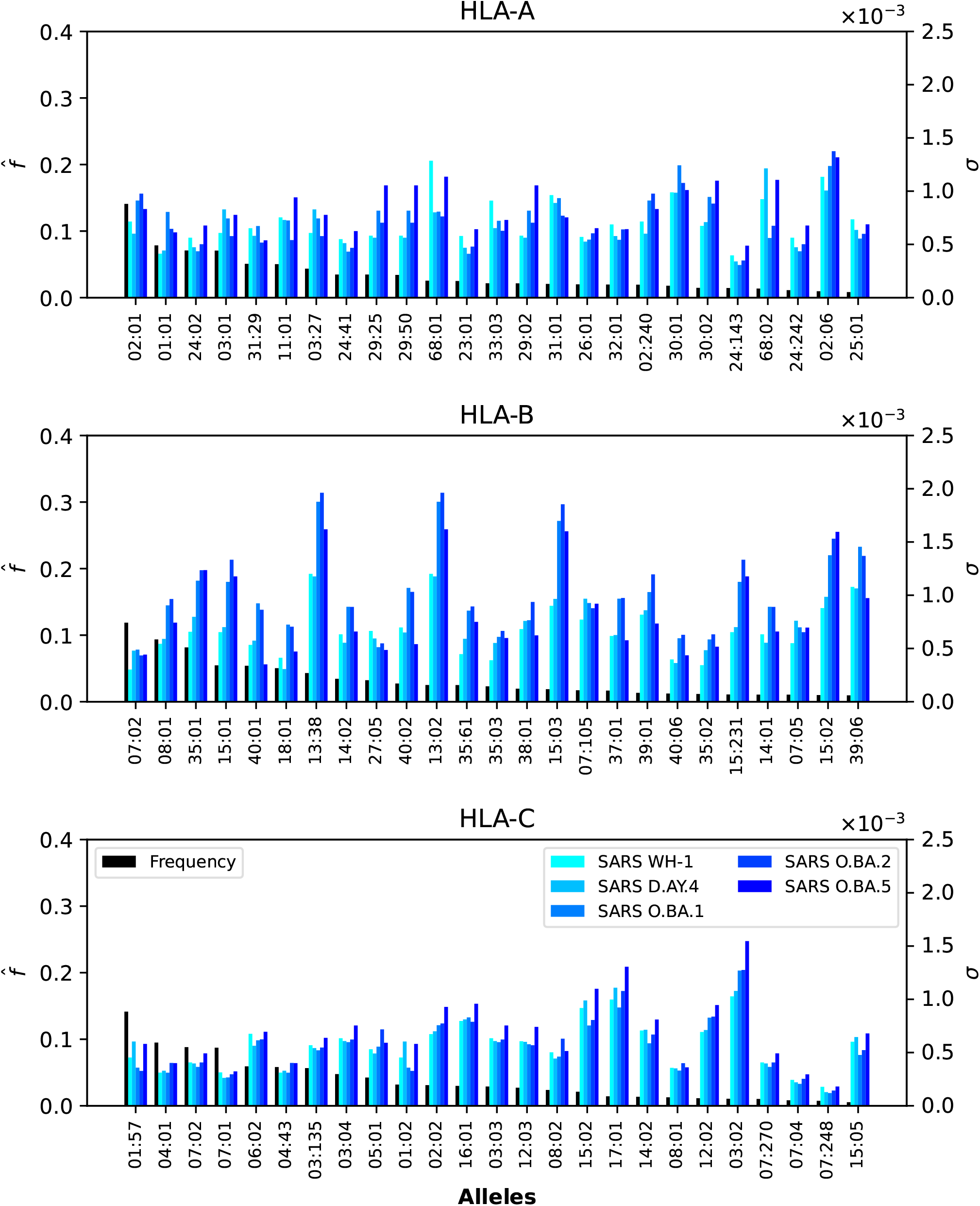
Normalized regional frequencies (left *y* axis), 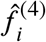, and SARS-CoV-2 σ_*i*_ values (right *y* axis) for the top 25 most frequent alleles of each type in North America (*x* axis). The top panel represents HLA-A, the middle HLA-B, and the bottom HLA-C alleles, respectively.

**Figure 7.**
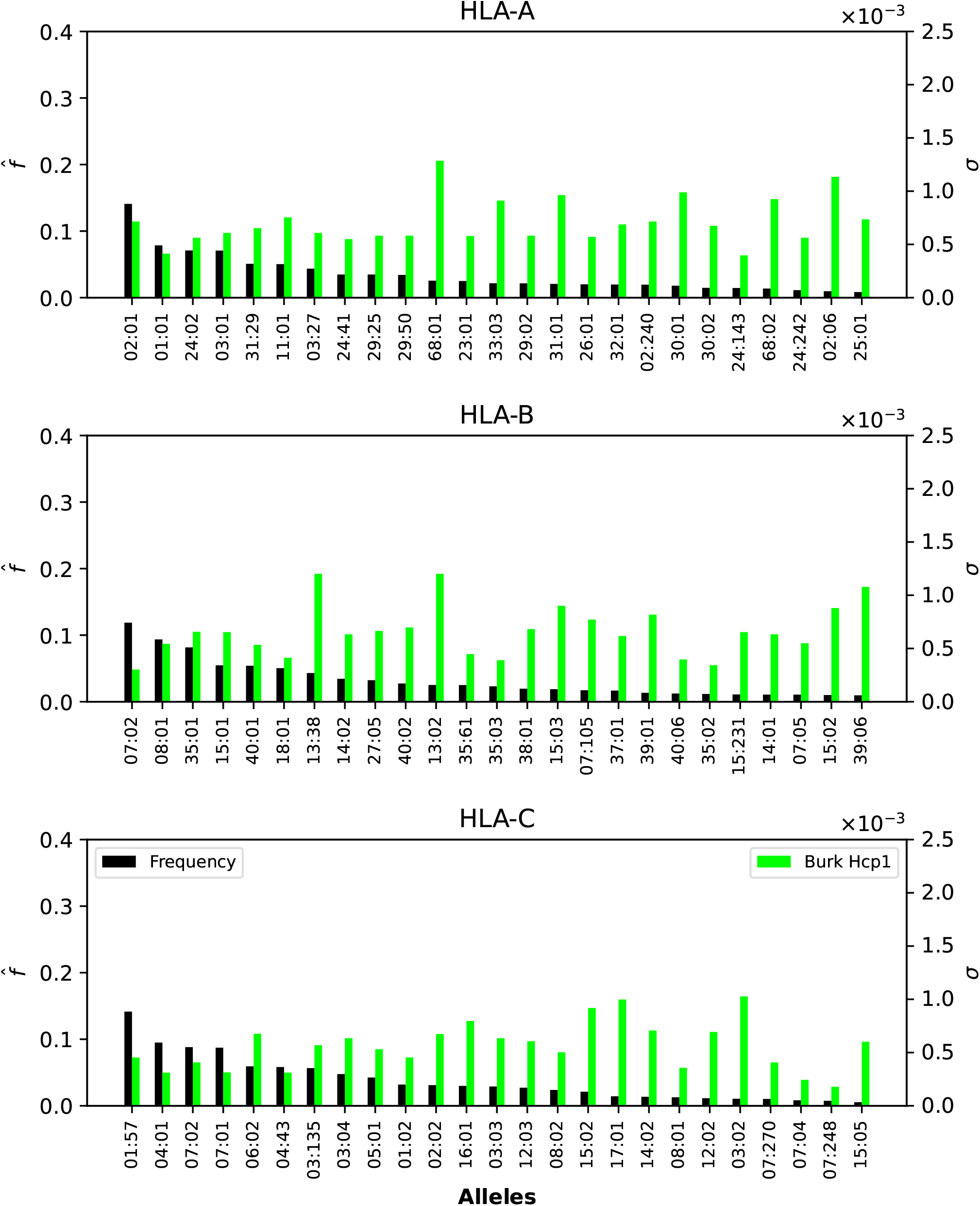
Normalized regional frequencies (left *y* axis), 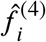, and Burkholderia σ_*i*_ values (right *y* axis) for the top 25 most frequent alleles of each type in North America (*x* axis). The top panel represents HLA-A, the middle HLA-B, and the bottom HLA-C alleles, respectively.

Fig. 5, Fig. 6, and Fig. 7 show that only one allele per type, HLA-A*02:01, HLA-B*07:02, HLA-C*01:57, has a frequency greater than 10%. For Ebola proteins, Fig. 5 shows that σ_*i*_ values are largest (overall) for HLA-B, then HLA-A, and HLA-C. This implies that CD8^+^ T cell responses to Ebola GP or NP proteins will be dominated by HLA-B restricted TCRs. Alleles HLA-A*68:01, HLA-A*30:01, HLA-A*68:02 and HLA-A*02:06 dominate the σ_*i*_ values. For HLA-A*68:01 and Ebola GP Zaire, its σ_*i*_ value is much larger than those of the other three Ebola proteins. In the case of HLA-B alleles, HLA-B*13:38, HLA-B*13:02 and HLA-B*15:03 have the largest σ_*i*_ values, followed by HLA-B*15:02 and HLA-B*39:06, for NP proteins (Sudan and Zaire).

In the case of SARS-CoV-2 spike protein, Fig. 6 shows, as was the case for Ebola, that CD8^+^ T cell responses will be dominated by HLA-B restricted TCRs. HLA-A*68:01 for Wuhan-Hu-1 has a larger σ_*i*_ value when compared to the other variants, and HLA-A*02:06 dominates the σ_*i*_ values for all five variants. The observed trend for HLA-B in Fig. 5 seems to be repeated for SARS-CoV-2, with HLA-B*13:38, HLA-B*13:02 and HLA-B*15:03 having the largest σ_*i*_ values, followed by HLA-B*15:02 and HLA-B*39:06. Contrary to HLA-A*68:01, it is now the Omicron variants that dominate the values. For HLA-C, it is HLA-C*03:02 that has the largest σ_*i*_ values, from lowest to highest as SARS-CoV-2 evolved from Wuhan-Hu-1 to Omicron BA.5.

Finally, Fig. 7 shows that HLA-A and HLA-B Burkholderia σ_*i*_ values are comparable, with HLA-C a bit lower (overall). Those alleles (A, B, or C) identified for their large σ_*i*_ values in Fig. 5 and Fig. 6 dominate as well in the case of Burkholderia Hcp1. It is, thus, interesting to observe that rather different proteins (from two viruses and one bacterium) seem to be binding better to a subset of HLA class I alleles.

### 3.3 Dissecting the individual regional coverage metric: allele pair analysis

We now turn our attention to the individual regional coverage metric for allele pairs. Fig. 8 shows the frequency and individual regional coverage score, 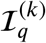, for each allele pair (see Eq. 7) in North America. The top row corresponds to allele frequencies (HLA-A, HLA-B, and HLA-C), the second, third, fourth and fifth to 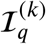 for Ebola GP Zaire, Ebola GP Sudan, Ebola NP Zaire, and Ebola NP Sudan, respectively. Each column thus corresponds to one HLA class I type, HLA-A (left), HLA-B (middle) and HLA-C (right). We observe that overall smaller coverage scores are obtained for HLA-C allele pairs, and that NP proteins and HLA-B allele pairs lead to the largest values, for both Sudan and Zaire variants. For HLA-A, similar coverage scores are obtained for GP and NP proteins, with a slight preference for Zaire versus Sudan. The HLA-B alleles identified in the previous section, HLA-B*13:38, HLA-B*13:02 and HLA-B*15:03, if paired with each other, lead to the largest scores.

**Figure 8.**
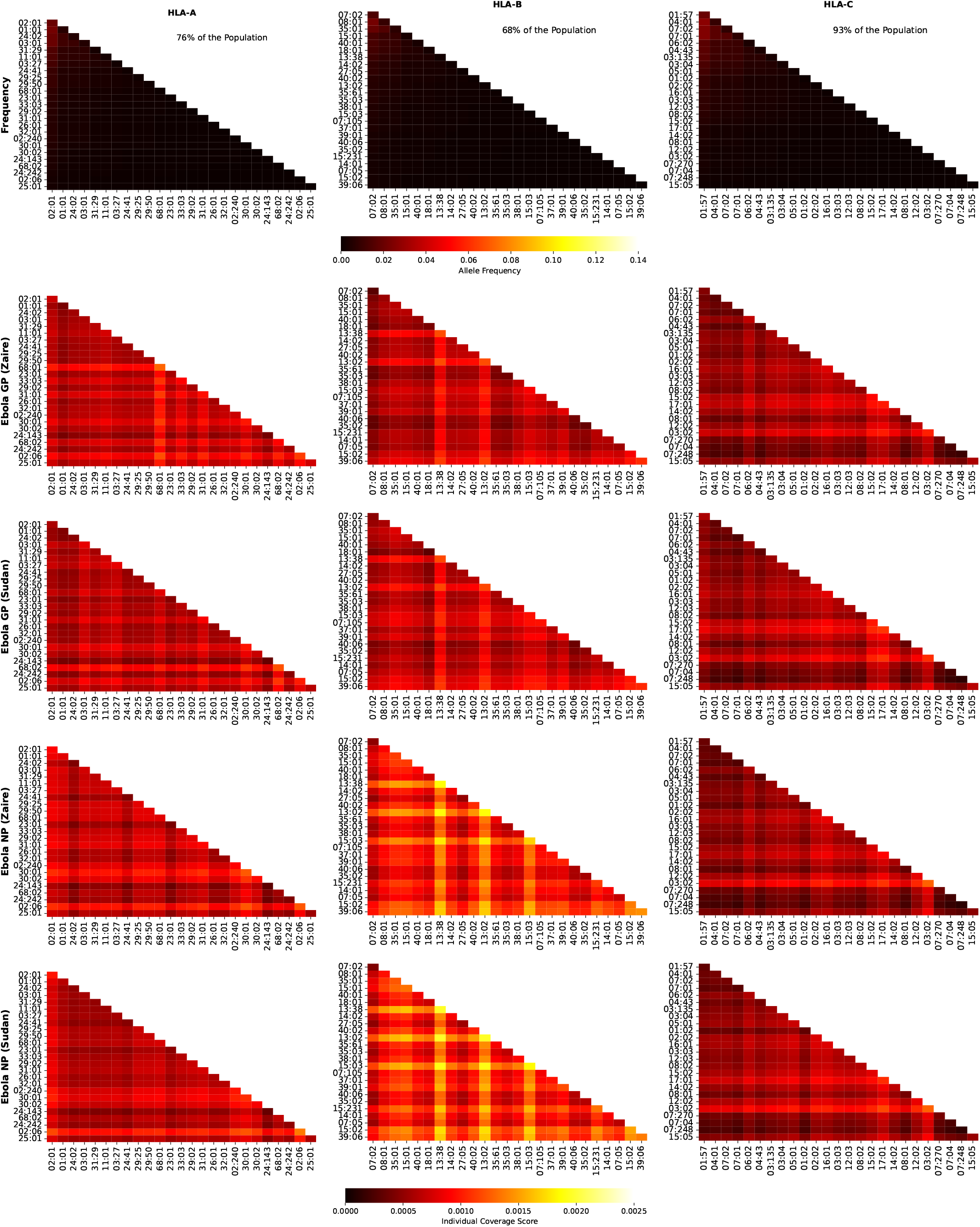
Frequency and individual regional coverage score, 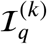, for each allele pair (see Eq. 7) in North America. The top row corresponds to allele frequencies (HLA-A, HLA-B, and HLA-C), the second, third, fourth and fifth to 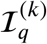 for Ebola GP Zaire, Ebola GP Sudan, Ebola NP Zaire, and Ebola NP Sudan, respectively. Left column corresponds to HLA-A alleles, middle to HLA-B, and right to HLA-C. The sum of the individual frequencies for each allele type is indicated on the panels in the top row.

Fig. 9 shows the frequency and individual regional coverage score, 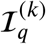, for each allele pair (see Eq. 7) in North America. The top row corresponds to allele frequencies (HLA-A, HLA-B, and HLA-C), the second and third to 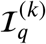 for SARS-CoV-2 spike Wuhan-Hu-1 and Delta AY.4, respectively Each column thus corresponds to one HLA class I type, HLA-A (left), HLA-B (middle) and HLA-C (right). We observe that overall smaller coverage scores are obtained for HLA-C allele pairs, followed by HLA-A, and then HLA-B. There is hardly any difference between the two variants, Wuhan-Hu-1 and Delta AY.4. The HLA-B alleles identified in the previous section, HLA-B*13:38, HLA-B*13:02 and HLA-B*15:03, if paired with each other, lead to the largest scores, which are lower when compared to those in Fig. 8.

**Figure 9.**
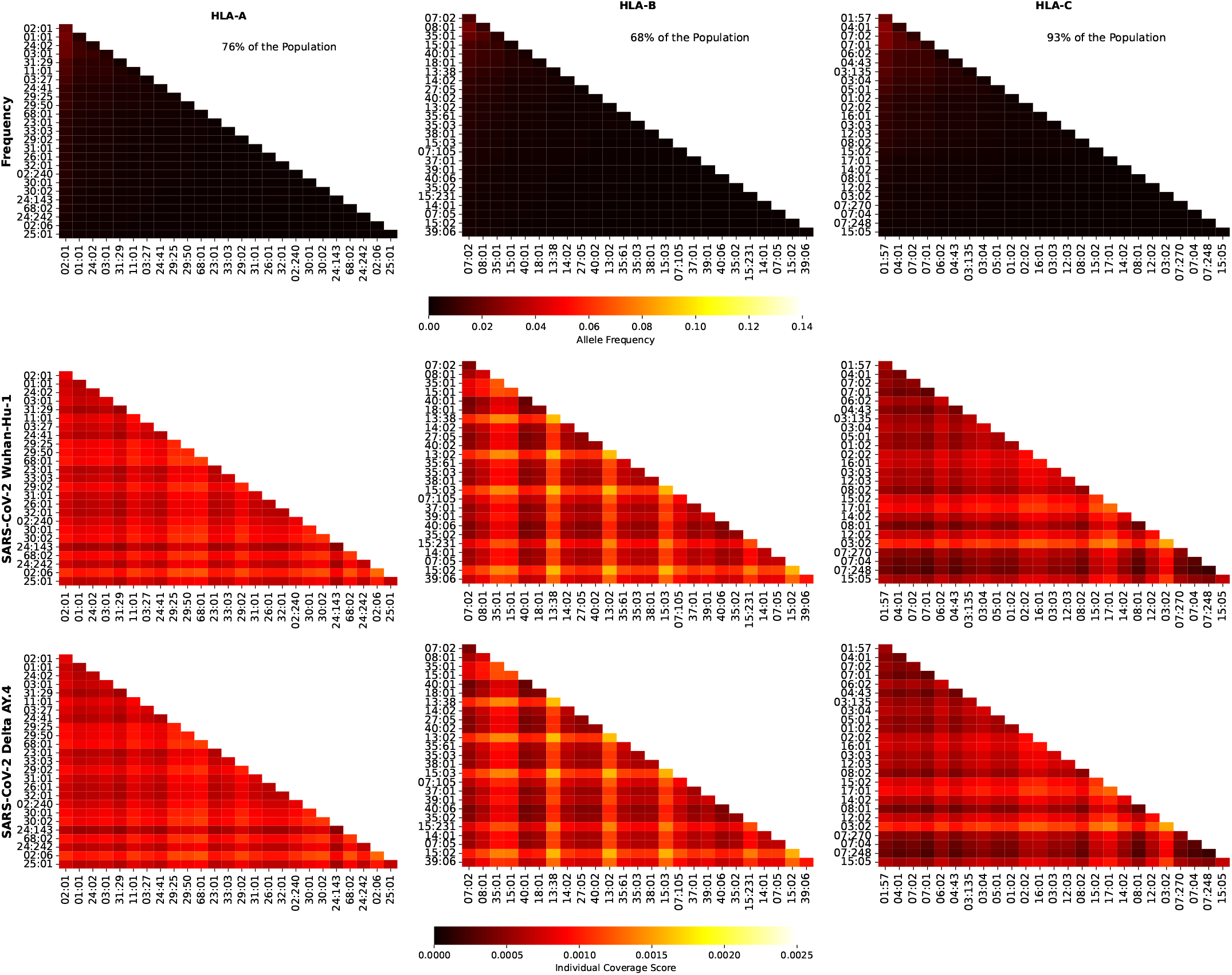
Frequency and individual regional coverage score, 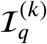, for each allele pair (see Eq. 7) in North America. The top row corresponds to allele frequencies (HLA-A, HLA-B, and HLA-C), the second and third to 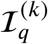 for SARS-CoV-2 spike Wuhan-Hu-1 and Delta AY.4, respectively Left column corresponds to HLA-A alleles, middle to HLA-B, and right to HLA-C. The sum of the individual frequencies for each allele type is indicated on the panels in the top row.

Fig. 10 shows the frequency and individual regional coverage score, 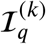, for each allele pair (see Eq. 7) in North America. The top row corresponds to allele frequencies (HLA-A, HLA-B, and HLA-C), the second, third, and fourth to 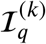 for SARS-CoV-2 spike Omicro BA.1, BA.2, and BA.5, respectively. Each column thus corresponds to one HLA class I type, HLA-A (left), HLA-B (middle) and HLA-C (right). No significant differences can be found between this figure and Fig. 9, indicating, in agreement with the results Ref. [37], that CD8^+^ T cell responses elicited by the SARS-CoV-2 spike vaccine (Wuhan ancestral sequence) will be protective and cross-reactive against Omicron variants.

**Figure 10.**
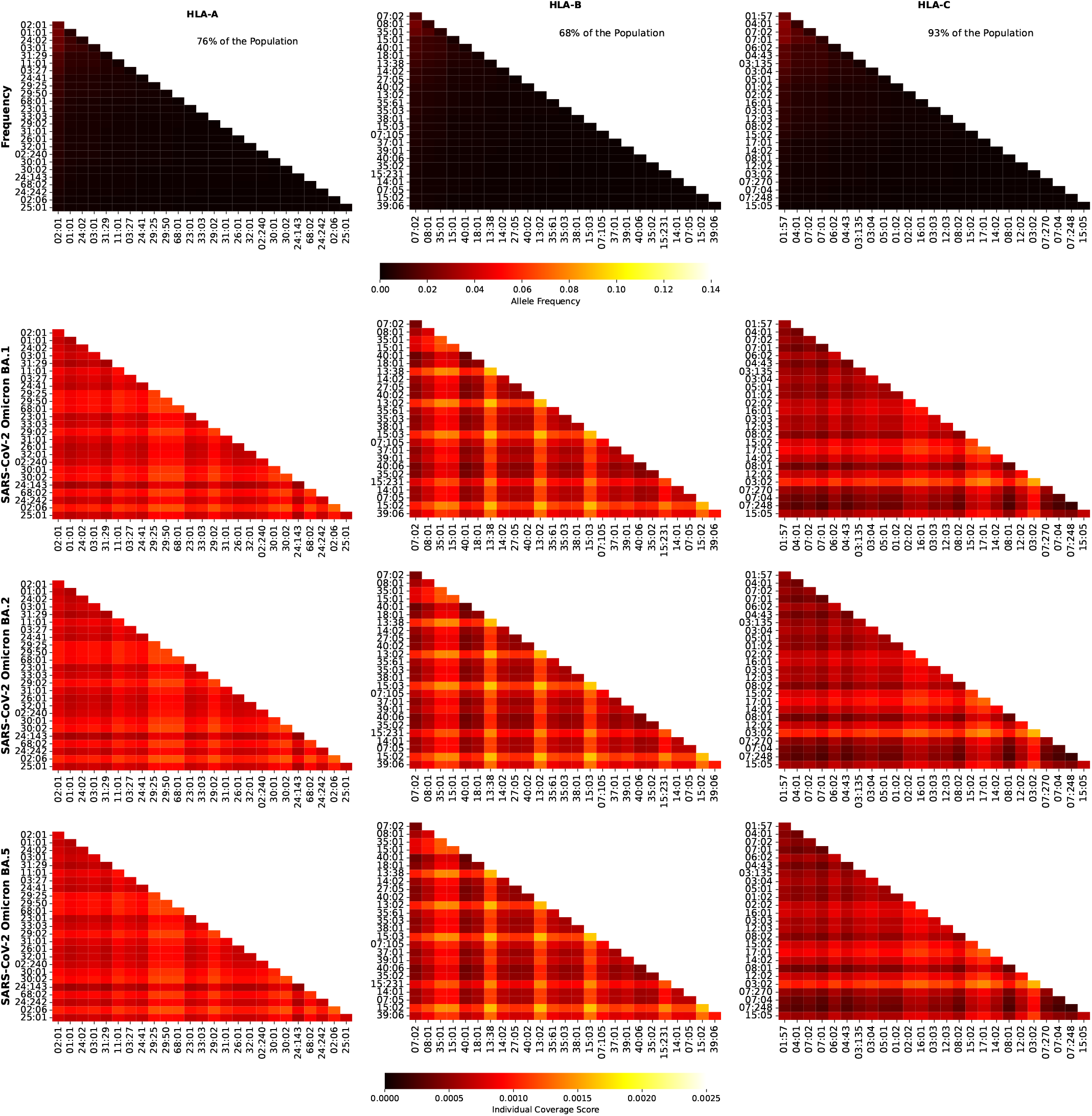
Frequency and individual regional coverage score, 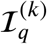, for each allele pair (see Eq. 7) in North America. The top row corresponds to allele frequencies (HLA-A, HLA-B, and HLA-C), the second, third, and fourth to 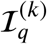 for SARS-CoV-2 spike Omicron BA.1, BA.2, and BA.5, respectively. Left column corresponds to HLA-A alleles, middle to HLA-B, and right to HLA-C. The sum of the individual frequencies for each allele type is indicated on the panels in the top row.

Fig. 11 shows the frequency and individual regional coverage score, 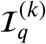, for each allele pair (see Eq. 7) in North America. The top row corresponds to allele frequencies (HLA-A, HLA-B, and HLA-C), and the bottom to 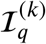 for *Burkholderia* Hcp1 protein. Each column thus corresponds to one HLA class I type, HLA-A (left), HLA-B (middle) and HLA-C (right). For the *Burkholderia* Hcp1 protein, we observe that the dominant individual coverage scores correspond to HLA-A, followed by HLA-B, and then HLA-C. The HLA-B alleles that were identified, both for Ebola NP and for SARS-CoV-2 spike, with high 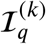 values, do not play such a significant role in the case of the Hcp1 protein.

**Figure 11.**
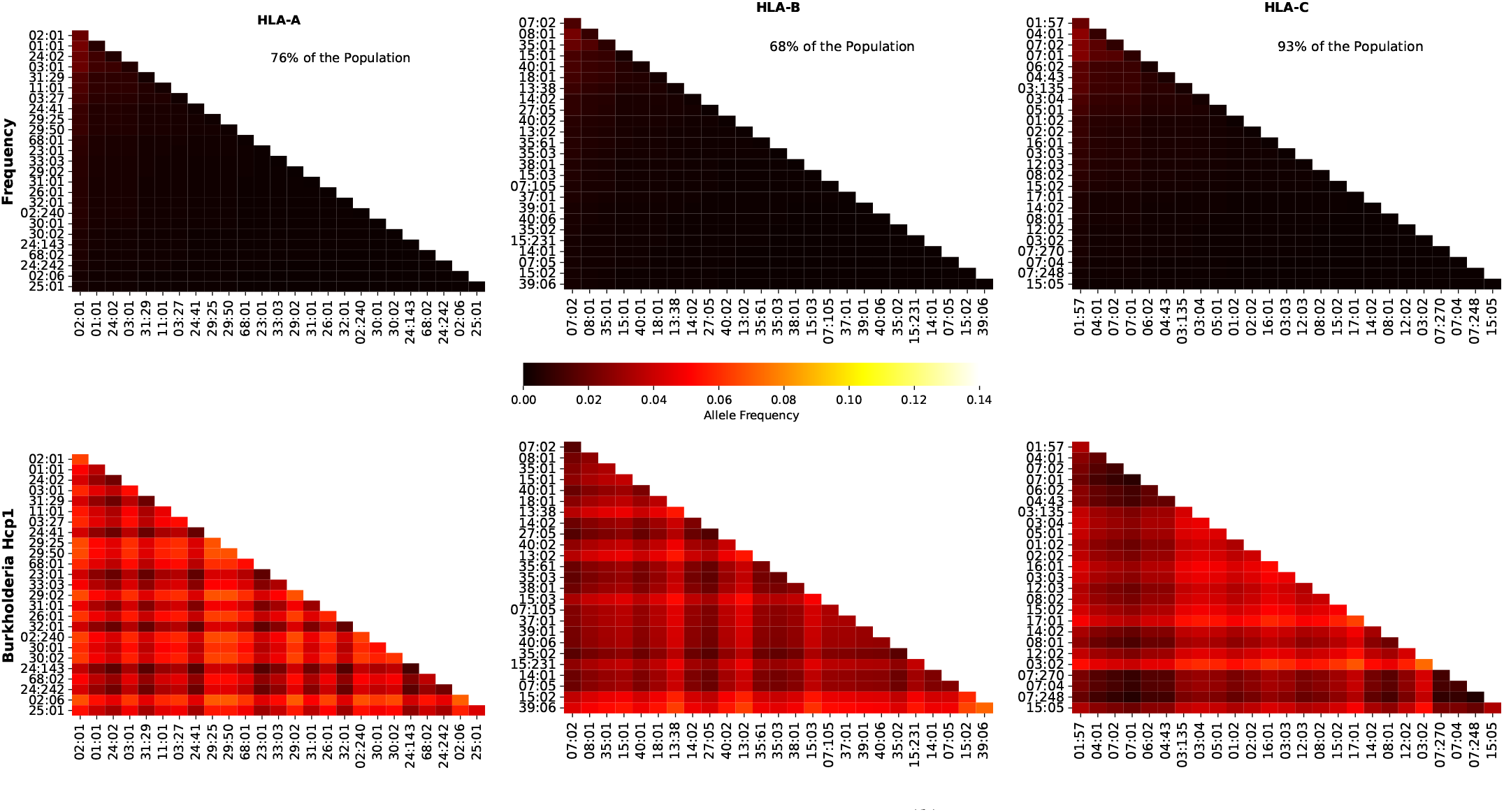
Frequency and individual regional coverage score, 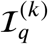, for each allele pair (see Eq. 7) in North America. The top row corresponds to allele frequencies (HLA-A, HLA-B, and HLA-C), and the bottom to 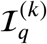 for *Burkholderia* Hcp1 protein. Left column corresponds to HLA-A alleles, middle to HLA-B, and right to HLA-C. The sum of the individual frequencies for each allele type is indicated on the panels in the top row.

### 3.4 Contribution of immuno-dominant epitopes to mean coverage metric

We next analyze the contribution of the immuno-dominant epitopes to the mean coverage metric, as defined by the ratio ℱ_*k*_ in Eq. (11). Immuno-dominant epitopes have been identified for Ebola GP (Zaire and Sudan) and SARS-CoV-2 spike protein in section 2.3.

Fig. 12 displays, per geographical region, the values of ℱ_*k*_ for the different proteins considered, and the three different HLA class I types, HLA-A (top), HLA-B (middle) and HLA-C (bottom), respectively. We note that the overall highest contributions from the immuno-dominant epitopes correspond to HLA-A alleles, with Ebola GP Zaire leading, for all regions, except for South and Central America. The contribution for the different SARS-CoV-2 immuno-dominant epitopes is largest for the Wuhan-Hu-1 variant, decreasing for Delta AY.4 and Omicron BA.1, and then increasing for both Omicron BA.2 and BA.5. For HLA-B alleles, ℱ_*k*_ is clearly largest for Ebola GP Zaire (around 6%), and lower for the SARS-CoV-2 spike immuno-dominant epitopes and Ebola GP Zaire (around 2%). The situation seems reversed for HLA-C alleles, where the SARS-CoV-2 spike immuno-dominant epitopes lead to the largest values of ℱ_*k*_ (around 5%). In this instance, Ebola GP Zaire is around 1% and much lower for the Ebola GP Sudan.

**Figure 12.**
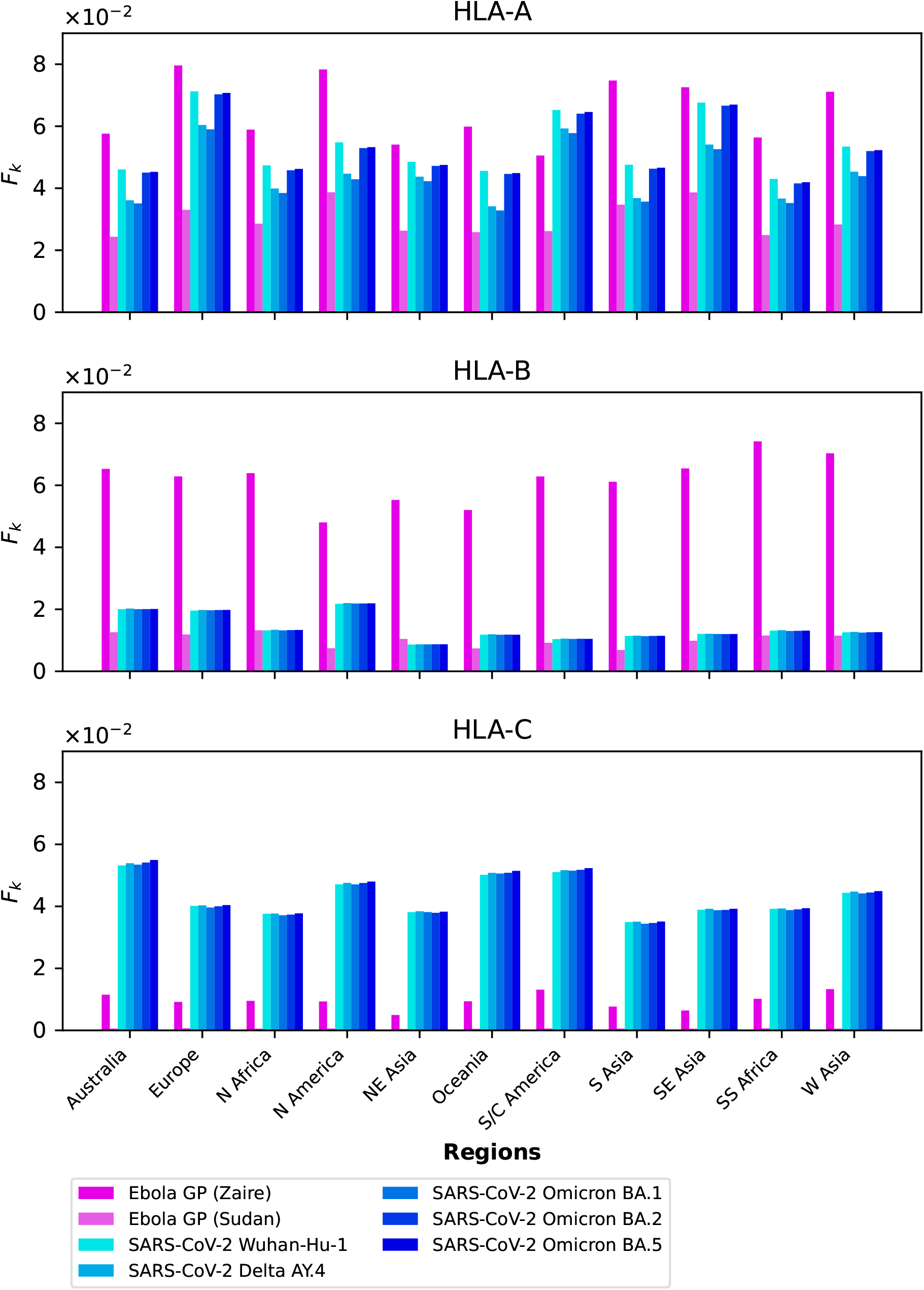
**ℱ**_*k*_ grouped by geographical region for Ebola GP and SARS-CoV-2 spike immuno-dominant epitopes, and for HLA-A (top), HLA-B (middle), and HLA-C (bottom).

Fig. 13 displays, per pathogen, the values of ℱ_*k*_ for the different proteins considered, and the three different HLA class I types, HLA-A (top), HLA-B (middle) and HLA-C (bottom), respectively. It is interesting to observe that for HLA-A alleles, and across proteins, the largest contribution from immuno-dominant epitopes to the mean regional coverage metric is achieved in Europe. Whereas for HLA-C alleles, the leading region is Australia, followed closely by South and Central America, Oceania, and North America.

**Figure 13.**
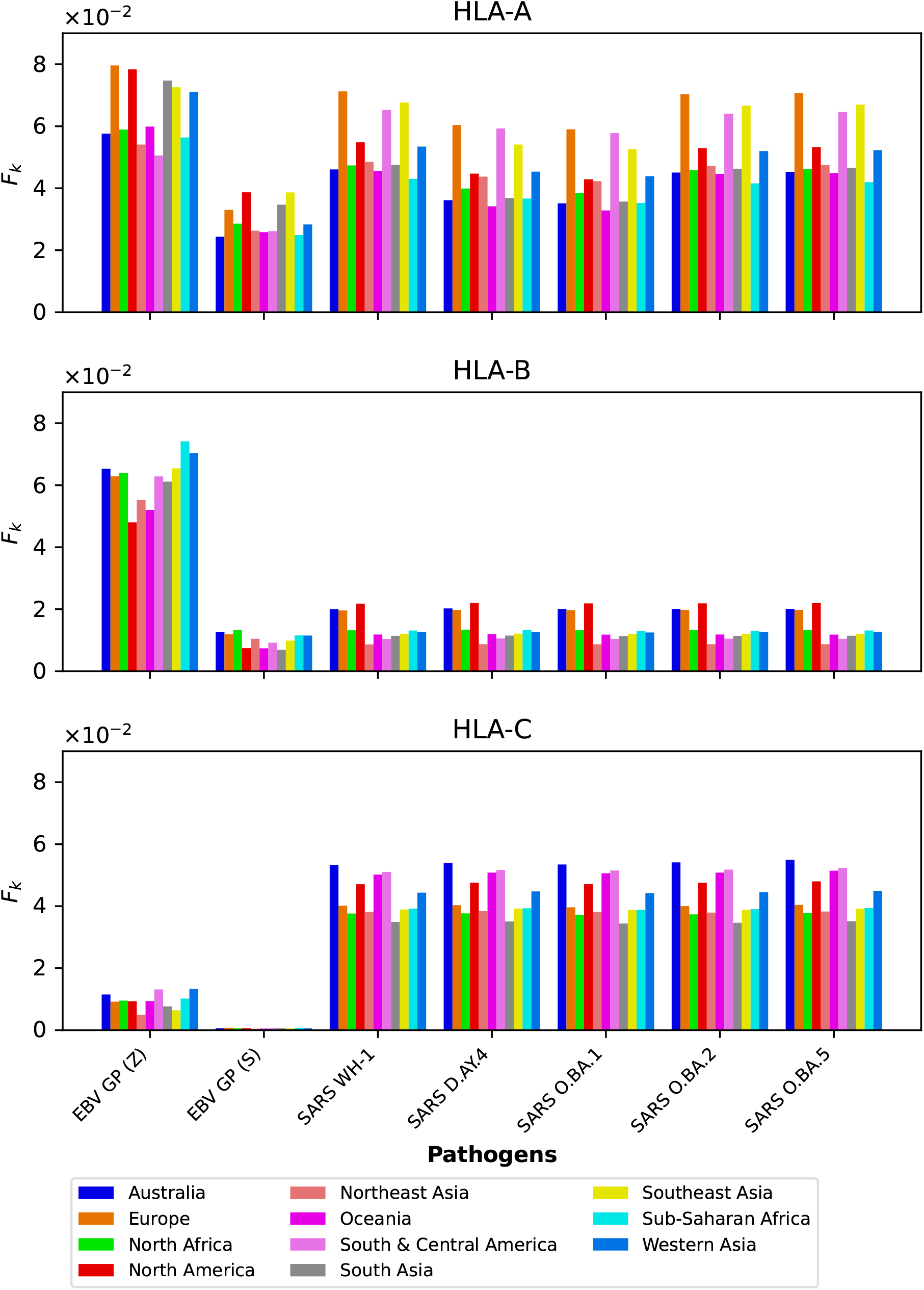
**ℱ**_*k*_ grouped by protein for the eleven different geographical regions, and for HLA-A (top), HLA-B (middle), and HLA-C (bottom).

### 3.5 Distributions of immuno-dominant epitopes

We now display the results from the analysis of the probability distributions for *g*_*j*_ and *ϕ*_*j*_ (see section 2.3).

Fig. 14 and Fig. 15 show the *g*_*j*_ probability distributions for Ebola GP and SARS-CoV-2 spike protein, respectively. We have identified individual values corresponding to the immuno-dominant epitopes. Our results indicate that the immuno-dominant epitopes do not have significantly larger immunogenicity values, when compared to non-immuno-dominant ones.

**Figure 14.**
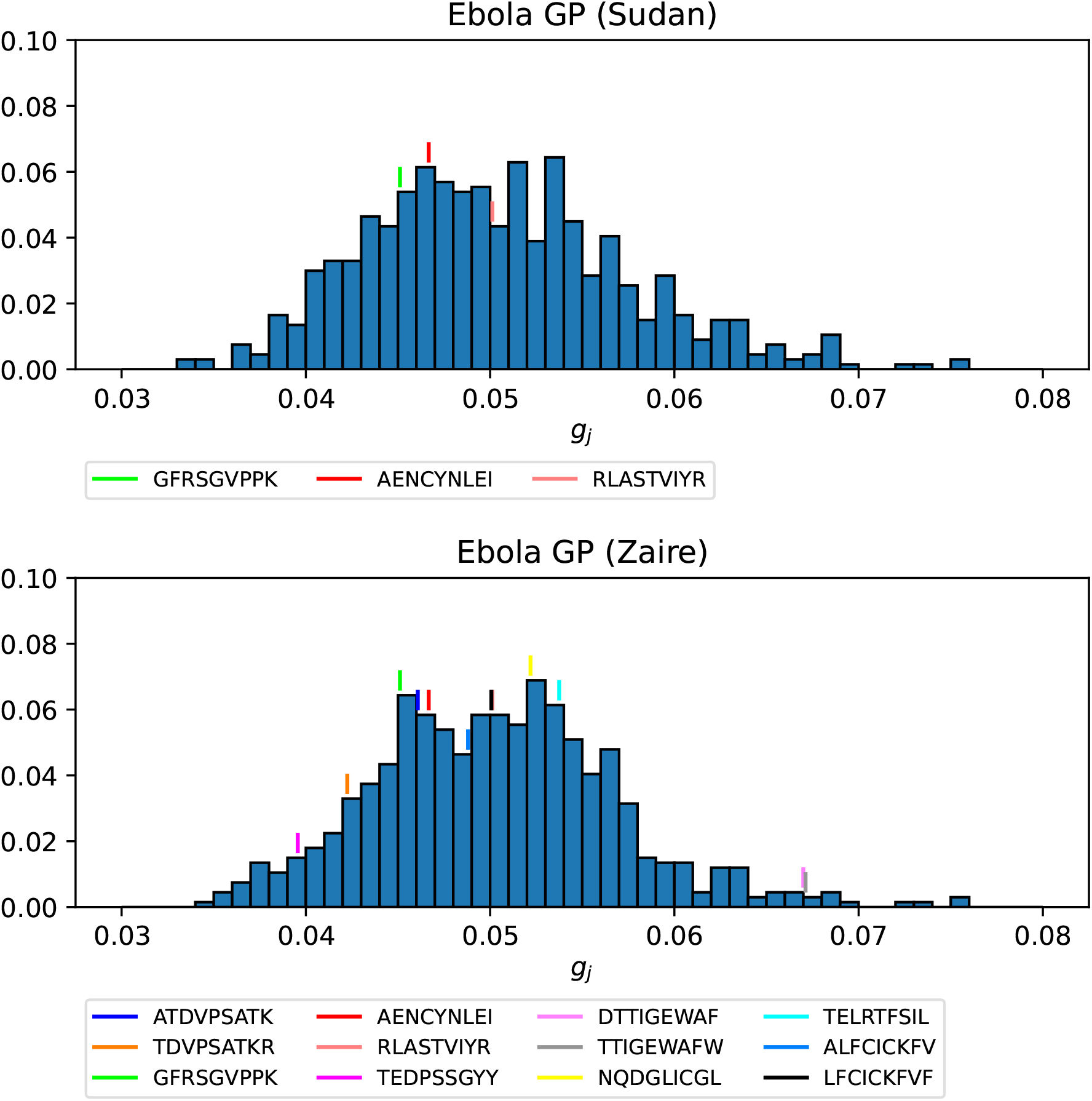
Probability distribution for the immunogenicity, *g*_*j*_, of the nonamers of Ebola GP Sudan (top) and Ebola GP Zaire (bottom). Individual values corresponding to the immuno-dominant epitopes have been identified.

**Figure 15.**
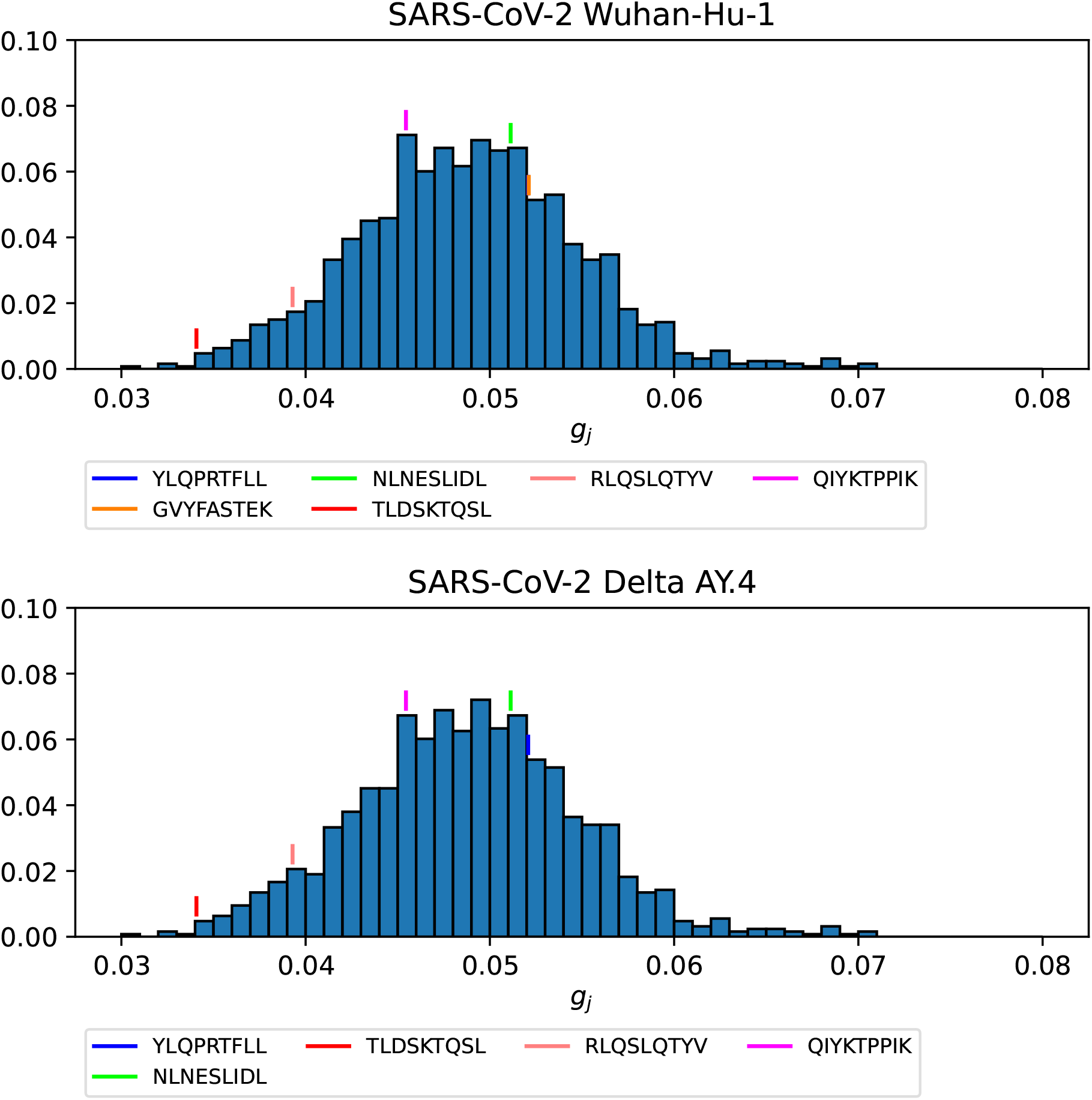
Probability distribution for the immunogenicity, *g*_*j*_, of the nonamers of SARS-CoV-2 Wuhan-Hu-1 spike (top) and SARS-CoV-2 Delta AY.4 spike (bottom). Individual values corresponding to the immuno-dominant epitopes have been identified.

Fig. 16, Fig. 17, and Fig. 18 show the *ϕ*_*j*_ probability distributions for Ebola GP Sudan, Ebola GP Zaire, and SARS-CoV-2 spike proteins, respectively, for North America, and for the three HLA class I types. We have identified individual values corresponding to the immuno-dominant epitopes. Our results indicate that the immuno-dominant epitopes have a significantly larger *ϕ*_*j*_ value, when compared to non-immuno-dominant ones. For instance, Fig. 16 shows that for HLA-A nonamer RLASTVIYR belongs to the tail of the distribution, and the same is true for HLA-B nonamer TELRTFSIL (see Fig. 17). In the case of immuno-dominant epitopes for SARS-CoV-2 spike protein, Fig. 18 indicates that nonamer YLQPRTFLL belongs to the tail of the distribution for HLA-A, as well as HLA-B and HLA-C, and so does nonamer TLDSKTQSL for HLA-B and HLA-C. These results indicate that the immuno-dominance of the nonamers is determined not so much by their immunogenicity, as defined by Eq. (4), but by their associated binding scores to HLA-class alleles (see Eq. (15)). Furthermore, since our results indicate that immuno-dominant epitopes belong to the tail of certain probability distributions, they provide an indirect validation of the methods proposed here to characterize vaccine coverage.

**Figure 16.**
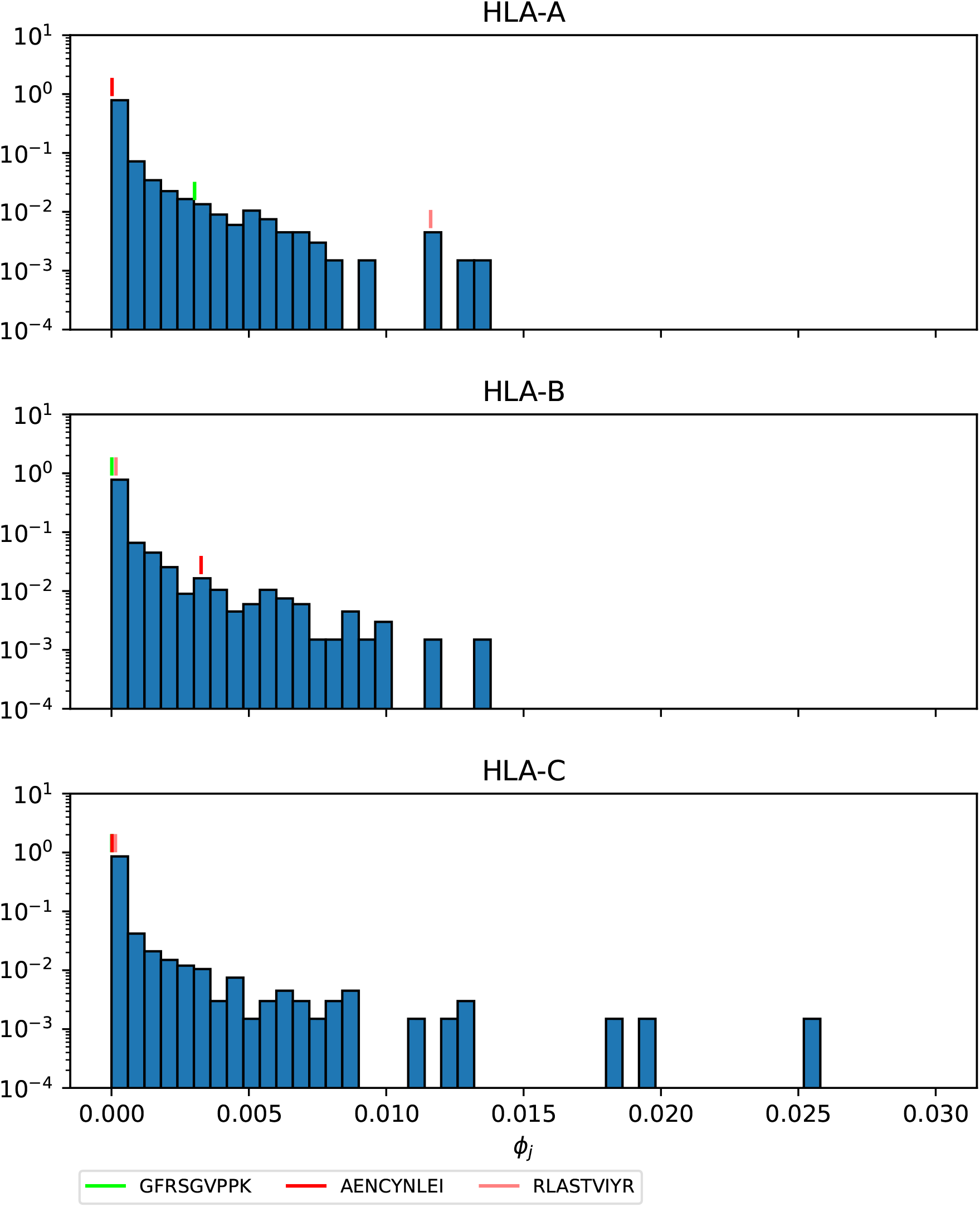
*ϕ*_*j*_ probability distribution in North America of the nonamers for Ebola GP Sudan, with HLA-A (top), HLA-B (middle), and HLA-C (bottom). Individual values corresponding to the immuno-dominant epitopes have been identified.

**Figure 17.**
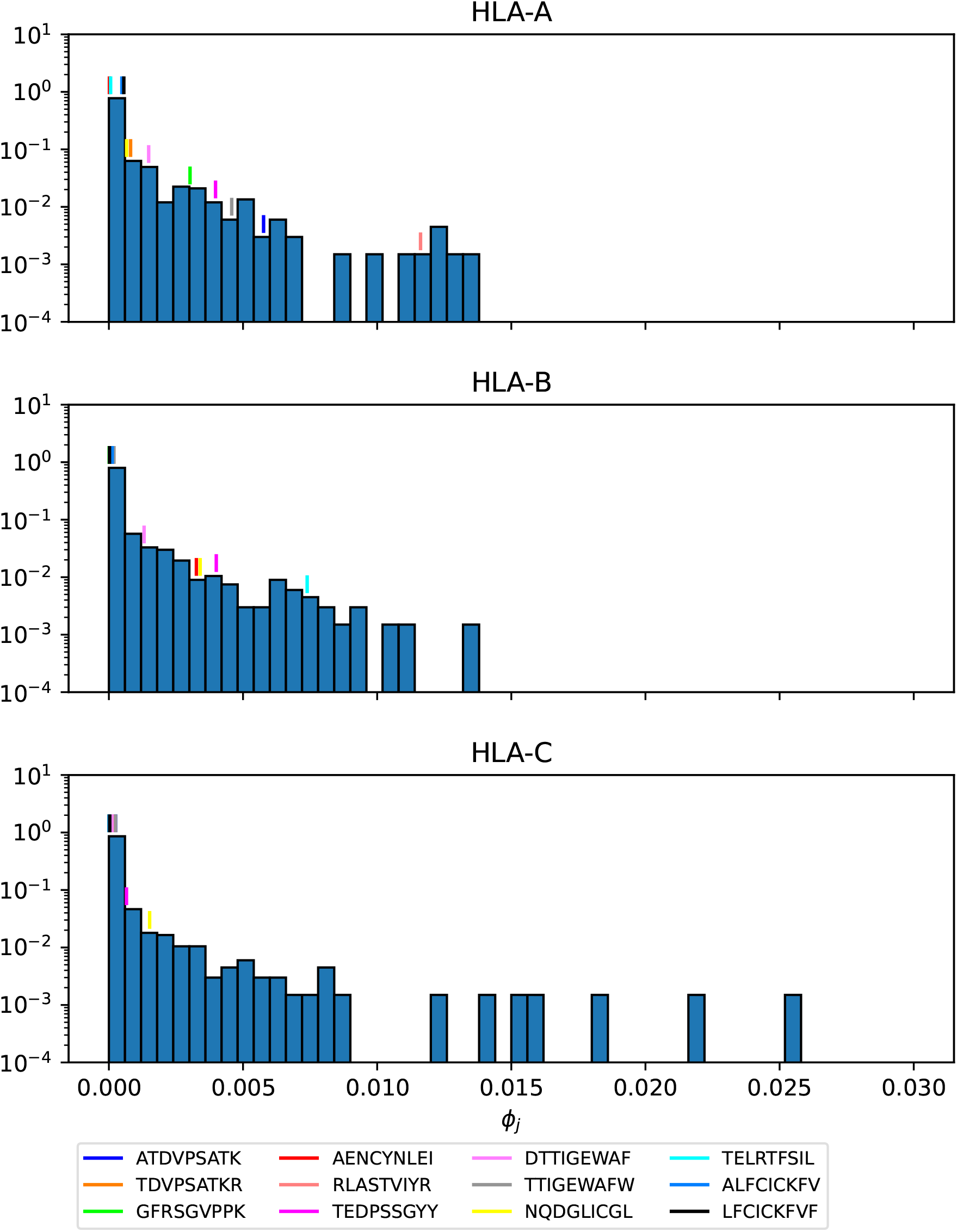
*ϕ*_*j*_ probability distribution in North America of the nonamers for Ebola GP Zaire, with HLA-A (top), HLA-B (middle), and HLA-C (bottom). Individual values corresponding to the immuno-dominant epitopes have been identified.

**Figure 18.**
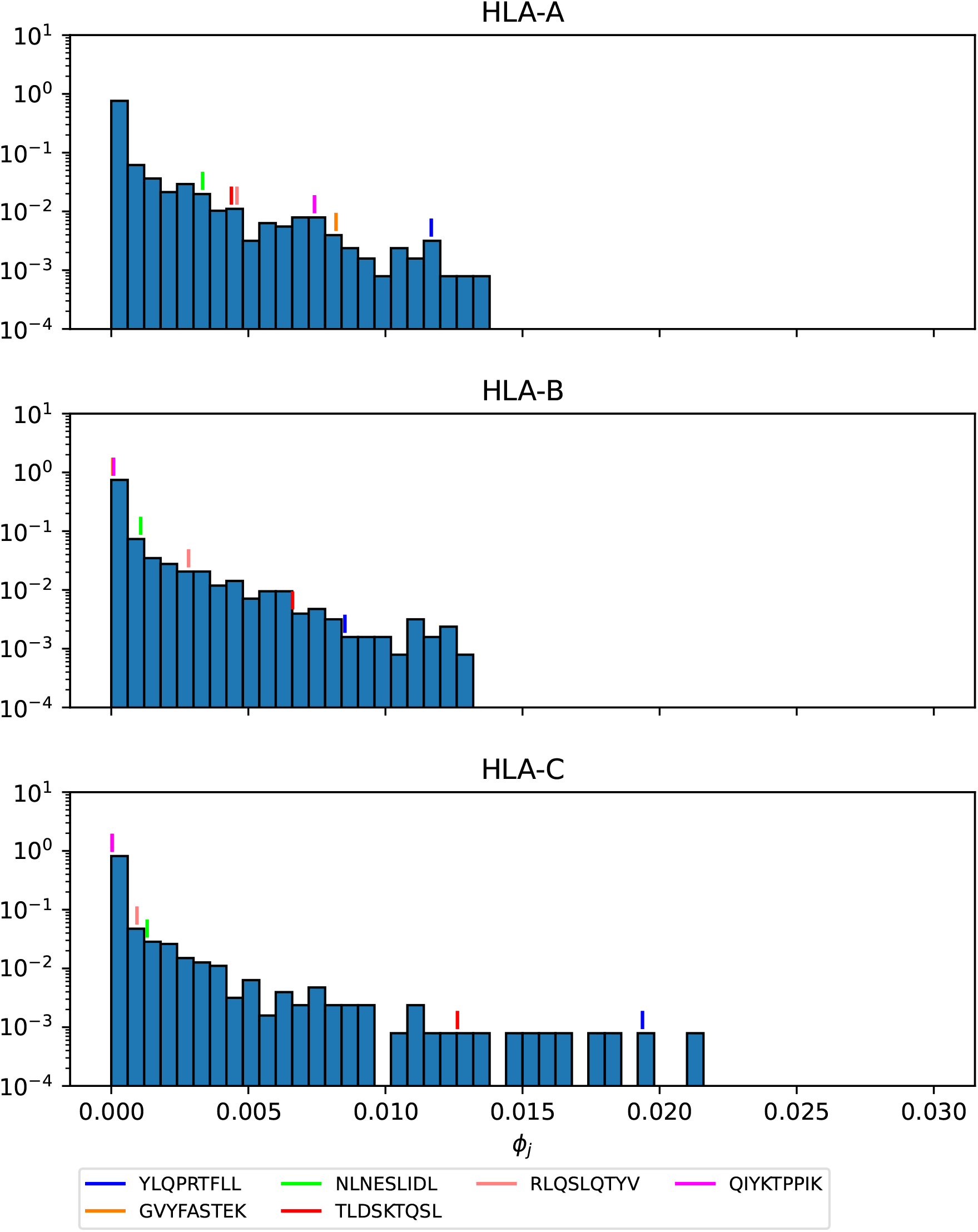
*ϕ*_*j*_ probability distribution in North America of the nonamers for SARS-CoV-2 Wuhan-Hu-1 spike, with HLA-A (top), HLA-B (middle), and HLA-C (bottom). Individual values corresponding to the immuno-dominant epitopes have been identified.

## 4 DISCUSSION

Sterilizing immunity, provided by (pre-exisiting) neutralizing antibodies, has been recognized as the ideal immune response and primary goal of vaccine design to control pathogens, viruses or bacteria [38]. Important human pathogens such as herpes viruses, *Mycobacterium tuberculosis*, malaria, and HIV pose a challenge in light of antigenic evolution and antibody immune escape, since vaccines which induce antibody responses (humoral immune responses) are ineffective against them [39, 38]. CD8^+^ T cells, elements of the adaptive cellular arm of the immune system [1], have been shown to mediate protection during infection with these pathogens, as reviewed in Refs. [39, 38]. More recently, substantial evidence has emerged of the protective role of CD8^+^ T cell-mediated responses to *conserved regions* of the genome of HIV-1 [4], Lassa virus [5, 40], SARS-CoV-2 [6, 7], pandemic influenza [8], and Ebola virus [9]. Yet, we still do not have a single metric to define protective T cell immune responses. This is a huge challenge given the phenotypic and multi-functional heterogeneity of T cell responses, and TCR diversity and cross-reactivity [39, 14].

In this paper, we aim to develop a novel framework to quantify the potential of CD8^+^ T cells to induce vaccine-mediated immune responses, and in turn, propose such a metric. The MHC-restriction of T cell receptor antigen recognition brings an additional and crucial consideration, since the HLA locus is the most polymorphic gene cluster of the entire human genome [10]. Our proposed solution is based on the hypothesis that a multi-partite graph (see Fig. 2) is the natural framework to consider: *1)* viral genetic diversity of the pathogen as represented in the set of peptides, 𝒫, so that wild type and all circulating (or predicted) variants can be analyzed, *2)* HLA variability as considered with regard to geographical regions ℛ, HLA alleles 𝒜, and their frequencies within each region, and *3)* TCR recognition variability as accounted for by *peptide immunogenicity* [16].

The multi-partite graph, together with HLA class I frequencies (for HLA-A, HLA-B, and HLA-C types) in eleven different geographical regions (see section 2.1.1), binding scores of HLA class I alleles to nonamers (see section 2.1.2), and peptide immunogenicity [16] (see section 2.1.3), allow us to define a mean regional coverage metric in Eq. (5) for a given vaccine protein. Fig. 3 and Fig. 4 show our results for the ten different proteins considered here: Ebola virus (GP and NP, Sudan and Zaire), SARS-CoV-2 spike (five variants), and *Burkholderia pseudomallei* Hcp1. We then argue that the mean regional coverage metric does not capture the fact that an individual carries two alleles, and not *M* different ones. Thus, we propose the individual regional coverage metric in Eq. (7), and the mean individual regional coverage metric in Eq. (8) to account for this important difference. In the absence of allele associations, we show that both metrics are the same. We conclude that were we to obtain true allele pair frequencies, instead of the individual allele frequencies used here, the mean individual regional coverage metric would be the true metric for CD8^+^ T cell immune responses. Finally, we discuss immuno-dominance and immuno-dominant epitopes [10], in light of recent studies for Ebola GP and SARS-CoV-2 spike protein [36, 35]. We make use of the immuno-dominant epitopes identified in these studies (see Table 4 and Table 3), together with our approaches, to calculate the contribution of the immuno-dominant epitopes to the mean regional coverage metric (see section 3.4), and to show that for suitably defined probability distributions (see section 2.3) the immuno-dominant peptides belong to the tail of the distribution. In fact, Fig. 12 and Fig. 13 show that the subset of *η* different immuno-dominant epitopes make a significant contribution to the mean regional coverage metric, which is of the order of 5% for HLA-A and Ebola GP Zaire and SARS-CoV-2 spike across regions, as well as for HLA-B and Ebola GP Zaire, and HLA-C and SARS-CoV-2 spike. We note that for Ebola GP Zaire there are *η* = 12 different immuno-dominant nonamers, out of a total of *P* = 676; that is, the set of immuno-dominant nonamers is less than 2% of the total nonamer set. In the case of SARS-CoV-2 Wuhan-Hu-1 spike protein *η* = 6 and *P* = 1273, which implies the set of immuno-dominant nonamers is less than 0.5% of the total nonamer set. These results and the figures included in section 3.5 provide a first validation of the metrics defined here, since they capture the *singular* nature of the small subset of immuno-dominant epitopes.

There are a number of limitations to our study. First of all, the multi-partite graph does not include important processes such as the processing and presentation of CD8^+^ T cell epitopes, or the expression levels of different MHC molecules (HLA-A, HLA-B, or HLA-C). These could be considered in our methods as node weights; for instance, the level of expression of allele *a*_*i*_ (the level of processing and presentation of peptide *p*_*j*_) could be included in the graph as a node weight *e*_*i*_ (node weight *π*_*j*_). Secondly, and as a proxy for TCR diversity, we have made use of the concept of nonamer immunogenicity [16]. This is clearly not the full story, and methods such as TCRdist [41], together with single cell, paired *α* and *β* TCR sequencing, are providing us with extremely valuable insights into the identification of public T cell receptors which mediate protection against SARS-CoV-2 infection [42]. Furthermore, recent work by Chen *et al*. has shown that TCR sequences are the most important and quantitative factor determining both the phenotype and persistence of specific CD8^+^ T cells against immunogenic viral antigens from SARS-CoV-2, cytomegalovirus, and influenza virus [43]. Thus, our future work will be along this direction to include the role of the full set 𝒯, as well as the edges between elements of 𝒫 and 𝒯. The metrics proposed here can be (easily) generalized to account for TCR diversity.

Looking forward there is a lot of work ahead of us. We will take advantage of the multi-partite graph approach to evaluate differences in vaccine platform antigen presentation. To generate effective CD8^+^ T cells, the cross-presentation of antigen on the MHC class I molecule is critical. Generally, cross-presentation depends on delivery to lymph nodes, uptake by dendritic cells (DCs), and the ability to get antigen into the cytosol of antigen presenting cells (APCs), primarily DCs [44]. In a typical antigen presentation process, proteins in the cytosol of APCs are broken down into peptides and delivered to the endoplasmic reticulum for loading and presentation in MHC class I molecules by a transporter associated with antigen presentation (TAP). To generate cross-presentation, one must enhance both vacuolar and cytosolic pathways [45]. Here, sequence and conformation of the antigens and their lifetimes could affect the cross-presentation process. Along with the chosen adjuvant, a given vaccine platform that is used for antigen presentation can influence or alter the efficiency of these processes. Therefore, we intend to use this model to better inform us on the ability of a chosen vaccine platform to favor cross-presentation.

As mentioned above, we want to explore the role of allele associations and aim to obtain allele pair frequencies to compare the two metrics proposed [46]. We would like to apply our methods to other pathogens of public health relevance such as Lassa virus and Crimean Congo hemorrhagic fever virus, with the viral sequences provided in Refs. [47, 48] Another avenue we have failed to explore is that of immune evasion and the role of MHC-restriction [17] in eliciting HLA-mediated selective pressure [11, 12, 13]. We plan to make use of the computational methods developed by Hertz *et al*. [17] and the approaches adopted here to quantify the potential of a vaccine protein to exert immune pressure and drive viral evolution in different human populations, as well as to identify HLA generalists and specialists [29]. Finally, the CD8^+^ T cell metrics proposed here do not account for T cell function (cytokine secretion, proliferative capacity, or cytotoxic killing activity) or T cell half-life (of particular relevance for central and effector memory T cells). We propose to make use of the multi-partite graph developed here, together with mathematical models of viral and immune dynamics [49, 50, 51, 52, 53], to identify and quantify other potential correlates of immune protection, such as half-lives of cellular subsets of interest, as well as their function and phenotype [54].

## ACKNOWLEDGMENTS

This manuscript has been reviewed at Los Alamos National Laboratory and assigned report number LA-UR-24-23493.

## SUPPLEMENTAL DATA

A separate file, Supplementary Material, includes our extended analysis to all geographical regions other than North America.

## Footnotes

1 A data set is gold standard if allele frequency sums to 1, sample size is greater than 50, and it has four digit resolution. A data set is silver standard if allele frequency sums to 1, sample size is any, and it has mixed two/four or more digits [26].

2 The number of locations is different for each region.

3 We also note that the set 𝒫 depends on the choice of pathogen; for instance, the set for Ebola (Sudan) GP protein is different from that of Ebola (Zaire) GP. The same is true for each of the five different SARS-CoV-2 spike variants considered here.

