## Supplementary file for "Quantification of heterogeneity in human CD8^+^ T cell responses to vaccine antigens: an HLA-guided perspective"

### *Supplementary Material*

#### 1 ANALYSIS OF IMMUNO-DOMINANT EPITOPES: PROBABILITY DISTRIBUTIONS FOR $g_j$ AND $\phi_j$

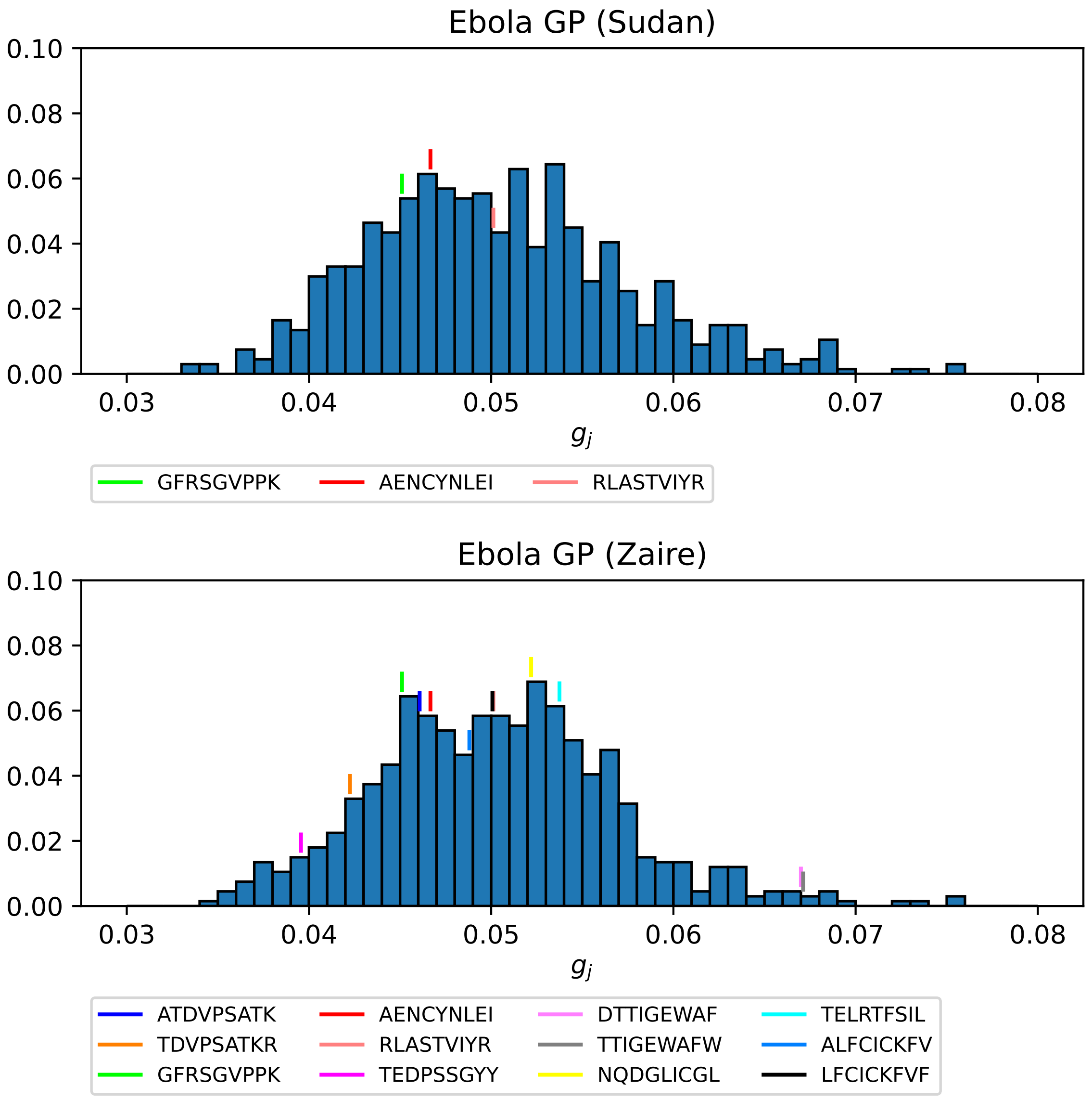


**Figure S1.** Top: Ebola GP (Sudan), Bottom: Ebola GP (Zaire)

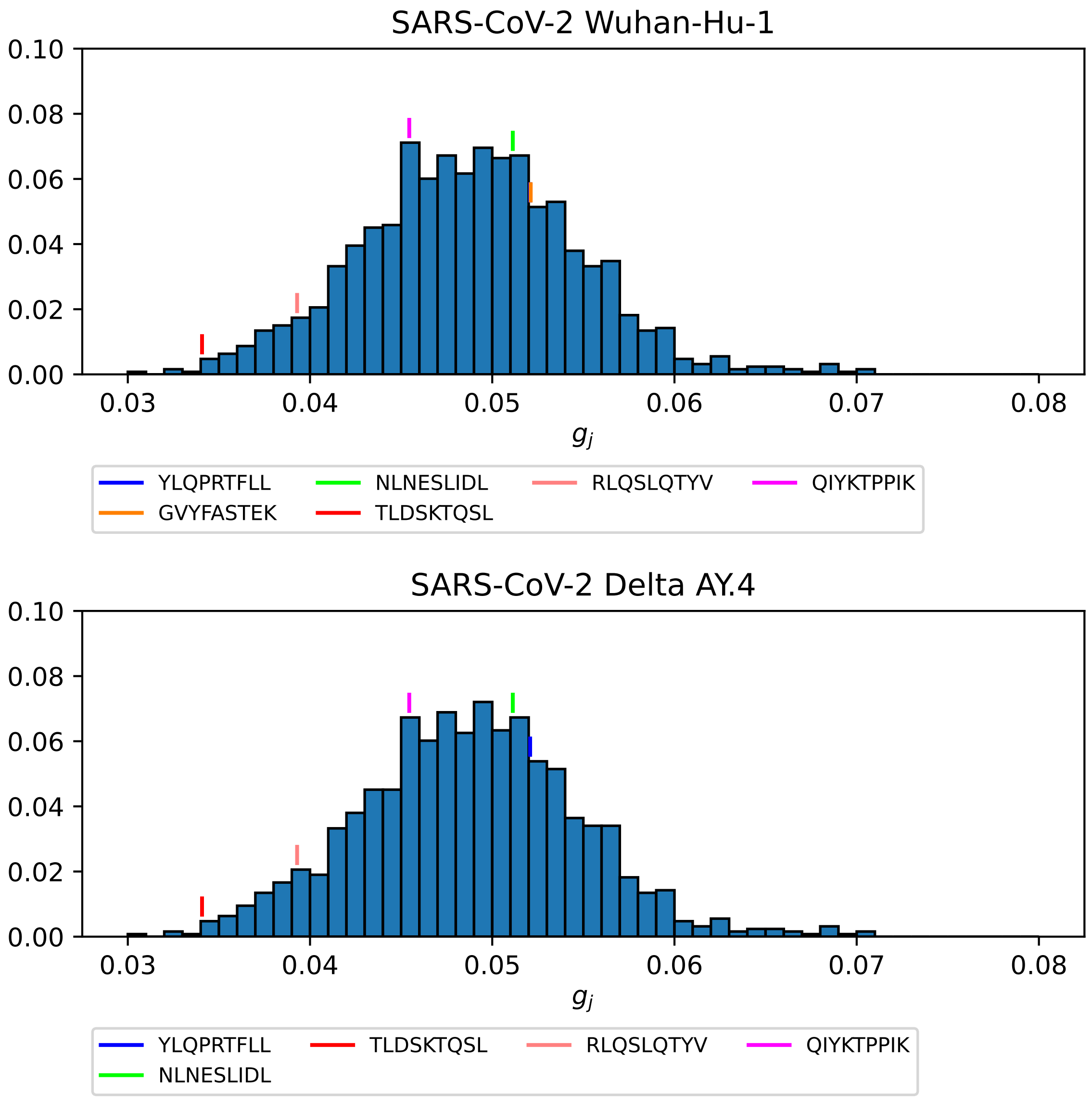


**Figure S2.** Top: SARS-CoV-2 Wuhan-Hu-1, Bottom: SARS-CoV-2 Delta AY.4

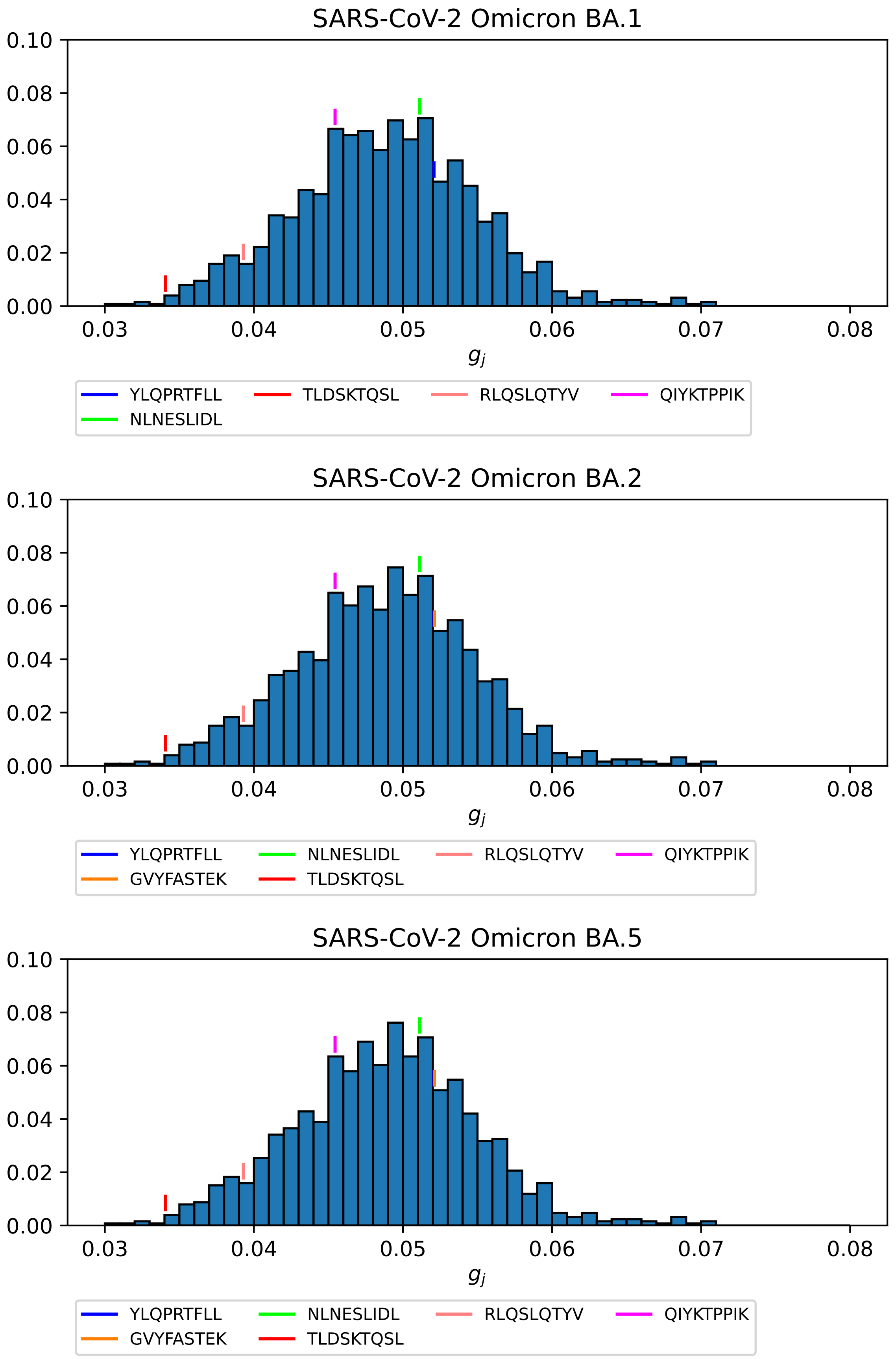


**Figure S3.** Top: SARS-CoV-2 Omicron BA.1, Middle: SARS-CoV-2 Omicron BA.2, Bottom: SARS-CoV-2 Omicron BA.5

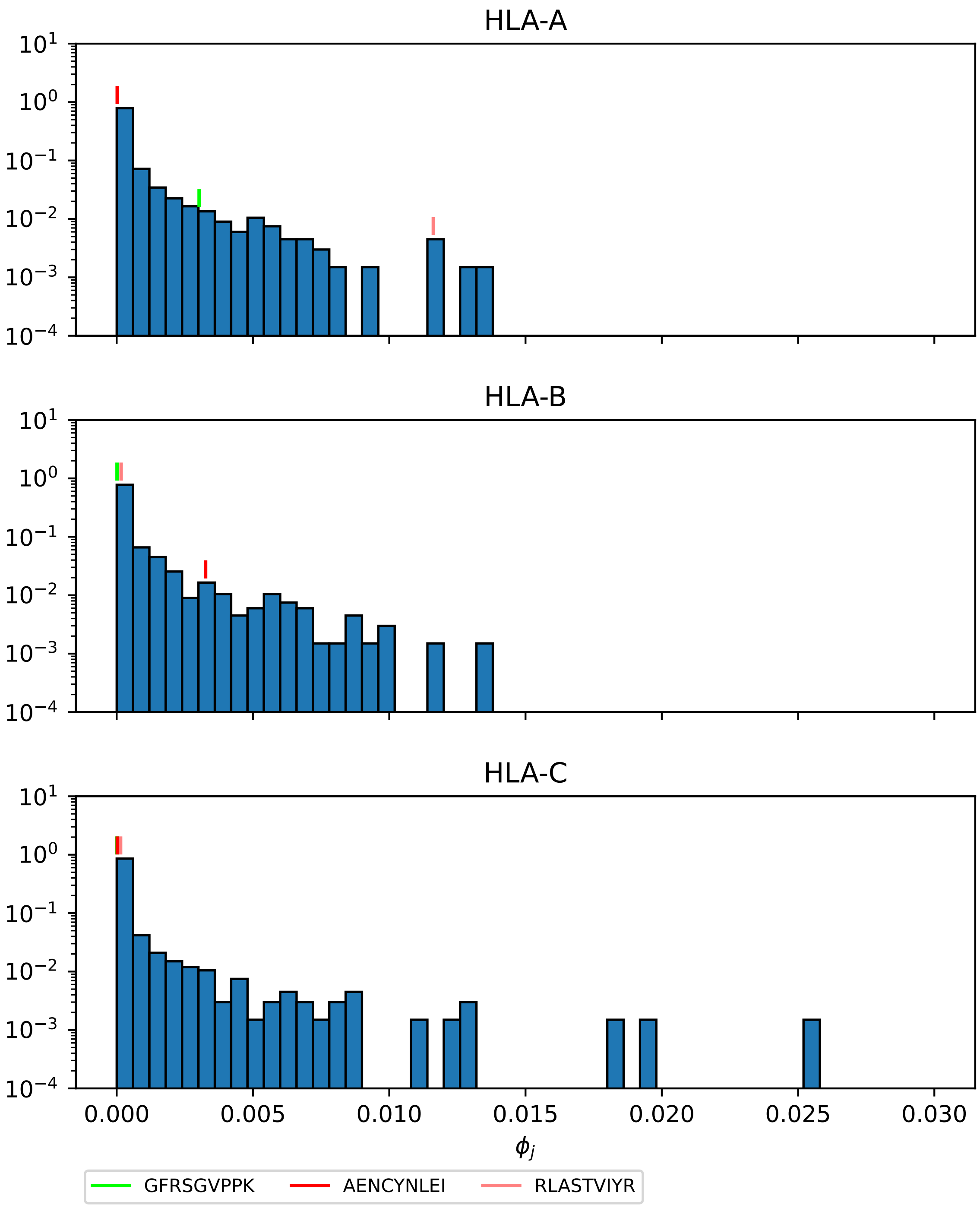


**Figure S4.** North America, Ebola GP (Sudan).

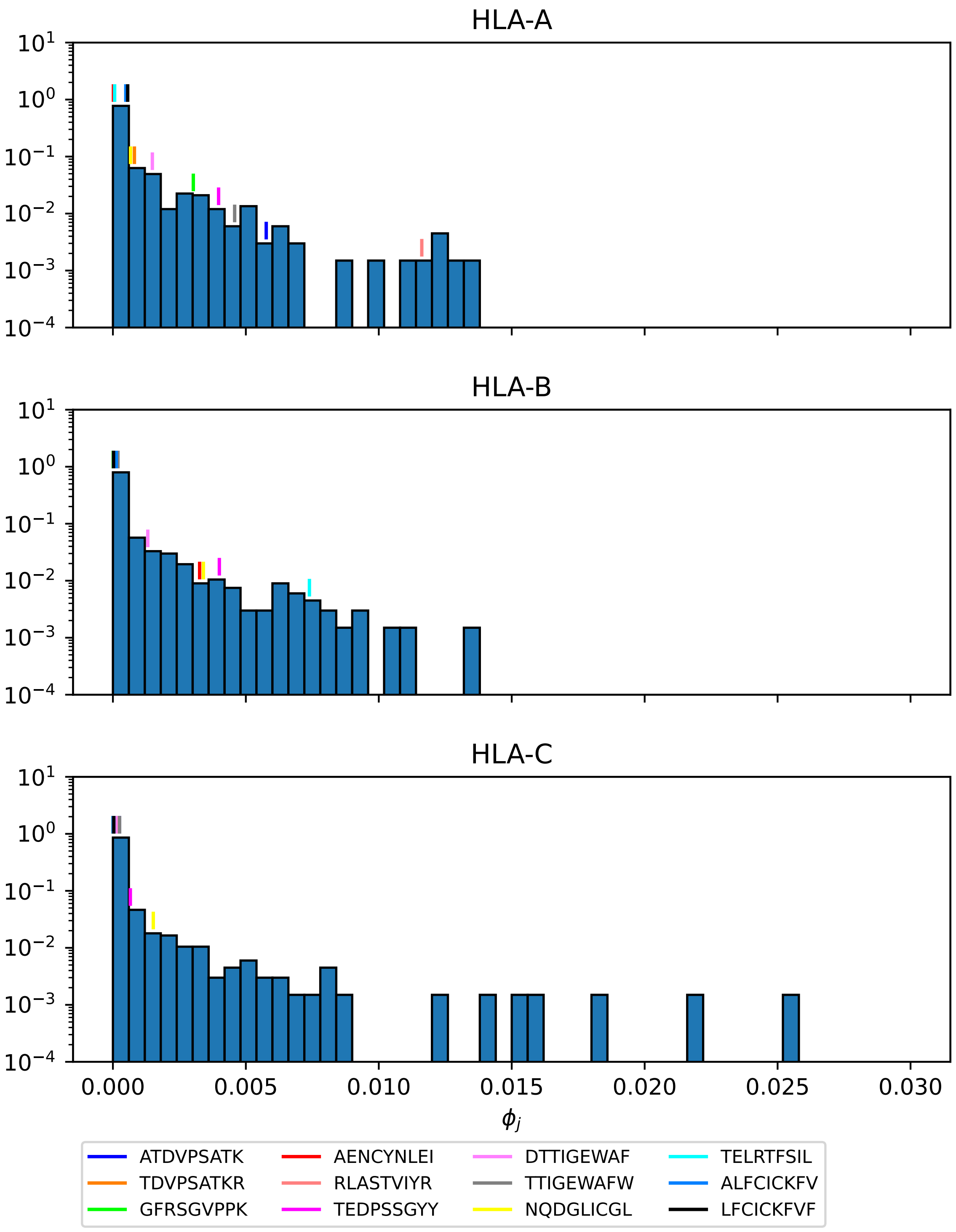


**Figure S5.** North America, Ebola GP (Zaire).

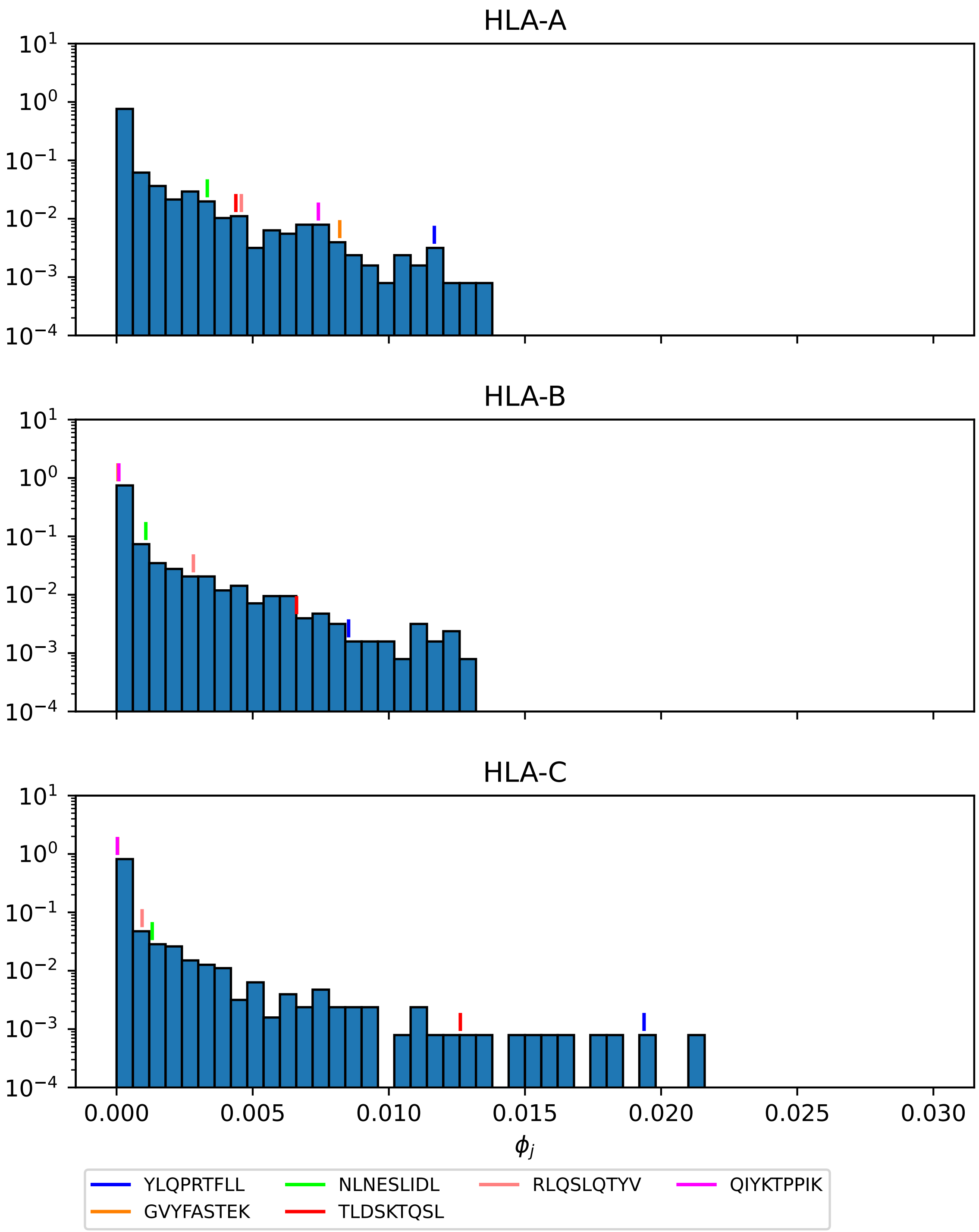


**Figure S6.** North America, SARS-CoV-2 Wuhan-Hu-1.

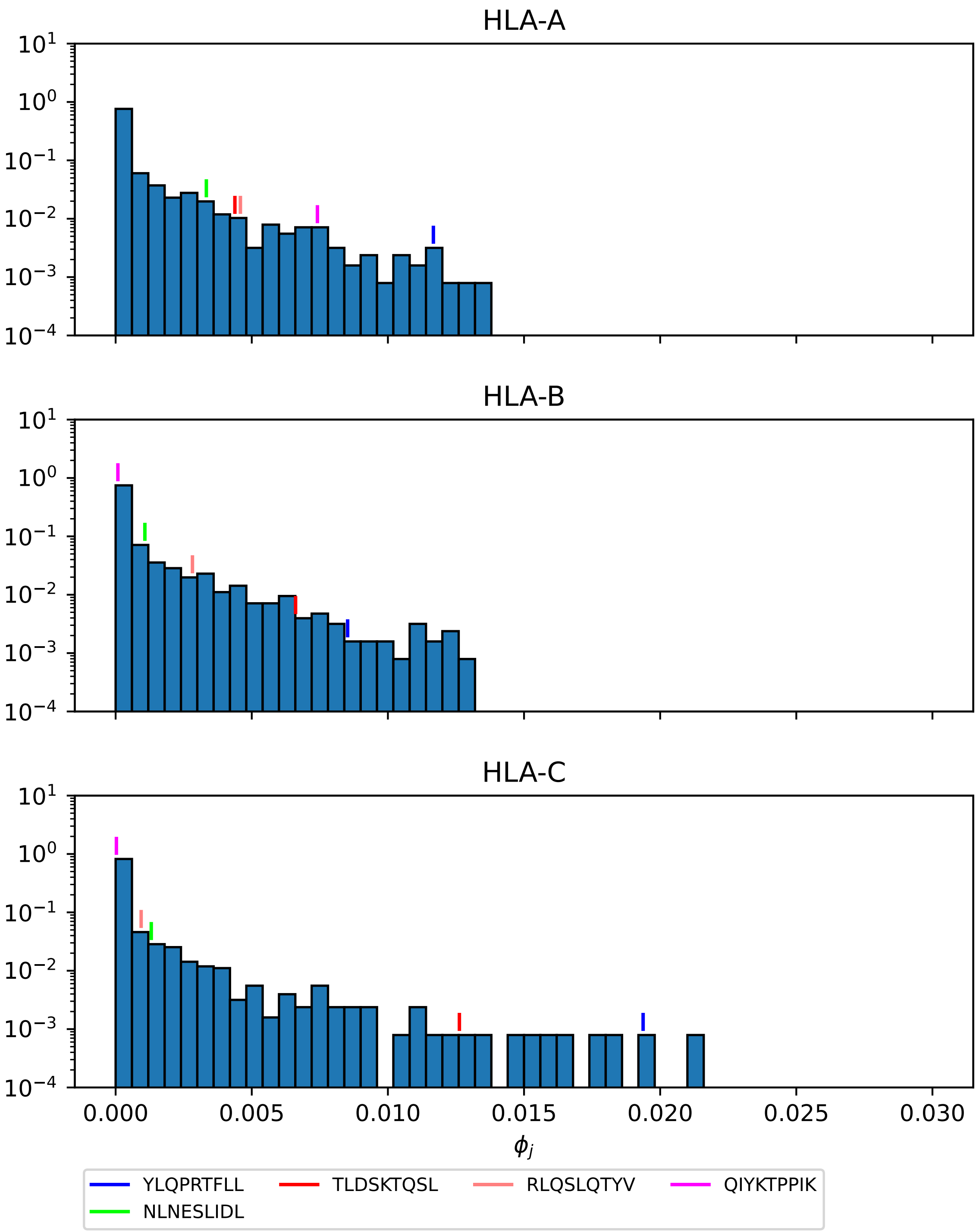


**Figure S7.** North America, SARS-CoV-2 Delta AY.4.

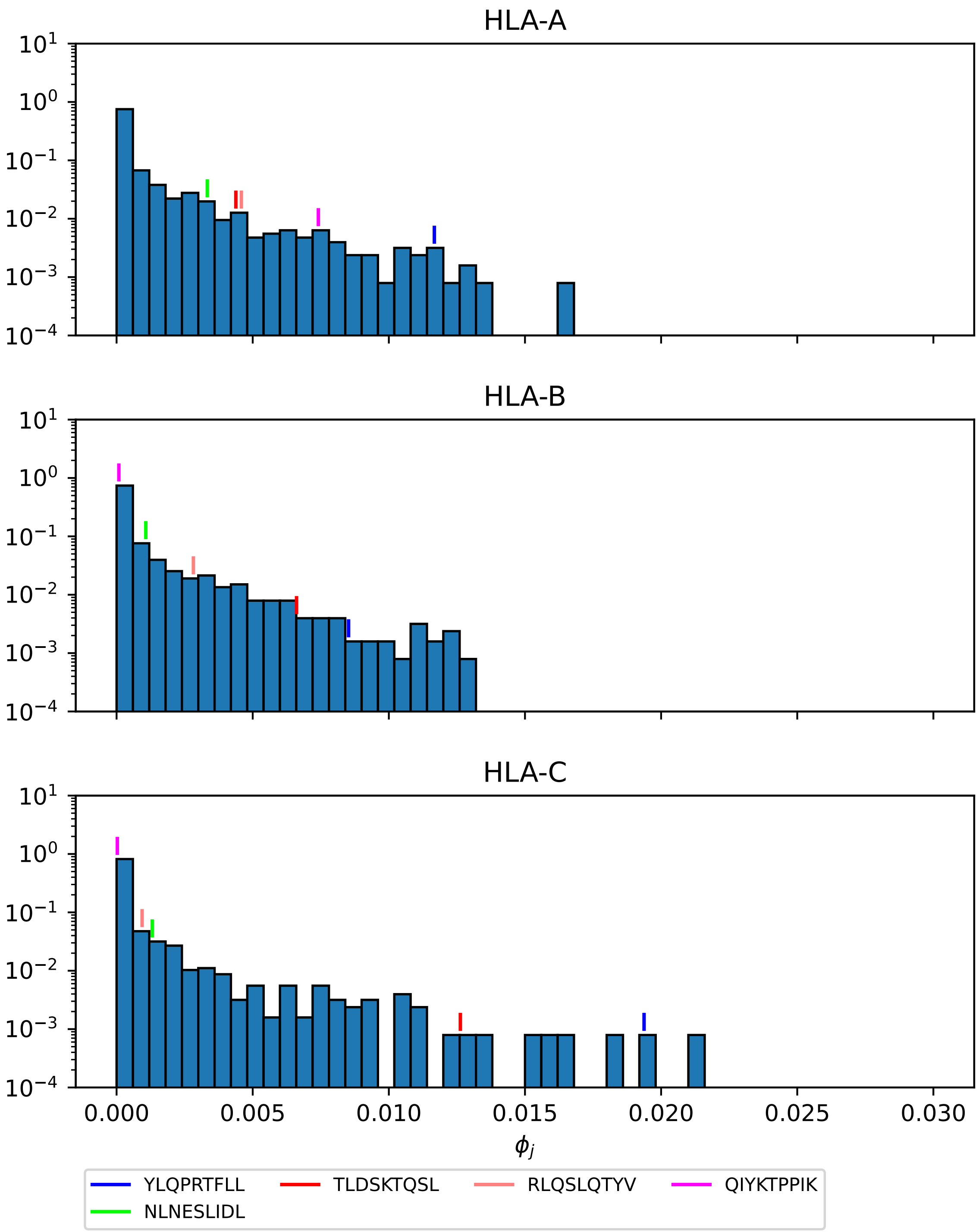


**Figure S8.** North America, SARS-CoV-2 Omicron BA.1.

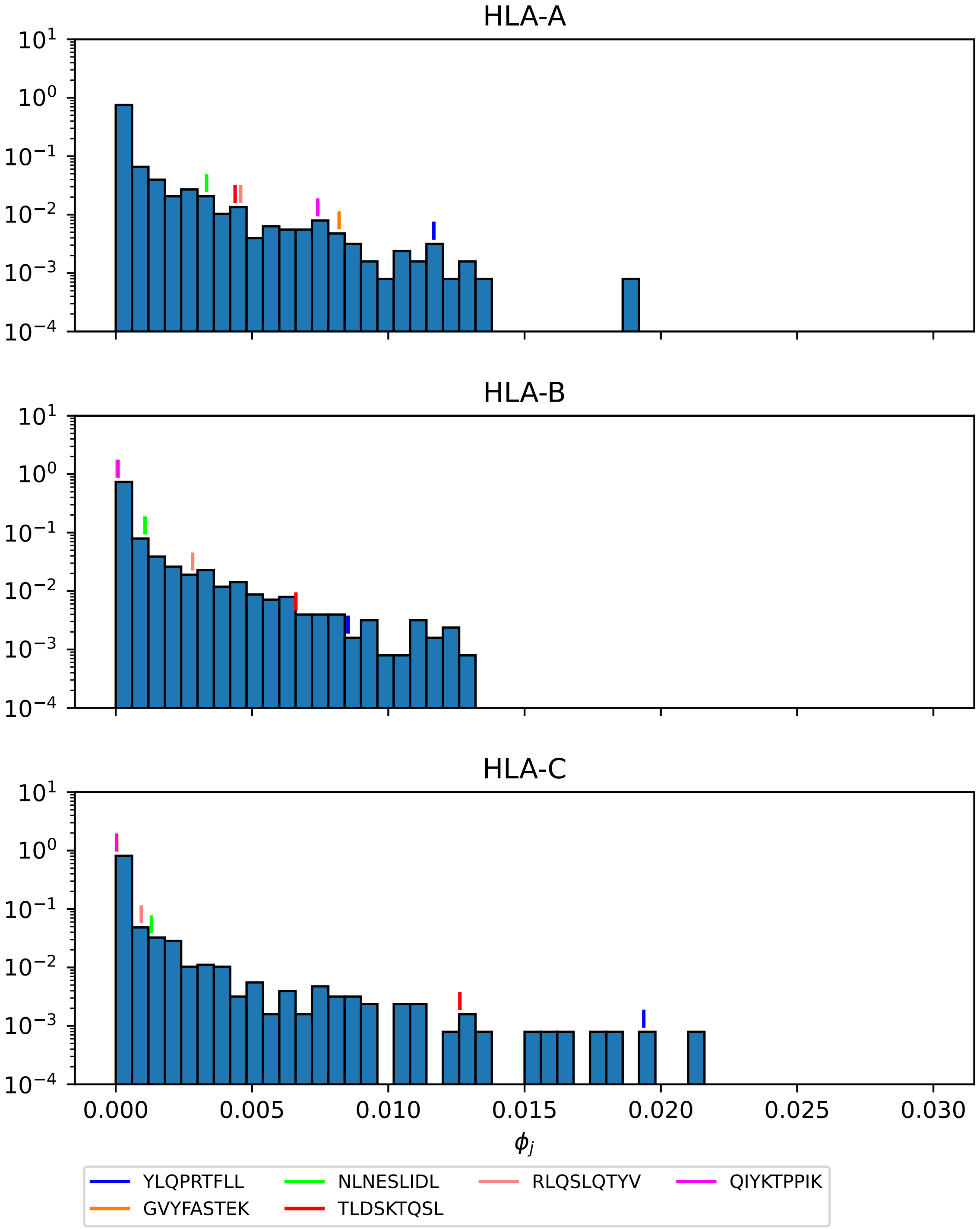


**Figure S9.** North America, SARS-CoV-2 Omicron BA.2.

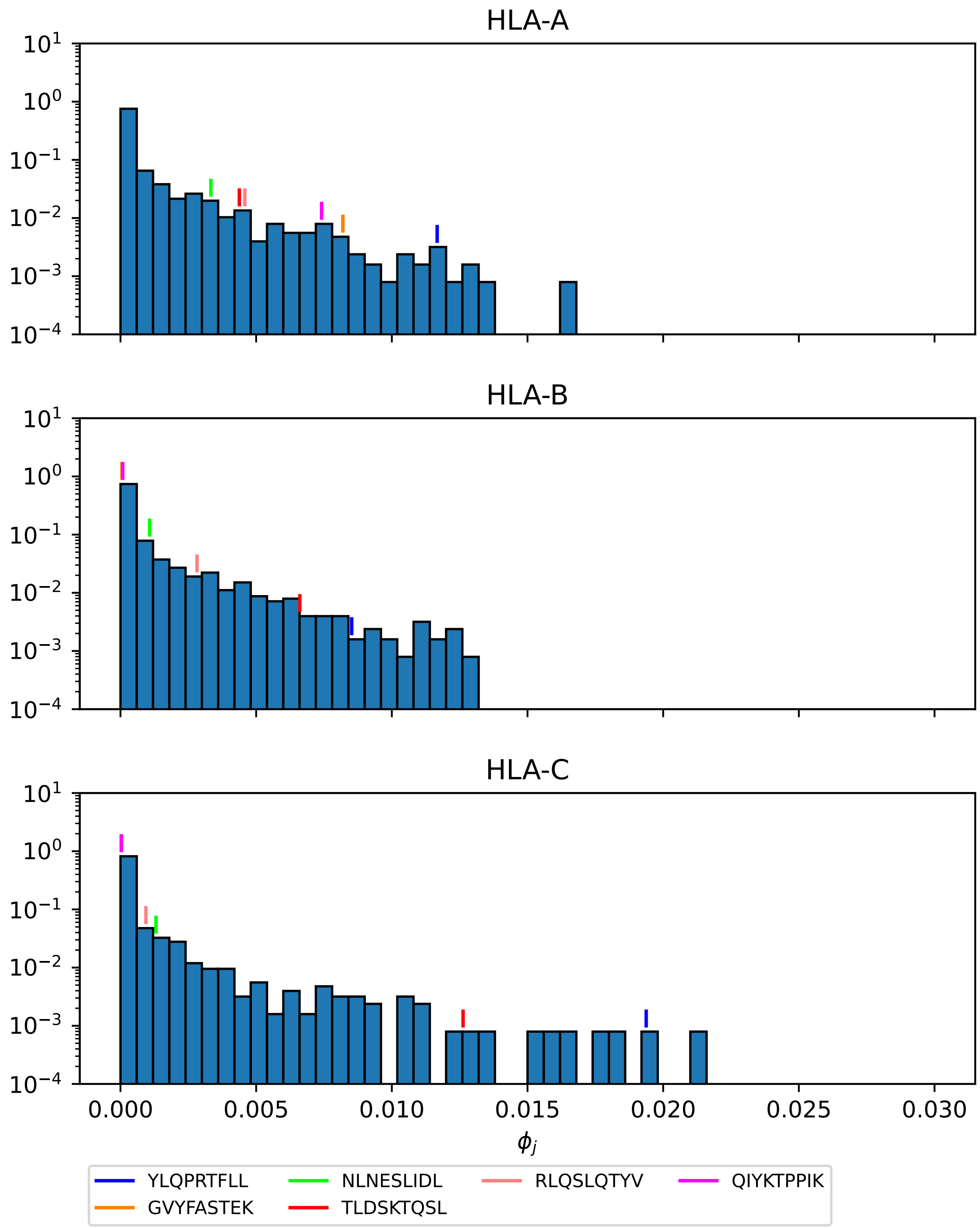


**Figure S10.** North America, SARS-CoV-2 Omicron BA.5.

#### 2 DISSECTING MEAN REGIONAL COVERAGE METRIC: ALL REGIONS

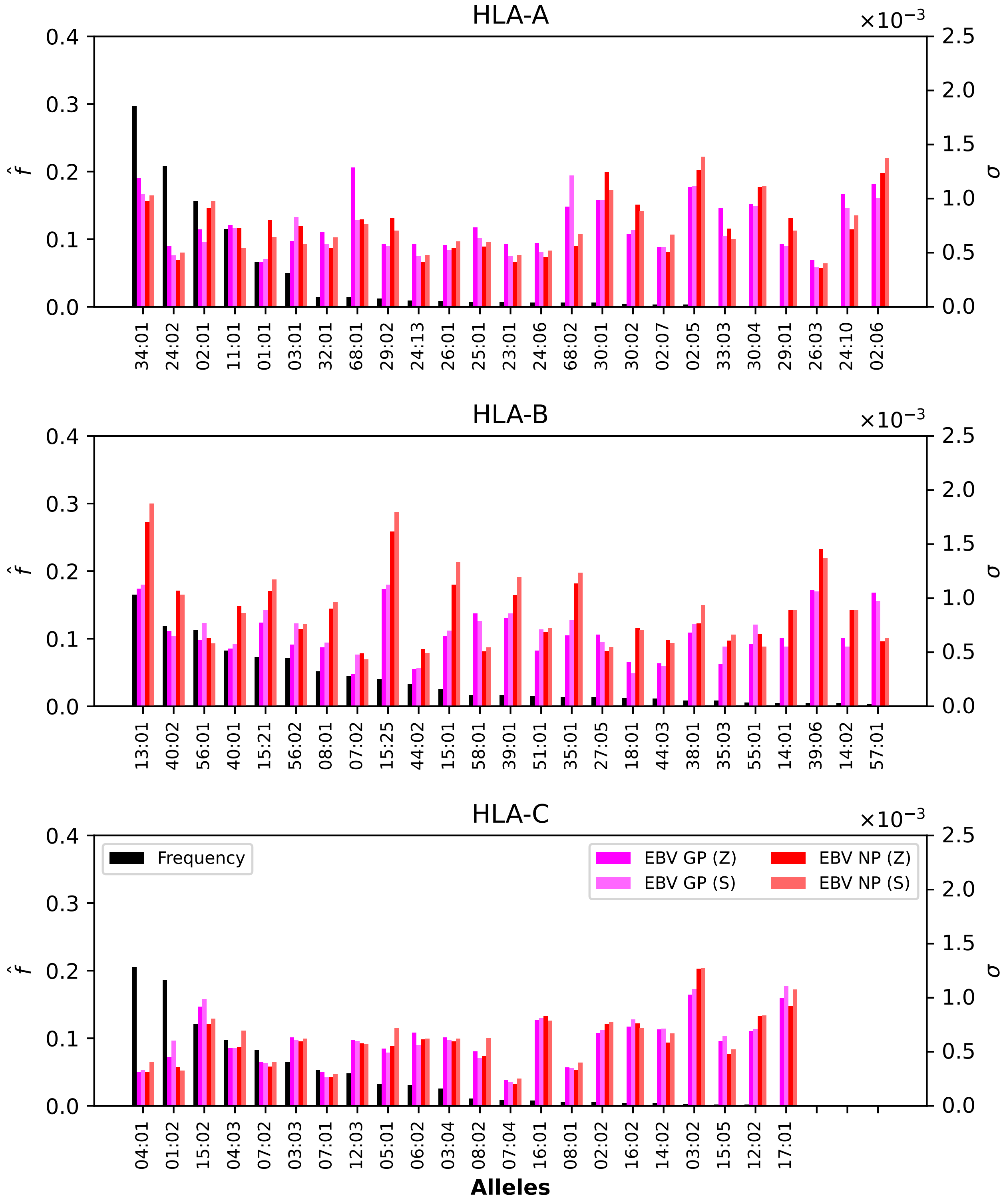


**Figure S11.** Normalized regional frequencies ($\hat{f}_i^{(1)}$) and Ebola $\sigma_i$ values for the top 25 (22 for HLA-C) most frequent alleles of each type in Australia. The top panel represents HLA-A alleles, the middle HLA-B, and the bottom HLA-C. From left to right, the bars in each group represent frequency, Ebola GP1 (Zaire), Ebola GP1 (Sudan), Ebola NP (Zaire), and Ebola NP (Sudan).

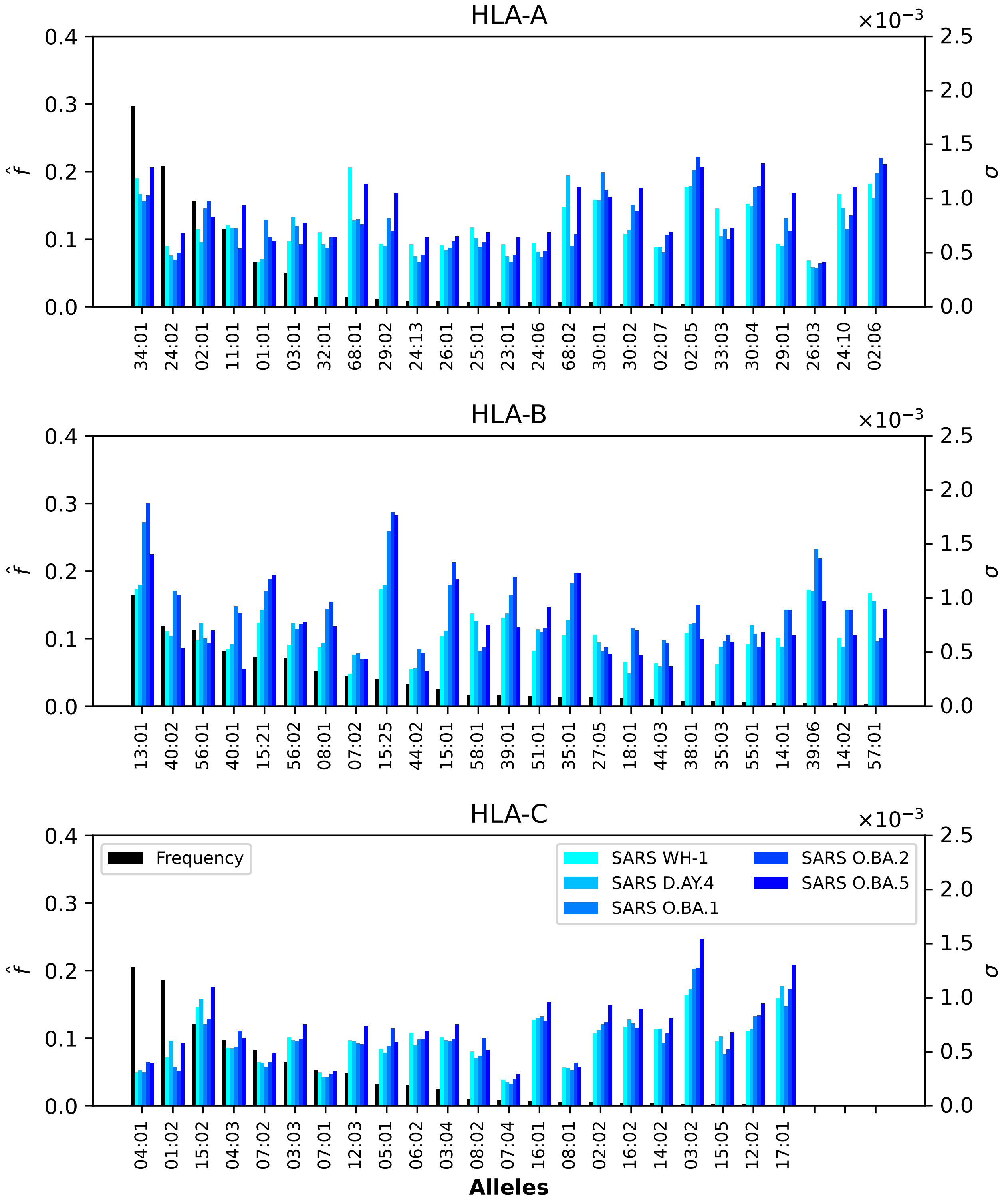


**Figure S12.** Normalized regional frequencies ($\hat{f}_i^{(1)}$) and SARS-CoV-2 $\sigma_i$ values for the top 25 (22 for HLA-C) most frequent alleles of each type in Australia. The top panel represents HLA-A alleles, the middle HLA-B, and the bottom HLA-C. From left to right, the bars in each group represent frequency, SARS-CoV-2 Wuhan-Hu-1, SARS-CoV-2 Delta AY.4, SARS-CoV-2 Omicron BA.1, SARS-CoV-2 Omicron BA.2, SARS-CoV-2 Omicron BA.5.

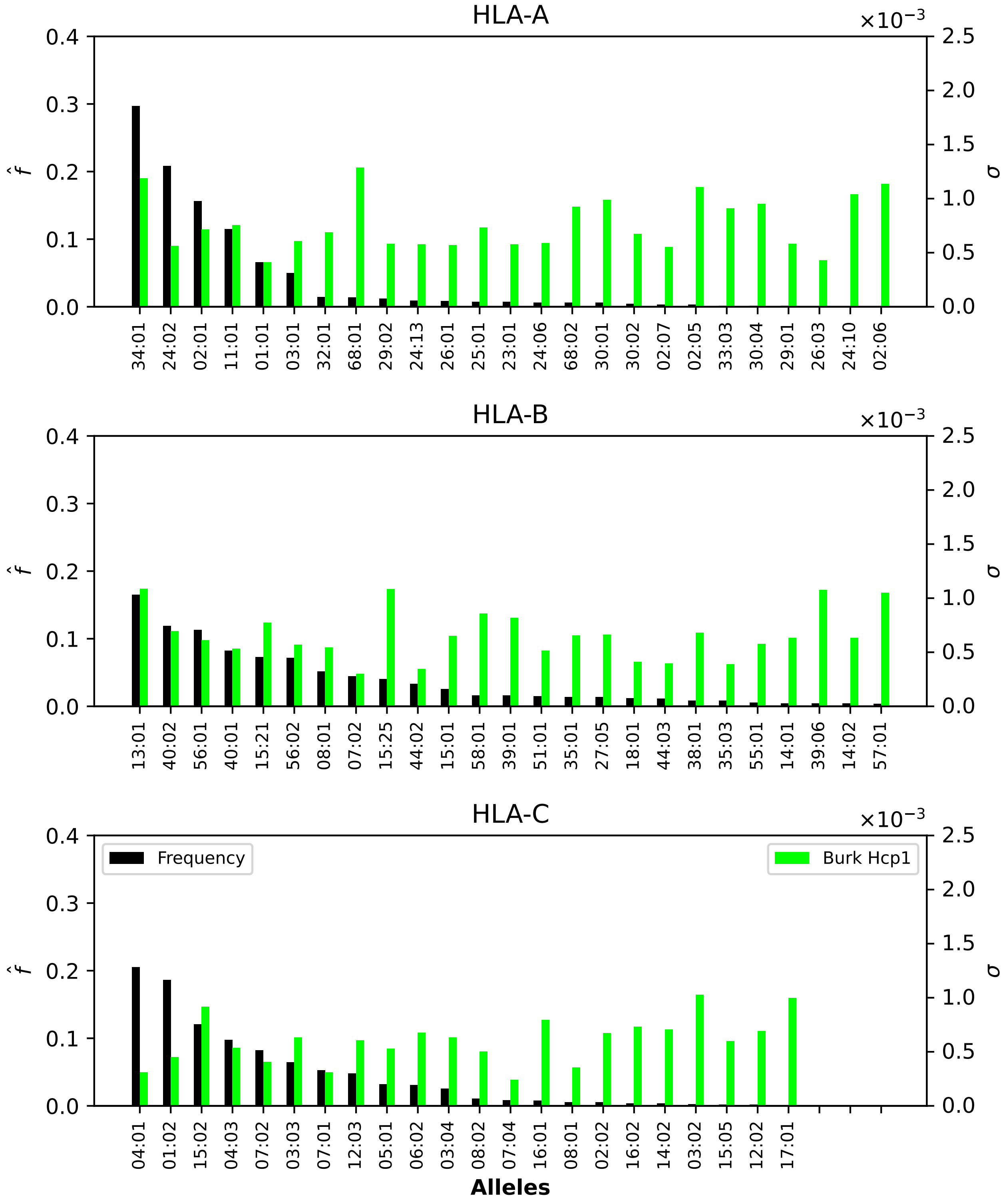


**Figure S13.** Normalized regional frequencies ($\hat{f}_i^{(1)}$) and Burkholderia $\sigma_i$ values for the top 25 (22 for HLA-C) most frequent alleles of each type in Australia. The top panel represents HLA-A alleles, the middle HLA-B, and the bottom HLA-C. From left to right, the bars in each group represent frequency and Burkholderia HCP1.

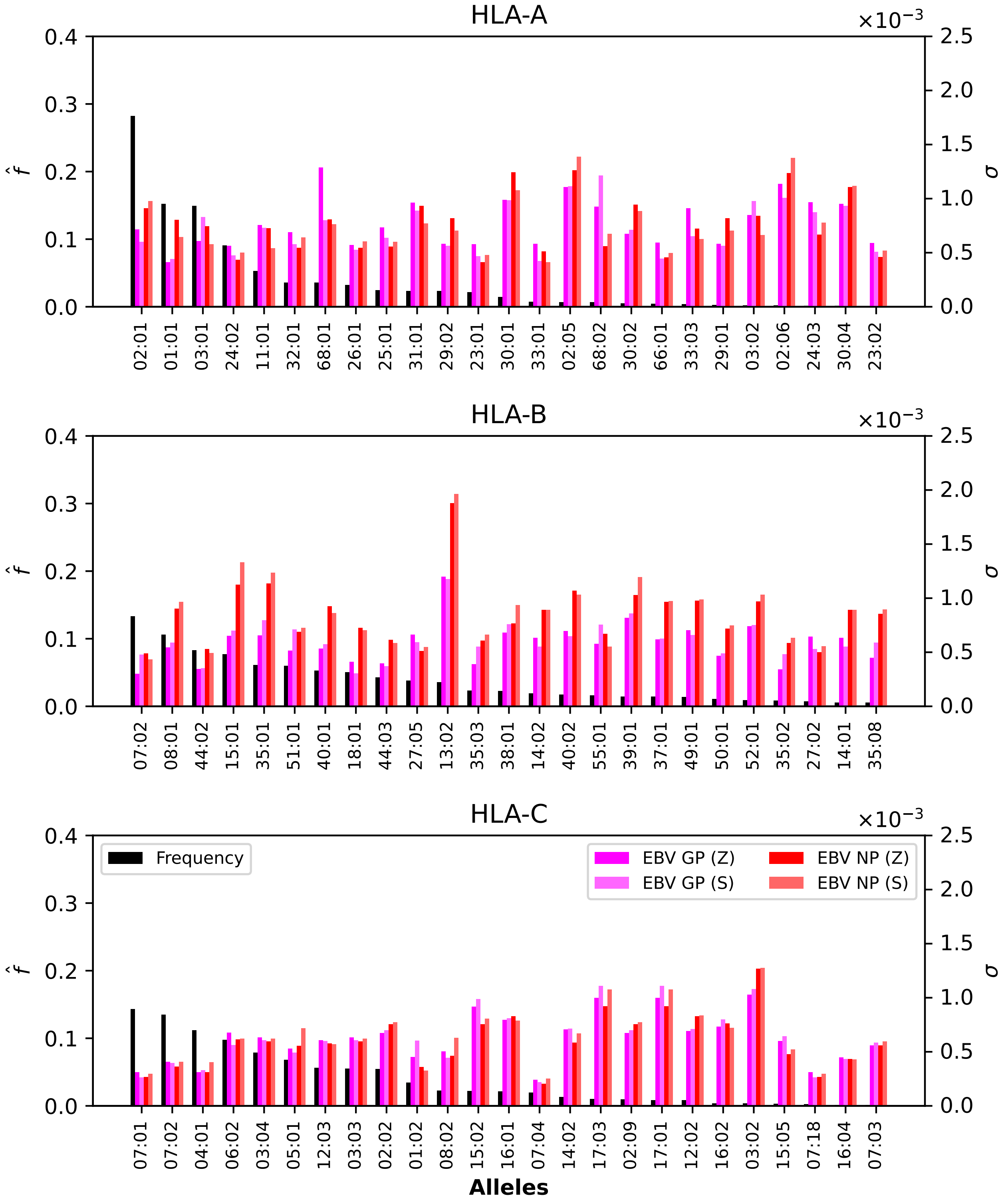


**Figure S14.** Normalized regional frequencies ($\hat{f}_i^{(2)}$) and Ebola $\sigma_i$ values for the top 25 most frequent alleles of each type in Europe. The top panel represents HLA-A alleles, the middle HLA-B, and the bottom HLA-C. From left to right, the bars in each group represent frequency, Ebola GP1 (Zaire), Ebola GP1 (Sudan), Ebola NP (Zaire), and Ebola NP (Sudan).

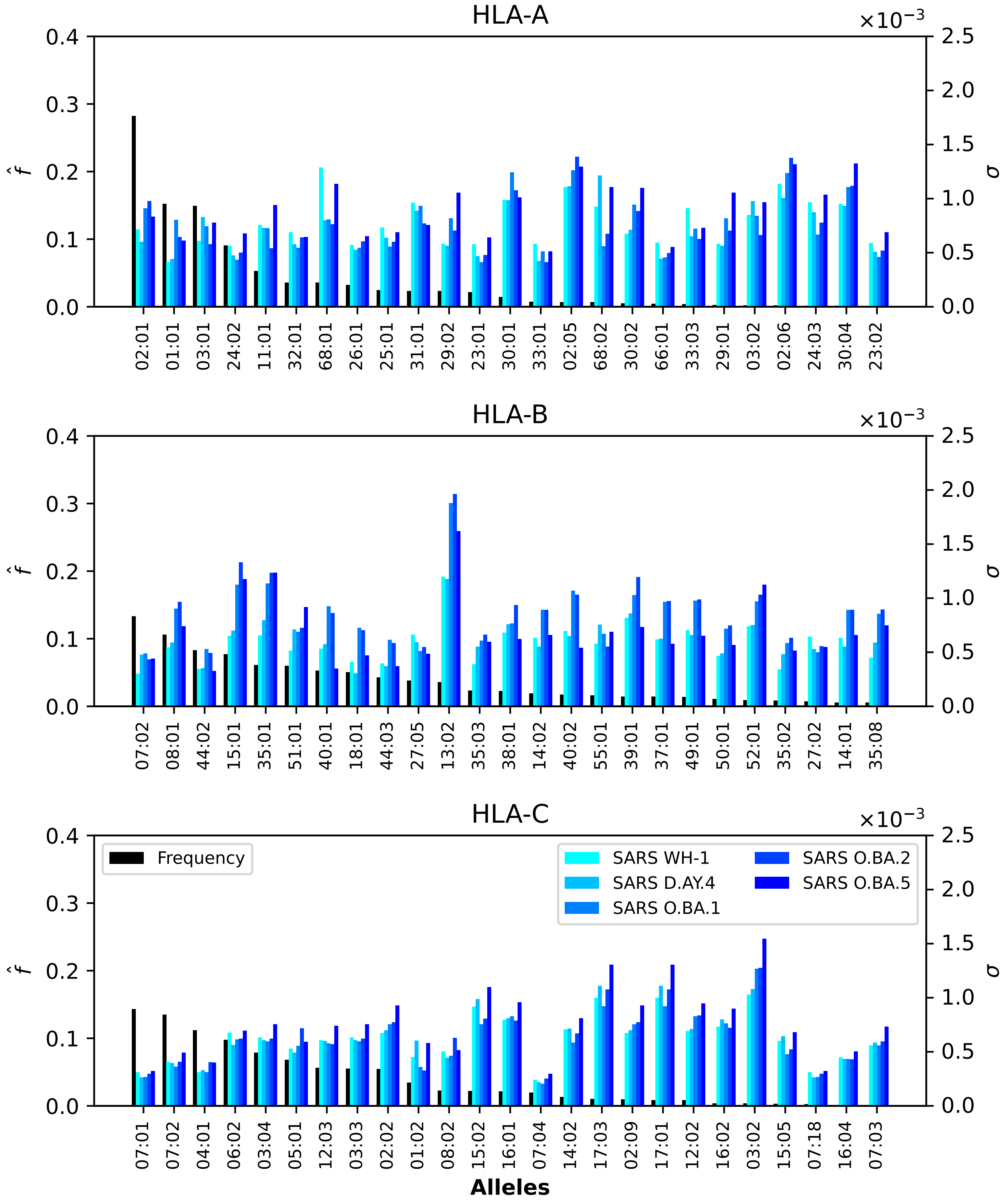


**Figure S15.** Normalized regional frequencies ($\hat{f}_i^{(2)}$) and SARS-CoV-2 $\sigma_i$ values for the top 25 most frequent alleles of each type in Europe. The top panel represents HLA-A alleles, the middle HLA-B, and the bottom HLA-C. From left to right, the bars in each group represent frequency, SARS-CoV-2 Wuhan-Hu-1, SARS-CoV-2 Delta AY.4, SARS-CoV-2 Omicron BA.1, SARS-CoV-2 Omicron BA.2, SARS-CoV-2 Omicron BA.5.

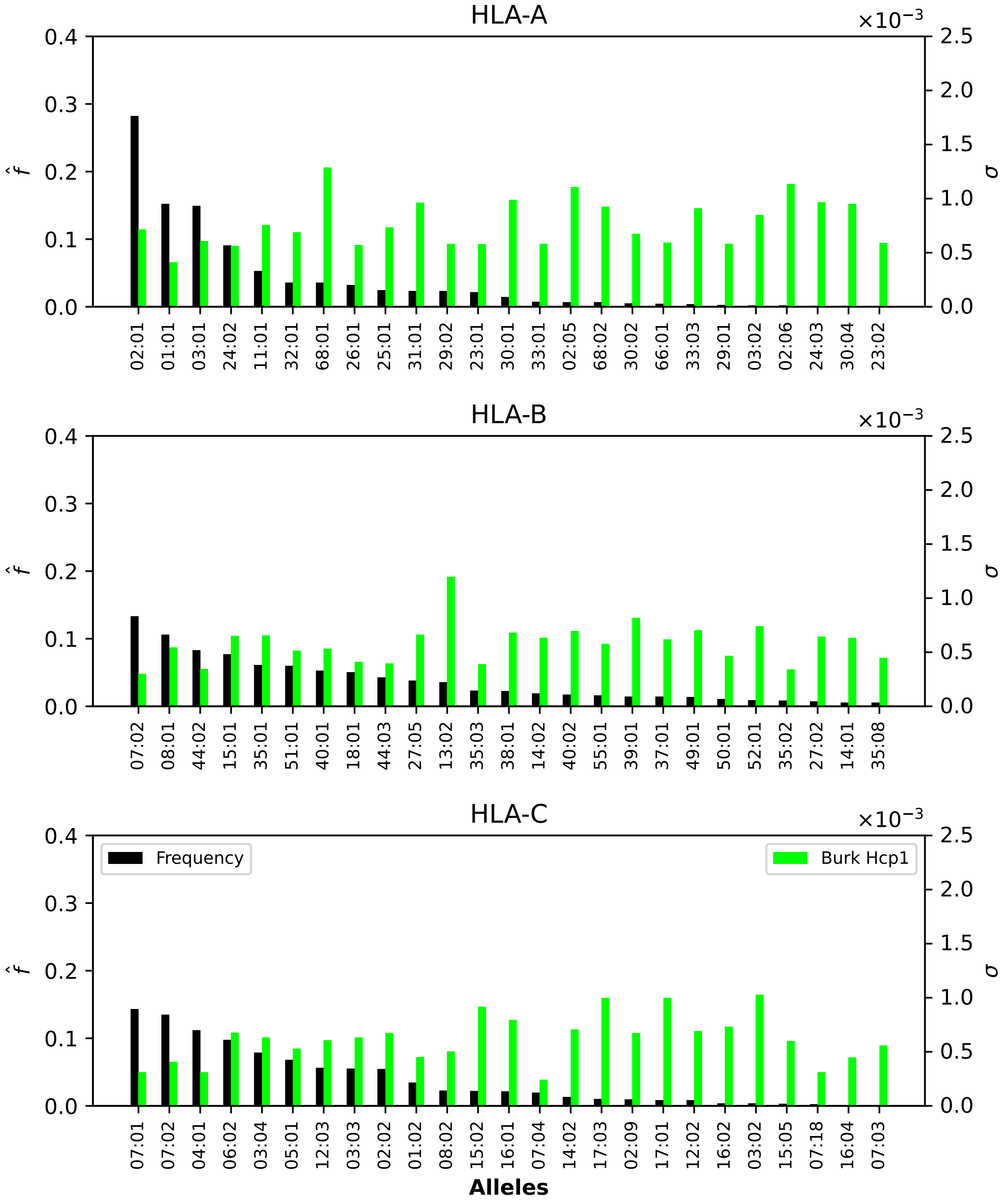


**Figure S16.** Normalized regional frequencies ($\hat{f}_i^{(2)}$) and Burkholderia $\sigma_i$ values for the top 25 most frequent alleles of each type in Europe. The top panel represents HLA-A alleles, the middle HLA-B, and the bottom HLA-C. From left to right, the bars in each group represent frequency and Burkholderia HCP1.

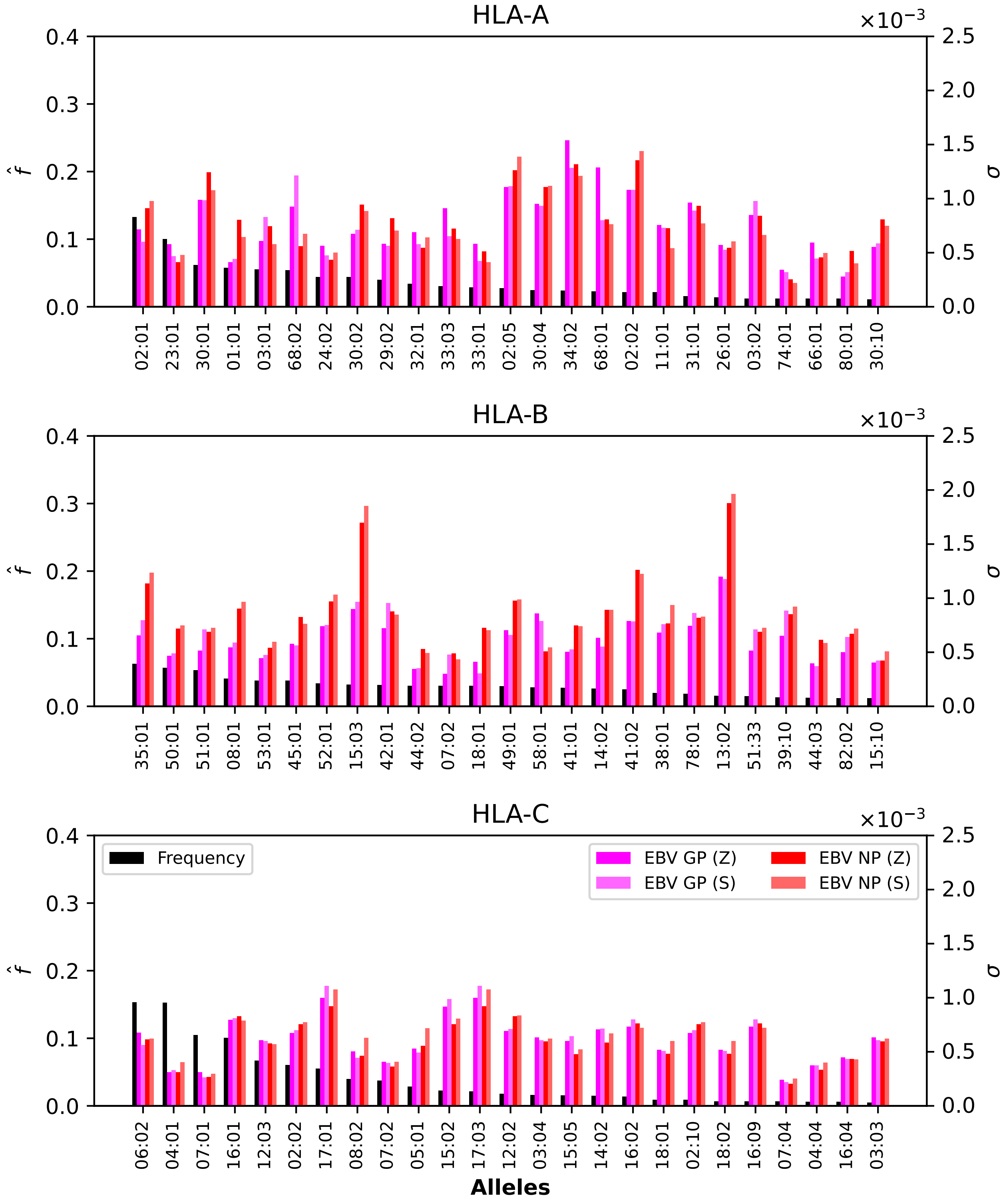


**Figure S17.** Normalized regional frequencies ($\hat{f}_i^{(3)}$) and Ebola $\sigma_i$ values for the top 25 most frequent alleles of each type in North Africa. The top panel represents HLA-A alleles, the middle HLA-B, and the bottom HLA-C. From left to right, the bars in each group represent frequency, Ebola GP1 (Zaire), Ebola GP1 (Sudan), Ebola NP (Zaire), and Ebola NP (Sudan).

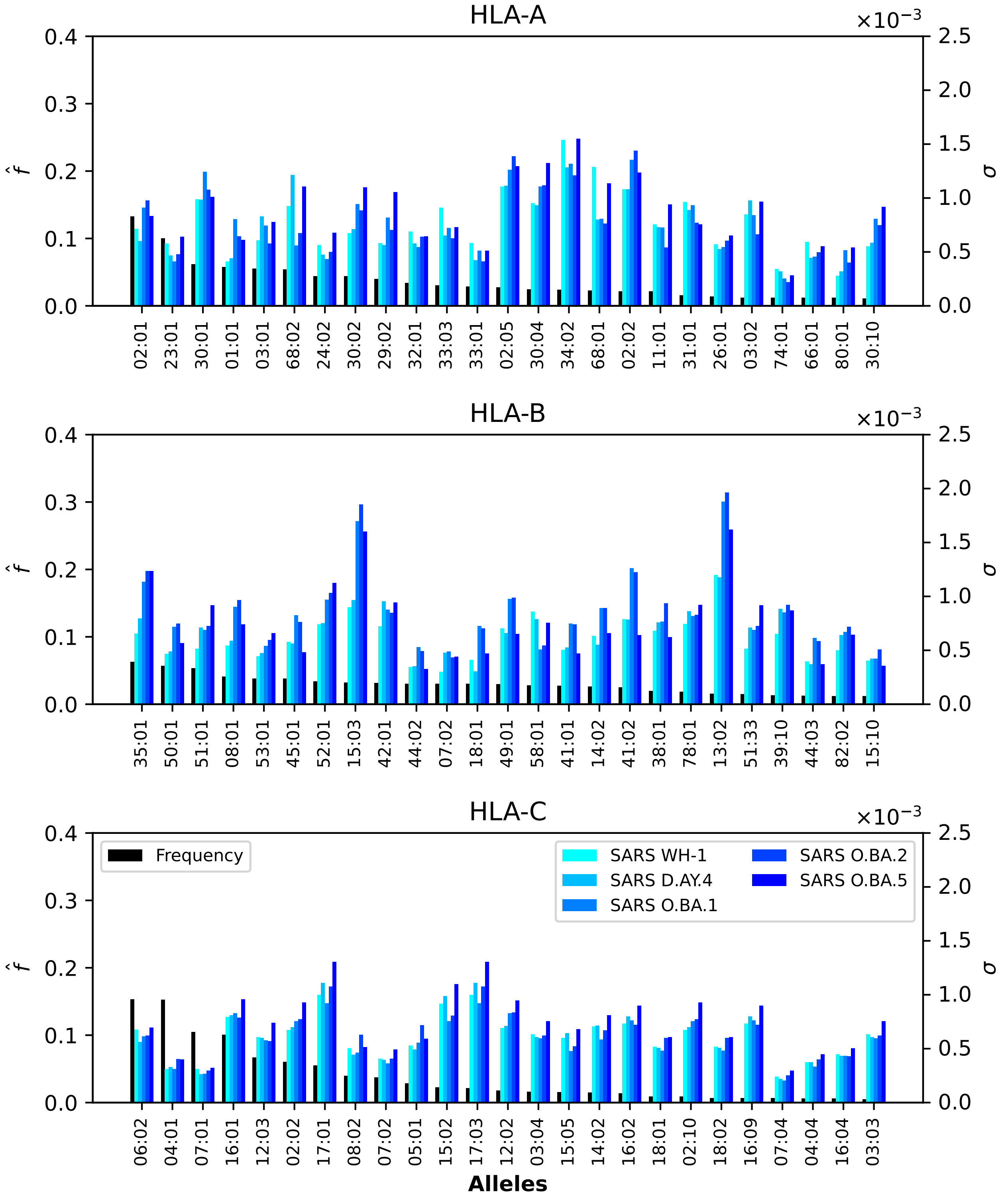


**Figure S18.** Normalized regional frequencies ($\hat{f}_i^{(3)}$) and SARS-CoV-2 $\sigma_i$ values for the top 25 most frequent alleles of each type in North Africa. The top panel represents HLA-A alleles, the middle HLA-B, and the bottom HLA-C. From left to right, the bars in each group represent frequency, SARS-CoV-2 Wuhan-Hu-1, SARS-CoV-2 Delta AY.4, SARS-CoV-2 Omicron BA.1, SARS-CoV-2 Omicron BA.2, SARS-CoV-2 Omicron BA.5.

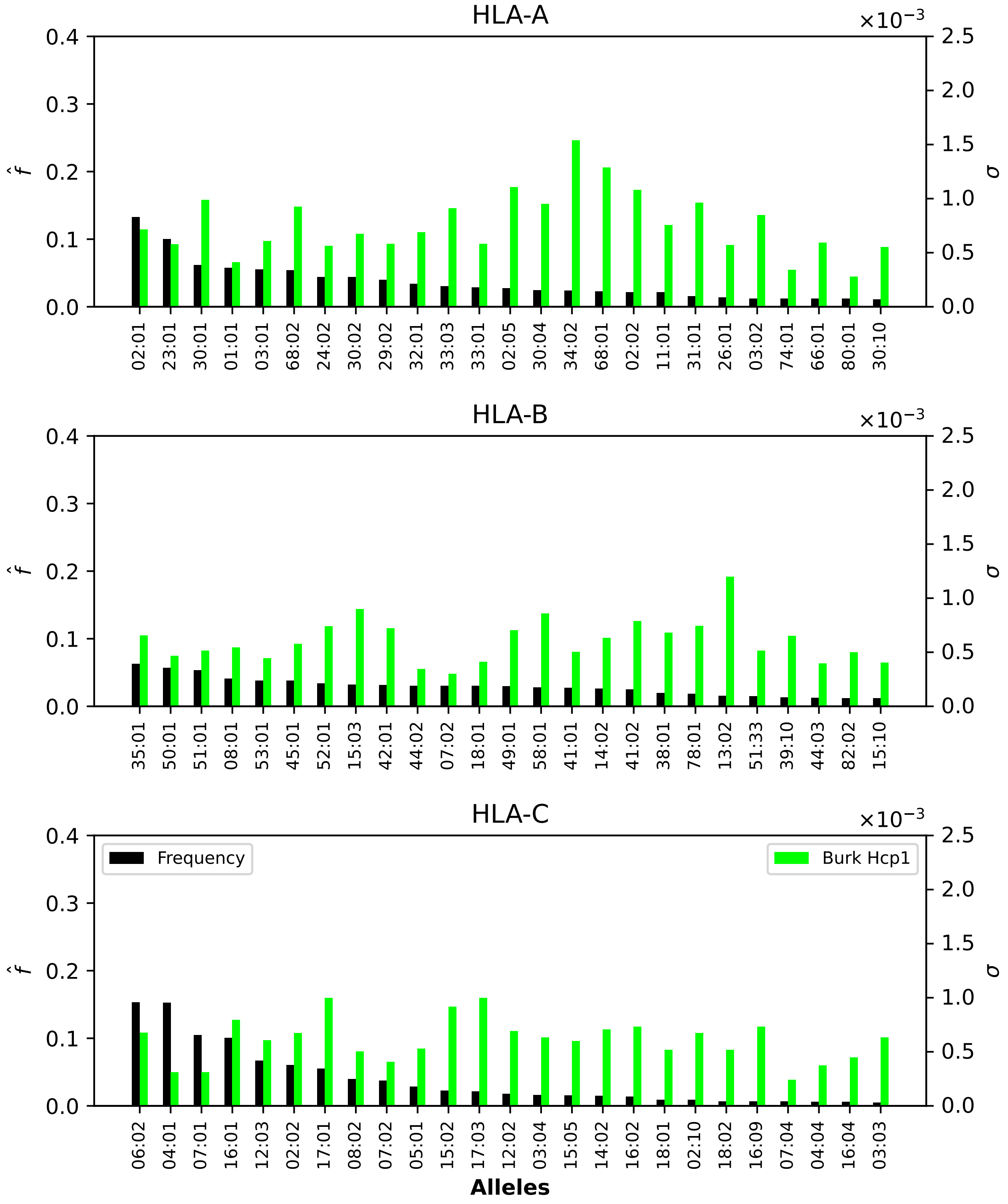


**Figure S19.** Normalized regional frequencies ($\hat{f}_i^{(3)}$) and Burkholderia $\sigma_i$ values for the top 25 most frequent alleles of each type in North Africa. The top panel represents HLA-A alleles, the middle HLA-B, and the bottom HLA-C. From left to right, the bars in each group represent frequency and Burkholderia HCP1.

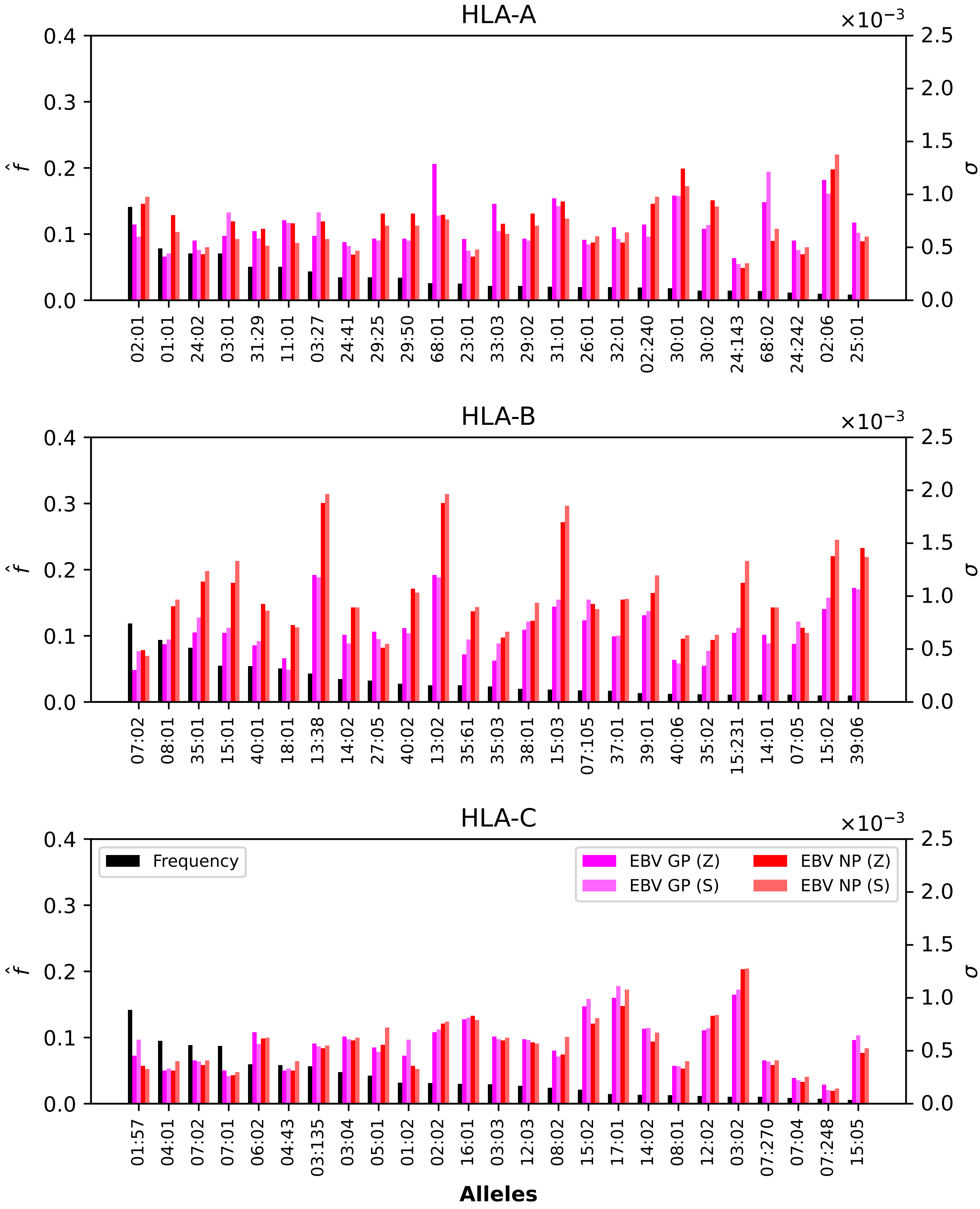


**Figure S20.** Normalized regional frequencies ($\hat{f}_i^{(4)}$) and Ebola $\sigma_i$ values for the top 25 most frequent alleles of each type in North America. The top panel represents HLA-A alleles, the middle HLA-B, and the bottom HLA-C. From left to right, the bars in each group represent frequency, Ebola GP1 (Zaire), Ebola GP1 (Sudan), Ebola NP (Zaire), and Ebola NP (Sudan).

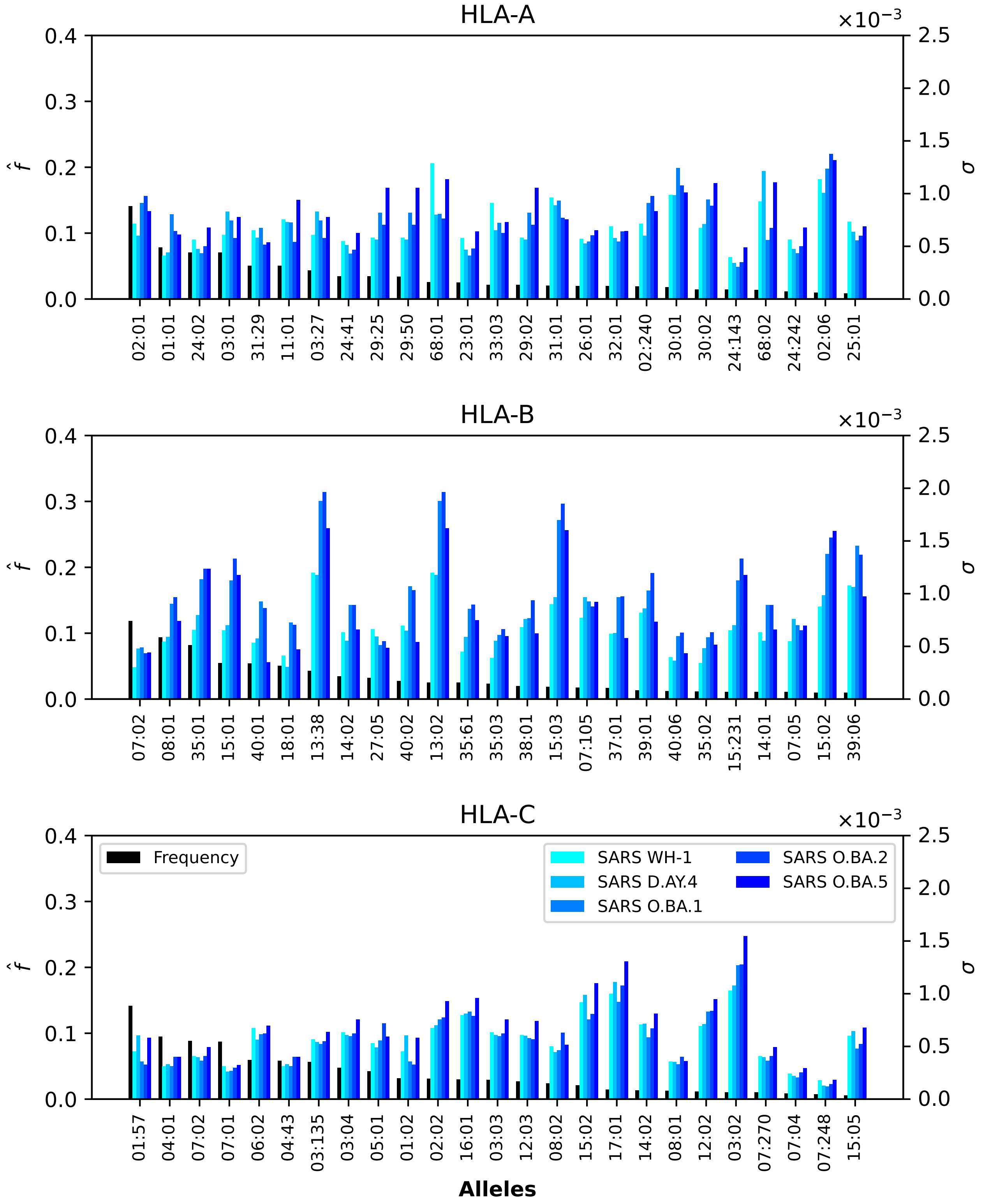


**Figure S21.** Normalized regional frequencies ($\hat{f}_i^{(4)}$) and SARS-CoV-2 $\sigma_i$ values for the top 25 most frequent alleles of each type in North America. The top panel represents HLA-A alleles, the middle HLA-B, and the bottom HLA-C. From left to right, the bars in each group represent frequency, SARS-CoV-2 Wuhan-Hu-1, SARS-CoV-2 Delta AY.4, SARS-CoV-2 Omicron BA.1, SARS-CoV-2 Omicron BA.2, SARS-CoV-2 Omicron BA.5.

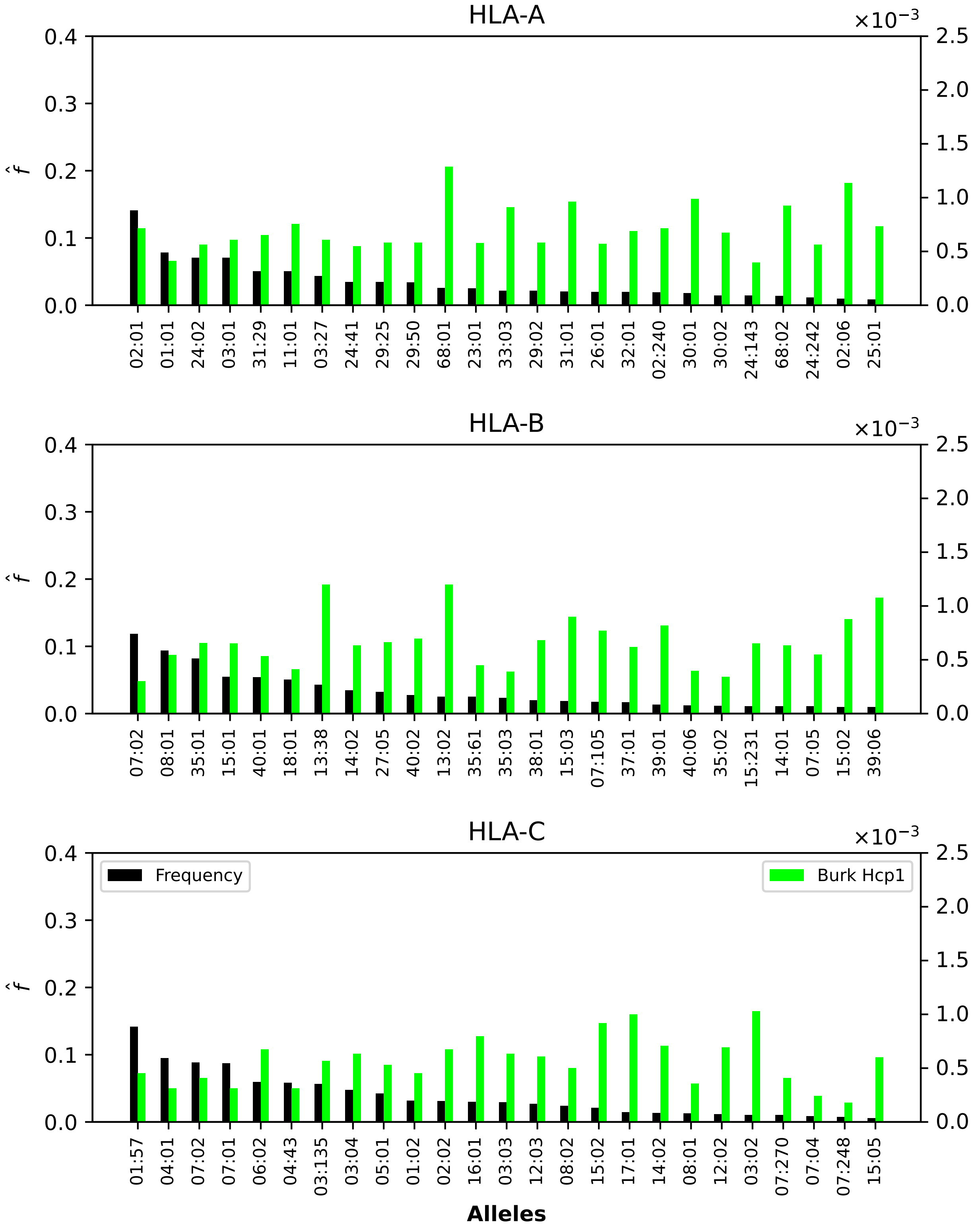


**Figure S22.** Normalized regional frequencies ($\hat{f}_i^{(4)}$) and Burkholderia $\sigma_i$ values for the top 25 most frequent alleles of each type in North America. The top panel represents HLA-A alleles, the middle HLA-B, and the bottom HLA-C. From left to right, the bars in each group represent frequency and Burkholderia HCP1.

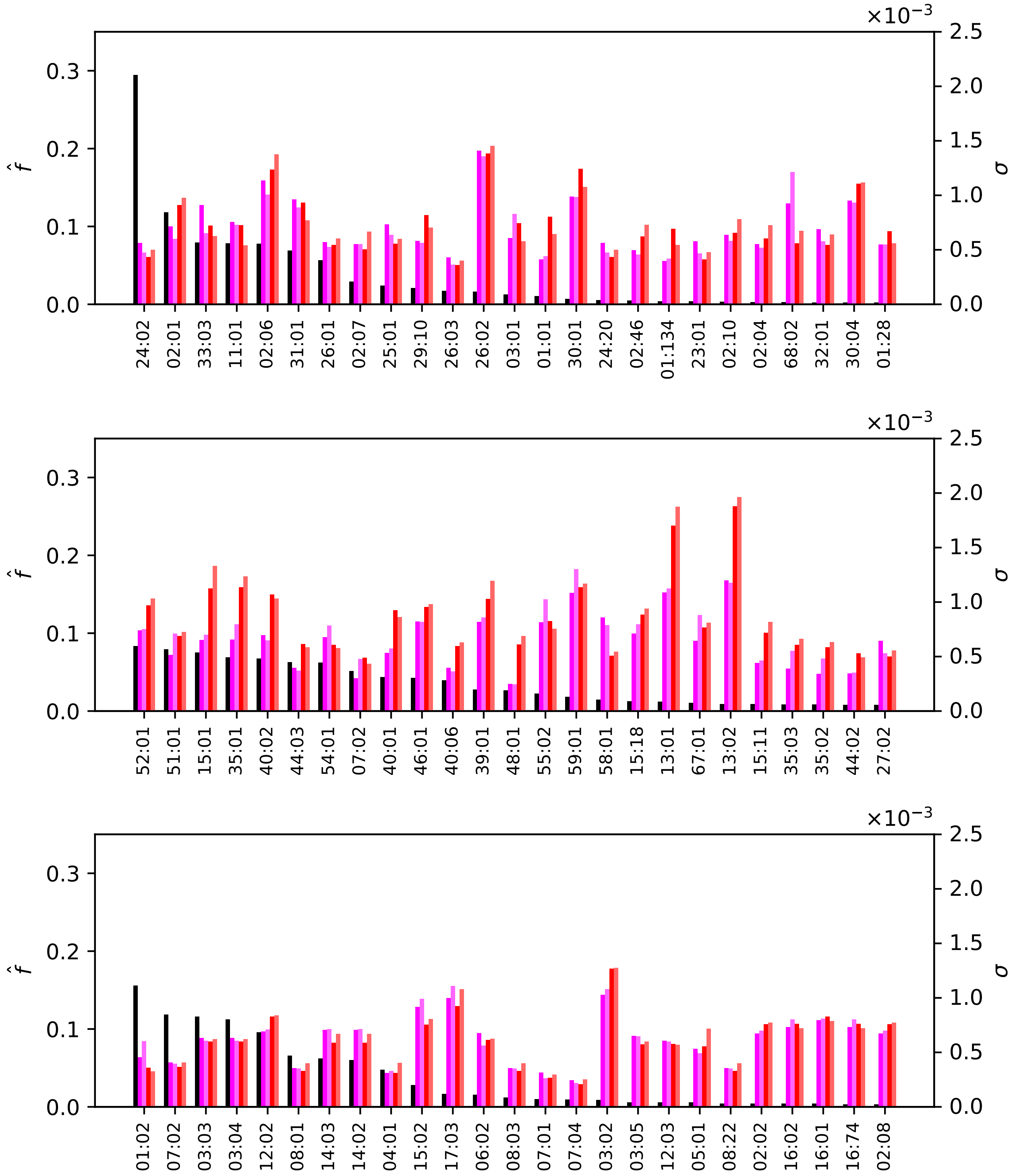


**Figure S23.** Normalized regional frequencies ($\hat{f}_i^{(5)}$) and Ebola $\sigma_i$ values for the top 25 most frequent alleles of each type in Northeast Asia. The top panel represents HLA-A alleles, the middle HLA-B, and the bottom HLA-C. From left to right, the bars in each group represent frequency, Ebola GP1 (Zaire), Ebola GP1 (Sudan), Ebola NP (Zaire), and Ebola NP (Sudan).

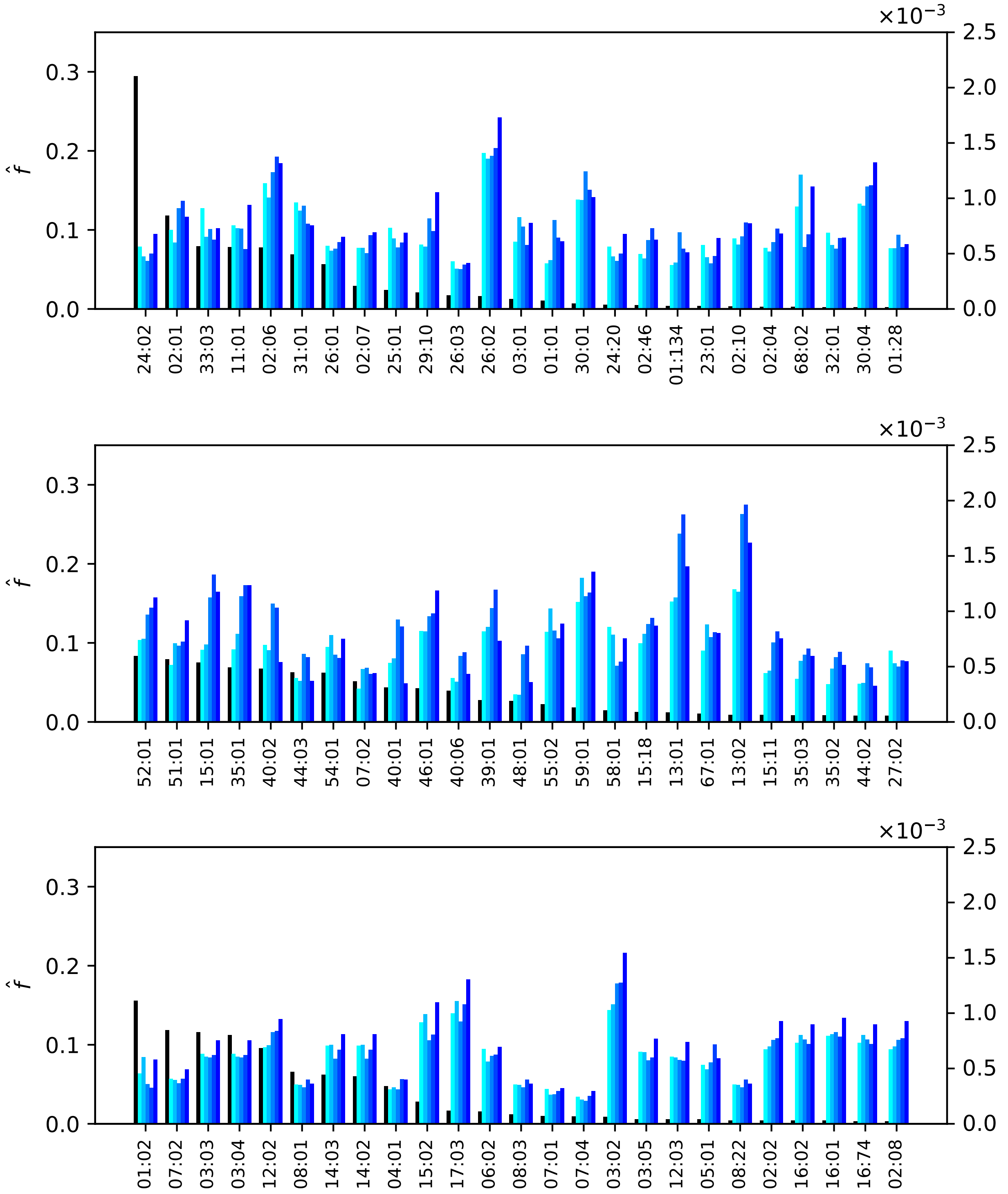


**Figure S24.** Normalized regional frequencies ($\hat{f}_i^{(5)}$) and SARS-CoV-2 $\sigma_i$ values for the top 25 most frequent alleles of each type in Northeast Asia. The top panel represents HLA-A alleles, the middle HLA-B, and the bottom HLA-C. From left to right, the bars in each group represent frequency, SARS-CoV-2 Wuhan-Hu-1, SARS-CoV-2 Delta AY.4, SARS-CoV-2 Omicron BA.1, SARS-CoV-2 Omicron BA.2, SARS-CoV-2 Omicron BA.5.

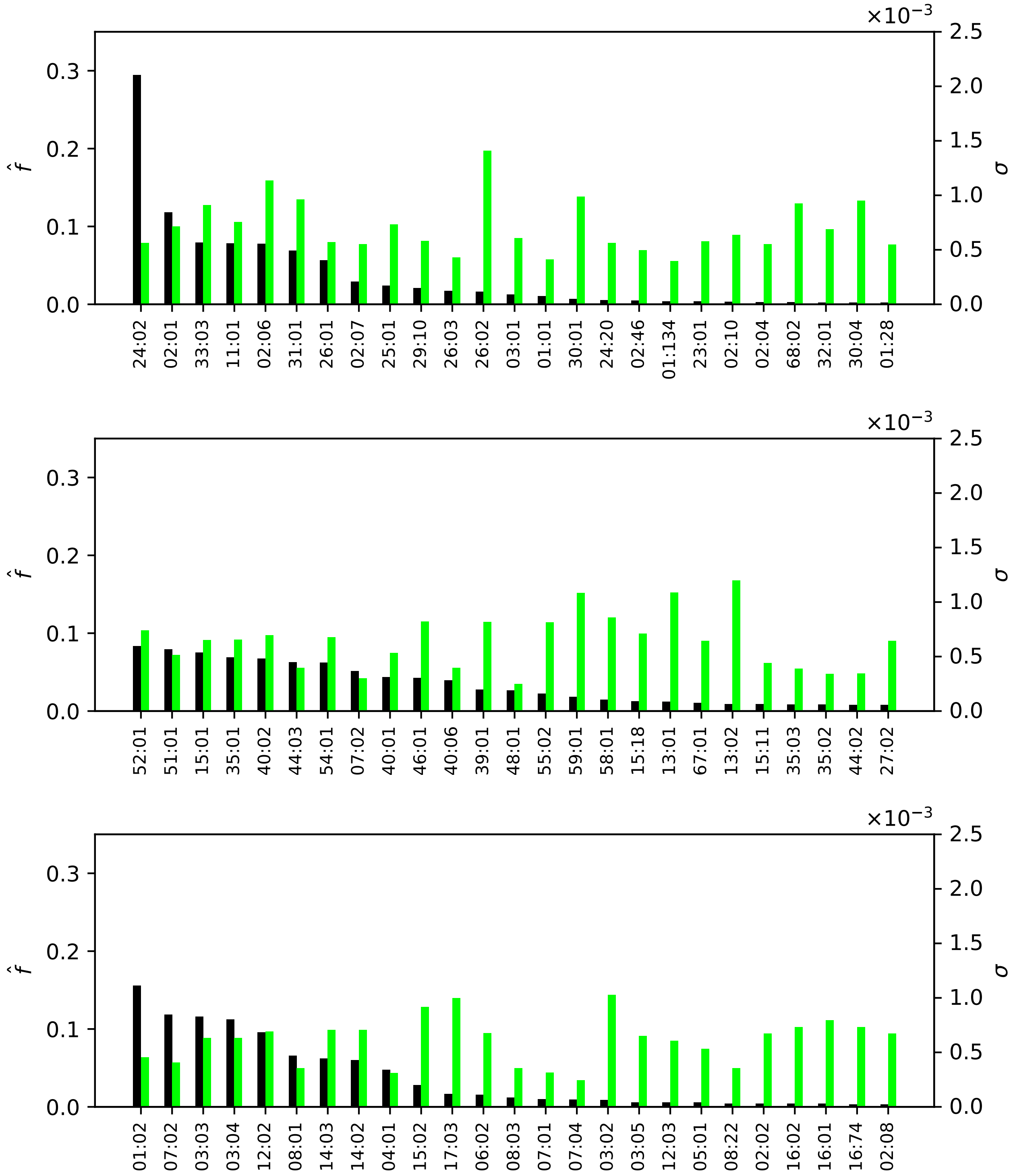


**Figure S25.** Normalized regional frequencies ($\hat{f}_i^{(5)}$) and Burkholderia $\sigma_i$ values for the top 25 most frequent alleles of each type in Northeast Asia. The top panel represents HLA-A alleles, the middle HLA-B, and the bottom HLA-C. From left to right, the bars in each group represent frequency and Burkholderia HCP1.

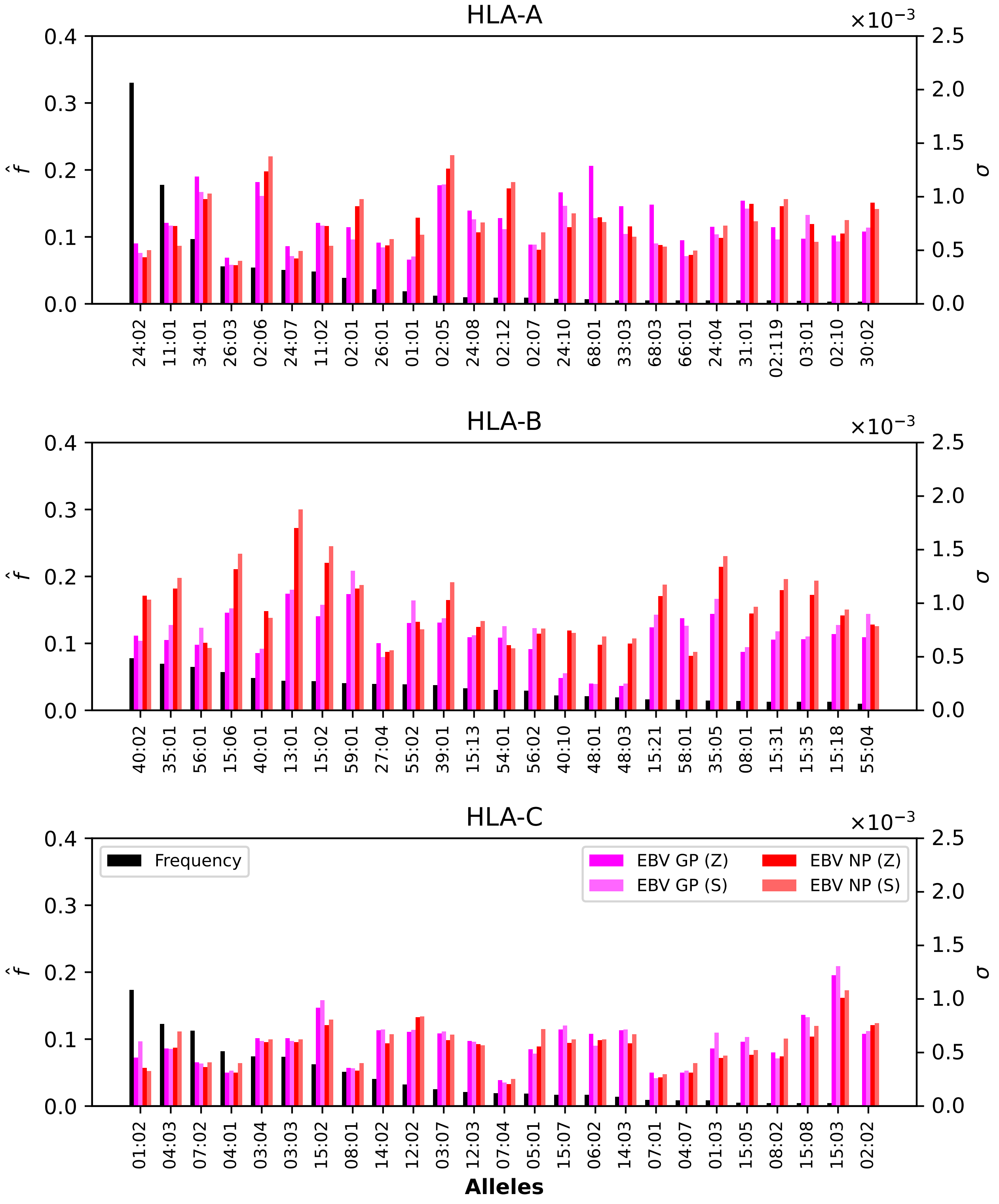


**Figure S26.** Normalized regional frequencies ($\hat{f}_i^{(6)}$) and Ebola $\sigma_i$ values for the top 25 most frequent alleles of each type in Oceania. The top panel represents HLA-A alleles, the middle HLA-B, and the bottom HLA-C. From left to right, the bars in each group represent frequency, Ebola GP1 (Zaire), Ebola GP1 (Sudan), Ebola NP (Zaire), and Ebola NP (Sudan).

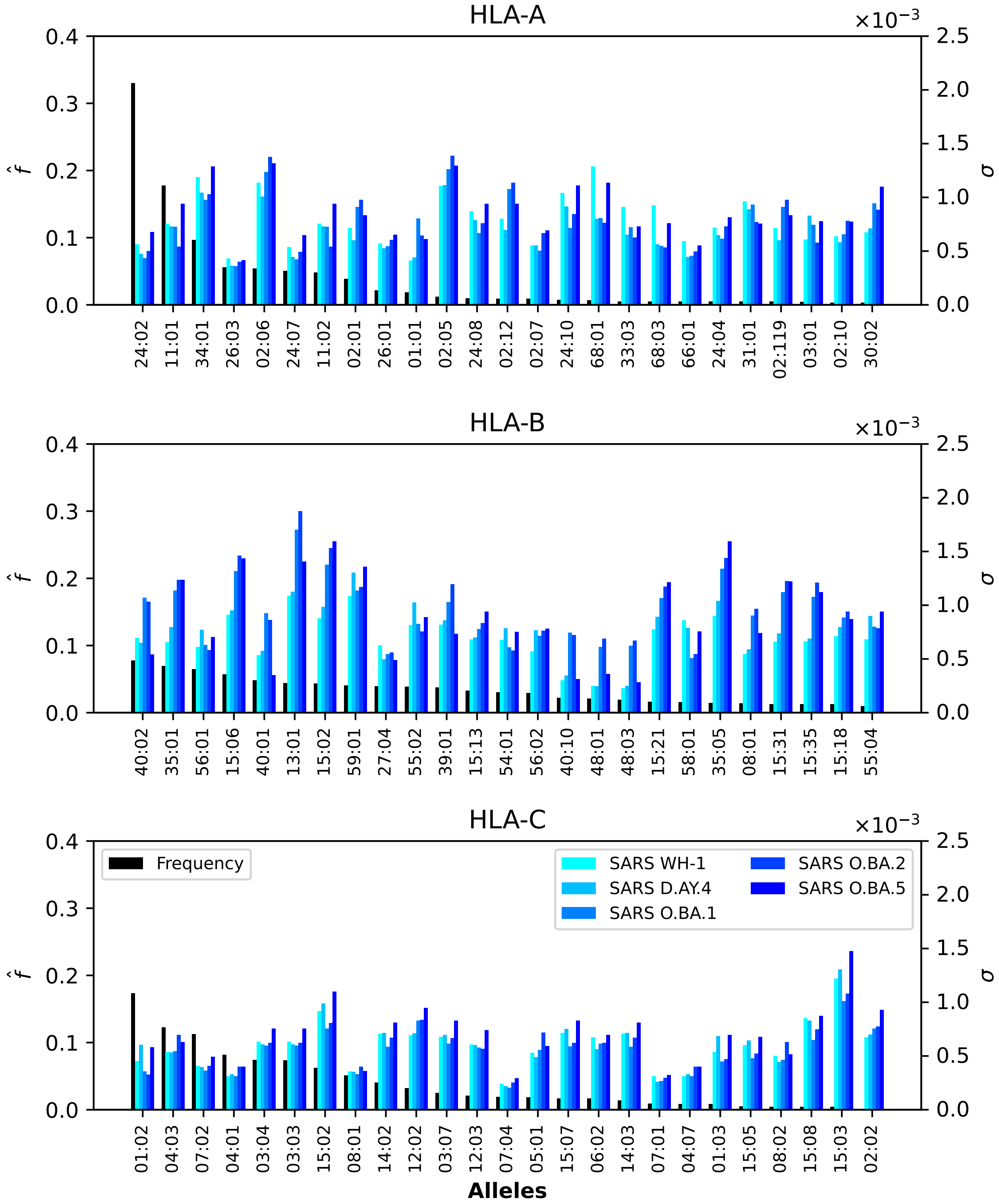


**Figure S27.** Normalized regional frequencies ($\hat{f}_i^{(6)}$) and SARS-CoV-2 $\sigma_i$ values for the top 25 most frequent alleles of each type in Oceania. The top panel represents HLA-A alleles, the middle HLA-B, and the bottom HLA-C. From left to right, the bars in each group represent frequency, SARS-CoV-2 Wuhan-Hu-1, SARS-CoV-2 Delta AY.4, SARS-CoV-2 Omicron BA.1, SARS-CoV-2 Omicron BA.2, SARS-CoV-2 Omicron BA.5.

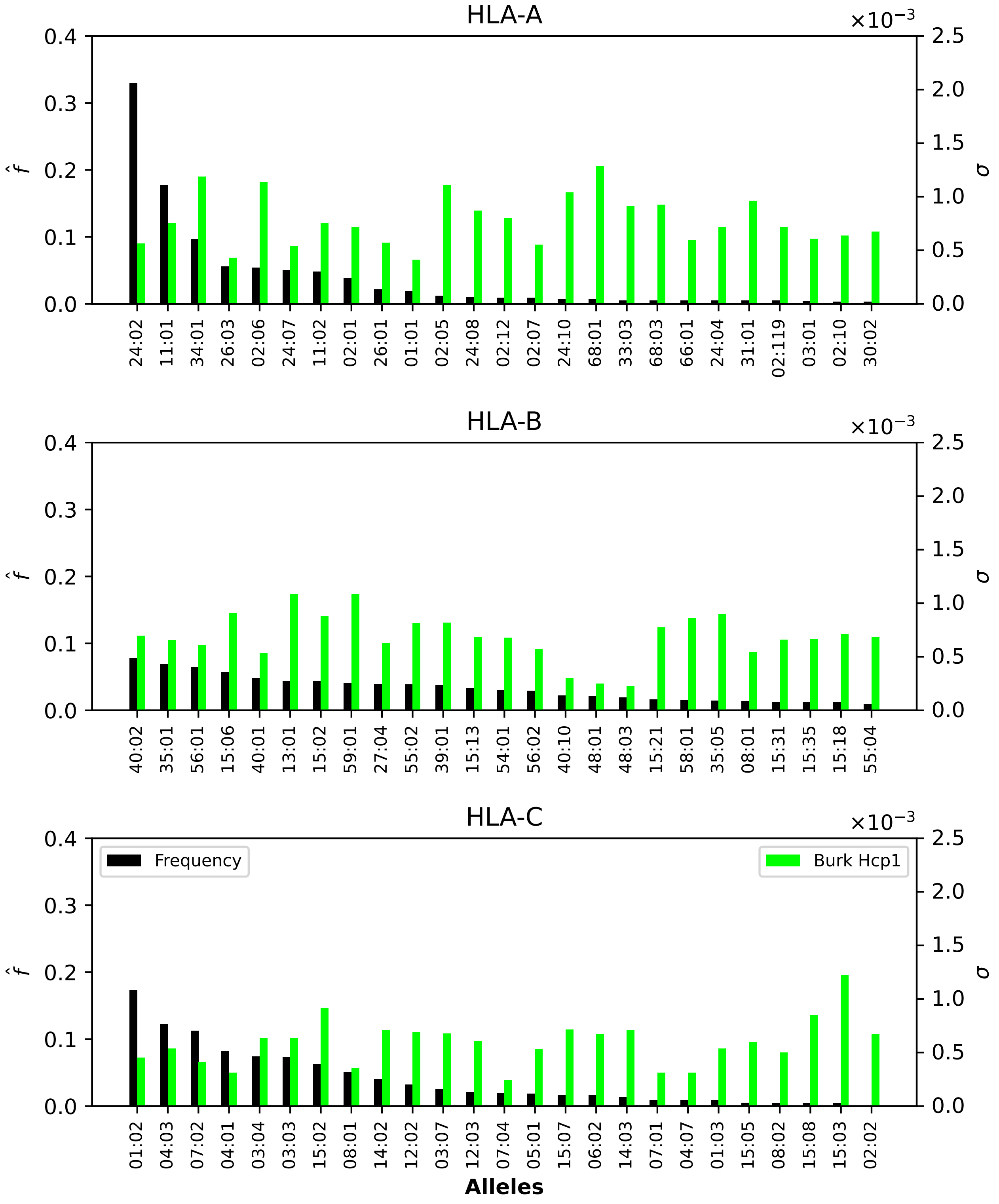


**Figure S28.** Normalized regional frequencies ($\hat{f}_i^{(6)}$) and Burkholderia $\sigma_i$ values for the top 25 most frequent alleles of each type in Oceania. The top panel represents HLA-A alleles, the middle HLA-B, and the bottom HLA-C. From left to right, the bars in each group represent frequency and Burkholderia HCP1.

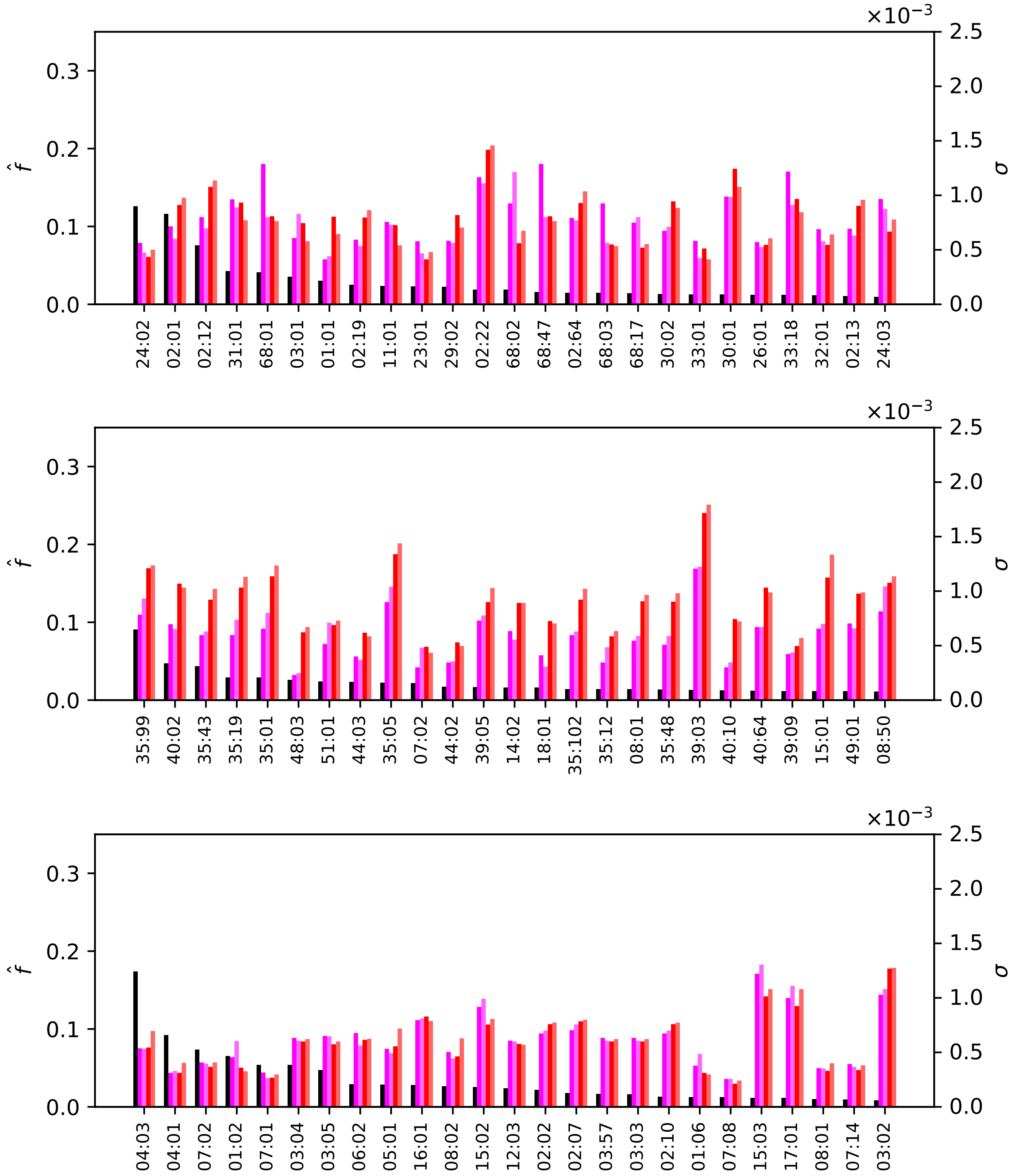


**Figure S29.** Normalized regional frequencies ($\hat{f}_i^{(7)}$) and Ebola $\sigma_i$ values for the top 25 most frequent alleles of each type in South and Central America. The top panel represents HLA-A alleles, the middle HLA-B, and the bottom HLA-C. From left to right, the bars in each group represent frequency, Ebola GP1 (Zaire), Ebola GP1 (Sudan), Ebola NP (Zaire), and Ebola NP (Sudan).

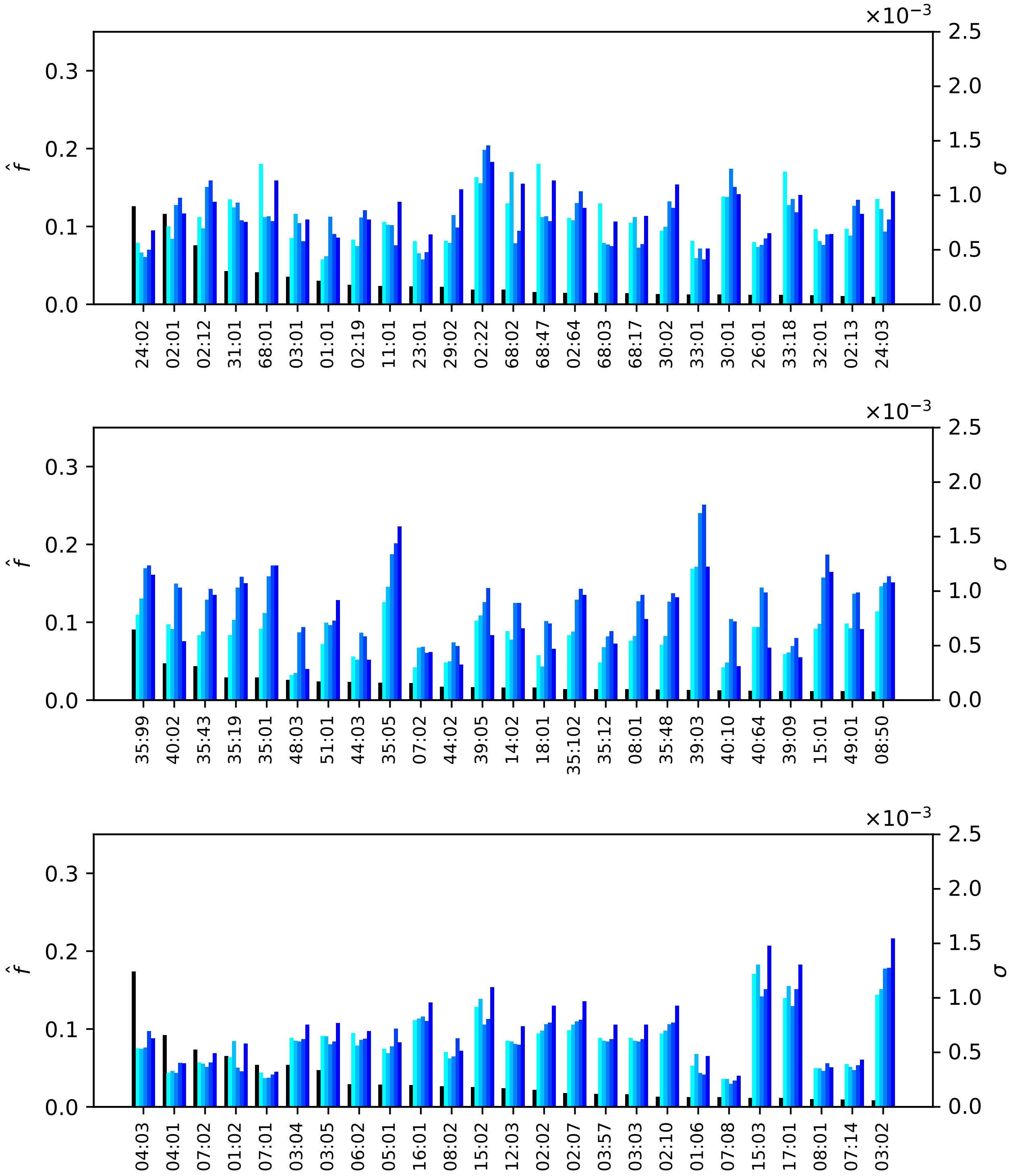


**Figure S30.** Normalized regional frequencies ($\hat{f}_i^{(7)}$) and SARS-CoV-2 $\sigma_i$ values for the top 25 most frequent alleles of each type in South and Central America. The top panel represents HLA-A alleles, the middle HLA-B, and the bottom HLA-C. From left to right, the bars in each group represent frequency, SARS-CoV-2 Wuhan-Hu-1, SARS-CoV-2 Delta AY.4, SARS-CoV-2 Omicron BA.1, SARS-CoV-2 Omicron BA.2, SARS-CoV-2 Omicron BA.5.

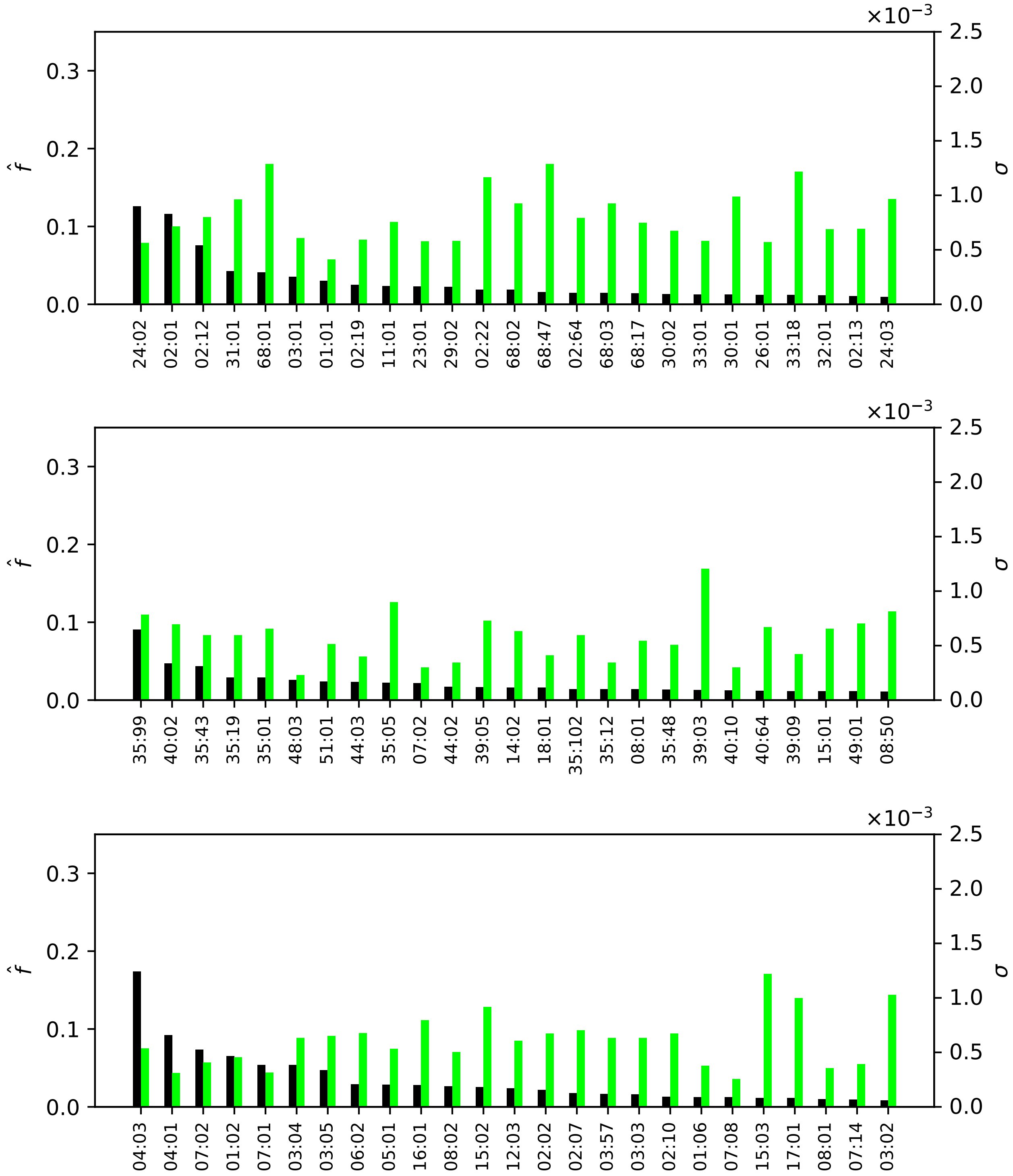


**Figure S31.** Normalized regional frequencies ($\hat{f}_i^{(7)}$) and Burkholderia $\sigma_i$ values for the top 25 most frequent alleles of each type in South and Central America. The top panel represents HLA-A alleles, the middle HLA-B, and the bottom HLA-C. From left to right, the bars in each group represent frequency and Burkholderia HCP1.

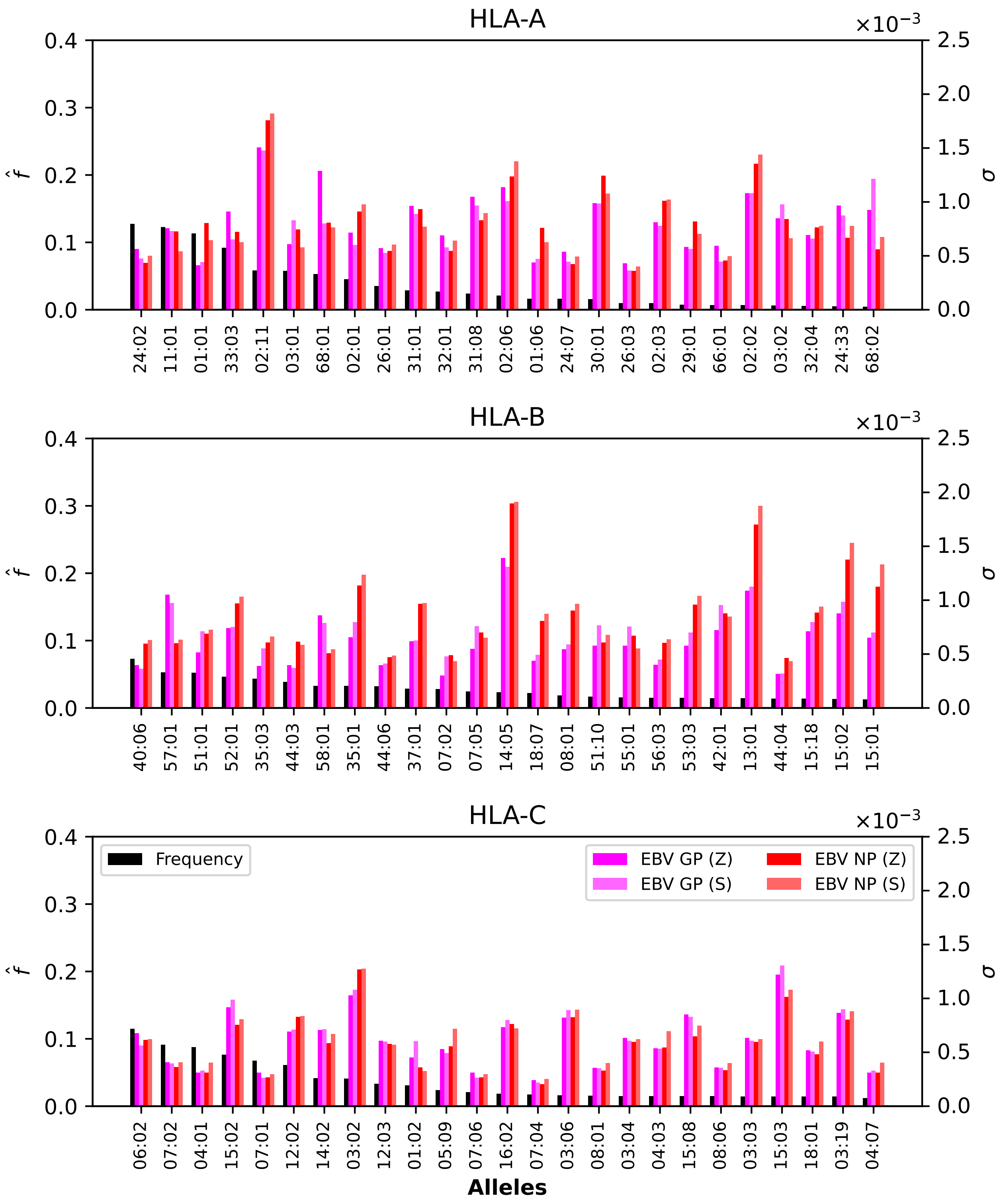


**Figure S32.** Normalized regional frequencies ($\hat{f}_i^{(8)}$) and Ebola $\sigma_i$ values for the top 25 most frequent alleles of each type in South Asia. The top panel represents HLA-A alleles, the middle HLA-B, and the bottom HLA-C. From left to right, the bars in each group represent frequency, Ebola GP1 (Zaire), Ebola GP1 (Sudan), Ebola NP (Zaire), and Ebola NP (Sudan).

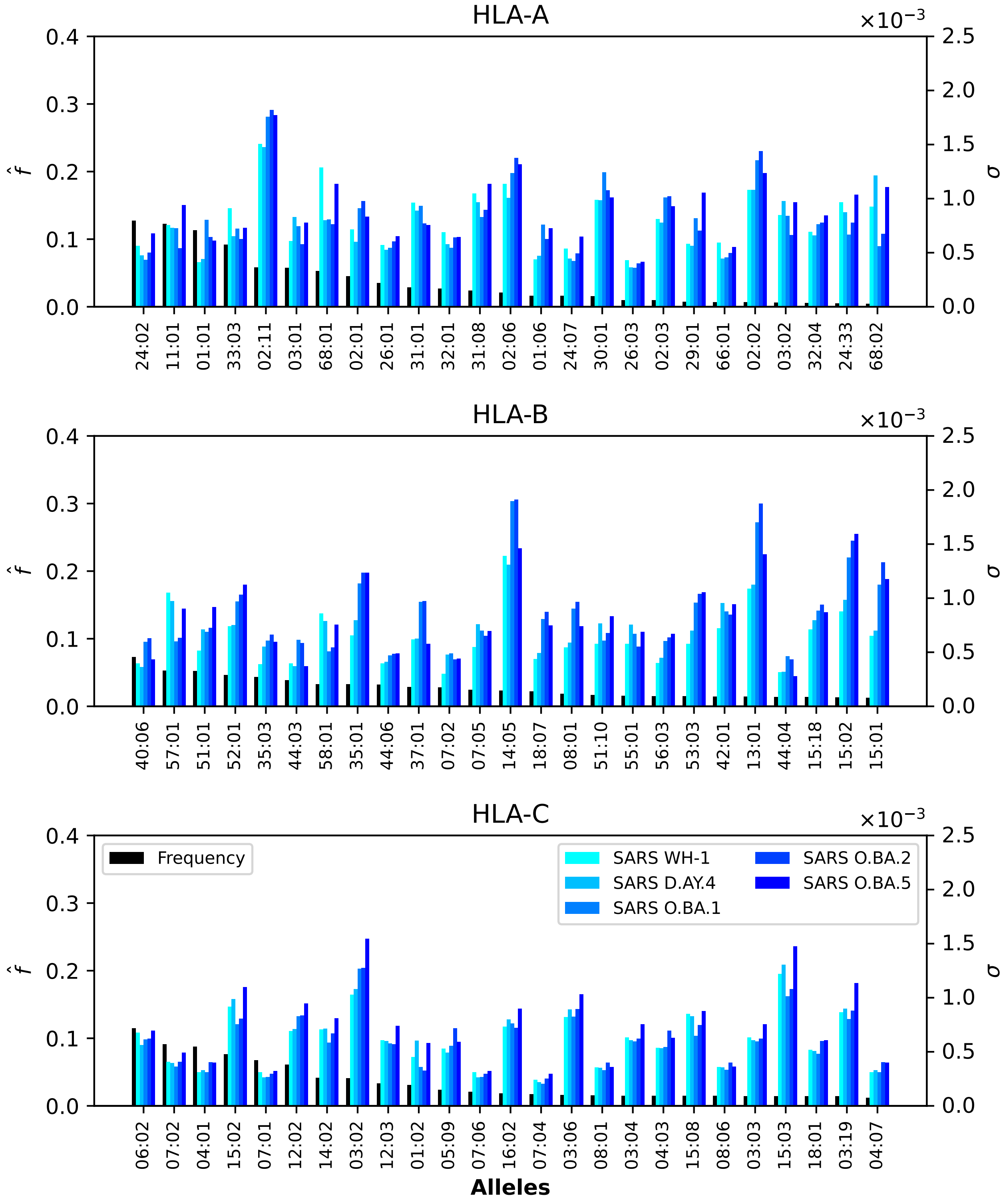


**Figure S33.** Normalized regional frequencies ($\hat{f}_i^{(8)}$) and SARS-CoV-2 $\sigma_i$ values for the top 25 most frequent alleles of each type in South Asia. The top panel represents HLA-A alleles, the middle HLA-B, and the bottom HLA-C. From left to right, the bars in each group represent frequency, SARS-CoV-2 Wuhan-Hu-1, SARS-CoV-2 Delta AY.4, SARS-CoV-2 Omicron BA.1, SARS-CoV-2 Omicron BA.2, SARS-CoV-2 Omicron BA.5.

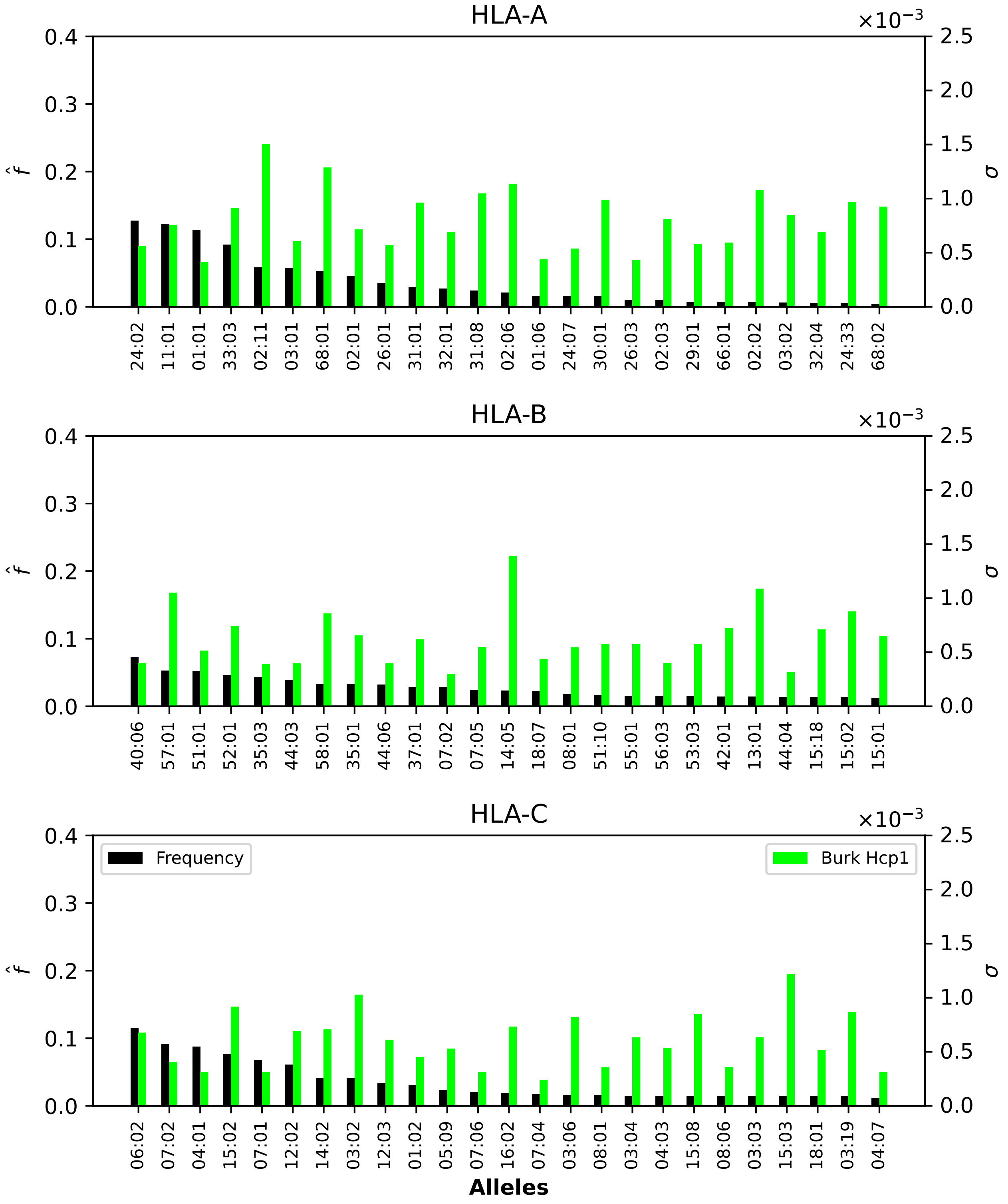


**Figure S34.** Normalized regional frequencies ($\hat{f}_i^{(8)}$) and Burkholderia $\sigma_i$ values for the top 25 most frequent alleles of each type in South Asia. The top panel represents HLA-A alleles, the middle HLA-B, and the bottom HLA-C. From left to right, the bars in each group represent frequency and Burkholderia HCP1.

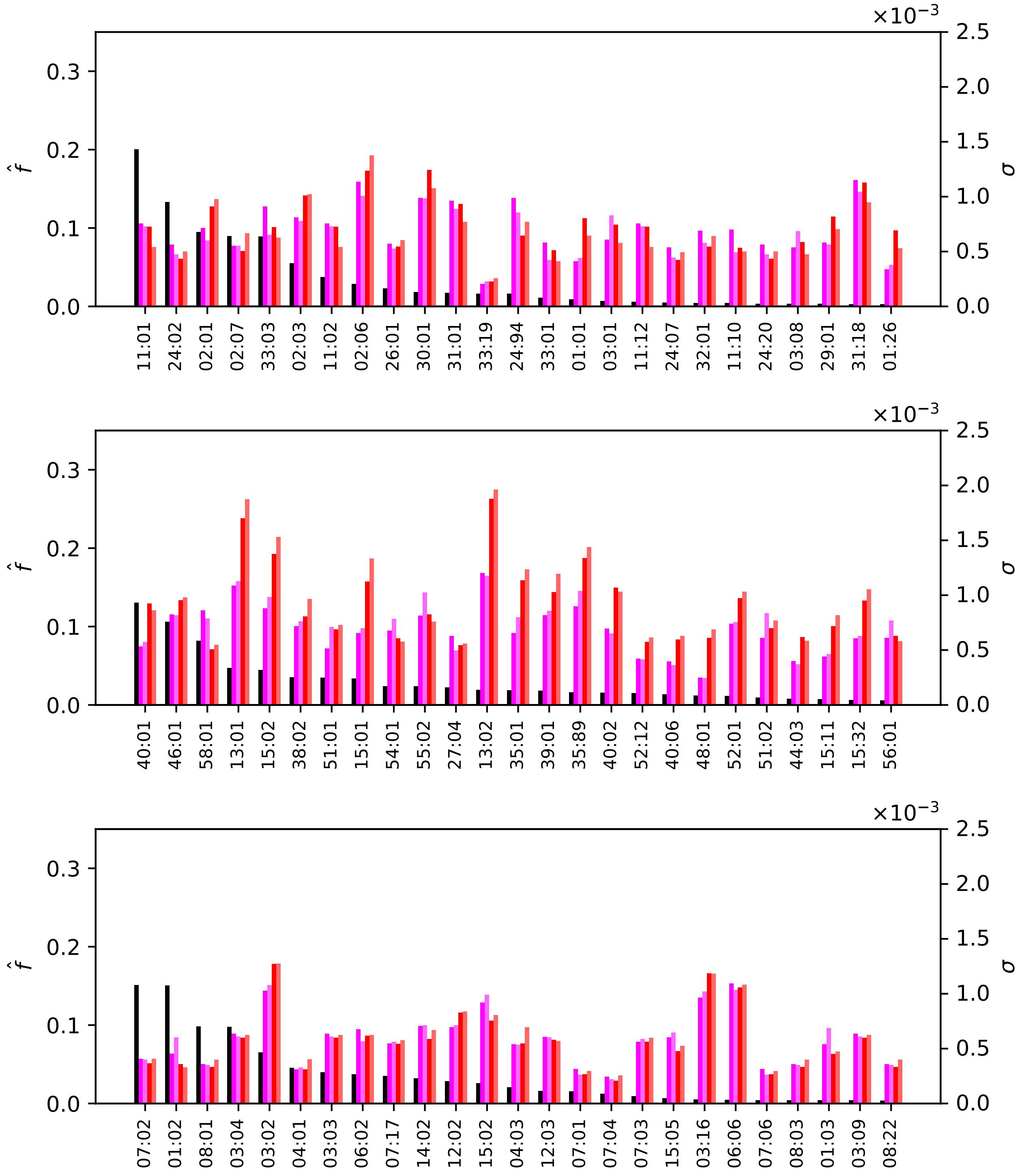


**Figure S35.** Normalized regional frequencies ($\hat{f}_i^{(9)}$) and Ebola $\sigma_i$ values for the top 25 most frequent alleles of each type in Southeast Asia. The top panel represents HLA-A alleles, the middle HLA-B, and the bottom HLA-C. From left to right, the bars in each group represent frequency, Ebola GP1 (Zaire), Ebola GP1 (Sudan), Ebola NP (Zaire), and Ebola NP (Sudan).

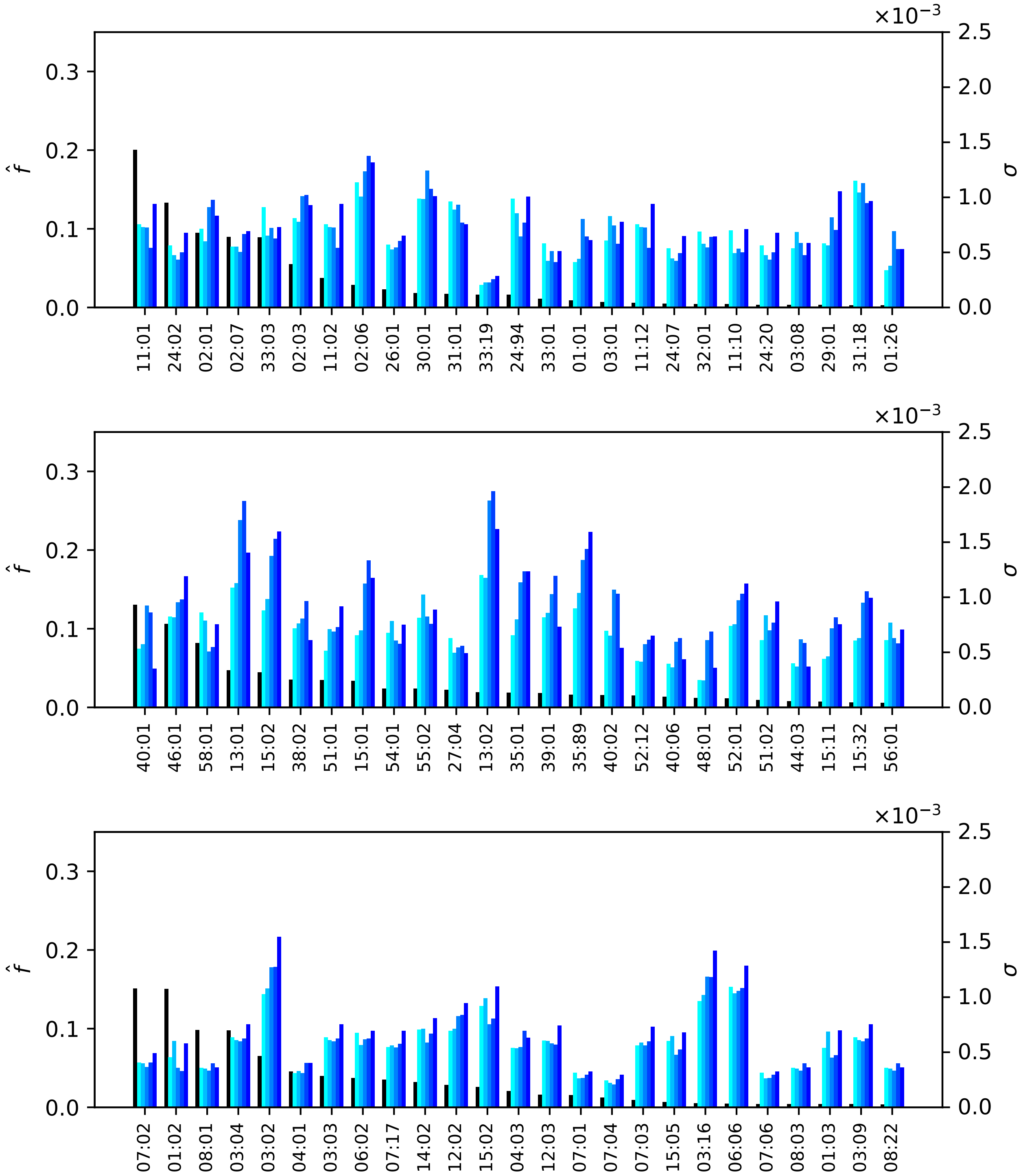


**Figure S36.** Normalized regional frequencies ($\hat{f}_i^{(9)}$) and SARS-CoV-2 $\sigma_i$ values for the top 25 most frequent alleles of each type in Southeast Asia. The top panel represents HLA-A alleles, the middle HLA-B, and the bottom HLA-C. From left to right, the bars in each group represent frequency, SARS-CoV-2 Wuhan-Hu-1, SARS-CoV-2 Delta AY.4, SARS-CoV-2 Omicron BA.1, SARS-CoV-2 Omicron BA.2, SARS-CoV-2 Omicron BA.5.

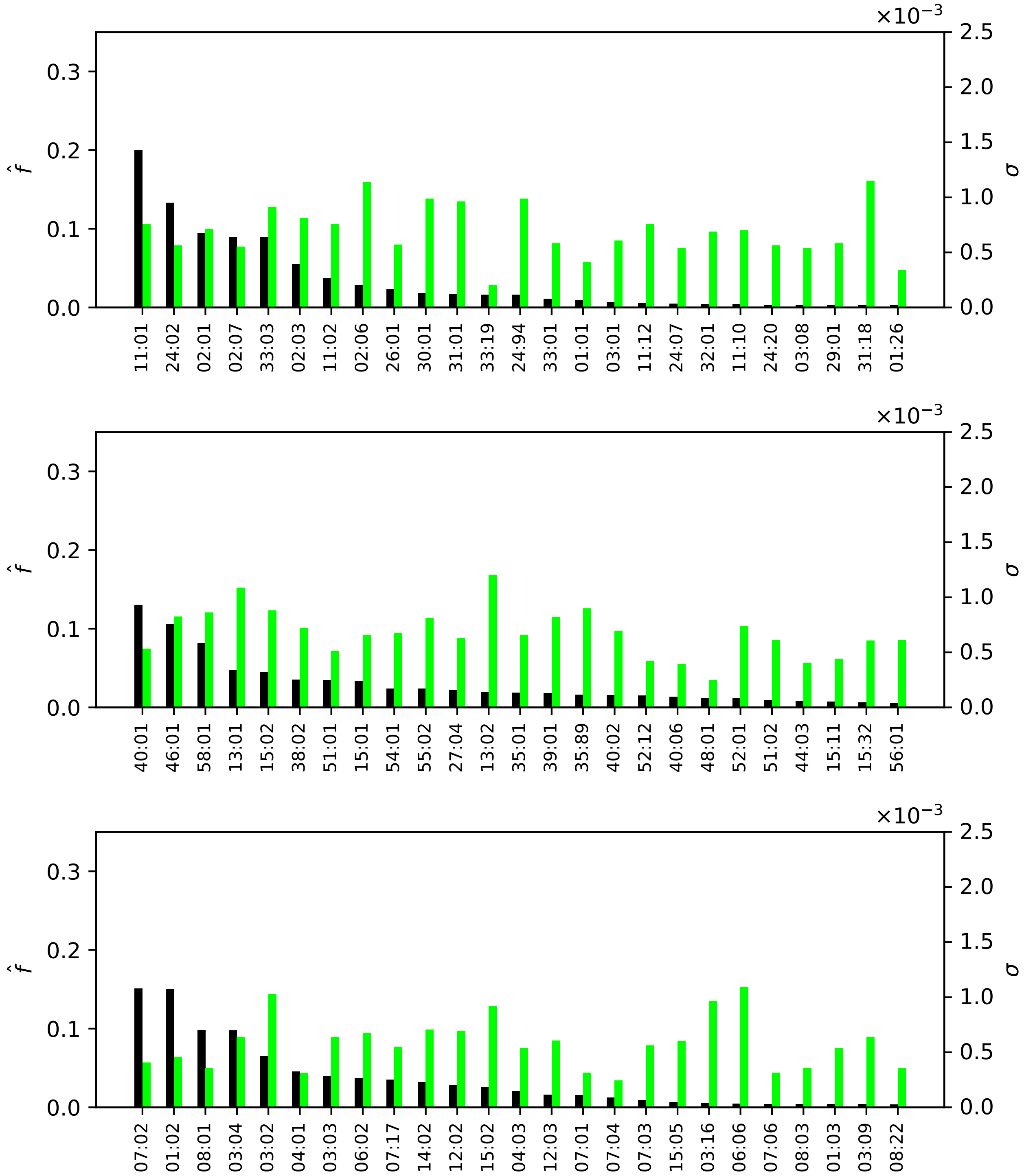


**Figure S37.** Normalized regional frequencies ($\hat{f}_i^{(9)}$) and Burkholderia $\sigma_i$ values for the top 25 most frequent alleles of each type in Southeast Asia. The top panel represents HLA-A alleles, the middle HLA-B, and the bottom HLA-C. From left to right, the bars in each group represent frequency and Burkholderia HCP1.

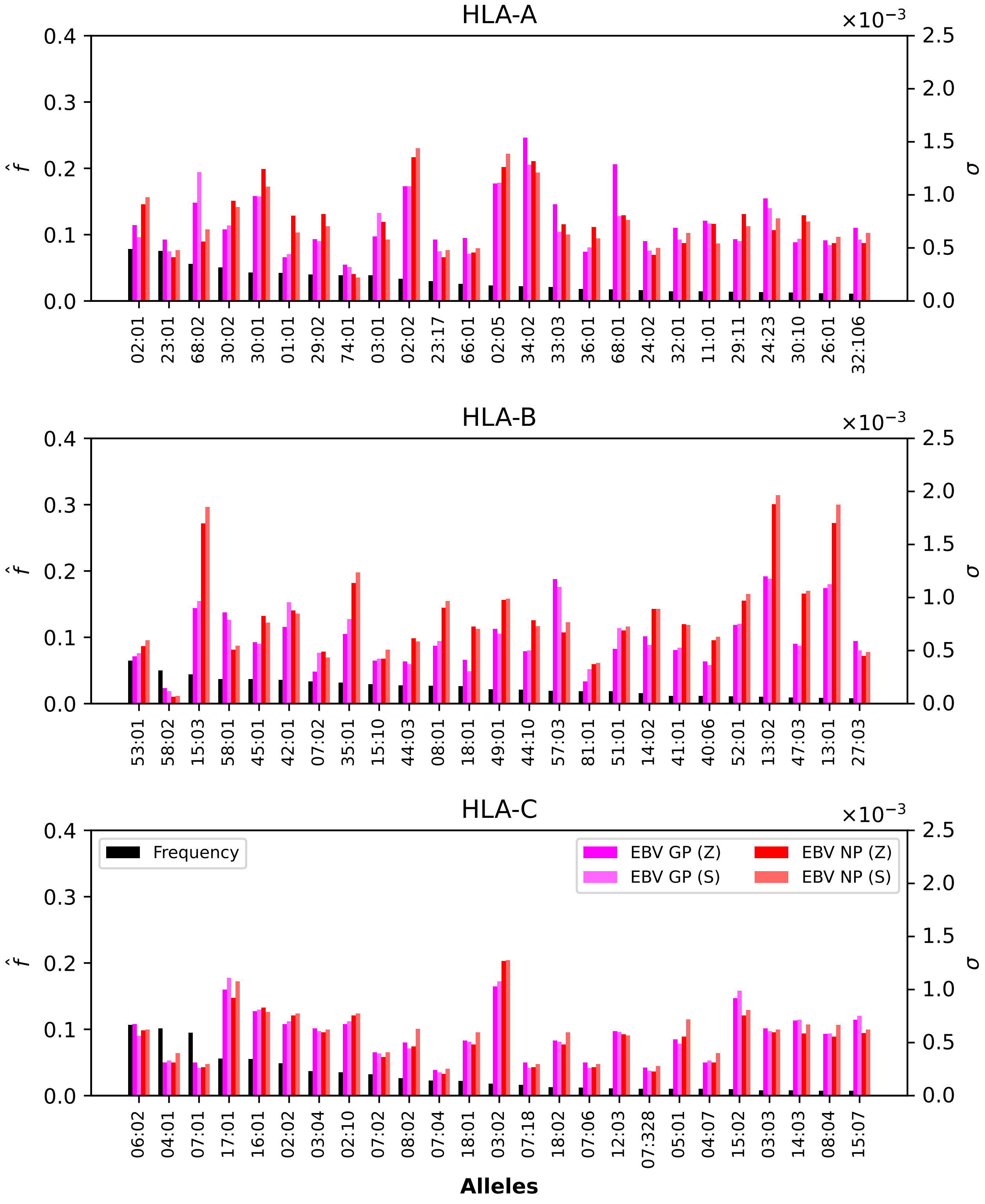


**Figure S38.** Normalized regional frequencies ($\hat{f}_i^{(10)}$) and Ebola $\sigma_i$ values for the top 25 most frequent alleles of each type in Sub-Saharan Africa. The top panel represents HLA-A alleles, the middle HLA-B, and the bottom HLA-C. From left to right, the bars in each group represent frequency, Ebola GP1 (Zaire), Ebola GP1 (Sudan), Ebola NP (Zaire), and Ebola NP (Sudan).

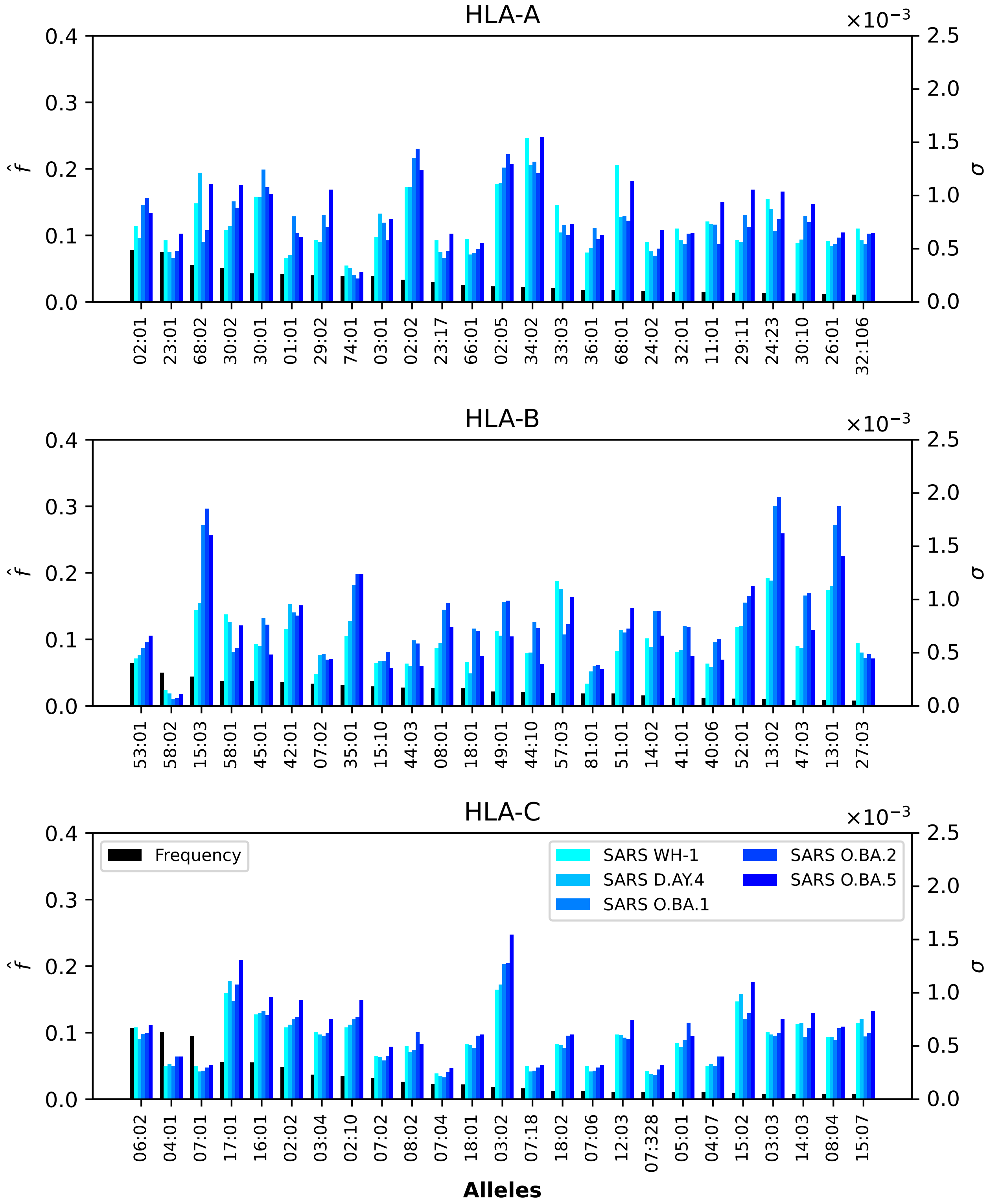


**Figure S39.** Normalized regional frequencies ($\hat{f}_i^{(10)}$) and SARS-CoV-2 $\sigma_i$ values for the top 25 most frequent alleles of each type in Sub-Saharan Africa. The top panel represents HLA-A alleles, the middle HLA-B, and the bottom HLA-C. From left to right, the bars in each group represent frequency, SARS-CoV-2 Wuhan-Hu-1, SARS-CoV-2 Delta AY.4, SARS-CoV-2 Omicron BA.1, SARS-CoV-2 Omicron BA.2, SARS-CoV-2 Omicron BA.5.

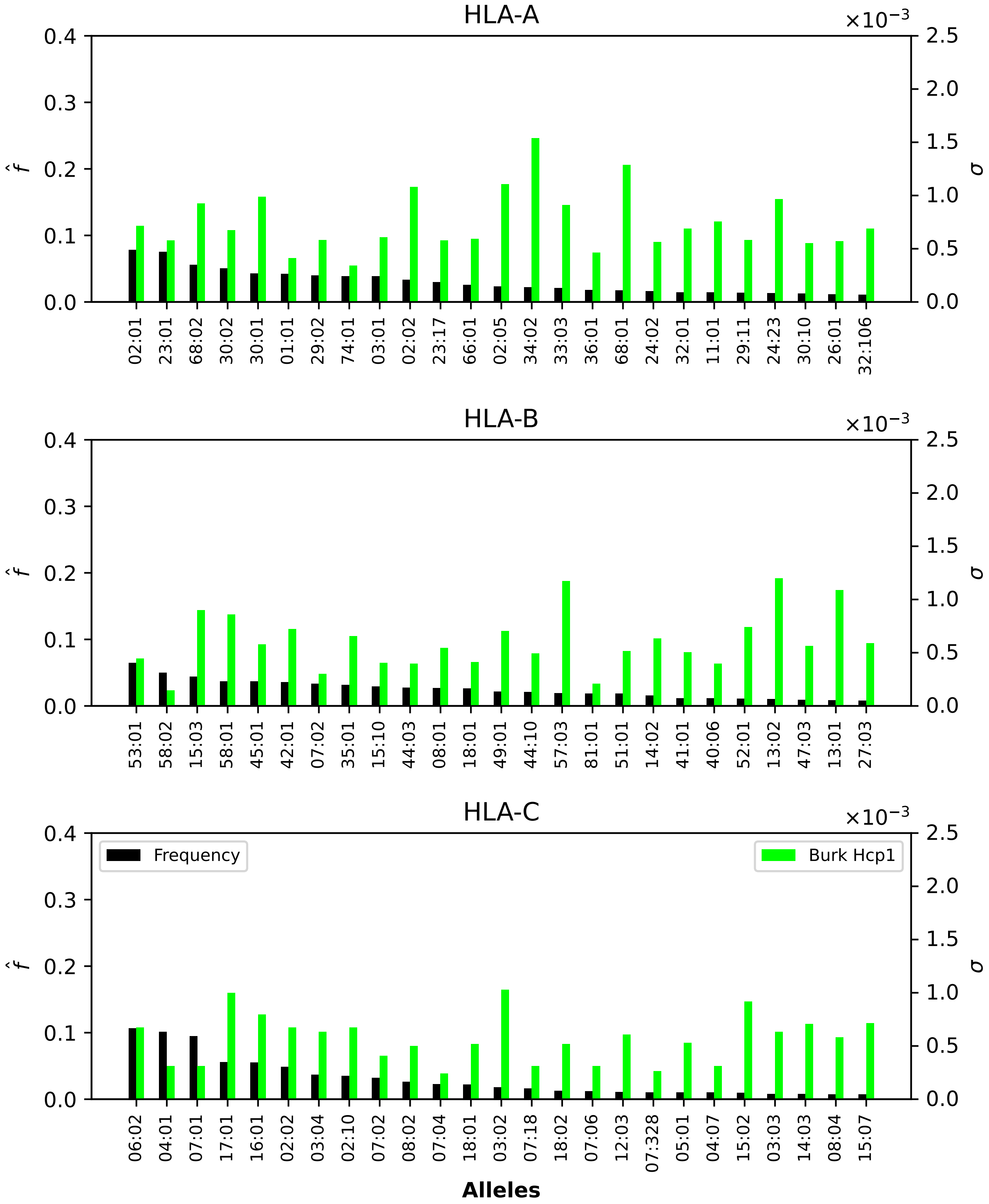


**Figure S40.** Normalized regional frequencies ($\hat{f}_i^{(10)}$) and Burkholderia $\sigma_i$ values for the top 25 most frequent alleles of each type in Sub-Saharan Africa. The top panel represents HLA-A alleles, the middle HLA-B, and the bottom HLA-C. From left to right, the bars in each group represent frequency and Burkholderia HCP1.

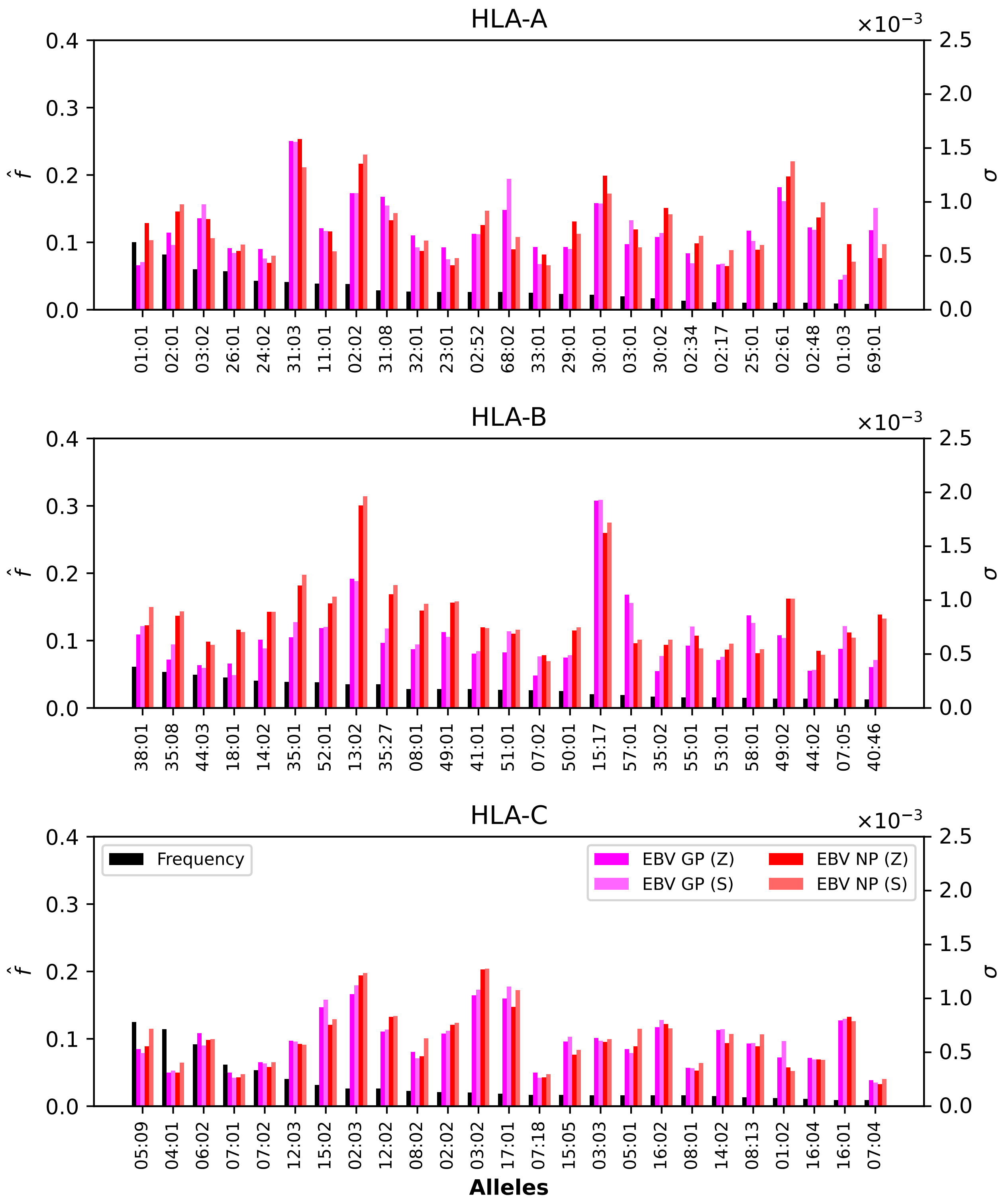


**Figure S41.** Normalized regional frequencies ($\hat{f}_i^{(11)}$) and Ebola $\sigma_i$ values for the top 25 most frequent alleles of each type in Western Asia. The top panel represents HLA-A alleles, the middle HLA-B, and the bottom HLA-C. From left to right, the bars in each group represent frequency, Ebola GP1 (Zaire), Ebola GP1 (Sudan), Ebola NP (Zaire), and Ebola NP (Sudan).

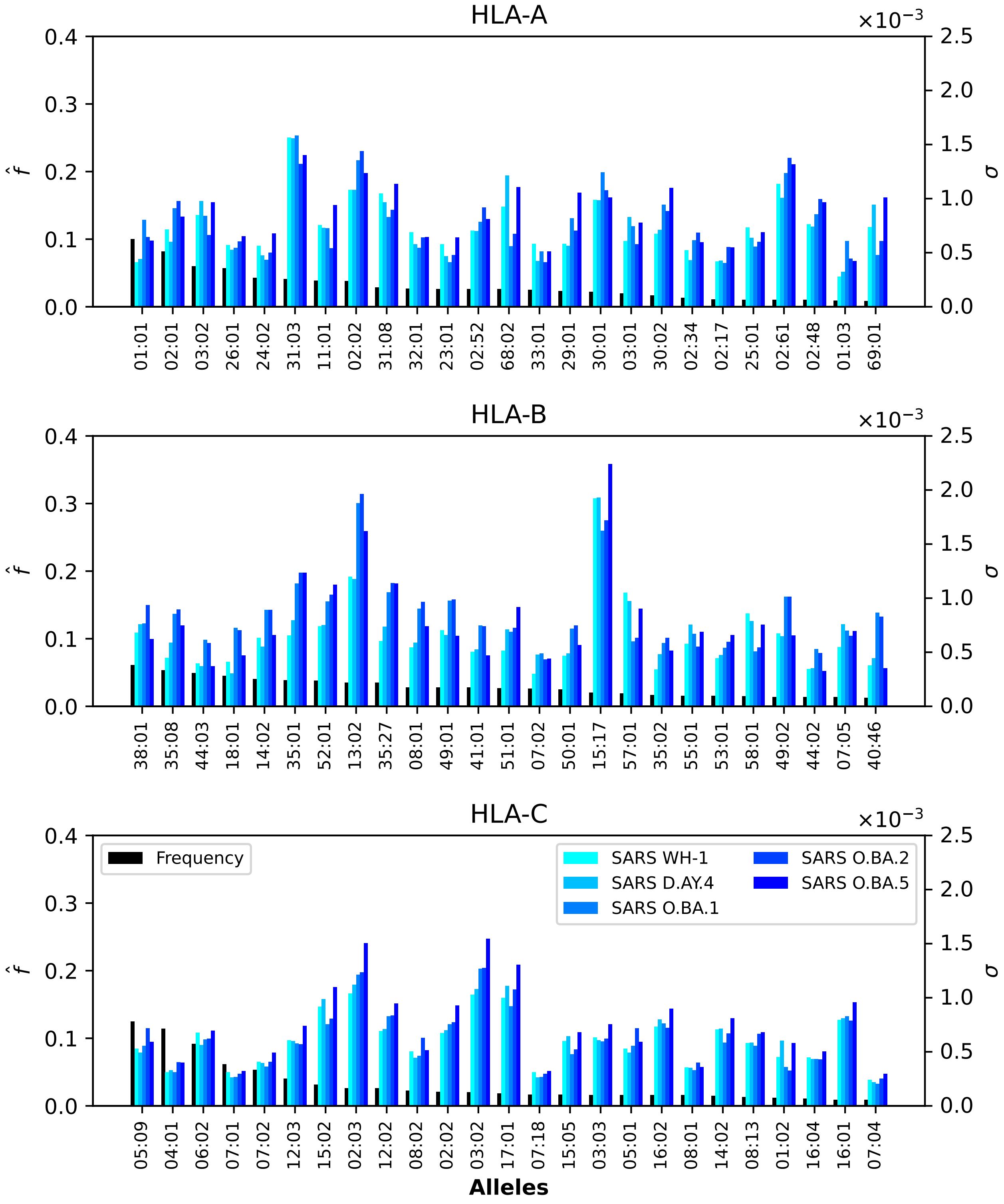


**Figure S42.** Normalized regional frequencies ($\hat{f}_i^{(11)}$) and SARS-CoV-2 $\sigma_i$ values for the top 25 most frequent alleles of each type in Western Asia. The top panel represents HLA-A alleles, the middle HLA-B, and the bottom HLA-C. From left to right, the bars in each group represent frequency, SARS-CoV-2 Wuhan-Hu-1, SARS-CoV-2 Delta AY.4, SARS-CoV-2 Omicron BA.1, SARS-CoV-2 Omicron BA.2, SARS-CoV-2 Omicron BA.5.

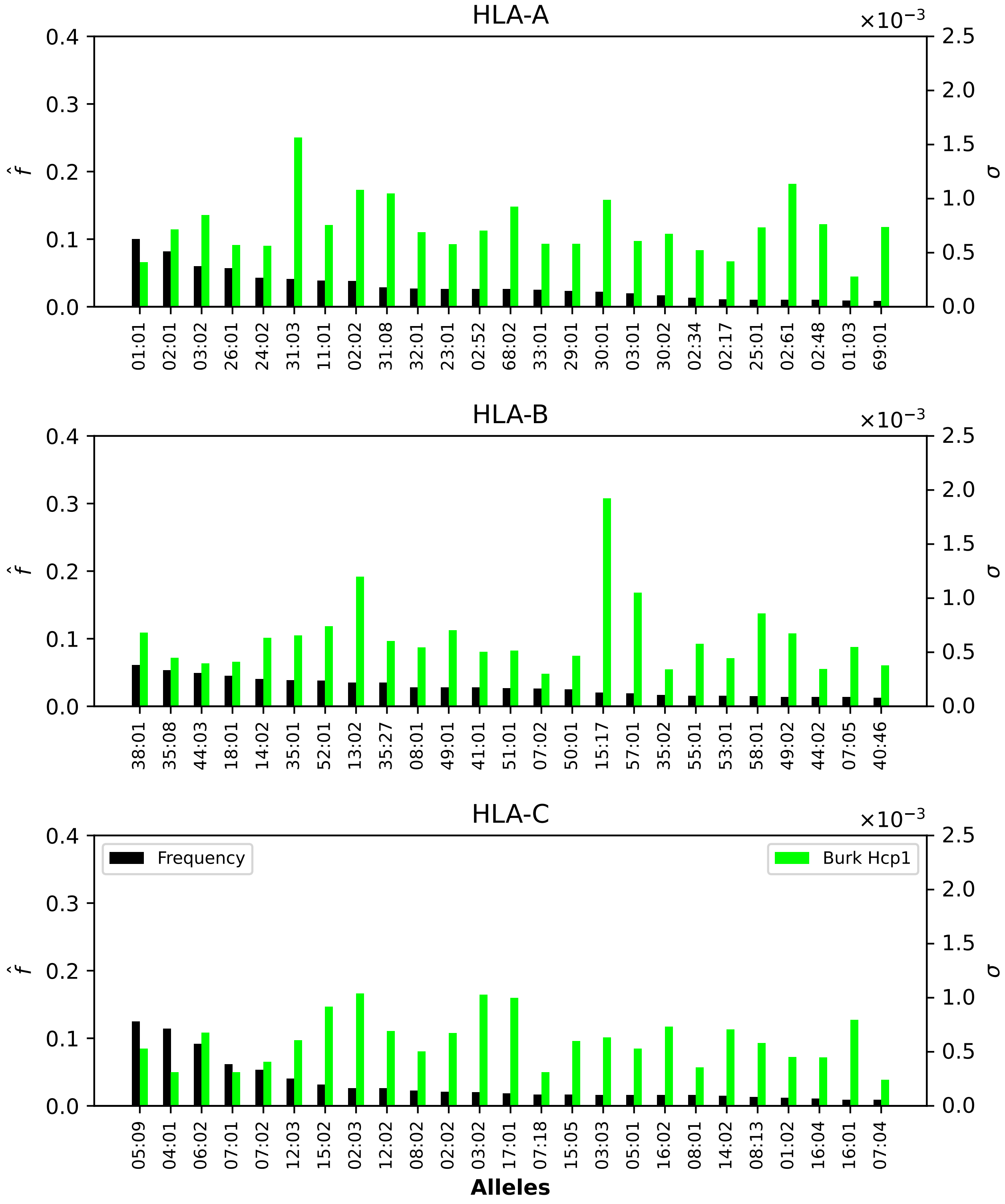


**Figure S43.** Normalized regional frequencies ($\hat{f}_i^{(11)}$) and Burkholderia $\sigma_i$ values for the top 25 most frequent alleles of each type in Western Asia. The top panel represents HLA-A alleles, the middle HLA-B, and the bottom HLA-C. From left to right, the bars in each group represent frequency and Burkholderia HCP1.

### 3 DISSECTING THE CONTRIBUTION TO THE INDIVIDUAL COVERAGE METRIC: ALLELE PAIR ANALYSIS FOR ALL REGIONS

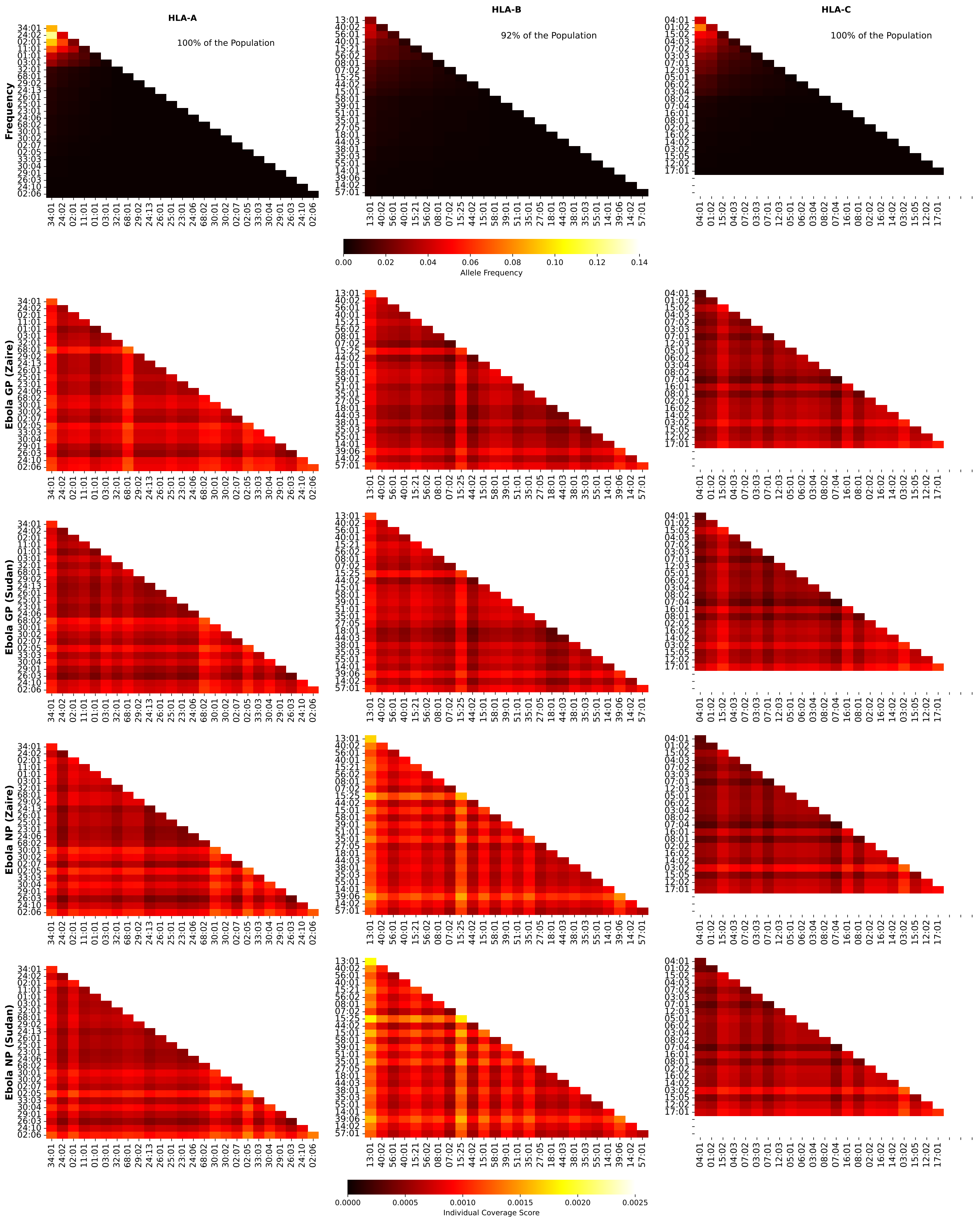


**Figure S44.** Frequencies and Ebola coverage scores for individuals in Australia. The 1st row corresponds to allele frequencies, the 2nd to GP1 Zaire, the 3rd to GP1 Sudan, the 4th to NP Zaire, and the 5th to NP Sudan. The 1st column is associated with HLA-A alleles, the 2nd to HLA-B, and the 3rd to HLA-C. The sum of the individual frequencies for each allele type is indicated on the panels in the 1st row.

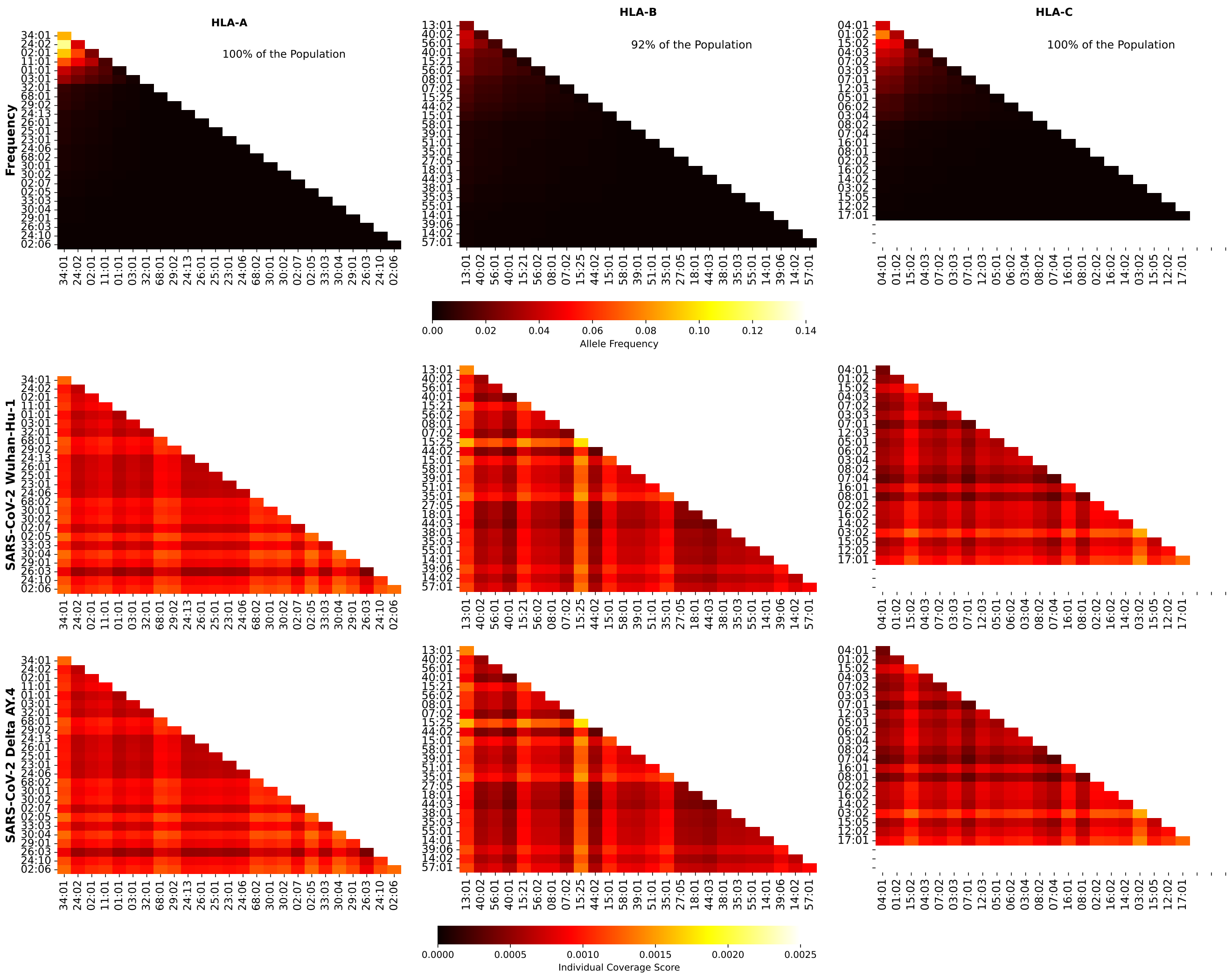


**Figure S45.** Frequencies and SARS-CoV-2 (Wuhan-Hu-1 and Delta AY.4 variants) coverage scores for individuals in Australia. The 1st row corresponds to allele frequencies, the 2nd to Wuhan-Hu-1, and the 3rd to Delta AY.4. The 1st column is associated with HLA-A alleles, the 2nd to HLA-B, and the 3rd to HLA-C. The sum of the individual frequencies for each allele type is indicated on the panels in the 1st row.

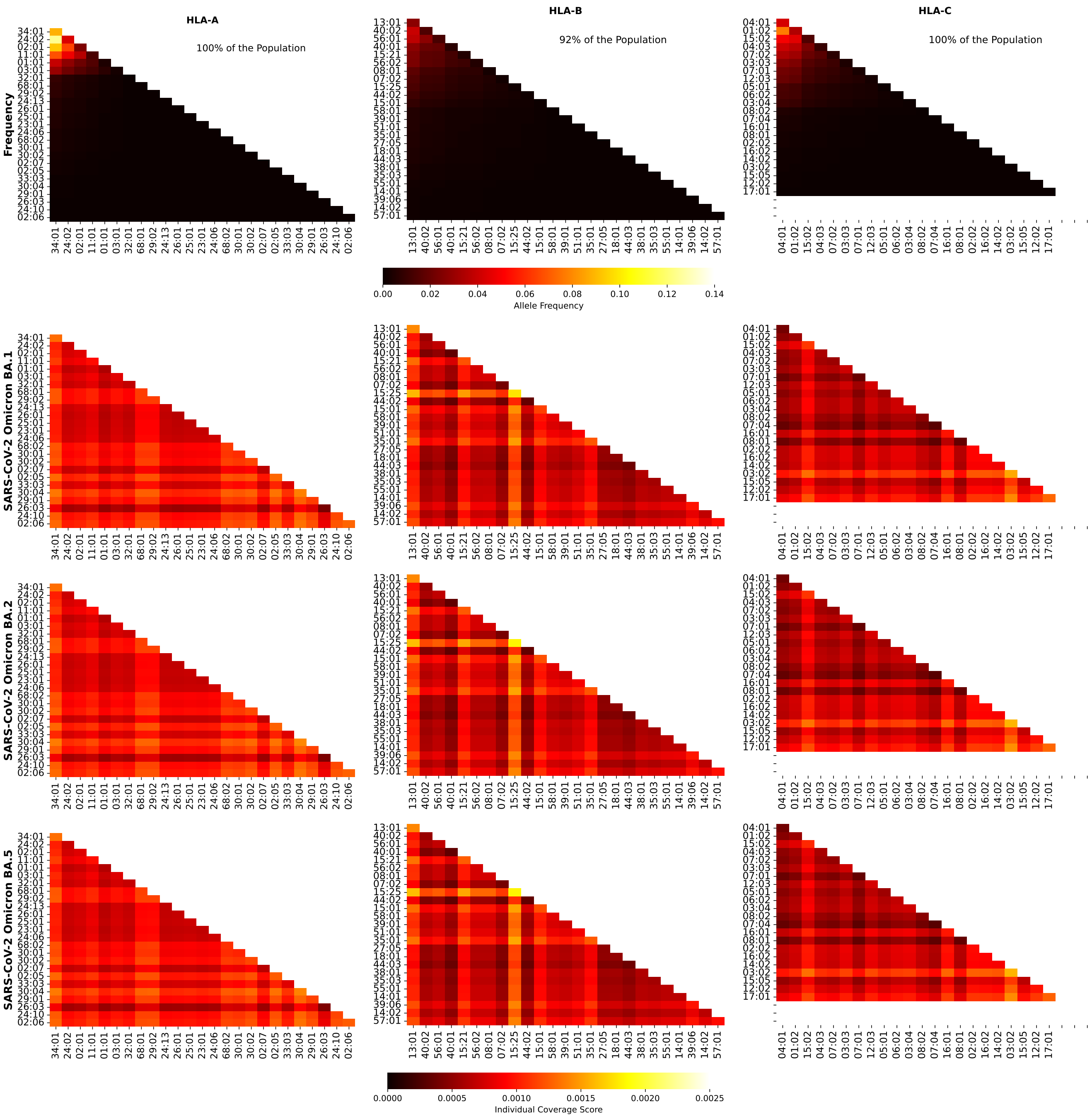


**Figure S46.** Frequencies and SARS-CoV-2 (Omicron variants) coverage scores for individuals in Australia. The 1st row corresponds to allele frequencies, the 2nd to BA.1, and the 3rd to BA.2, and the 4th to BA.5. The 1st column is associated with HLA-A alleles, the 2nd to HLA-B, and the 3rd to HLA-C. The sum of the individual frequencies for each allele type is indicated on the panels in the 1st row.

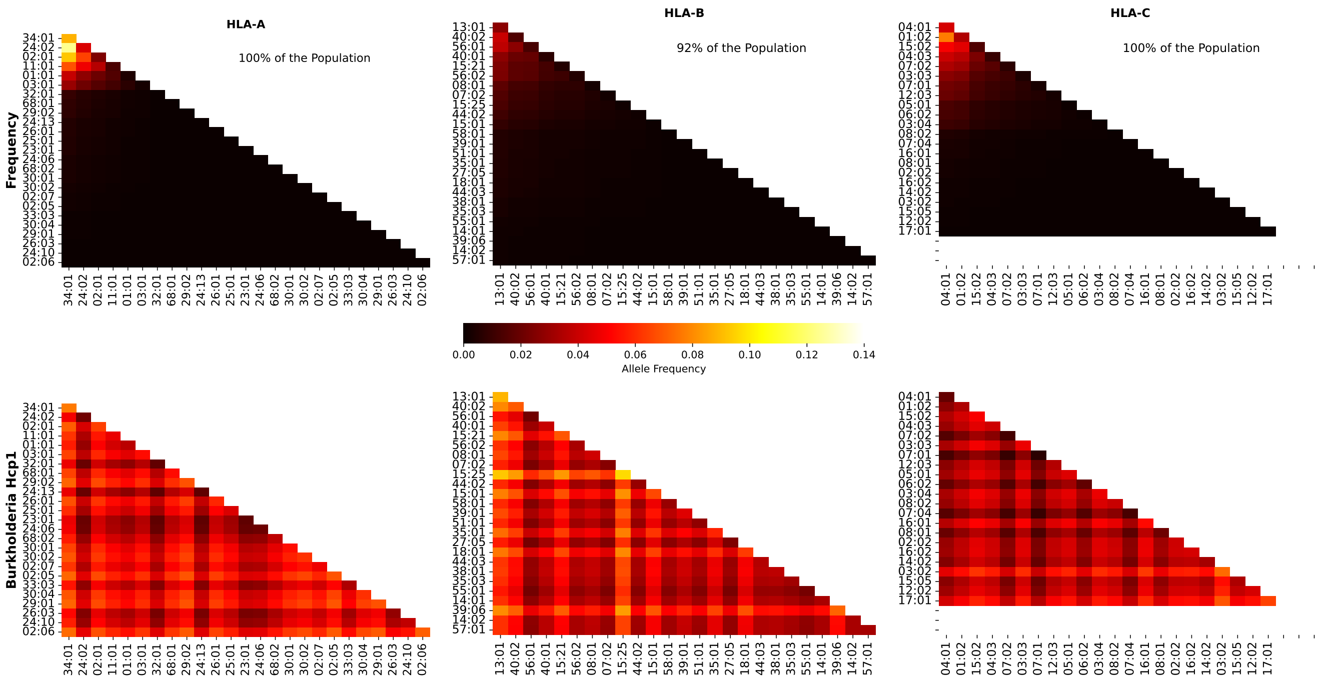


**Figure S47.** Frequencies and Burkholderia coverage scores for individuals in Australia. The 1st row corresponds to allele frequencies and the 2nd row to Burkholderia coverage score. The 1st column is associated with HLA-A alleles, the 2nd to HLA-B, and the 3rd to HLA-C. The sum of the individual frequencies for each allele type is indicated on the panels in the 1st row.

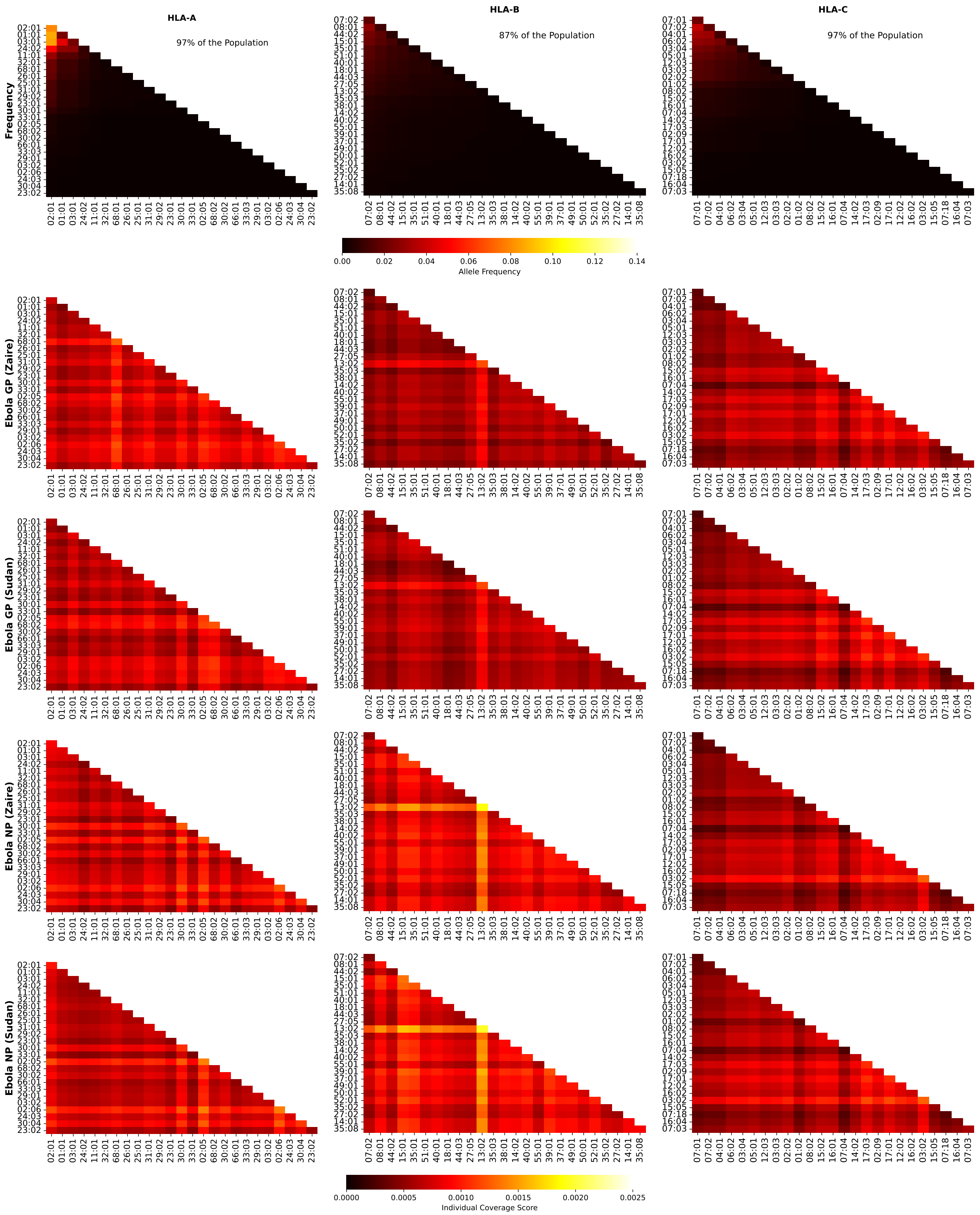


**Figure S48.** Frequencies and Ebola coverage scores for individuals in Europe. The 1st row corresponds to allele frequencies, the 2nd to GP1 Zaire, the 3rd to GP1 Sudan, the 4th to NP Zaire, and the 5th to NP Sudan. The 1st column is associated with HLA-A alleles, the 2nd to HLA-B, and the 3rd to HLA-C. The sum of the individual frequencies for each allele type is indicated on the panels in the 1st row.

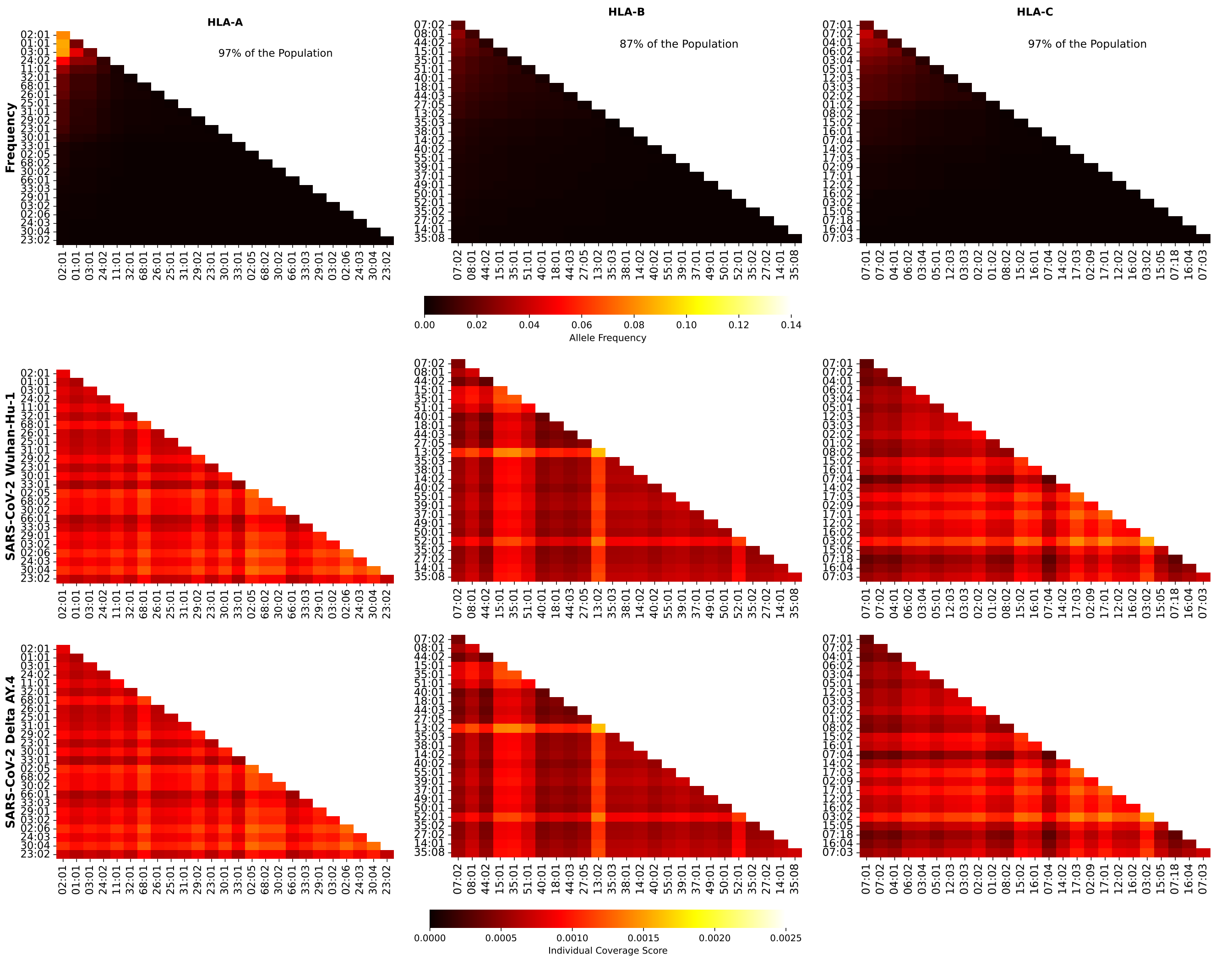


**Figure S49.** Frequencies and SARS-CoV-2 (Wuhan-Hu-1 and Delta AY.4 variants) coverage scores for individuals in Europe. The 1st row corresponds to allele frequencies, the 2nd to Wuhan-Hu-1, and the 3rd to Delta AY.4. The 1st column is associated with HLA-A alleles, the 2nd to HLA-B, and the 3rd to HLA-C. The sum of the individual frequencies for each allele type is indicated on the panels in the 1st row.

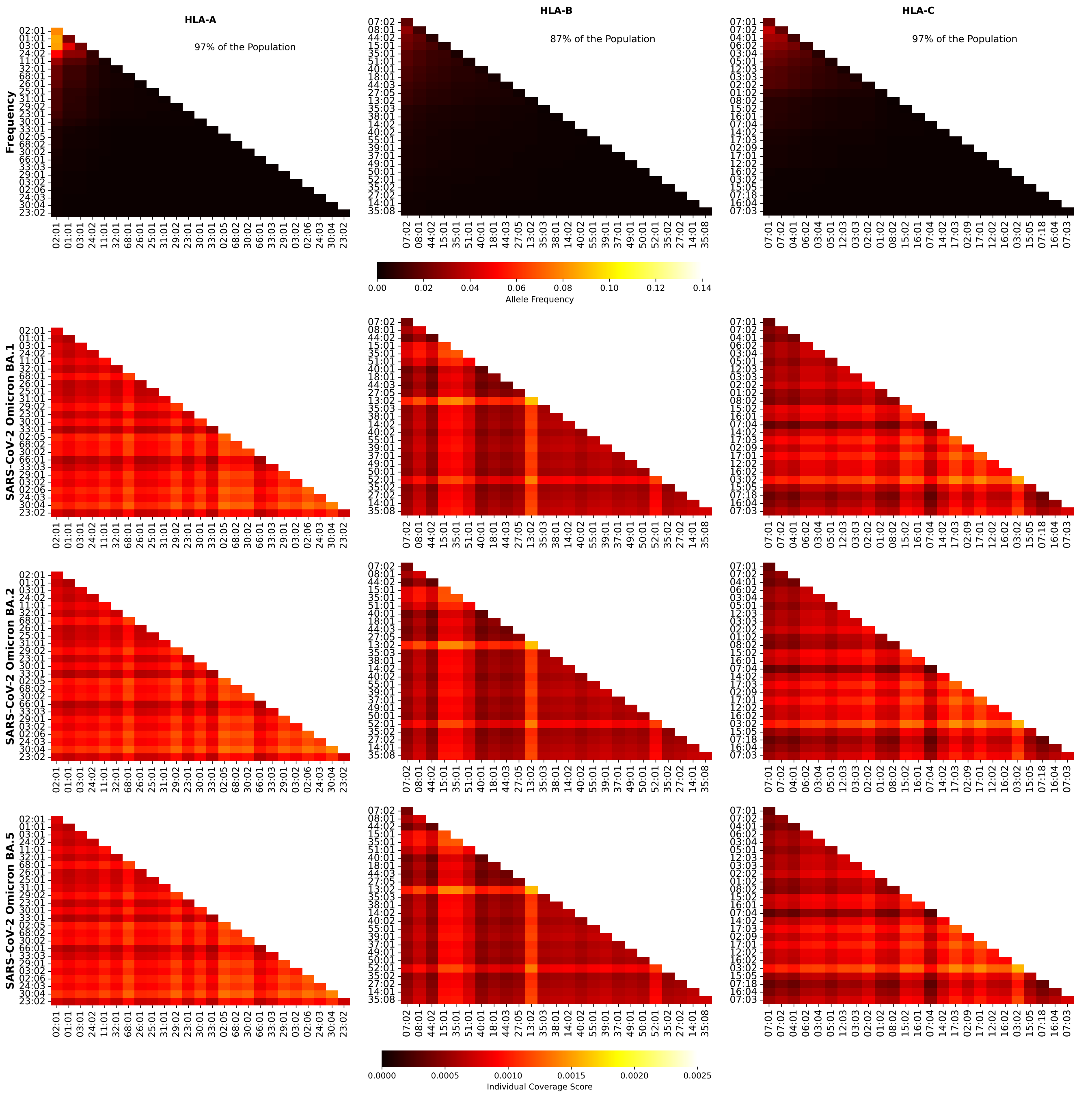


**Figure S50.** Frequencies and SARS-CoV-2 (Omicron variants) coverage scores for individuals in Europe. The 1st row corresponds to allele frequencies, the 2nd to BA.1, and the 3rd to BA.2, and the 4th to BA.5. The 1st column is associated with HLA-A alleles, the 2nd to HLA-B, and the 3rd to HLA-C. The sum of the individual frequencies for each allele type is indicated on the panels in the 1st row.

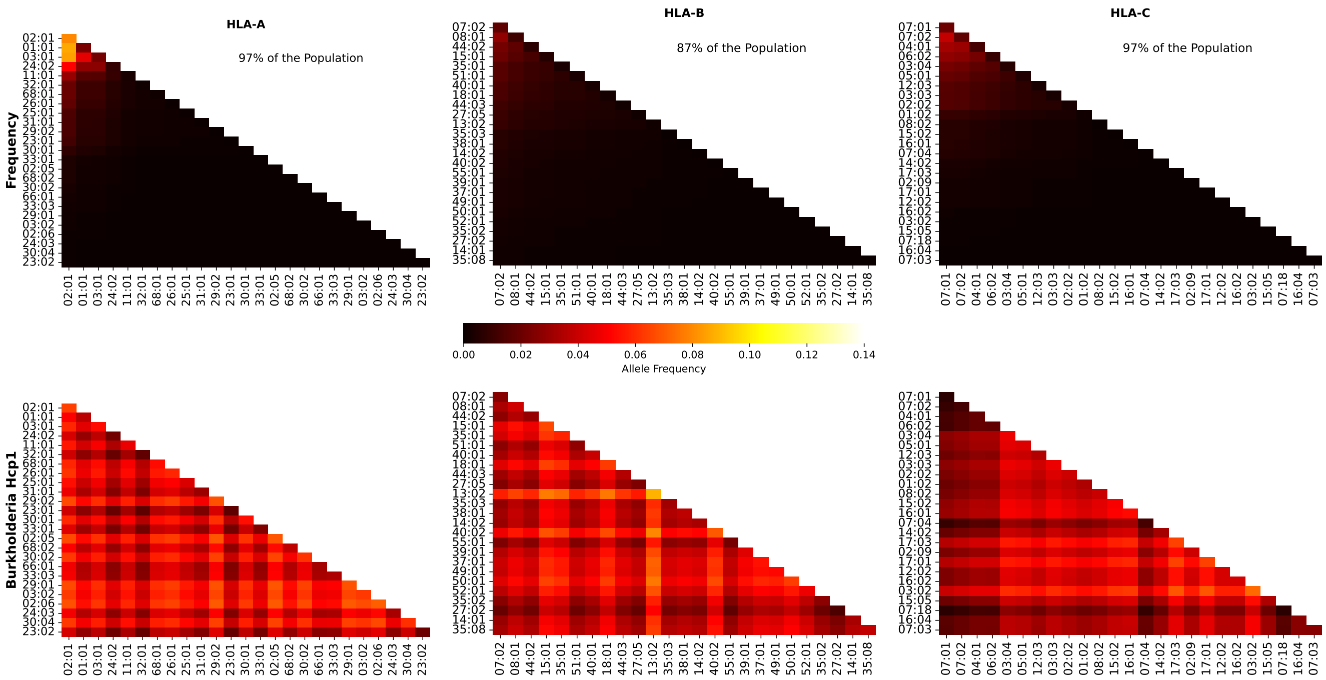


**Figure S51.** Frequencies and Burkholderia coverage scores for individuals in Europe. The 1st row corresponds to allele frequencies and the 2nd row to Burkholderia coverage score. The 1st column is associated with HLA-A alleles, the 2nd to HLA-B, and the 3rd to HLA-C. The sum of the individual frequencies for each allele type is indicated on the panels in the 1st row.

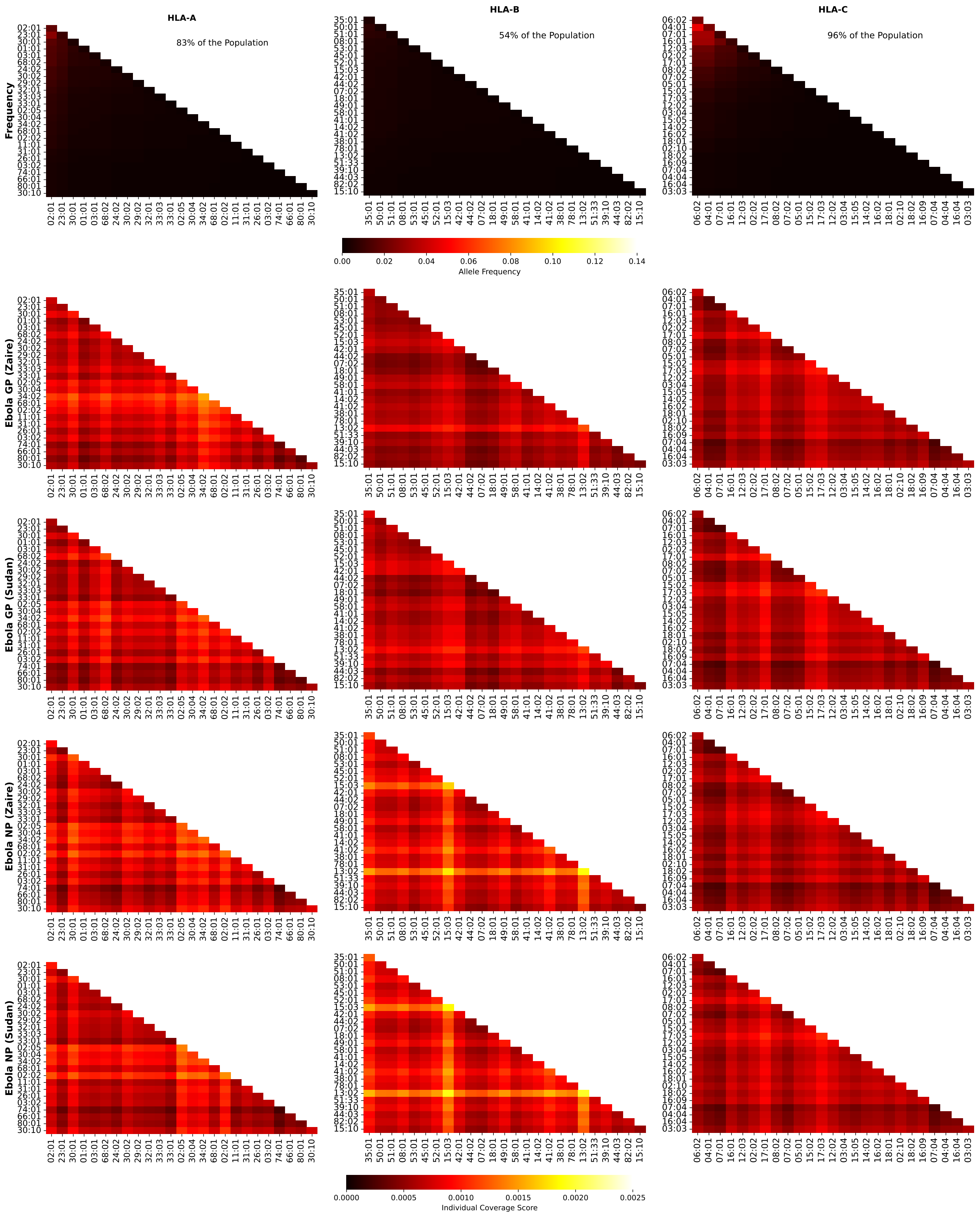


**Figure S52.** Frequencies and Ebola coverage scores for individuals in North Africa. The 1st row corresponds to allele frequencies, the 2nd to GP1 Zaire, the 3rd to GP1 Sudan, the 4th to NP Zaire, and the 5th to NP Sudan. The 1st column is associated with HLA-A alleles, the 2nd to HLA-B, and the 3rd to HLA-C. The sum of the individual frequencies for each allele type is indicated on the panels in the 1st row.

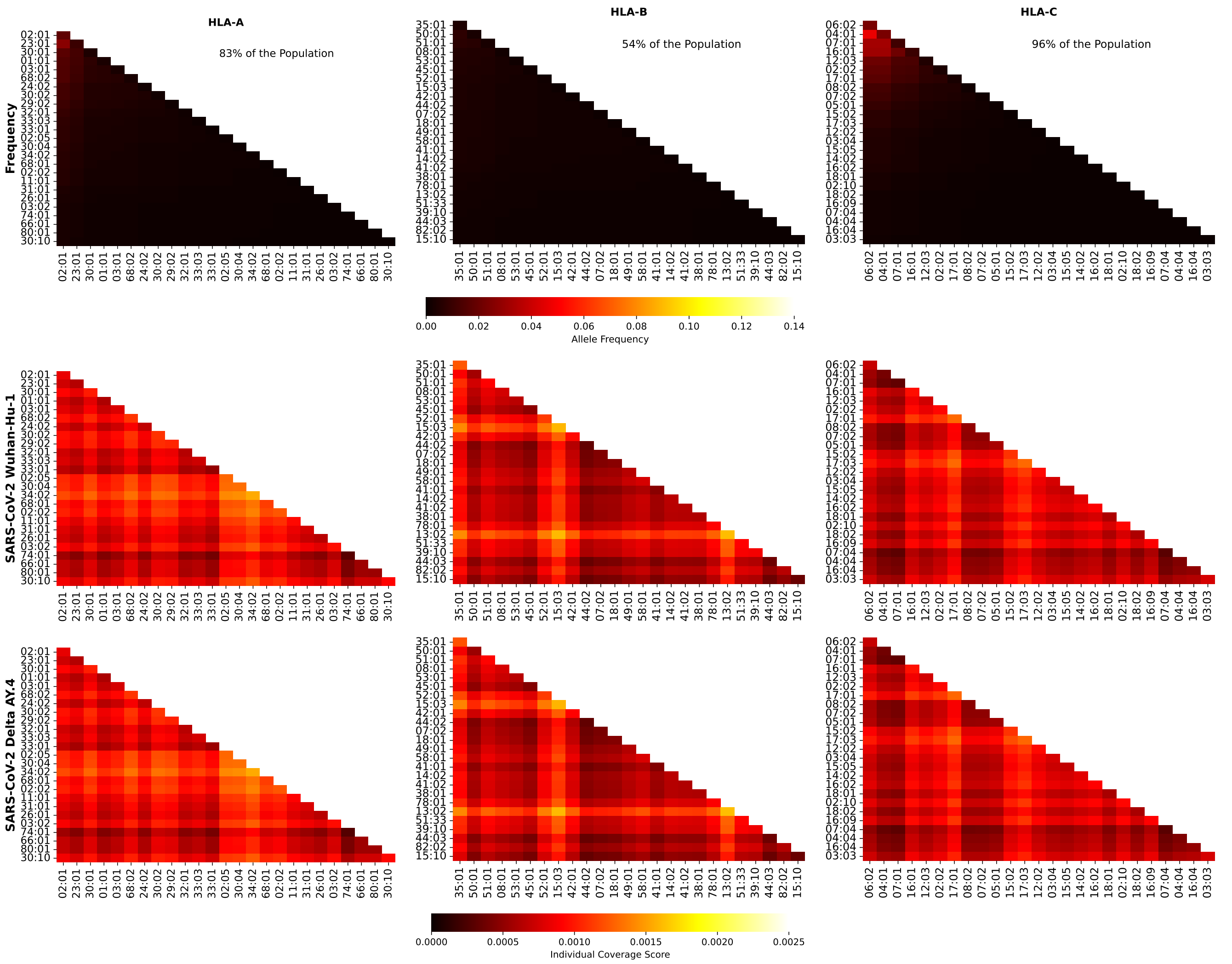


**Figure S53.** Frequencies and SARS-CoV-2 (Wuhan-Hu-1 and Delta AY.4 variants) coverage scores for individuals in North Africa. The 1st row corresponds to allele frequencies, the 2nd to Wuhan-Hu-1, and the 3rd to Delta AY.4. The 1st column is associated with HLA-A alleles, the 2nd to HLA-B, and the 3rd to HLA-C. The sum of the individual frequencies for each allele type is indicated on the panels in the 1st row.

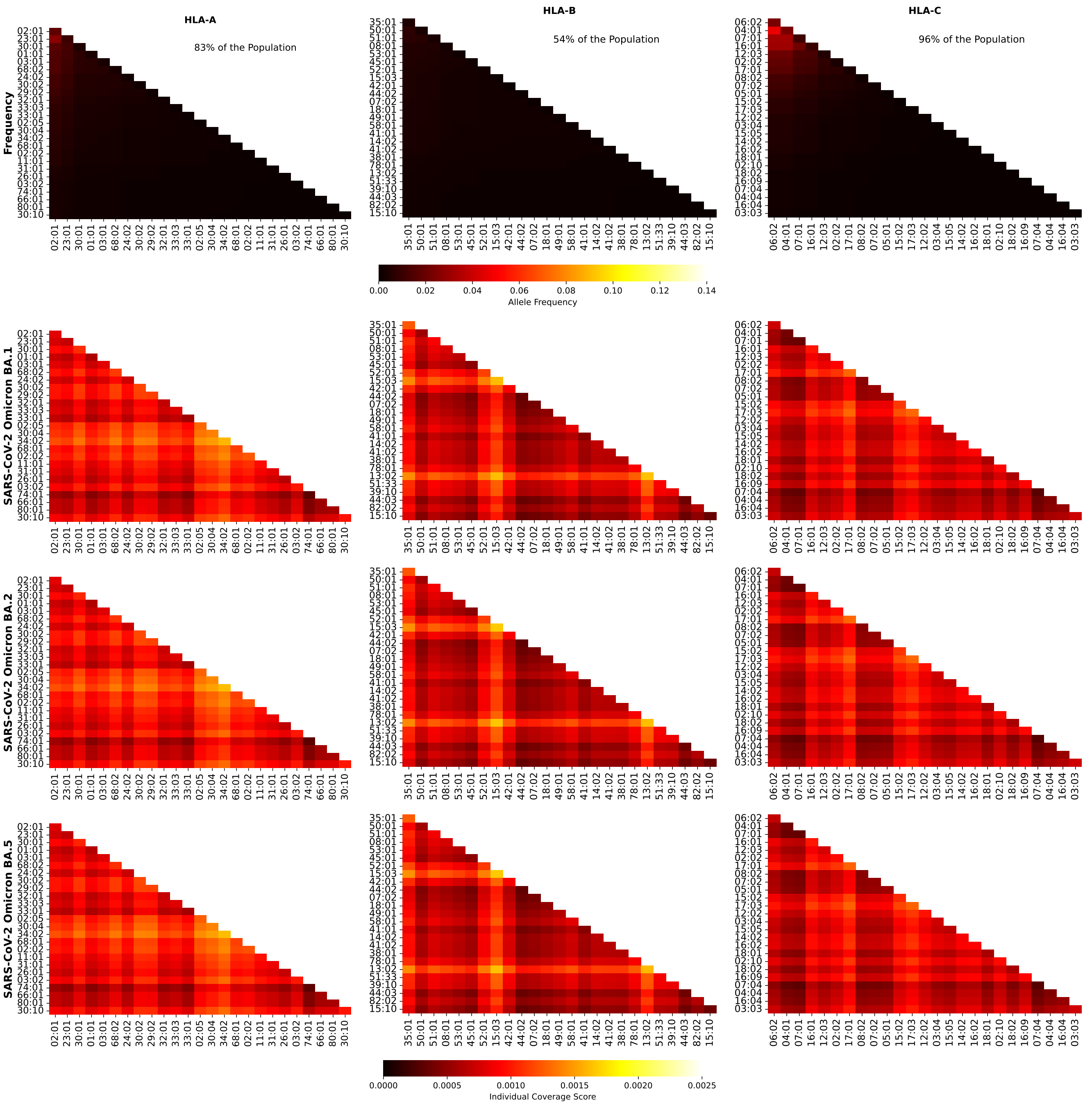


**Figure S54.** Frequencies and SARS-CoV-2 (Omicron variants) coverage scores for individuals in North Africa. The 1st row corresponds to allele frequencies, the 2nd to BA.1, and the 3rd to BA.2, and the 4th to BA.5. The 1st column is associated with HLA-A alleles, the 2nd to HLA-B, and the 3rd to HLA-C. The sum of the individual frequencies for each allele type is indicated on the panels in the 1st row.

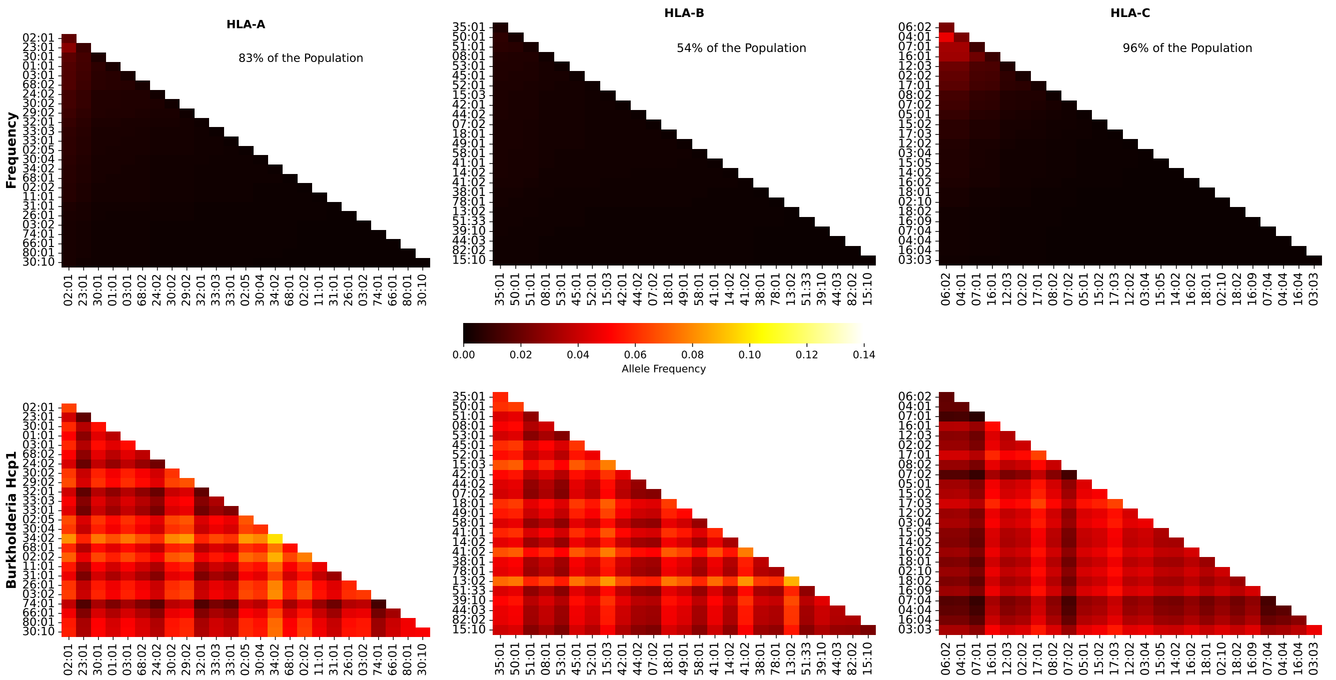


**Figure S55.** Frequencies and Burkholderia coverage scores for individuals in North Africa. The 1st row corresponds to allele frequencies and the 2nd row to Burkholderia coverage score. The 1st column is associated with HLA-A alleles, the 2nd to HLA-B, and the 3rd to HLA-C. The sum of the individual frequencies for each allele type is indicated on the panels in the 1st row.

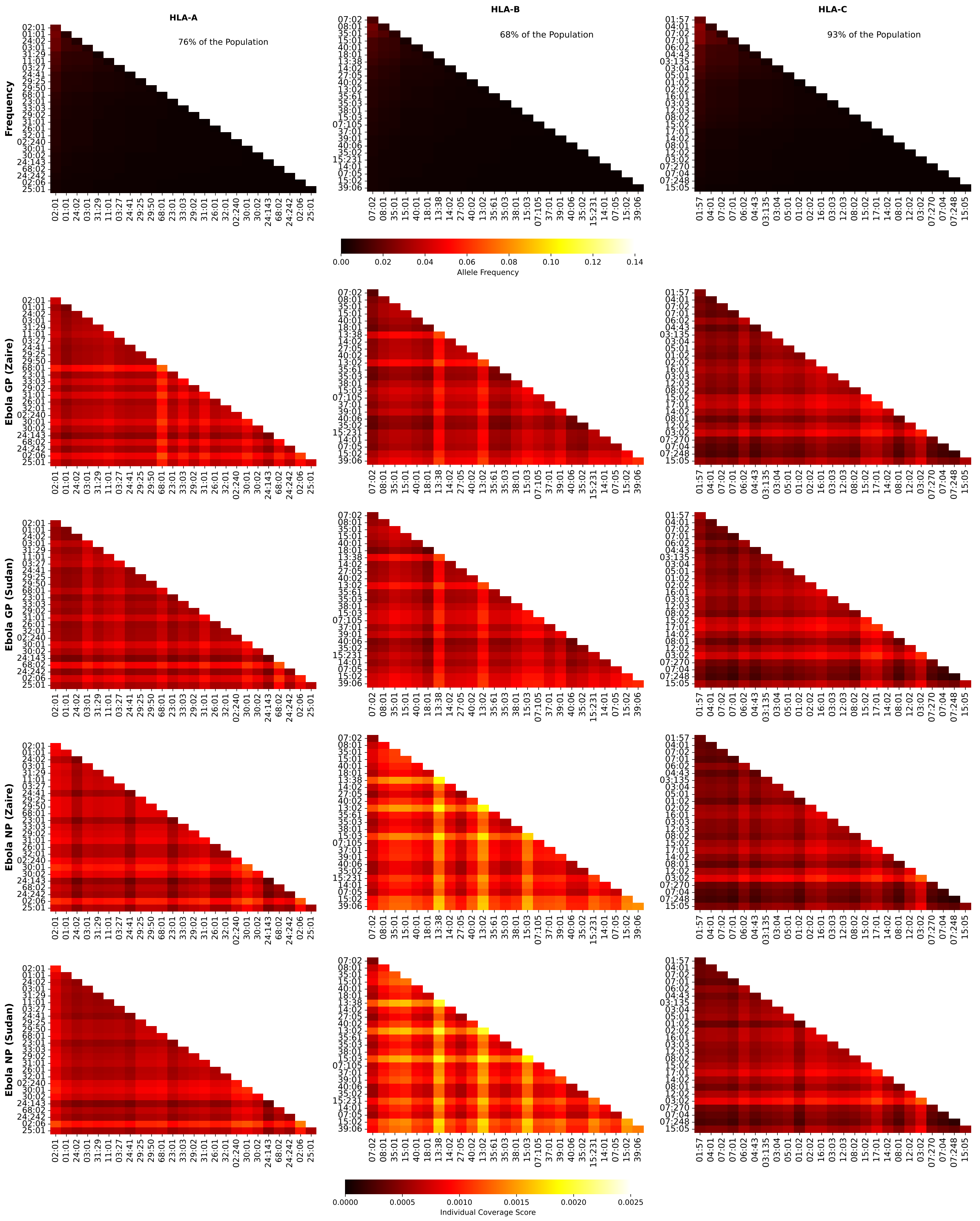

**Figure S56.** Frequencies and Ebola coverage scores for individuals in North America. The 1st row corresponds to allele frequencies, the 2nd to GP1 Zaire, the 3rd to GP1 Sudan, the 4th to NP Zaire, and the 5th to NP Sudan. The 1st column is associated with HLA-A alleles, the 2nd to HLA-B, and the 3rd to HLA-C. The sum of the individual frequencies for each allele type is indicated on the panels in the 1st row.

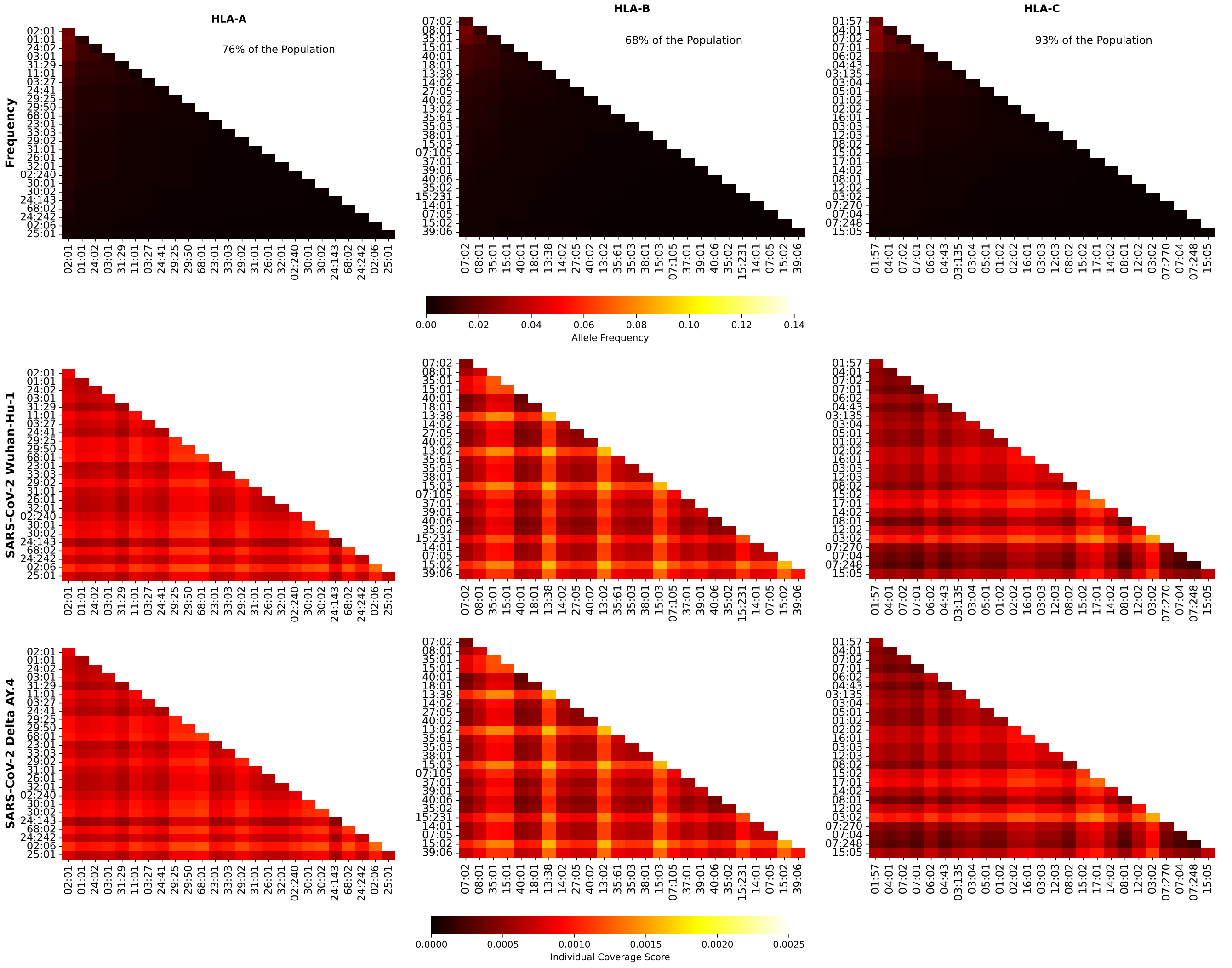


**Figure S57.** Frequencies and SARS-CoV-2 (Wuhan-Hu-1 and Delta AY.4 variants) coverage scores for individuals in North America. The 1st row corresponds to allele frequencies, the 2nd to Wuhan-Hu-1, and the 3rd to Delta AY.4. The 1st column is associated with HLA-A alleles, the 2nd to HLA-B, and the 3rd to HLA-C. The sum of the individual frequencies for each allele type is indicated on the panels in the 1st row.

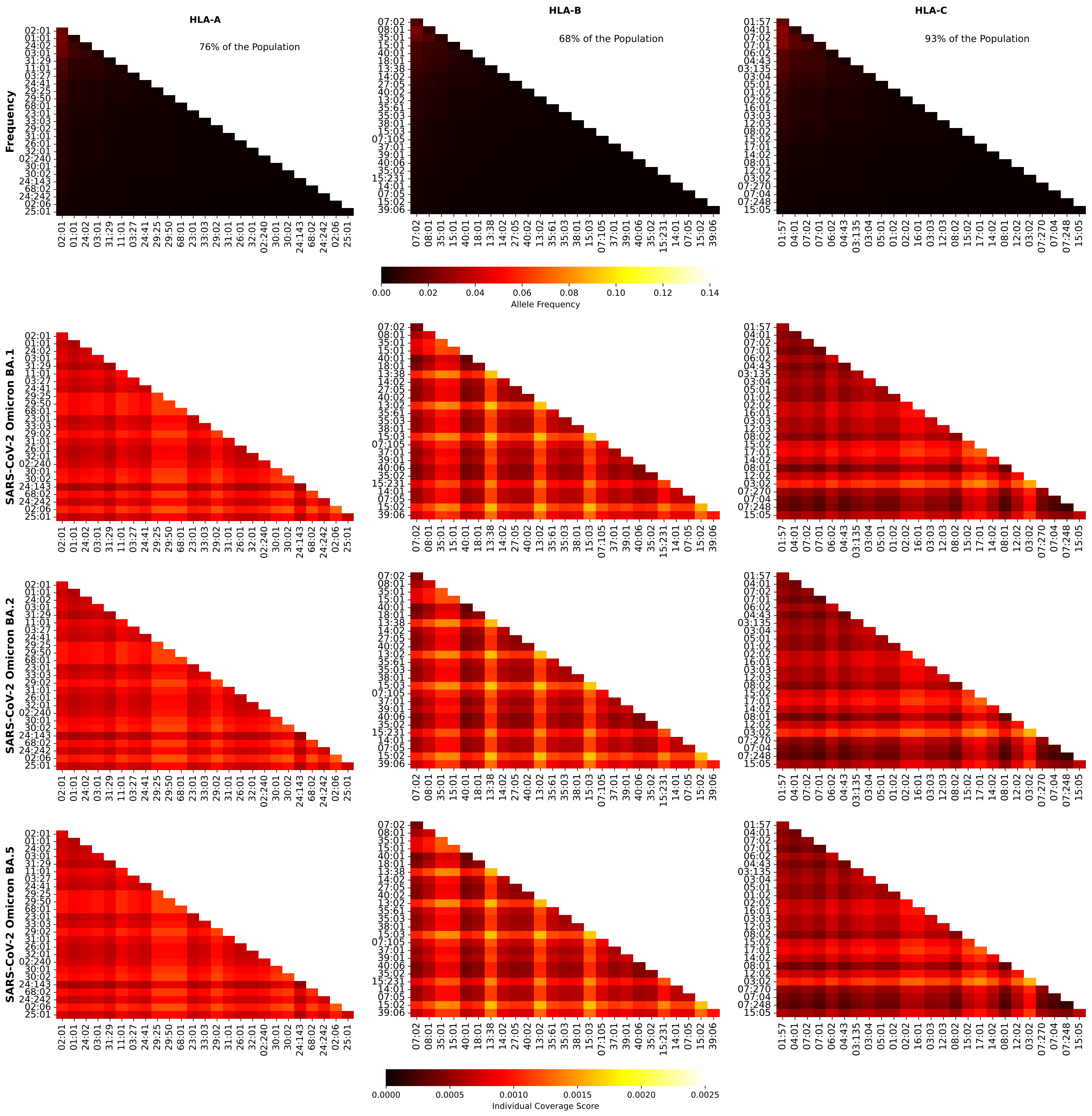


**Figure S58.** Frequencies and SARS-CoV-2 (Omicron variants) coverage scores for individuals in North America. The 1st row corresponds to allele frequencies, the 2nd to BA.1, and the 3rd to BA.2, and the 4th to BA.5. The 1st column is associated with HLA-A alleles, the 2nd to HLA-B, and the 3rd to HLA-C. The sum of the individual frequencies for each allele type is indicated on the panels in the 1st row.

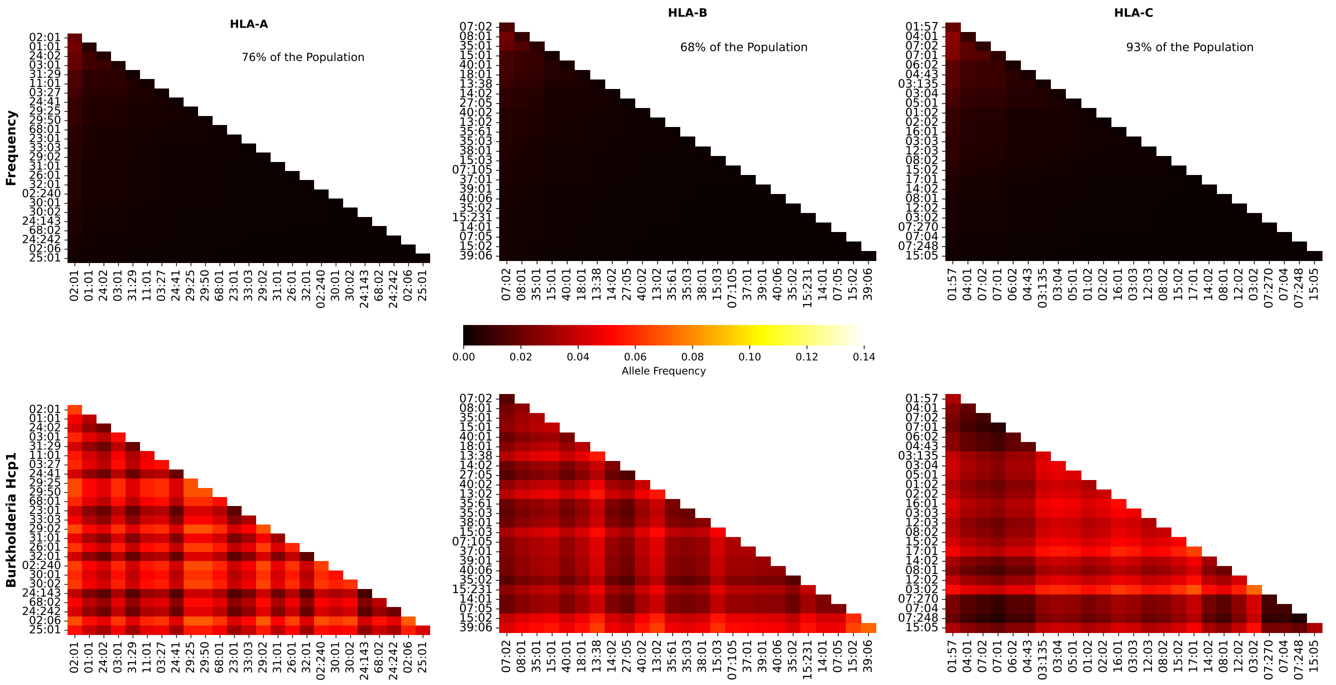


**Figure S59.** Frequencies and Burkholderia coverage scores for individuals in North America. The 1st row corresponds to allele frequencies and the 2nd row to Burkholderia coverage score. The 1st column is associated with HLA-A alleles, the 2nd to HLA-B, and the 3rd to HLA-C. The sum of the individual frequencies for each allele type is indicated on the panels in the 1st row.

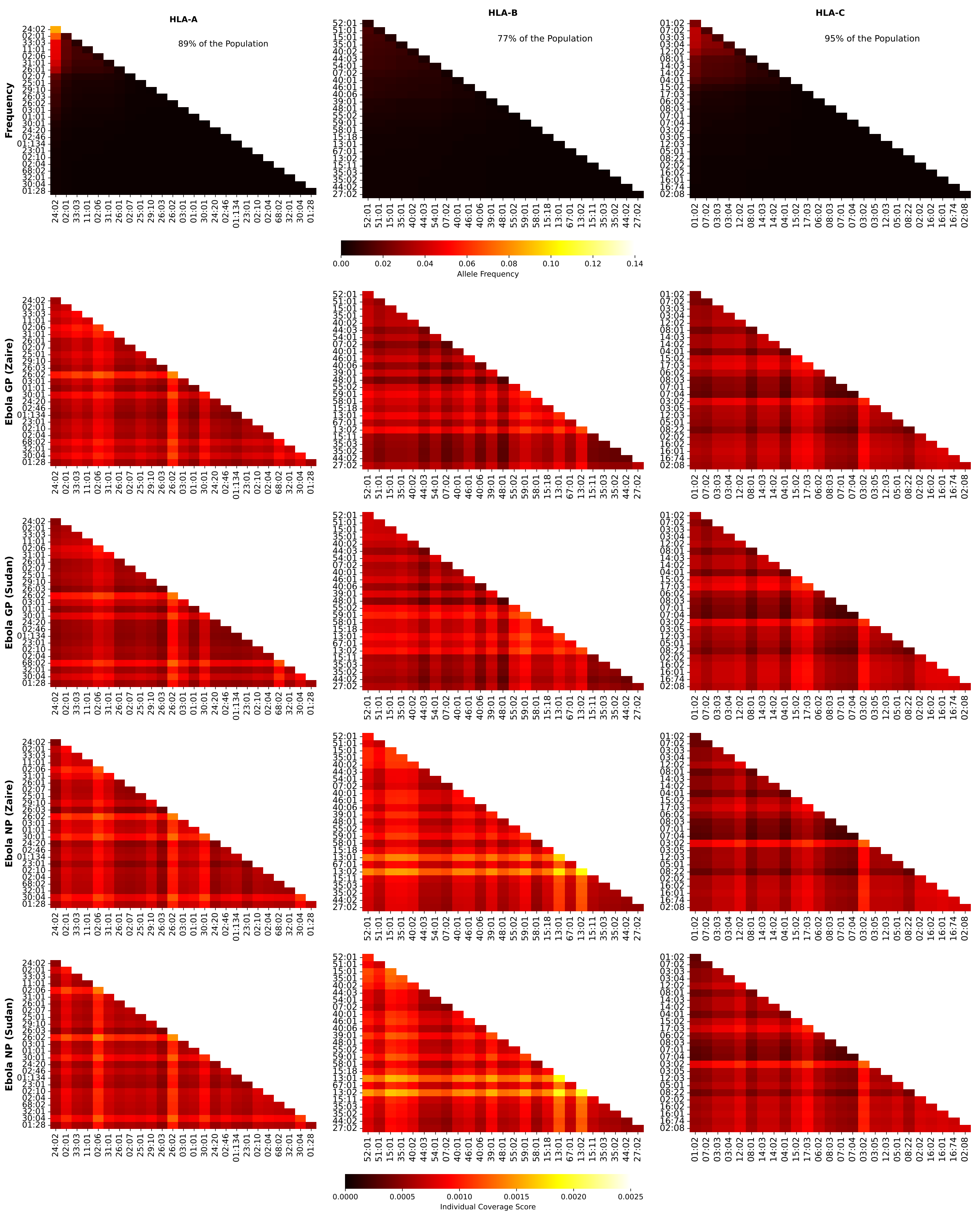


**Figure S60.** Frequencies and Ebola coverage scores for individuals in Northeast Asia. The 1st row corresponds to allele frequencies, the 2nd to GP1 Zaire, the 3rd to GP1 Sudan, the 4th to NP Zaire, and the 5th to NP Sudan. The 1st column is associated with HLA-A alleles, the 2nd to HLA-B, and the 3rd to HLA-C. The sum of the individual frequencies for each allele type is indicated on the panels in the 1st row.

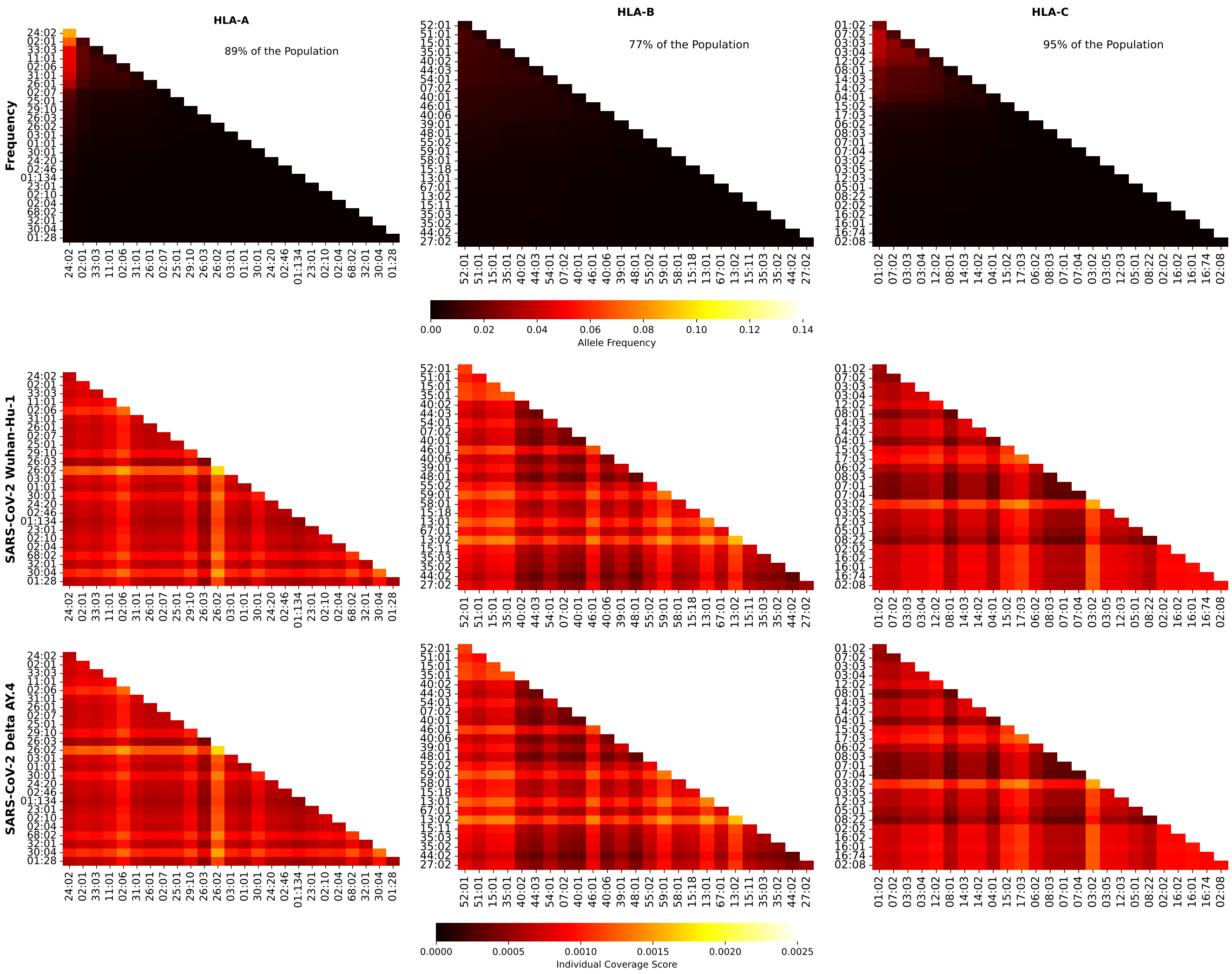


**Figure S61.** Frequencies and SARS-CoV-2 (Wuhan-Hu-1 and Delta AY.4 variants) coverage scores for individuals in Northeast Asia. The 1st row corresponds to allele frequencies, the 2nd to Wuhan-Hu-1, and the 3rd to Delta AY.4. The 1st column is associated with HLA-A alleles, the 2nd to HLA-B, and the 3rd to HLA-C. The sum of the individual frequencies for each allele type is indicated on the panels in the 1st row.

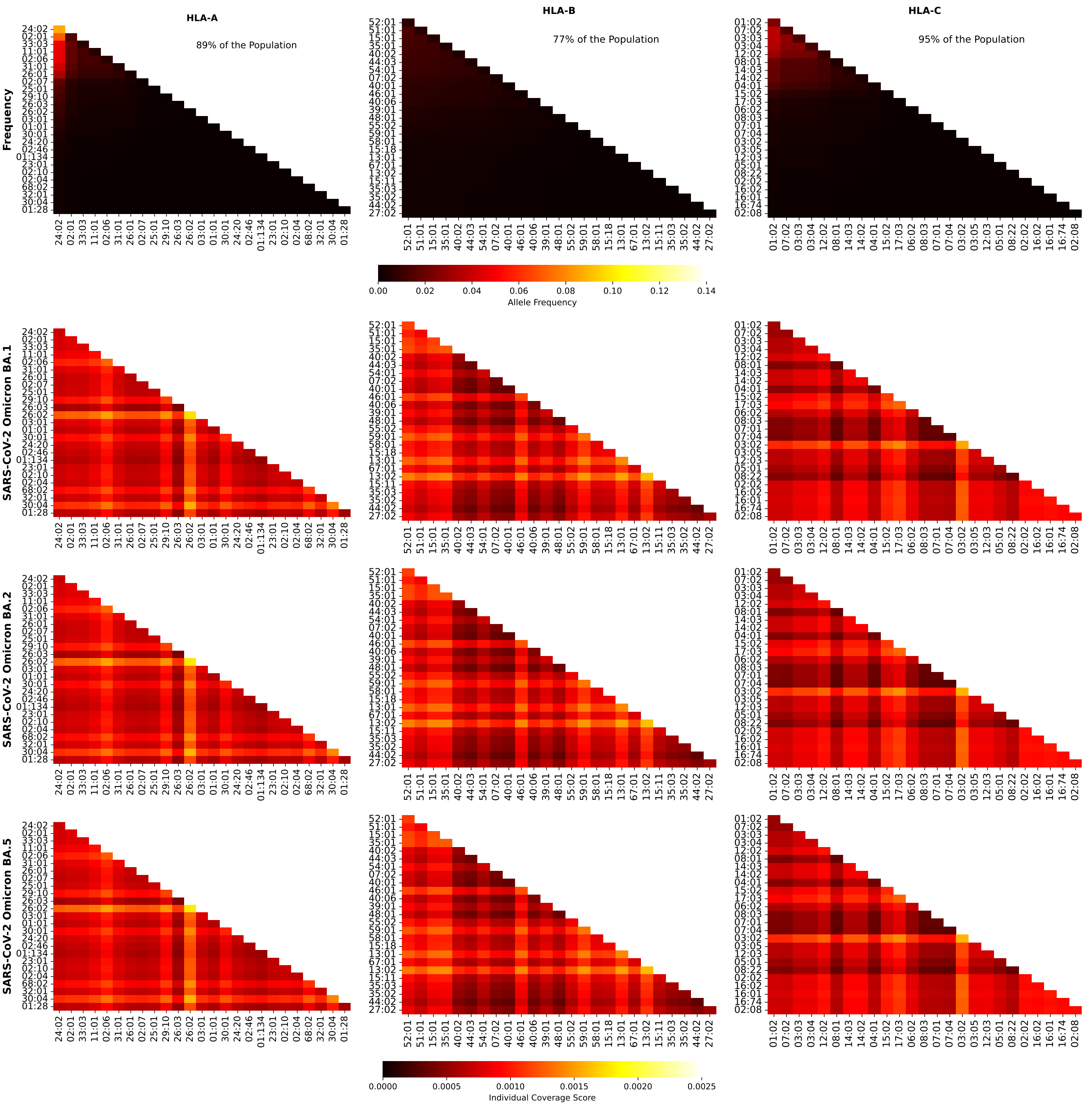


**Figure S62.** Frequencies and SARS-CoV-2 (Omicron variants) coverage scores for individuals in Northeast Asia. The 1st row corresponds to allele frequencies, the 2nd to BA.1, and the 3rd to BA.2, and the 4th to BA.5. The 1st column is associated with HLA-A alleles, the 2nd to HLA-B, and the 3rd to HLA-C. The sum of the individual frequencies for each allele type is indicated on the panels in the 1st row.

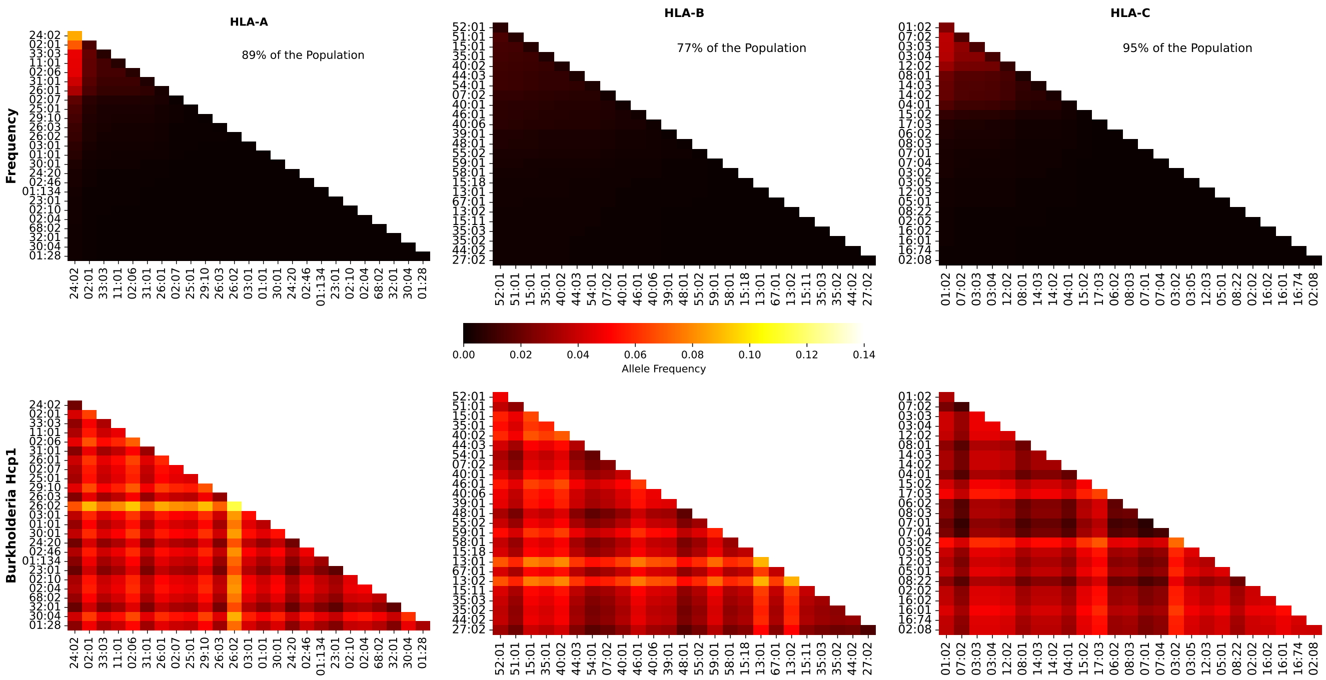


**Figure S63.** Frequencies and Burkholderia coverage scores for individuals in Northeast Asia. The 1st row corresponds to allele frequencies and the 2nd row to Burkholderia coverage score. The 1st column is associated with HLA-A alleles, the 2nd to HLA-B, and the 3rd to HLA-C. The sum of the individual frequencies for each allele type is indicated on the panels in the 1st row.

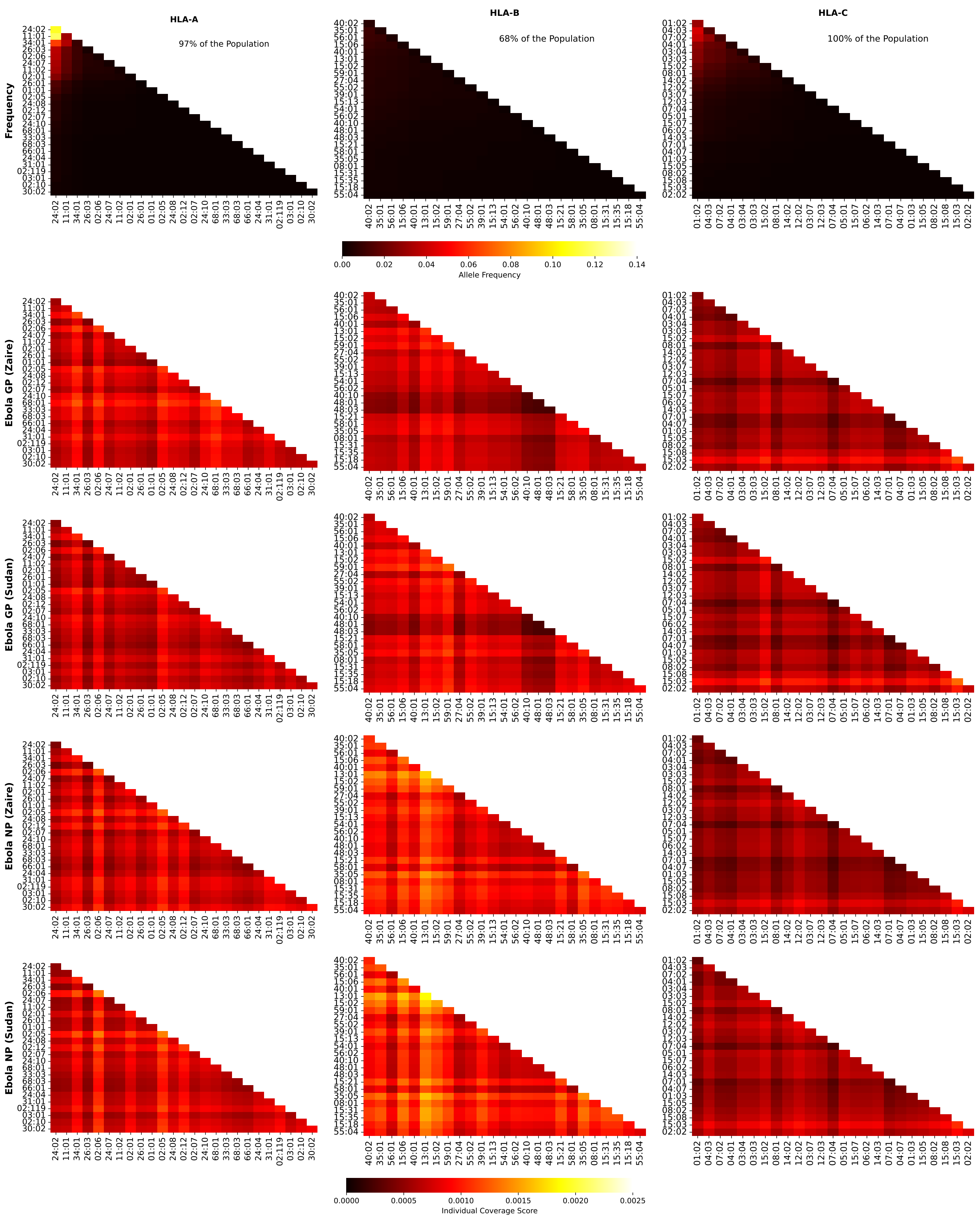


**Figure S64.** Frequencies and Ebola coverage scores for individuals in Oceania. The 1st row corresponds to allele frequencies, the 2nd to GP1 Zaire, the 3rd to GP1 Sudan, the 4th to NP Zaire, and the 5th to NP Sudan. The 1st column is associated with HLA-A alleles, the 2nd to HLA-B, and the 3rd to HLA-C. The sum of the individual frequencies for each allele type is indicated on the panels in the 1st row.

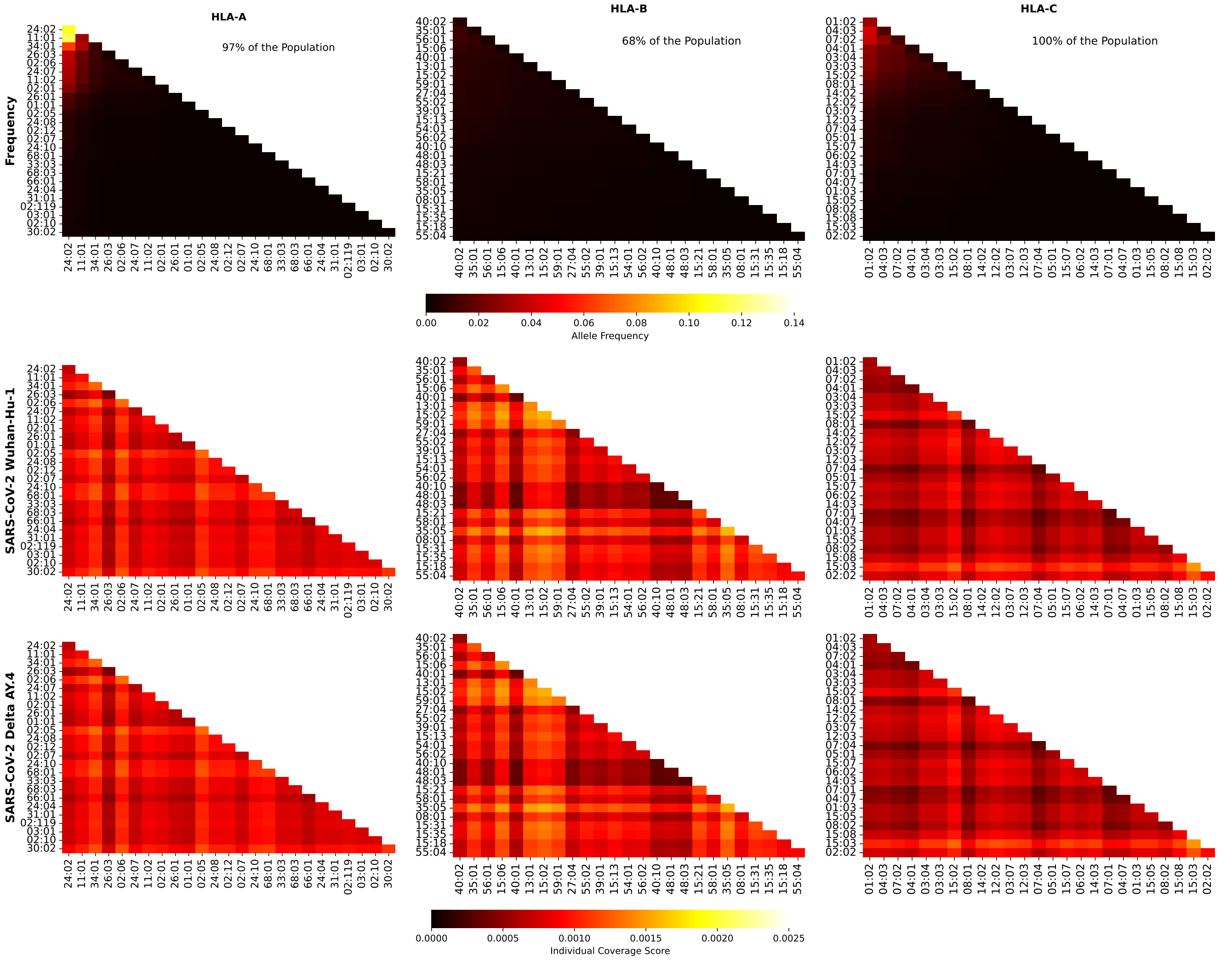


**Figure S65.** Frequencies and SARS-CoV-2 (Wuhan-Hu-1 and Delta AY.4 variants) coverage scores for individuals in Oceania. The 1st row corresponds to allele frequencies, the 2nd to Wuhan-Hu-1, and the 3rd to Delta AY.4. The 1st column is associated with HLA-A alleles, the 2nd to HLA-B, and the 3rd to HLA-C. The sum of the individual frequencies for each allele type is indicated on the panels in the 1st row.

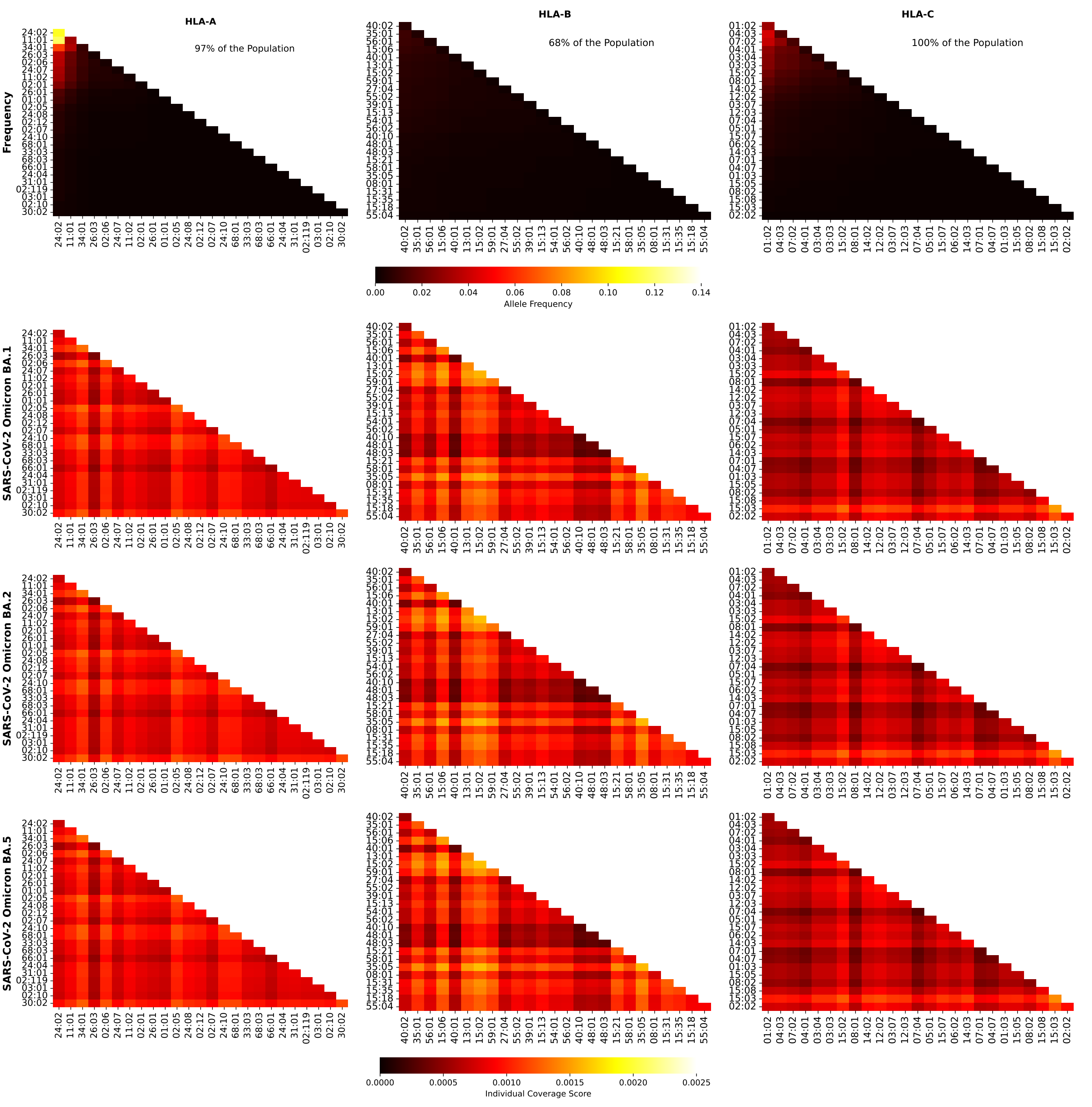


**Figure S66.** Frequencies and SARS-CoV-2 (Omicron variants) coverage scores for individuals in Oceania. The 1st row corresponds to allele frequencies, the 2nd to BA.1, and the 3rd to BA.2, and the 4th to BA.5. The 1st column is associated with HLA-A alleles, the 2nd to HLA-B, and the 3rd to HLA-C. The sum of the individual frequencies for each allele type is indicated on the panels in the 1st row.

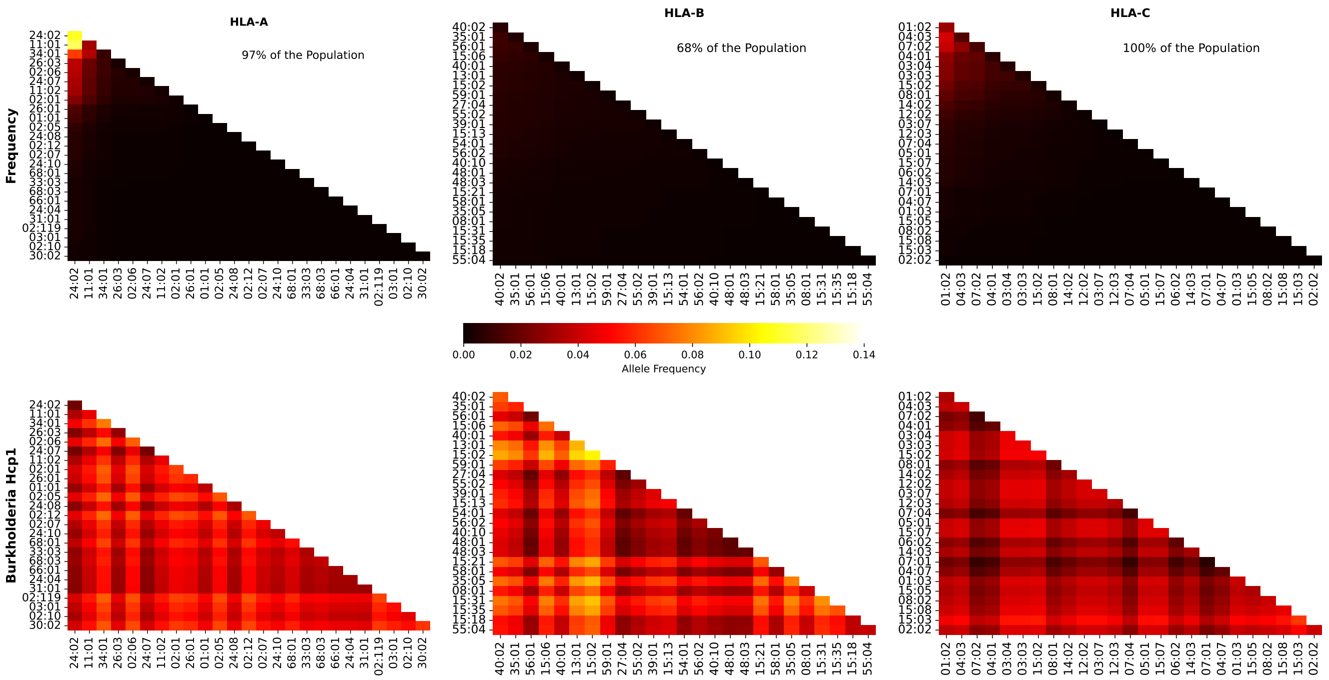


**Figure S67.** Frequencies and Burkholderia coverage scores for individuals in Oceania. The 1st row corresponds to allele frequencies and the 2nd row to Burkholderia coverage score. The 1st column is associated with HLA-A alleles, the 2nd to HLA-B, and the 3rd to HLA-C. The sum of the individual frequencies for each allele type is indicated on the panels in the 1st row.

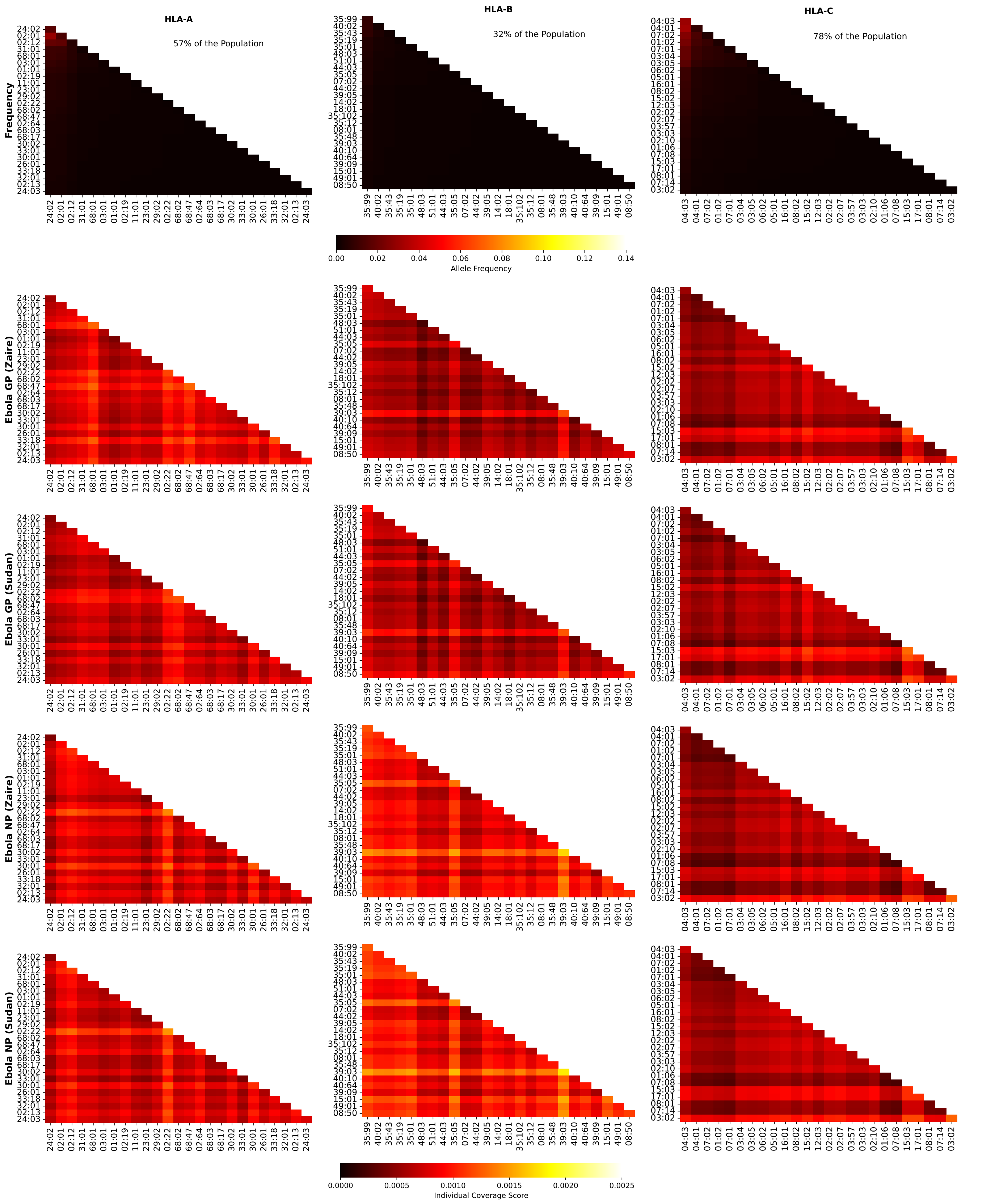


**Figure S68.** Frequencies and Ebola coverage scores for individuals in South and Central America. The 1st row corresponds to allele frequencies, the 2nd to GP1 Zaire, the 3rd to GP1 Sudan, the 4th to NP Zaire, and the 5th to NP Sudan. The 1st column is associated with HLA-A alleles, the 2nd to HLA-B, and the 3rd to HLA-C. The sum of the individual frequencies for each allele type is indicated on the panels in the 1st row.

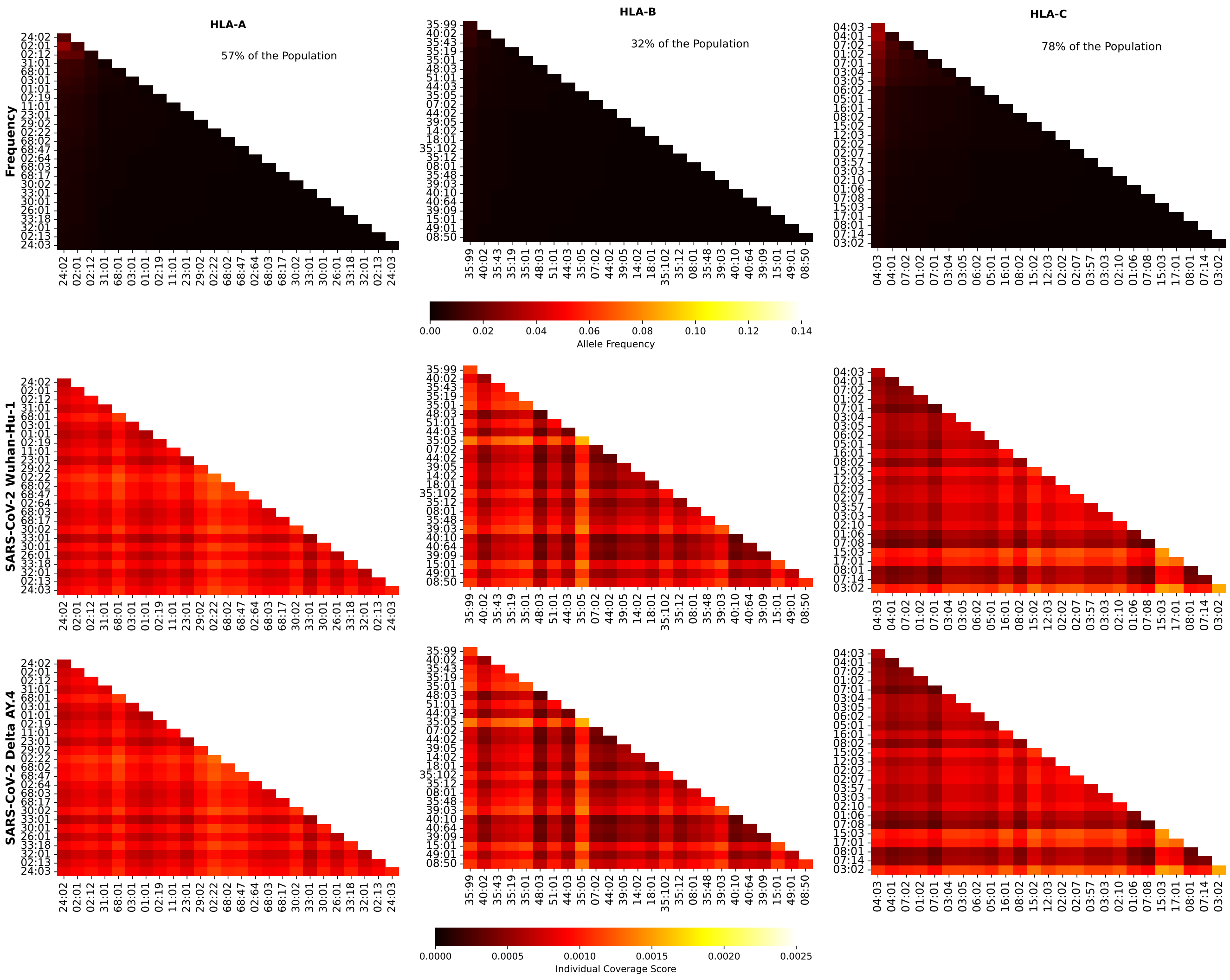


**Figure S69.** Frequencies and SARS-CoV-2 (Wuhan-Hu-1 and Delta AY.4 variants) coverage scores for individuals in South and Central America. The 1st row corresponds to allele frequencies, the 2nd to Wuhan-Hu-1, and the 3rd to Delta AY.4. The 1st column is associated with HLA-A alleles, the 2nd to HLA-B, and the 3rd to HLA-C. The sum of the individual frequencies for each allele type is indicated on the panels in the 1st row.

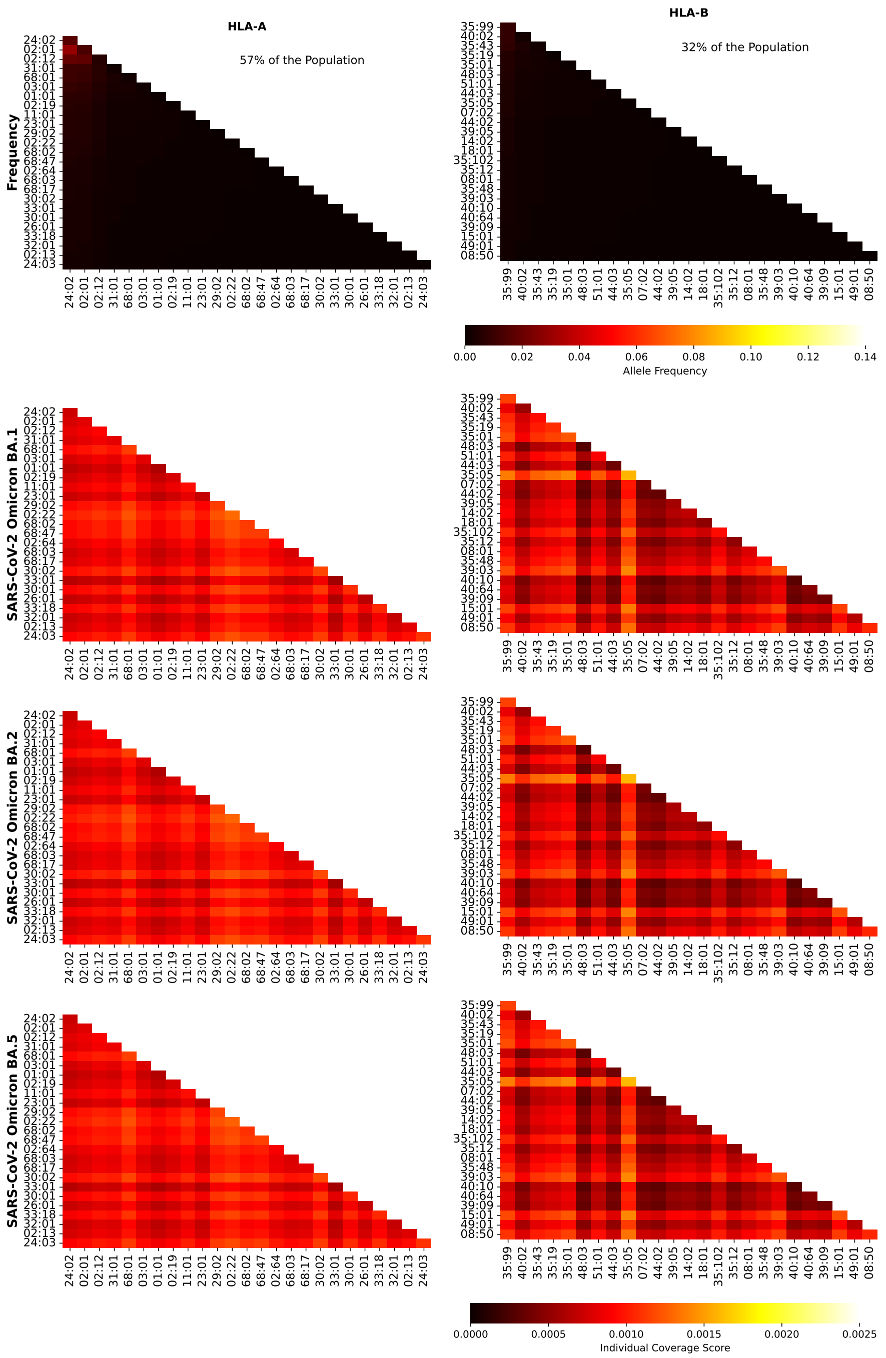


**Figure S70.** Frequencies and SARS-CoV-2 (Omicron variants) coverage scores for individuals in South and Central America. The 1st row corresponds to allele frequencies, the 2nd to BA.1, and the 3rd to BA.2, and the 4th to BA.5. The 1st column is associated with HLA-A alleles, the 2nd to HLA-B, and the 3rd to HLA-C. The sum of the individual frequencies for each allele type is indicated on the panels in the 1st row.

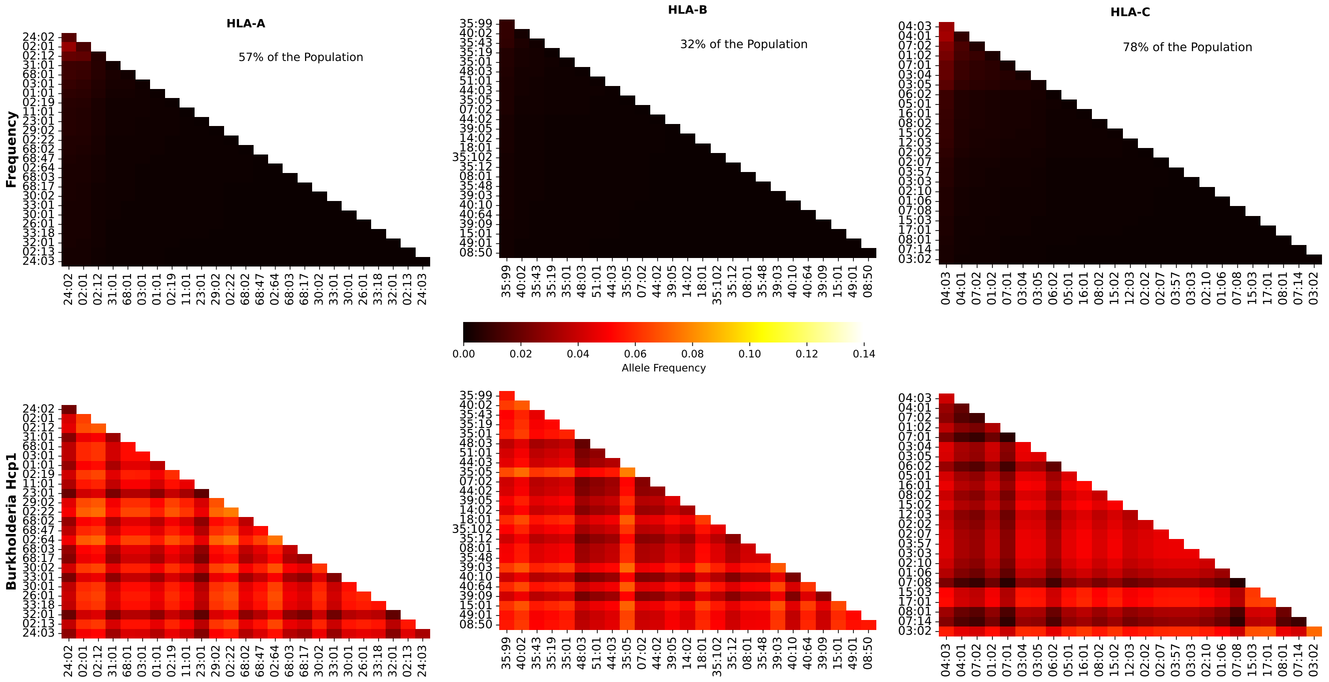


**Figure S71.** Frequencies and Burkholderia coverage scores for individuals in South and Central America. The 1st row corresponds to allele frequencies and the 2nd row to Burkholderia coverage score. The 1st column is associated with HLA-A alleles, the 2nd to HLA-B, and the 3rd to HLA-C. The sum of the individual frequencies for each allele type is indicated on the panels in the 1st row.

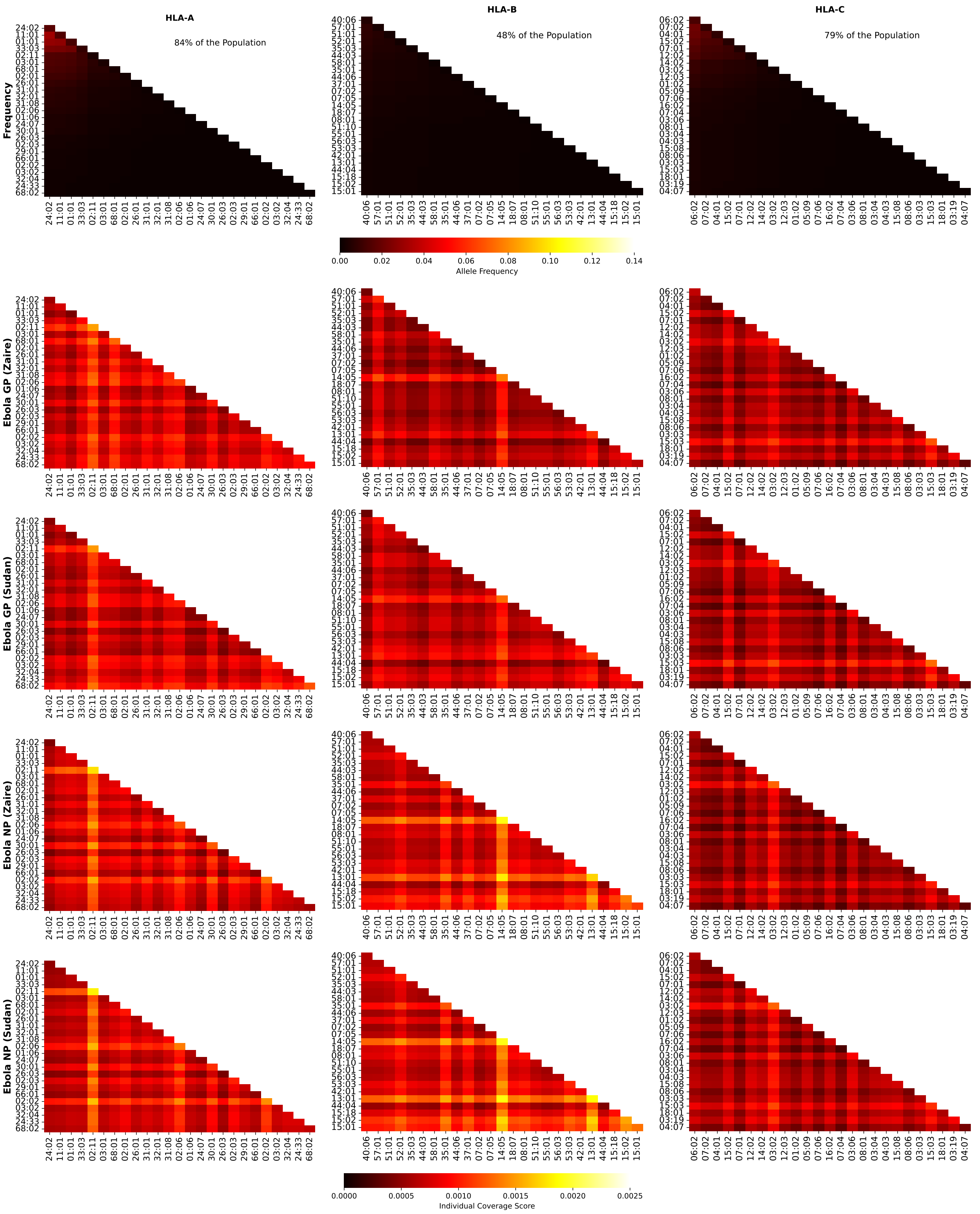


**Figure S72.** Frequencies and Ebola coverage scores for individuals in South Asia. The 1st row corresponds to allele frequencies, the 2nd to GP1 Zaire, the 3rd to GP1 Sudan, the 4th to NP Zaire, and the 5th to NP Sudan. The 1st column is associated with HLA-A alleles, the 2nd to HLA-B, and the 3rd to HLA-C. The sum of the individual frequencies for each allele type is indicated on the panels in the 1st row.

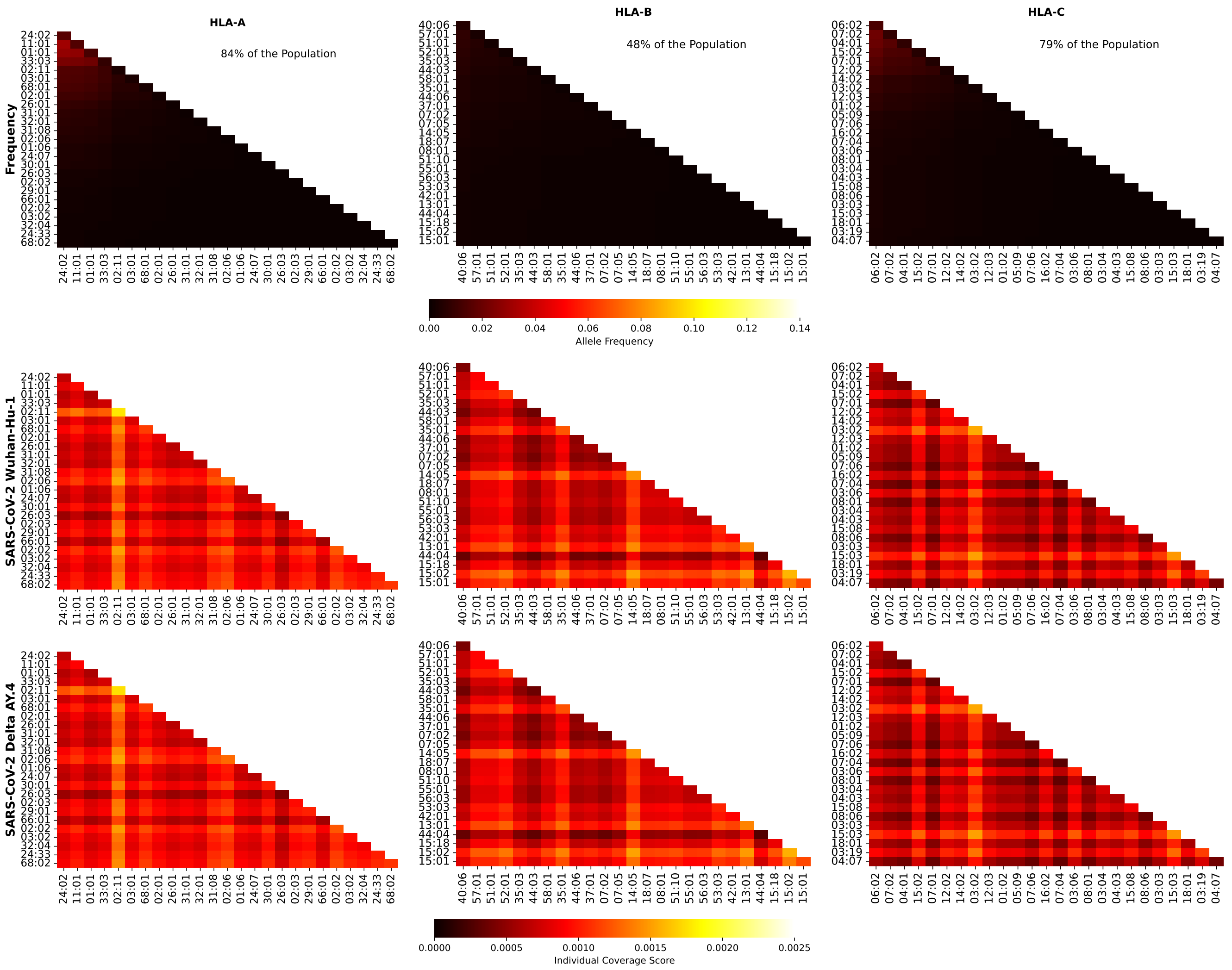


**Figure S73.** Frequencies and SARS-CoV-2 (Wuhan-Hu-1 and Delta AY.4 variants) coverage scores for individuals in South Asia. The 1st row corresponds to allele frequencies, the 2nd to Wuhan-Hu-1, and the 3rd to Delta AY.4. The 1st column is associated with HLA-A alleles, the 2nd to HLA-B, and the 3rd to HLA-C. The sum of the individual frequencies for each allele type is indicated on the panels in the 1st row.

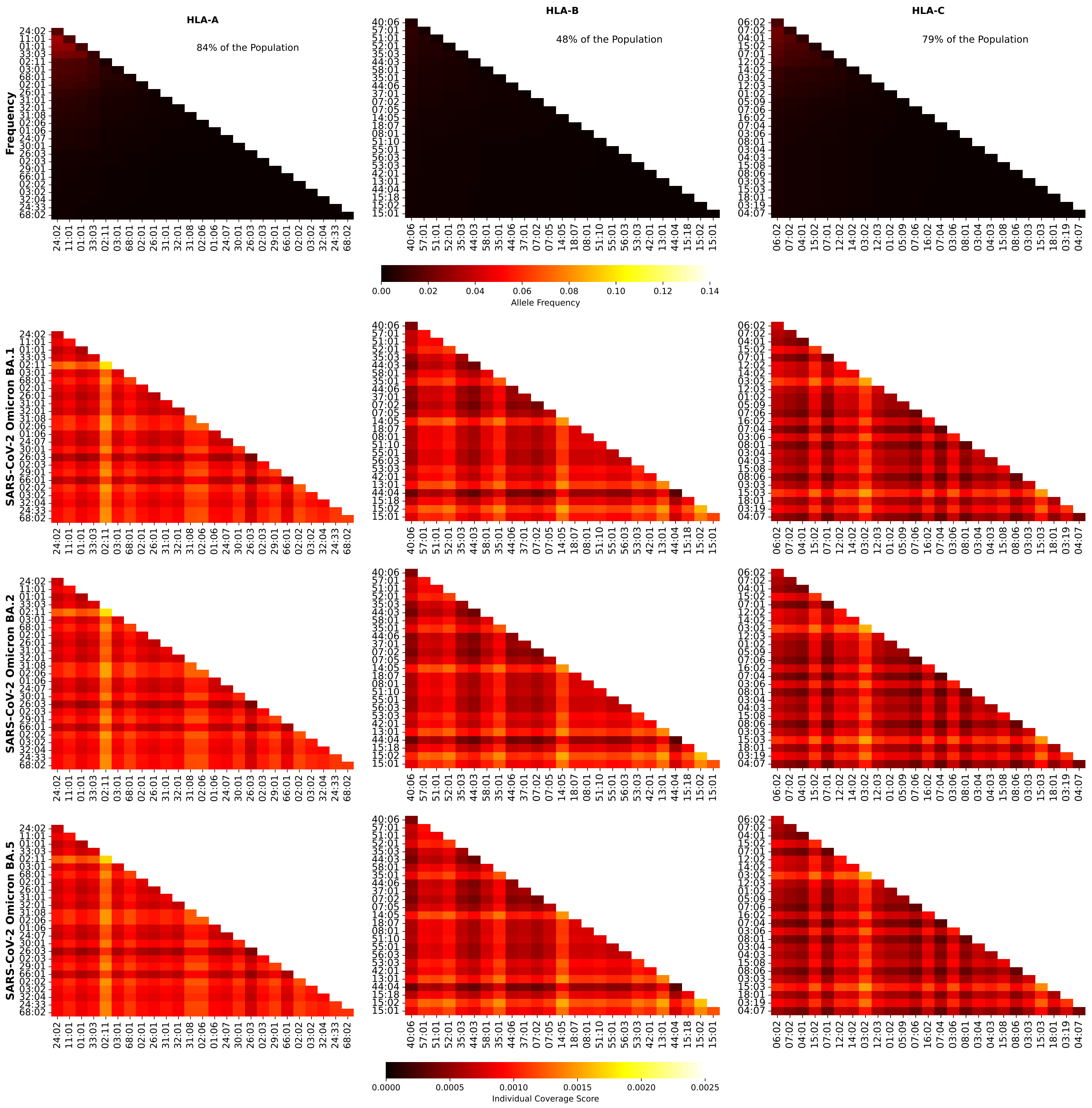


**Figure S74.** Frequencies and SARS-CoV-2 (Omicron variants) coverage scores for individuals in South Asia. The 1st row corresponds to allele frequencies, the 2nd to BA.1, and the 3rd to BA.2, and the 4th to BA.5. The 1st column is associated with HLA-A alleles, the 2nd to HLA-B, and the 3rd to HLA-C. The sum of the individual frequencies for each allele type is indicated on the panels in the 1st row.

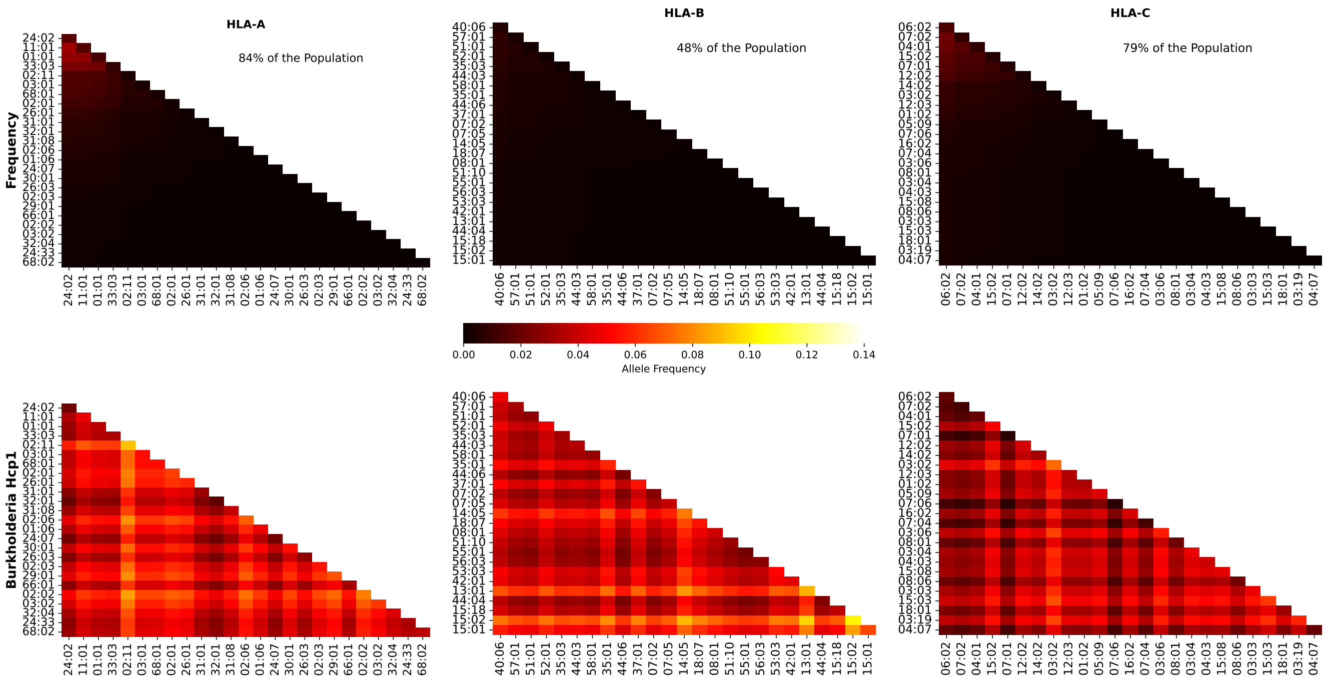


**Figure S75.** Frequencies and Burkholderia coverage scores for individuals in South Asia. The 1st row corresponds to allele frequencies and the 2nd row to Burkholderia coverage score. The 1st column is associated with HLA-A alleles, the 2nd to HLA-B, and the 3rd to HLA-C. The sum of the individual frequencies for each allele type is indicated on the panels in the 1st row.

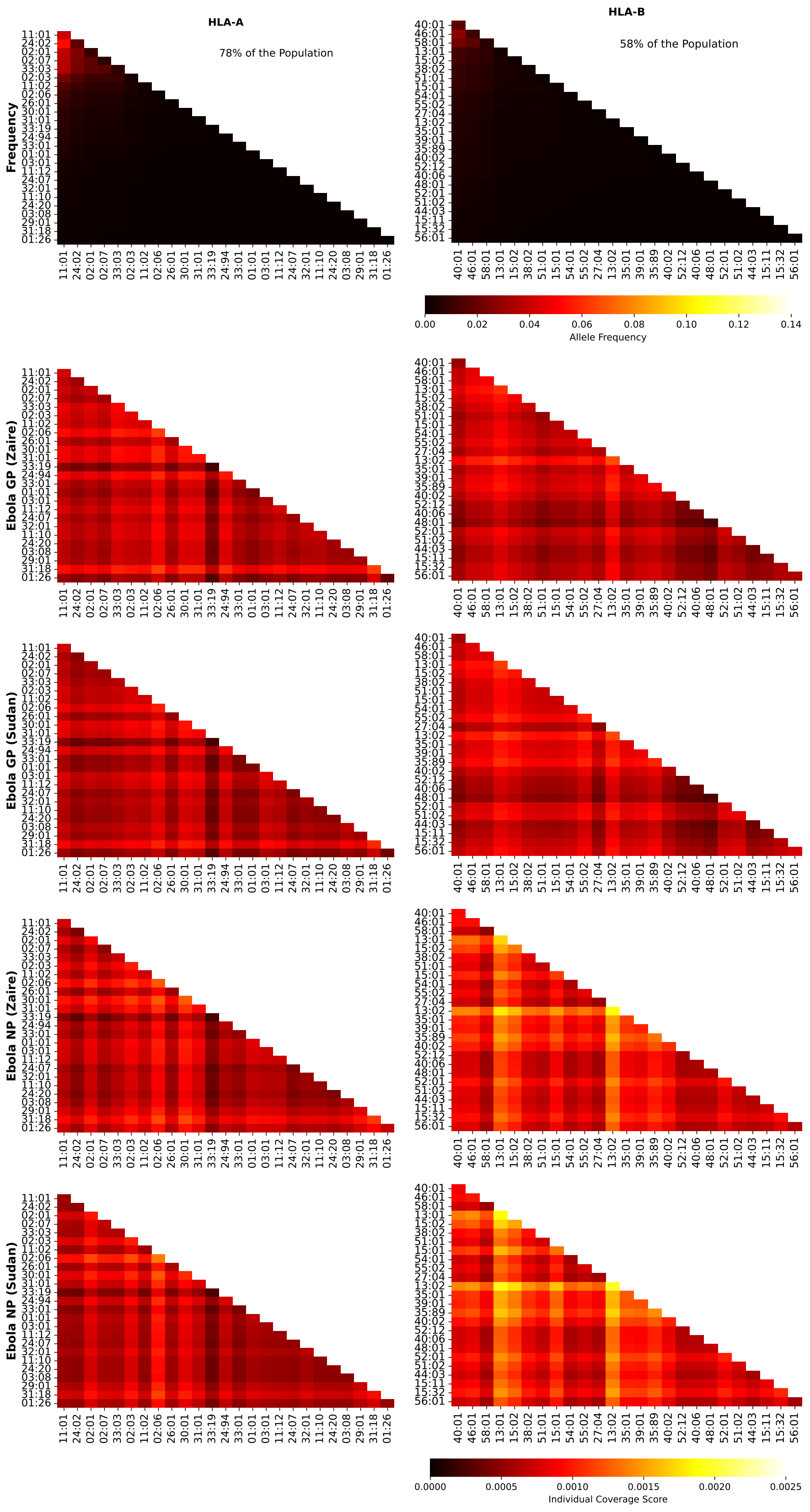


**Figure S76.** Frequencies and Ebola coverage scores for individuals in Southeast Asia. The 1st row corresponds to allele frequencies, the 2nd to GP1 Zaire, the 3rd to GP1 Sudan, the 4th to NP Zaire, and the 5th to NP Sudan. The 1st column is associated with HLA-A alleles, the 2nd to HLA-B, and the 3rd to HLA-C. The sum of the individual frequencies for each allele type is indicated on the panels in the 1st row.

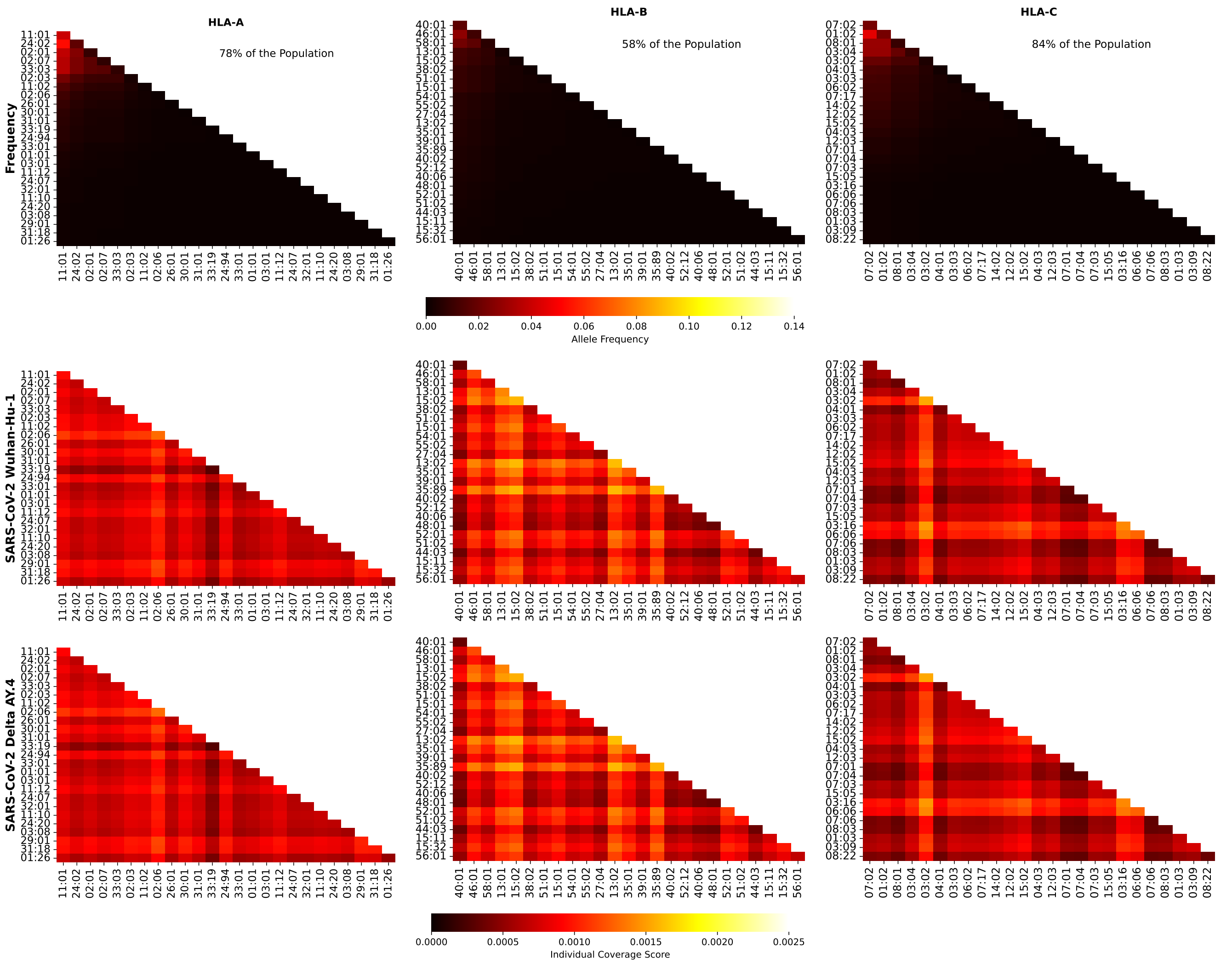


**Figure S77.** Frequencies and SARS-CoV-2 (Wuhan-Hu-1 and Delta AY.4 variants) coverage scores for individuals in Southeast Asia. The 1st row corresponds to allele frequencies, the 2nd to Wuhan-Hu-1, and the 3rd to Delta AY.4. The 1st column is associated with HLA-A alleles, the 2nd to HLA-B, and the 3rd to HLA-C. The sum of the individual frequencies for each allele type is indicated on the panels in the 1st row.

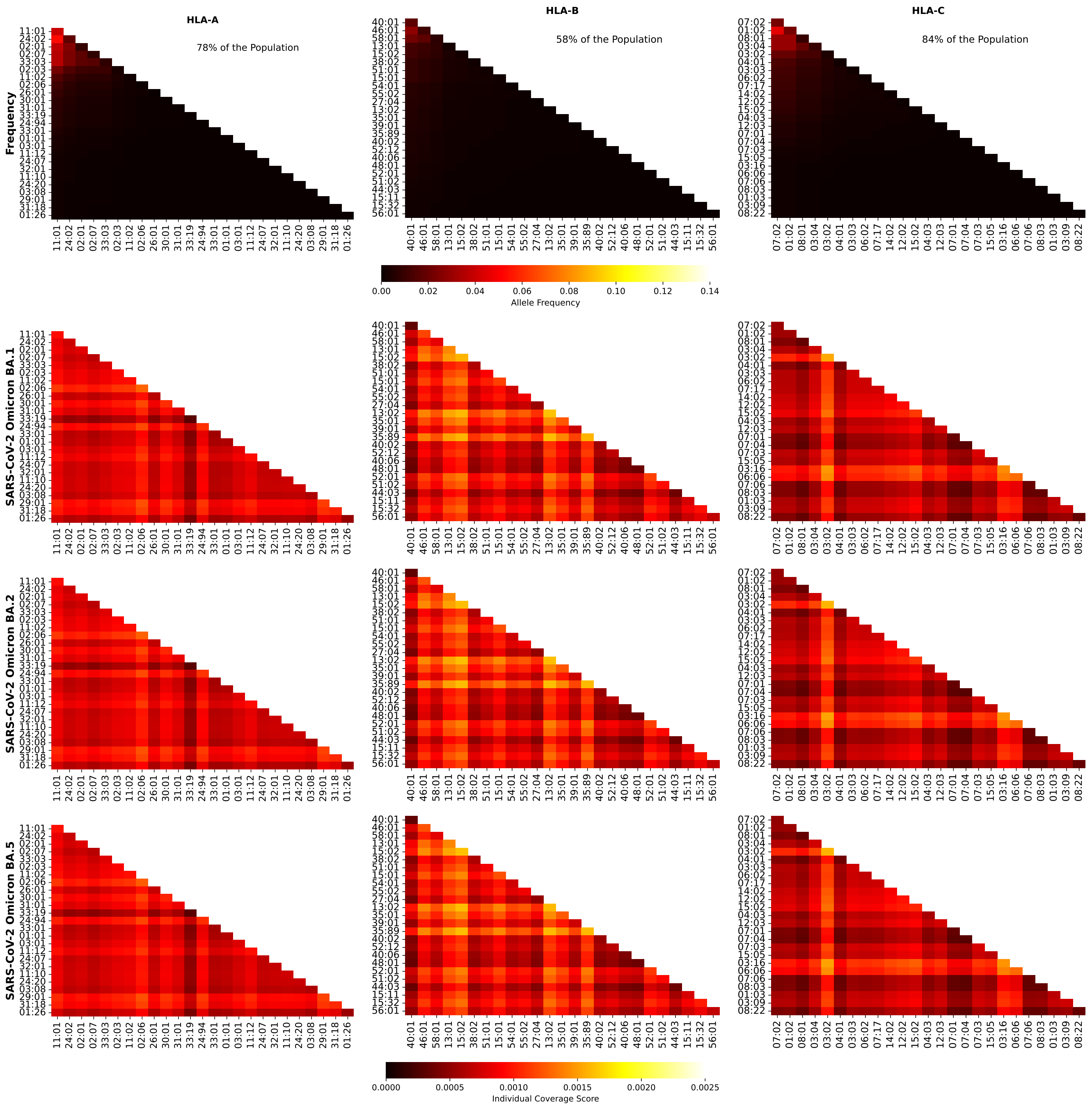


**Figure S78.** Frequencies and SARS-CoV-2 (Omicron variants) coverage scores for individuals in Southeast Asia. The 1st row corresponds to allele frequencies, the 2nd to BA.1, and the 3rd to BA.2, and the 4th to BA.5. The 1st column is associated with HLA-A alleles, the 2nd to HLA-B, and the 3rd to HLA-C. The sum of the individual frequencies for each allele type is indicated on the panels in the 1st row.

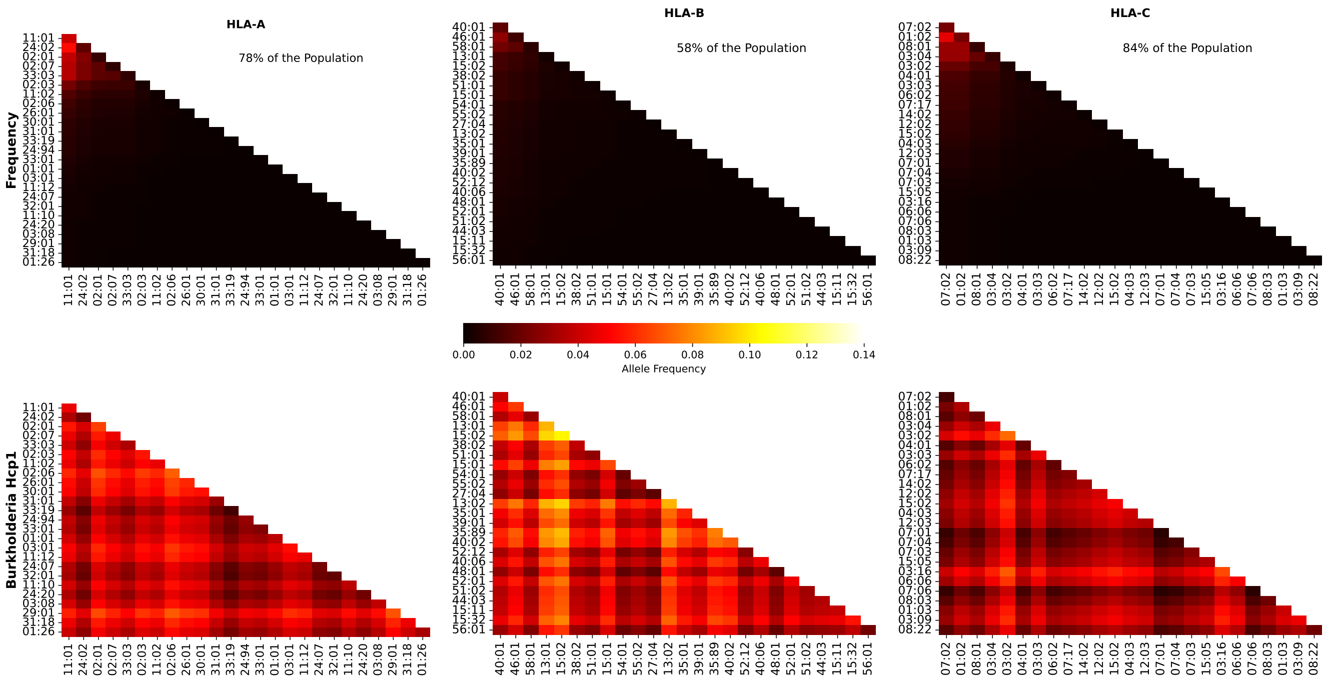


**Figure S79.** Frequencies and Burkholderia coverage scores for individuals in Southeast Asia. The 1st row corresponds to allele frequencies and the 2nd row to Burkholderia coverage score. The 1st column is associated with HLA-A alleles, the 2nd to HLA-B, and the 3rd to HLA-C. The sum of the individual frequencies for each allele type is indicated on the panels in the 1st row.

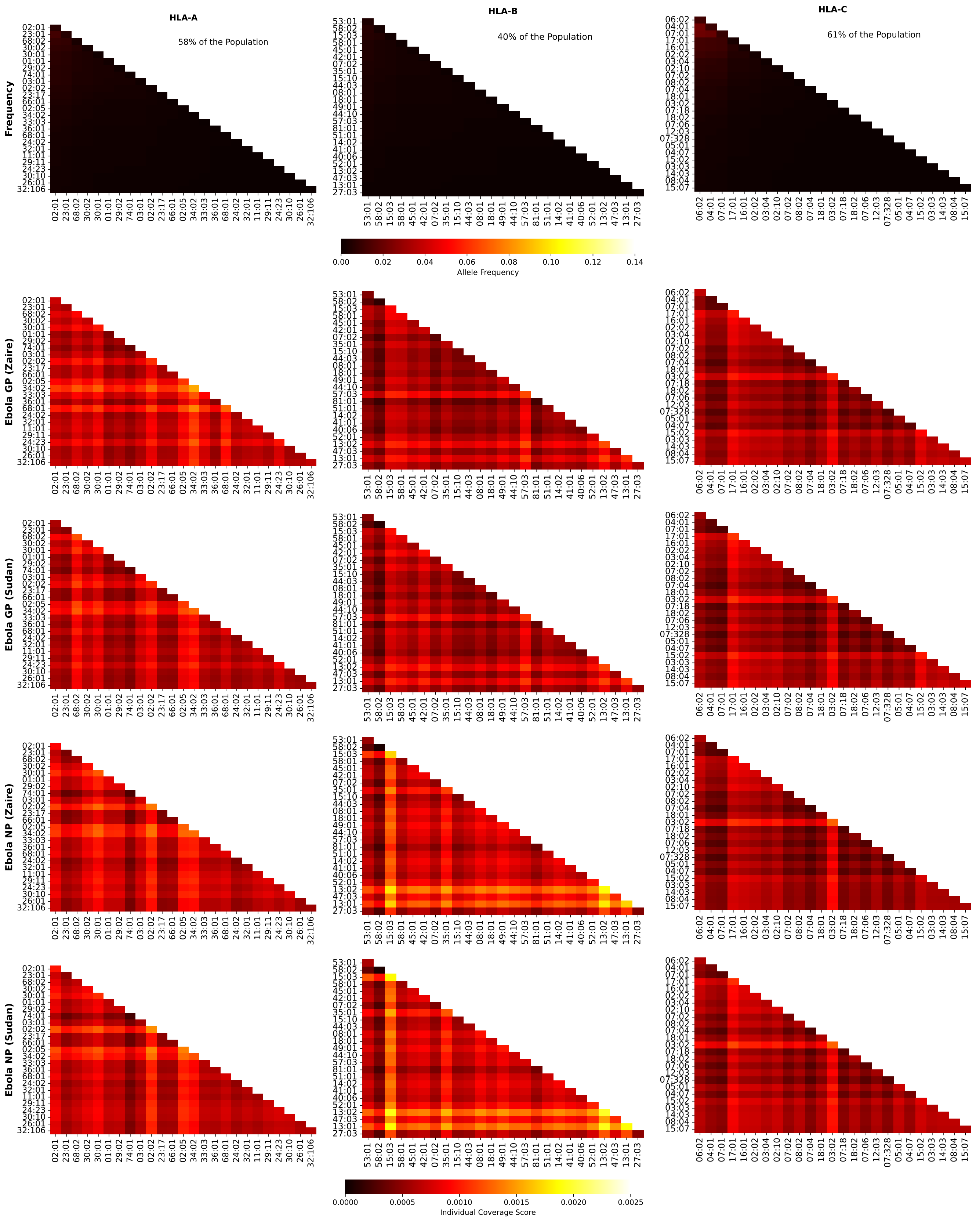


**Figure S80.** Frequencies and Ebola coverage scores for individuals in Sub-Saharan Africa. The 1st row corresponds to allele frequencies, the 2nd to GP1 Zaire, the 3rd to GP1 Sudan, the 4th to NP Zaire, and the 5th to NP Sudan. The 1st column is associated with HLA-A alleles, the 2nd to HLA-B, and the 3rd to HLA-C. The sum of the individual frequencies for each allele type is indicated on the panels in the 1st row.

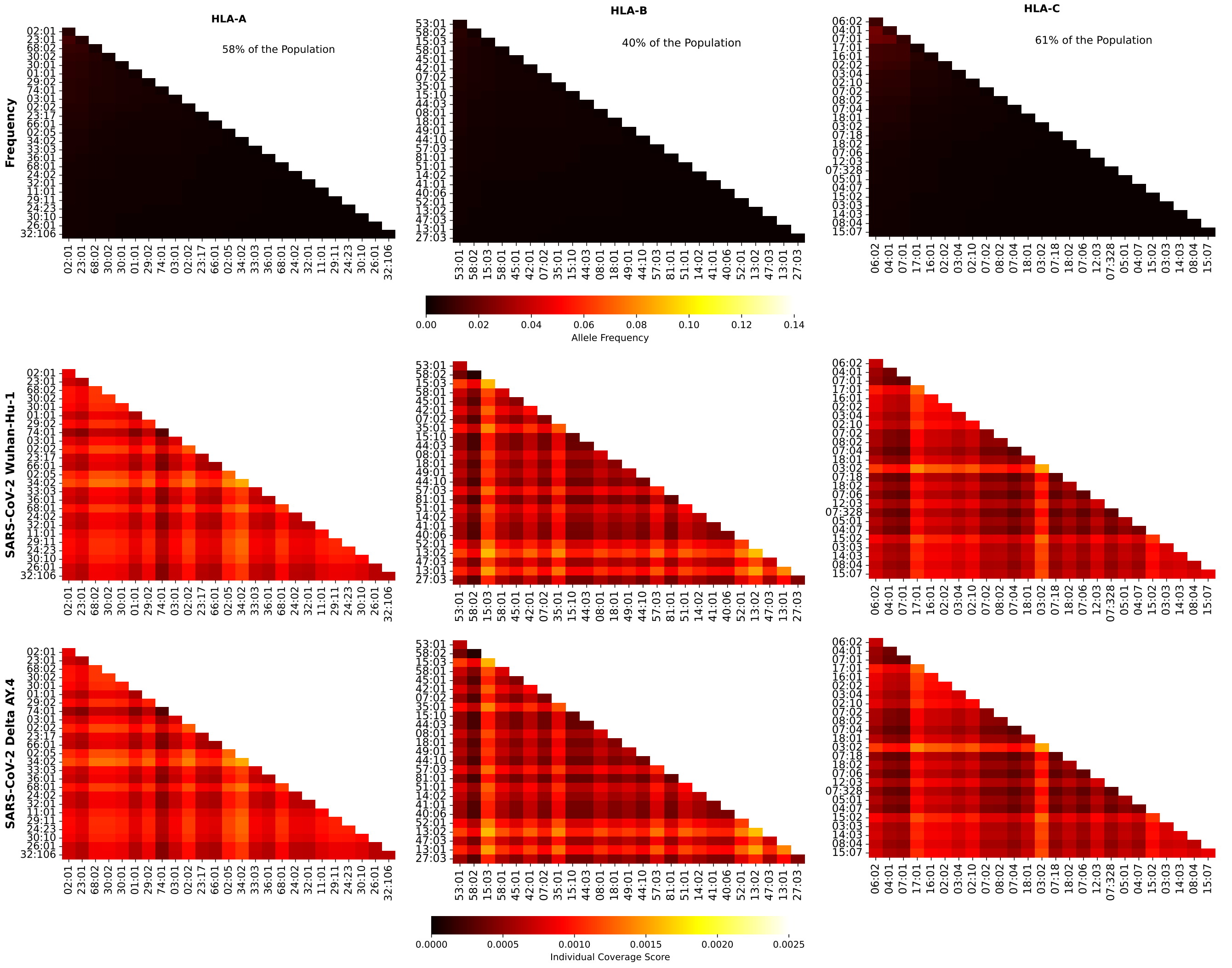


**Figure S81.** Frequencies and SARS-CoV-2 (Wuhan-Hu-1 and Delta AY.4 variants) coverage scores for individuals in Sub-Saharan Africa. The 1st row corresponds to allele frequencies, the 2nd to Wuhan-Hu-1, and the 3rd to Delta AY.4. The 1st column is associated with HLA-A alleles, the 2nd to HLA-B, and the 3rd to HLA-C. The sum of the individual frequencies for each allele type is indicated on the panels in the 1st row.

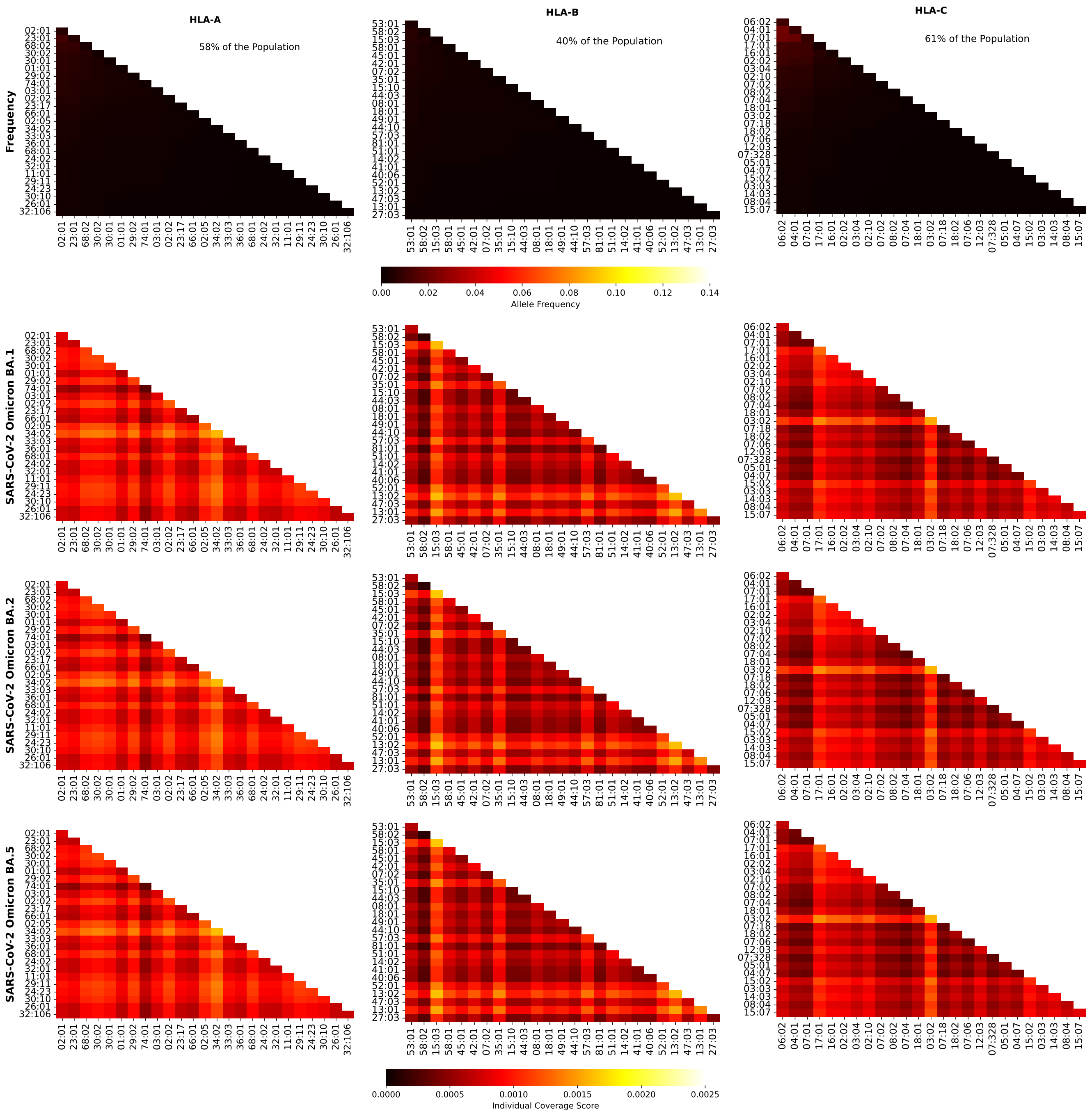


**Figure S82.** Frequencies and SARS-CoV-2 (Omicron variants) coverage scores for individuals in Sub-Saharan Africa. The 1st row corresponds to allele frequencies, the 2nd to BA.1, and the 3rd to BA.2, and the 4th to BA.5. The 1st column is associated with HLA-A alleles, the 2nd to HLA-B, and the 3rd to HLA-C. The sum of the individual frequencies for each allele type is indicated on the panels in the 1st row.

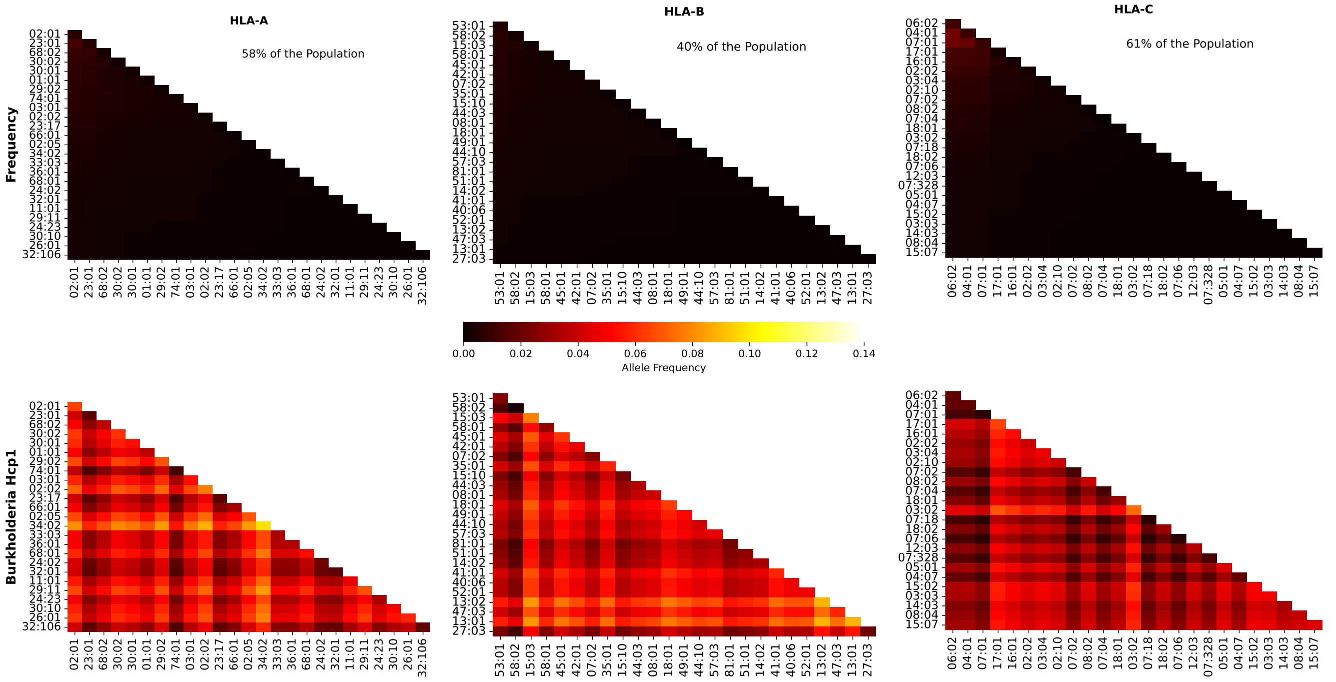


**Figure S83.** Frequencies and Burkholderia coverage scores for individuals in Sub-Saharan Africa. The 1st row corresponds to allele frequencies and the 2nd row to Burkholderia coverage score. The 1st column is associated with HLA-A alleles, the 2nd to HLA-B, and the 3rd to HLA-C. The sum of the individual frequencies for each allele type is indicated on the panels in the 1st row.

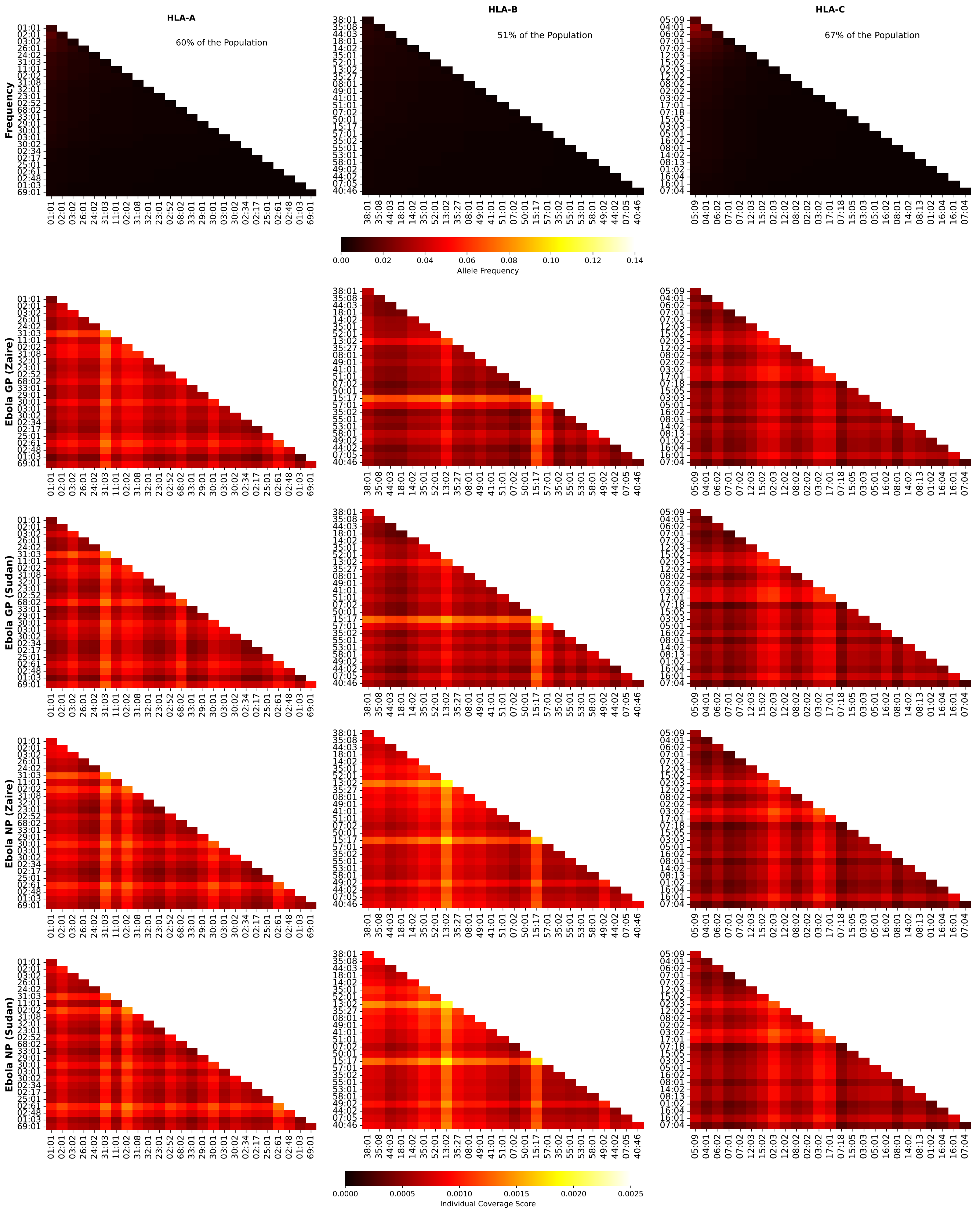


**Figure S84.** Frequencies and Ebola coverage scores for individuals in Western Asia. The 1st row corresponds to allele frequencies, the 2nd to GP1 Zaire, the 3rd to GP1 Sudan, the 4th to NP Zaire, and the 5th to NP Sudan. The 1st column is associated with HLA-A alleles, the 2nd to HLA-B, and the 3rd to HLA-C. The sum of the individual frequencies for each allele type is indicated on the panels in the 1st row.

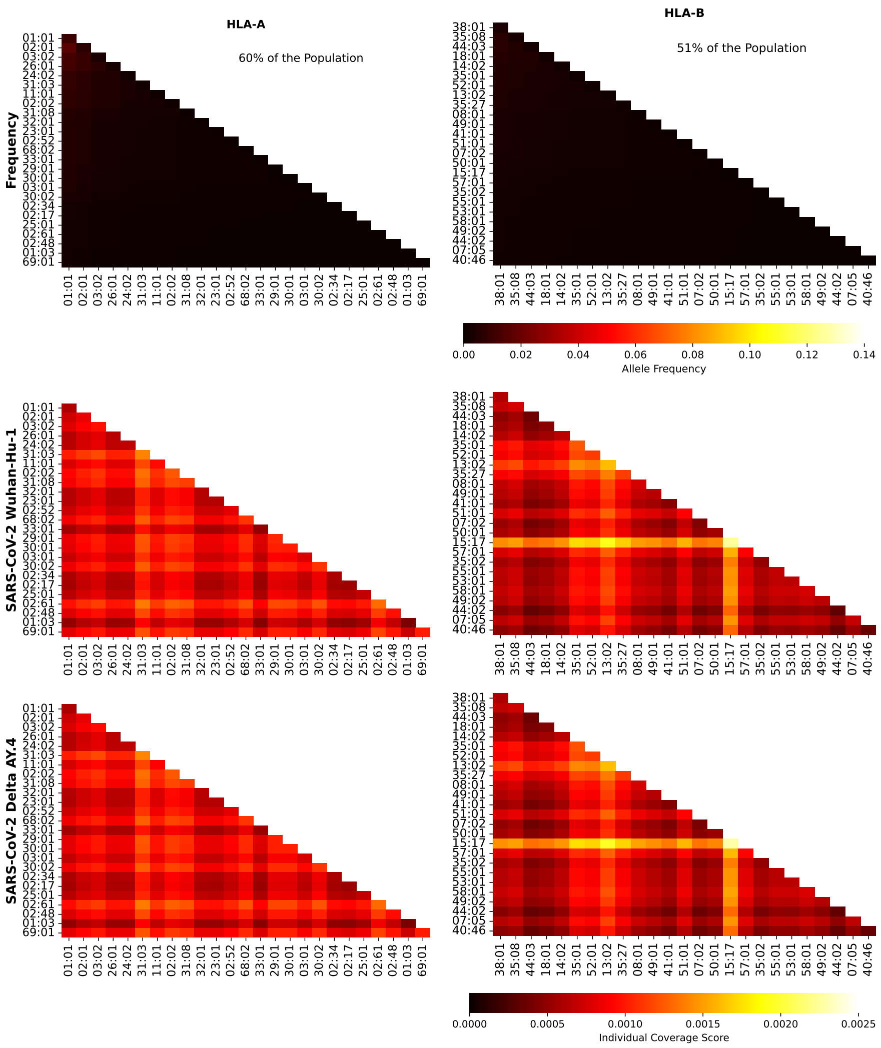


**Figure S85.** Frequencies and SARS-CoV-2 (Wuhan-Hu-1 and Delta AY.4 variants) coverage scores for individuals in Western Asia. The 1st row corresponds to allele frequencies, the 2nd to Wuhan-Hu-1, and the 3rd to Delta AY.4. The 1st column is associated with HLA-A alleles, the 2nd to HLA-B, and the 3rd to HLA-C. The sum of the individual frequencies for each allele type is indicated on the panels in the 1st row.

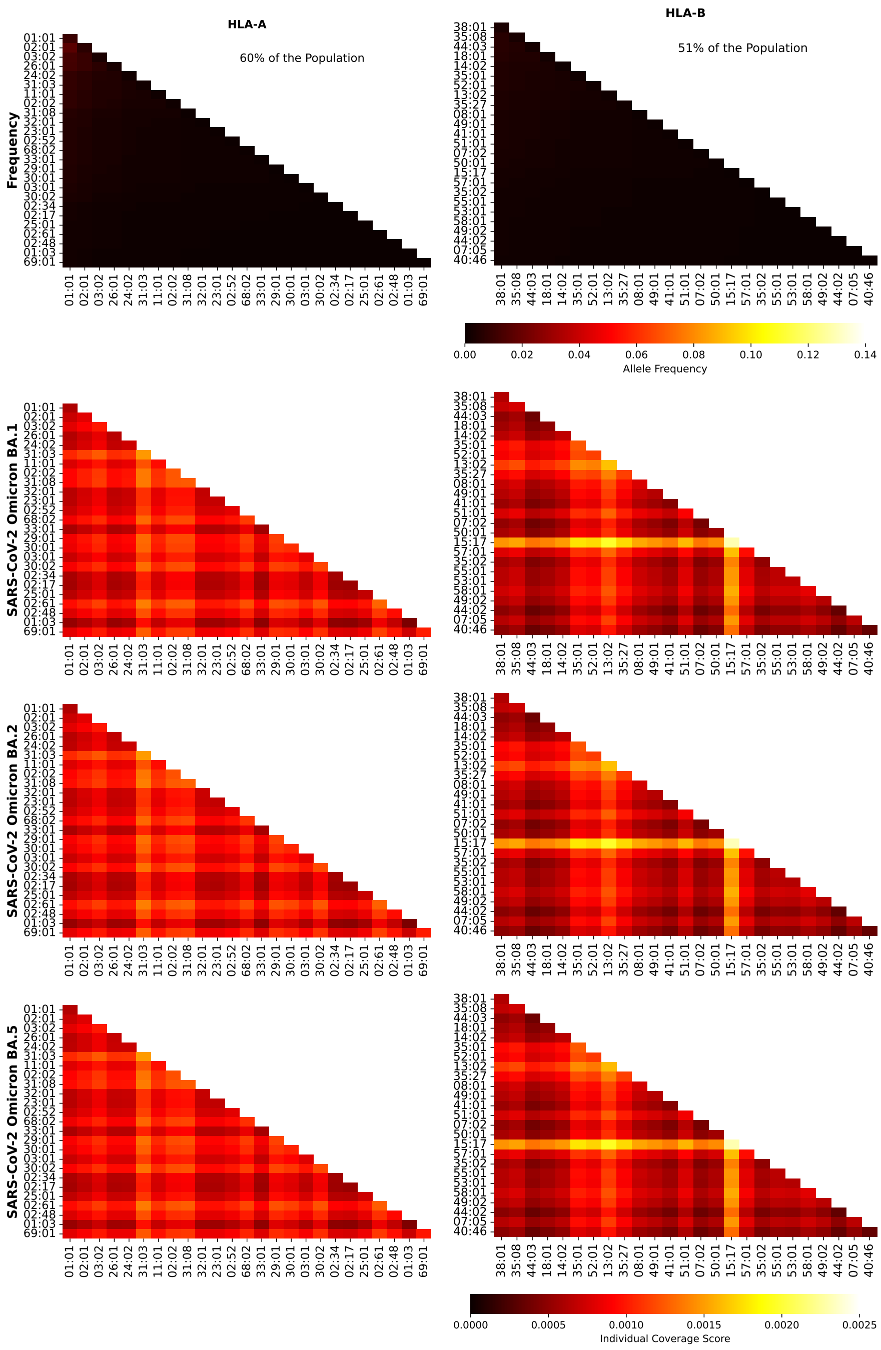


**Figure S86.** Frequencies and SARS-CoV-2 (Omicron variants) coverage scores for individuals in Western Asia. The 1st row corresponds to allele frequencies, the 2nd to BA.1, and the 3rd to BA.2, and the 4th to BA.5. The 1st column is associated with HLA-A alleles, the 2nd to HLA-B, and the 3rd to HLA-C. The sum of the individual frequencies for each allele type is indicated on the panels in the 1st row.

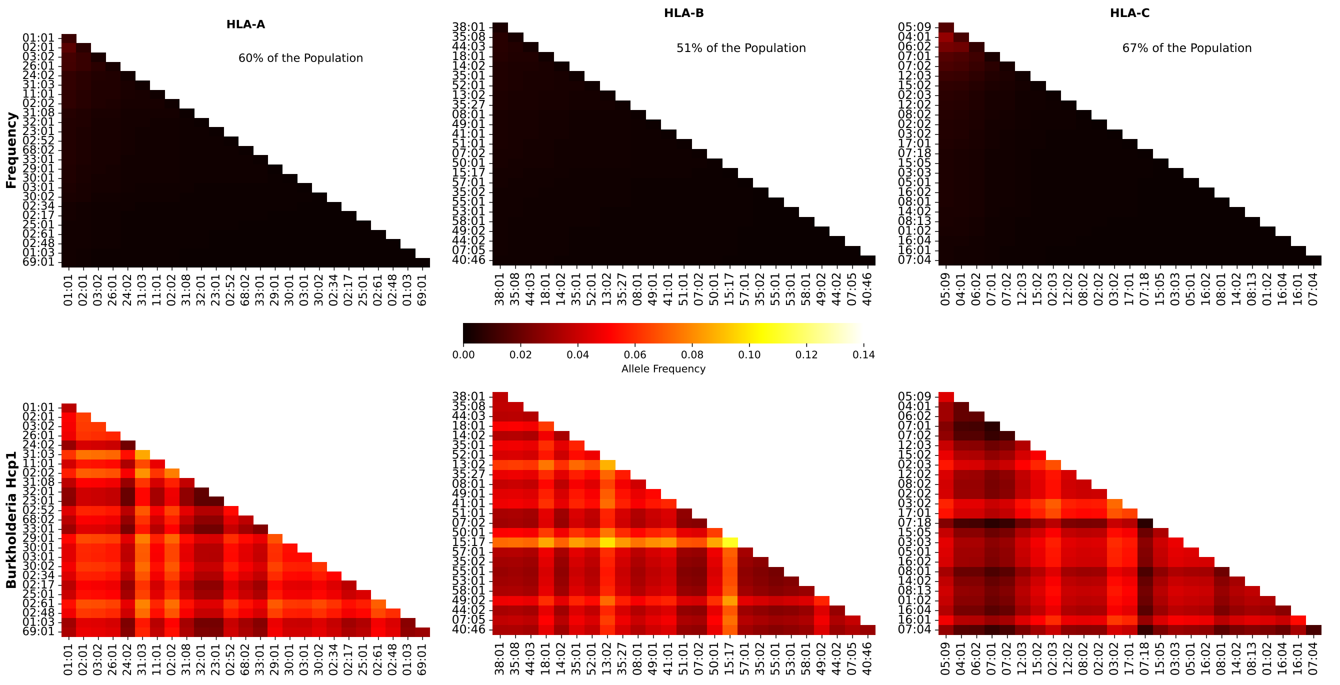


**Figure S87.** Frequencies and Burkholderia coverage scores for individuals in Western Asia. The 1st row corresponds to allele frequencies and the 2nd row to Burkholderia coverage score. The 1st column is associated with HLA-A alleles, the 2nd to HLA-B, and the 3rd to HLA-C. The sum of the individual frequencies for each allele type is indicated on the panels in the 1st row.
